# EDTP Loss of Function Impairs Longevity and Reproduction

**DOI:** 10.64898/2026.08.31.748413

**Authors:** Xuedou Lu, Taylor Barwell, Scott Edelman, Laurent Seroude

**Affiliations:** Department of Biology, Queen’s University, Kingston, Ontario, Canada

## Abstract

This study presents a comprehensive genetic characterization of *DJ694*, a viable GAL4 enhancer-trap allele of the age-regulated gene *EDTP*. *EDTP* transcript levels are reduced in DJ694, and homozygous flies exhibit reduced reproduction and shortened lifespan in both sexes. The fertility and longevity phenotypes are recessive and can be fully rescued by three independent UAS*-EDTP* insertions. Expression of UAS*-EDTP* into animals with wildtype phenotypes does not affect female fertility or lifespan. *DJ694* has a jumpy behaviour but it is not detected by a locomotion assay. Like its human homolog, Muscle-specific Inositol Phosphatase (*MIP*, also known as *MTMR14*), *EDTP* is mainly expressed in adult muscles. The structure of the muscle, myofibril, and sarcomere appears normal in homozygous *DJ694*. *DJ694* females have morphologically normal ovaries, implicating a functional rather than structural impairment. We demonstrate that *EDTP* is required during both development and adulthood to support normal fertility, with expression restricted to either stage alone being insufficient. Although it has been reported that *EDTP* can influence the accumulation of polyglutamine aggregates, we did not find evidence in its native tissue. Ectopic expression in the eye reduces the amount of aggregates but *EDTP* overexpression in muscles has no detectable effect on the accumulation or toxicity of two different kinds of polyglutamine aggregates.

## Introduction

*EDTP* (Egg-derived Tyrosine Phosphatase) was first identified from flesh fly eggs as a substrate of the cathepsin L protease with tyrosine phosphatase activity (Yamaguchi et al., 1999). The EDTP protein belongs to the Myotubularin and Myotubularin-related protein superfamily (Alonso et al., 2004; Tosch et al., 2006). Although Myotubularin has tyrosine phosphatase activity, it turned out to be a lipid phosphate with much higher specific activity toward phosphatidylinositol 3-phosphate (PI(3)P) than protein substrates (Taylor et al., 2000). The mammalian homolog of EDTP, MTMR14, also known as hJumpy, dephosphorylates PI(3)P and phosplatidylinositol-3,5-phosphate (PI(3,5)P_2_) at the D3 position instead of protein substrates (Alonso et al., 2004; Shen et al., 2009; Tosch et al., 2006).

In *Drosophila*, *EDTP* is expressed during oogenesis and embryogenesis (Yamaguchi et al., 2005). During oogenesis, *EDTP* mRNA is detected from stage 5 onward, predominantly in the cytoplasm of the nurse cells; by stage 10, it is also present in the oocyte cytoplasm (Yamaguchi et al., 2005). During embryogenesis, *EDTP* mRNA is present from freshly laid eggs through late embryonic stages (Yamaguchi et al., 2005). The mRNA and the protein are also expressed in the larval fat body (Manzéger et al., 2021; Papp et al., 2016). *EDTP* null allele are embryonic lethal, and germline clones exhibit severe defects in ovarian development (Yamaguchi et al., 2005). The *DJ694* enhancer-trap GAL4 strain has been isolated in a screen for changes in gene expression with age in adults (Seroude et al., 2002). The characterization of the expression profile of GAL4 with β-galactosidase reporters showed that it is localized in muscles and the level of expression increase from 0 to 30 days old. During development, GAL4 is expressed in embryos but not in the larvae (Barwell et al., 2017). The molecular mapping of the insertion showed that it is localized in the first intron of the *EDTP* gene (Seroude et al., 2002). The GAL4 expression has been independently confirmed with GFP reporter and the quantification of *EDTP* transcripts by real time PCR of 3 days- and 30 days-old adults demonstrated that it reflects *EDTP* expression (Singh et al., 2014). Since the *DJ694* insertion is homozygous viable, it provides a new *EDTP* allele that facilitate the determination of the role of *EDTP* in adult flies. In human, northern blot analyses demonstrate that *MTMR14* is mainly expressed in heart and skeletal muscle (Shen et al., 2009; Tosch et al., 2006) leading to renaming it *MIP* (Muscle-specific Inositol Phosphatase). *MTMR14* and myotubularin have been linked to myotubular myopathies, which are disorders characterized by progressive muscle weakness and wasting (Kim et al., 2002; Laporte et al., 2000; Laporte et al., 1996; Tosch et al., 2006). *MIP* knockout mice exhibit disrupted calcium homeostasis that affect muscle performance as well as muscle wasting (Dowling et al., 2010; Romero-Suarez et al., 2010; Shen et al., 2009; Wen et al., 2018). Mutations that cause muscle degeneration in *Drosophila* displays elevated expression of *EDTP*, suggesting that *EDTP* might serve a protective role in the muscle (Singh et al., 2014).

Mechanistically *EDTP/MTMR14/MIP* has been reported to be involved in autophagy (Vergne et al., 2009). Autophagy begins when the ULK1 initiation complex activates and recruits Atg9 and the PI3K III nucleation complex (Yamamoto et al., 2023). Atg9 is a lipid scramblase while the PI3K III nucleation complex produces PI(3)P in the endoplasmic reticulum (ER) membrane which can be bound by WIPI2. Two ubiquitin-like conjugation cascades are involved (Fleming et al., 2022). One cascade uses the E1-like enzyme Atg7 and the E2-like enzyme Atg10 to conjugate the ubiquitin-like Atg12 with its Atg5 substrate before recruiting the membrane/WIPI2 binding protein Atg16. The second cascade starts with Atg4 cleaving the ubiquitin-like Atg8/LC3 that is subsequently conjugated to phosphatidylethanolamine (PE) in the ER membrane by the E1-like Atg7, the E2-like Atg3 and the E3-like Atg5-Atg12-Atg16 complex. The conjugation to PE leads to the formation of the phagophore that will elongate and close to form the autophagosome that will subsequently fuse with the lysosome for degradation. Cargos are captured either non-selectively in bulk, or selectively via cargo receptors such as p62/ref(2)P or BNIP3. Because PI(3)P is a substrate of MTMR14, the involvement of MTMR14 in autophagy has been investigated. MTMR14 influences both early and late stages of autophagy. *MTMR14* knockdown in murine myoblasts and macrophages led to increased LC3-II levels and acceleration of p62 degradation, indicating stimulated autophagosome formation and increased autophagic flux (Vergne et al., 2009). Reciprocally, *MTMR14* overexpression led to p62 accumulation, suggesting that it blocks autophagic degradation. Deletion of *MTMR14* in mouse embryonic fibroblasts led to elevated LC3-II levels and increased autophagy-lysosome granules (Liu et al., 2014). *MTMR14* knockout mice also show elevated LC3-II in muscle (Hnia et al., 2012). In zebrafish, *MTMR14* morphants also exhibit significantly elevated LC3-II levels (Dowling et al., 2010).

Inhibitors of MTMR14 phosphatase activity causes an activation of autophagy in human cells and *Drosophila* larvae (Kovács et al., 2017; Papp et al., 2016). The effect of the inhibitors in the *Drosophila* larval fat is abolished by *EDTP* loss of function confirming that *EDTP* is the functional homologue of *MTMR14* in *Drosophila*. The genetic analysis of the role of *EDTP* in the larval fat body provided the best experimental evidence that *EDTP* is a negative regulator of autophagy in vivo in an intact organism. By inducing somatic clones in *Drosophila*, chimeric animals can be obtained in which GFP positive cells express higher (over-expression) or lower (RNAi) amount of *EDTP* surrounded by genetically identical wild-type control cells that are GFP negative. In RNAi clonal cells Atg8, Atg18, autolysosomes, and PI(3)P are elevated, while overexpression clones have reduced Atg8 levels (Manzéger et al., 2021; Papp et al., 2016). Loss-of-function mutant also showed increased Atg8 and autolysosomes, as well as decreased level of p62. Furthermore, Atg13 levels are unaffected by *EDTP* loss of function indicating that *EDTP* acts downstream of the induction complex (Manzéger et al., 2021; Papp et al., 2016). Unfortunately, the biological significance of the regulation of autophagy by EDTP in the fat body during the larval stage remains unknown since no autophagy is detected in wild-type larvae (Papp et al., 2016). Furthermore, the activation of autophagy in *EDTP* loss of function animals does not cause any visible phenotypes and only results in pupal lethality with low penetrance (Manzéger et al., 2021). The biological role of EDTP in adult muscles have not been reported. The flying ability of a loss of function allele is impaired (Komlos et al., 2023) but this allele also causes lethality during embryogenesis with 39% penetrance (Manzéger et al., 2021), and it cannot be excluded that the impairment is due to its role during development.

Here, we performed a genetic analysis of *DJ694*, an enhancer-trap allele of *EDTP*. We show that *DJ694* has reduced *EDTP* transcript levels and examined developmental lethality, reproduction, longevity, muscle integrity, and locomotion. *DJ694* mutants display decreased fertility and shortened longevity in both males and females, and both defects are rescued by several UAS-*EDTP* constructs. To investigate the basis of the reduced fertility phenotype, we examined ovarian morphology and determined at what stage of the life cycle EDTP is acting. Although no visible morphological defects are seen in *DJ694* females, EDTP is required during both developmental and adult stages to support normal fertility. Finally, we investigated if the regulation of autophagy by EDTP in adult muscles has biological consequences. A major outcome of autophagy is to cope with protein aggregates. Ubiquitin-positive aggregates accumulate naturally during aging in *Drosophila* muscles. (Demontis & Perrimon, 2010). The activation of autophagy reduces the amount of ubiquitin-positive muscle aggregates (Komlos et al., 2023) as well as polyglutamine aggregates in the nervous system (Billes et al., 2016). We recently demonstrated that the expression of polyQ aggregates in *Drosophila* muscles reduces longevity in a dose-dependent manner (Taylor Barwell et al., 2023). Furthermore, the level of expression during metamorphosis can lead to a drastically different kind of aggregate distribution that causes a much more severe reduction of longevity. We tested if EDTP affects the reduction of longevity by polyQ aggregates and if it affects the abundance of two kind of aggregates. We found no evidence that EDTP affect the shortened lifespan, nor did it alter the aggregates.

## Materials and Methods

### Fly Cultures and Strains

Flies were maintained on standard cornmeal medium (0.01% molasses, 8.2% cornmeal, 3.4% yeast, 0.94% agar, 0.18% benzoic acid, 0.66% propionic acid) at 25°C unless stated otherwise. *w^1118^* was used as a wildtype. The original *DJ694 w^1118^;P{w^+mW.hs^=GAWB}EDTP^DJ694^*strain was generated and described previously (Seroude et al., 2002). This strain was subsequently outcrossed and isogenized with a Canton-S background to generate *DJ694^P^*. Heterozygous *DJ694^O^* (BDSC: 8176) has been independently maintained by the Bloomington Drosophila Stock Center since 2002 and was reobtained in 2008. The homozygous *DJ694^H^*stock was generated from *DJ694^O^*. *DJ694^H^* was outcrossed and isogenized with a Canton-S background to generate *DJ694^L^*. Using multiple independently maintained lines of the same genotype reduces the risk that phenotypes reflect genetic drift or background effects accumulated within a single lineage rather than the mutation of interest, consistent with recommendations for rigorous aging studies (Partridge & Gems, 2007). The *GAL80^ts^ w^+^;P{w^+mc^=tubP-GAL80^ts^}2/TM2* (BDSC: 7017) and the *GMR w^+^;P{w^+mc^=GAL4-ninaE.GMR}12* (BDSC: 1104) strains were obtained from the Bloomington Drosophila Stock Center. The *DJ694;GAL80^ts^*strain was generated with standard genetic crosses. *UAS-Httex1-Q72-eGFP (Q72)* (Zhang et al., 2010) was obtained from N. Perrimon’s lab. The *Q72;;UAS-lacZ*, *Q72;;Mef2* and *Q72;;MHC* have been previously described (Taylor Barwell et al., 2023).

### UAS-EDTP Cloning

The cDNA clone LD39930 was obtained from the Drosophila Genomics Resource Centre (DGRC stock 1326332; https://dgrc.bio.indiana.edu//stock/1326332; RRID:DGRC 1326332). The cDNA was excised with EcoRI and XhoI and subcloned into the pBluescript SK+ vector to generate the pDJ134 plasmid. pDJ135 was constructed by subcloning the 2277bp BglII–SacII fragment (cDNA without 3’ UTR and coding region with 3’ deletion) from pDJ134 into pINDY5 (Xiao et al., 2007) digested with the same enzymes. pDJ134 was used as a PCR template with primers DJ105 (5’-CCTGCTGGTCAACGGATTGAC-3’) and DJ110 (5’-AGATCTAAACAGTTGGTCCGCCAGG-3’) to produce a 511 bp fragment containing the deleted coding region flanked by BglII sites. The PCR fragment and pDJ135 were digested with BglII and ligated to obtain the pDJ137 plasmid in which the EDTP open reading frame is under the control of a UAS promoter.

### Generation of Transgenic Animals

*UAS-EDTP* lines were generated by P-element transformation (Spradling & Rubin, 1982), without the removal of the chorion and dessication (Robertson et al., 1988). *w^1118^* embryo were injected with a mixture of pDJ137 plasmid (4.1 μg/μl) and plasmid pπ25.7 (0.584 μg/μl) (Karess & Rubin, 1984) in 1 mM sodium phosphate buffer at pH 7. Four independent insertions were obtained. All located on the second chromosome. Transposition was used to generate homozygous viable insertions located on the third chromosome. The strain with the corresponding insertions are: *w^1118^;; UAS-EDTP^A^*, *w^1118^;; UAS-EDTP^B^*, *w^1118^;; UAS-EDTP^D^*, *w^1118^;; UAS-EDTP^E^*. *DJ694;UAS-EDTP, Q72;;UAS-EDTP^D^, Q72;;UAS-EDTP^E^* strains were generated with standard genetic crosses.

### RNA Extraction and Semiquantitative RT-PCR

Total RNA was extracted from 60 freshly eclosed male flies per genotype (*w¹¹¹⁸* and *DJ694*) at 0– 1 day of age using the RNAqueous Kit (Ambion, Cat. No. AM1912) following the manufacturer’s instructions, with two independent biological replicates per genotype. Genomic DNA was removed using the DNA-free Kit (Ambion, Cat. No. AM1906). RNA quality was assessed visually by electrophoresis on a 0.7% TAE agarose gel, and RNA’s concentration was quantified by spectrophotometry at 260 nm. The removal of genomic DNA was verified by PCR using *EDTP* primers targeting exon 2 (DJ99: 5’-CAATATCCCTCGCAGCAGATG-3’) and exon 6 (DJ108: 5’-CAATGTTGATGAAATCTAAC-3’). The presence of genomic DNA yields a 2109bp product whereas the amplification of reverse-transcribed mRNA yields a 1836bp product. RT-PCR was performed using Ready-to-Go RT-PCR Beads (Amersham, cat# 27-9267-01) using 250ng (undiluted) or 25ng (10 fold dilution) RNA. Each reaction was supplemented with 1.2µM oligo d(T) (12-18nt), 400nM of each primer and RT-PCR Grade water (Ambion, cat# 9935). The reverse transcription was done at 42°C for 30 minutes. After an initial denaturation at 95°C for 5 minutes, 35 cycles PCR was performed in a Biometra thermocycler as follow: denaturation at 95°C for 30 seconds, annealing at 55°C for 1 minute, and extension at 72°C for 3 minutes. A final extension at 72°C for 10 minutes was performed after the last cycle.

### Development Lethality

To assess developmental lethality, crosses were established at 25°C using 40–50 males and 80– 120 females. Parental flies were transferred to egg collectors 24–48 hours after crosses were set up, and a drop of yeast was added to the egg collector to stimulate egg-laying. Females were allowed to lay eggs for 12-16 hours. Survival was assessed at three successive developmental transitions: egg to L1 larvae, L1 larvae to pupae, and pupae to adults. Survival rates at each stage were compared between genotypes using a two-sample, two-tailed, equal-variance t-test. The dataset and statistics are provided in Table S1.

### Reproduction assays

Female fertility was determined by transferring to a new vial daily and counting the number of eggs laid at 25°C for the entire lifespan for initial experiments, and for 40 days for the subsequent experiments. This assay used either 3 vials of 5 females and 5 males, or 3-8 vials of 3 females and 3 males. Female fertility was calculated daily by diving the number of eggs by the number of females remaining per vial. The cumulative number of eggs was obtained by adding the daily values, and the average of the vials is reported. After egg counting, vials were then stored for 10–12 days at 25°C, after which the number of eclosed adult progeny was scored to assess reproduction success and male fertility. Genotypes were compared using two-sample, two-tailed, equal-variance t-tests. Statistical outcomes from each independent biological replicate are represented by symbols on the graphs. A triangle indicates a statistically significant difference in egg output between the compared genotypes (p < 0.05), while a square indicates no significant difference (p ≥ 0.05). The direction of the triangle indicates whether egg output is higher or lower in the experimental genotype relative to the control, and symbol colour denotes the specific comparison being made, as detailed in the individual figure captions. A summary of the data and statistical results for each genotype comparison are provided in Table S2-4, and the corresponding daily fertility and eclosion data for each vial and replicate are provided in Table S5 and Table S6. A summary of mean egg output, standard deviation, and statistical outcomes, as well as daily fertility data, for the experiments expressing *EDTP* in a wild-type background are provided in Tables S7 and S8 respectively.

### Longevity assay

Longevity was assessed separately in males and females. Crosses were monitored daily, and upon observing the first few emerging adults, the bottle was emptied. Adults were collected and separated by sex less than 48h later. Flies were maintained at 25°C with a density between 20 to 35 per vial, with a minimum of two vials per genotype per sex. Flies were transferred to fresh food every 3 to 4 days, at which point deaths were recorded. Vials showing abnormally early mass mortality were excluded from analysis, as this is indicative of effect from experimental conditions rather than a genuine biological effect. The reduction in longevity was determined by calculating across vials the difference between the mean lifespan of the control minus its standard deviation and the mean lifespan of homozygous *DJ694* plus its standard deviation. Only 2 measurements in males and 3 in females do not show any change from the heterozygous control which is due to higher variability between vials. Initial experiments used *w¹¹¹⁸* as the control. Once the phenotype was confirmed to be recessive, heterozygous *DJ694/+* flies were used as the control for subsequent phenotype and rescue experiments. A successful rescue was defined as a significant improvement in lifespan relative to homozygous *DJ694*, with no significant difference from the heterozygous control. Log-rank tests were used to determine whether survival curves differed significantly between genotypes. All log-rank tests were performed with R Survival package. Statistical outcomes from each independent biological replicate are represented by symbols on the graphs, where each symbol corresponds to a single replicate. Filled symbols indicate p < 0.05 and unfilled symbols indicate p ≥ 0.05. A significant p-value from the log-rank test alone is not sufficient to conclude a meaningful change in lifespan, as the log-rank test is sensitive to differences in the shape of survival curves and can reach significance even when the difference in mean lifespan between genotypes is small. Symbol shape therefore reflects the relationship between the mean ± standard deviation ranges of the compared genotypes: a downward triangle indicates that the mutant mean plus its standard deviation is lower than the control’s mean minus its standard deviation, and a square indicates overlapping ranges. A filled square therefore indicates that although the survival curves differ significantly, the difference in mean lifespan falls within the natural variability of the data and is not considered biologically meaningful. Symbol colour denotes the specific comparison being made, as detailed in the individual figure captions. Summaries of mean lifespan, standard deviation, and statistical outcomes for each genotype and replicate are provided in Table S9 for the phenotype and rescue experiments, Table S10 for the effect of *EDTP* expression in a wild-type background, and Table S11 for the effect of *EDTP* on polyglutamine-mediated reduction of longevity. Flies expressing polyglutamine aggregates and EDTP with the MHC-Gene Switch driver were treated with 50µg/ml RU486 that we previously reported to yield the highest expression level (Barwell et al., 2023; Poirier et al., 2008). Crosses used to generate genotypes for longevity assays are indicates in Table S12.

### Locomotion

Locomotion assays were performed as described (Barwell et al., 2021, 2022). Females and males of both genotypes were recorded at 2, 4, 8, 15, 22, 30, 36, and 43 days of age. Three locomotion parameters were quantified: percentage of time spent immobile (%), total distance travelled (mm), and mean velocity (mm/s). At each age point, each *DJ694* background was compared to w^1118^ using a two-sample, two-tailed, equal-variance t-test. The dataset and statistics are provided in Table S13&S14.

### Muscle Sections for Light and Electron Microscopy

Adult tissue samples were processed for light microscopy following the protocol of (Renfranz & Benzer, 1989). Serial semi-thin plastic sections (1 µm) were stained with a solution of 1% toluidine blue and 1% borax in water. For electron microscopy, dorsal indirect flight muscles dissected from 20- and 40-day-old adult flies were fixed overnight at 4°C in a mixture of 2% paraformaldehyde and 1% glutaraldehyde, then post-fixed in 1% osmium tetroxide (OsO_4_) at room temperature. Samples were subsequently dehydrated through a graded ethanol series and embedded in Epon 812. Ultrathin sections of 80 nm were cut and examined at 100 kV using a Philips 420 electron microscope (Philips, Eindhoven, The Netherlands). A minimum of five individuals per genotype were analyzed at each time point.

### Ovary Dissection

Ovaries were dissected from 5-day-old mated females. Virgin females were housed with males in vials supplemented with yeast, and mating success was confirmed by the presence of larvae in the vials immediately prior to dissection. Four *DJ694* backgrounds (*DJ694^H^, DJ694^L^, DJ694^O^,* and *DJ694^P^*) were tested alongside *w¹¹¹⁸* controls. Two experimental conditions were used: *DJ694* females mated with *w¹¹¹⁸* males (2 replicates) and *DJ694* females mated with homozygous *DJ694* males (4 replicates), with *w¹¹¹⁸* females mated with *w¹¹¹⁸* males serving as the control in both cases. Ovaries were dissected in 1x PBS at room temperature and immediately mounted on glass slides with a drop of 1x PBS and covered with a coverslip for imaging. At least 6 females were dissected per genotype per replicate. Images were acquired using a Zeiss Axioskop 2 microscope at 50x magnification equipped with a Levenhuk M800 Plus camera and the LevenhukLite imaging software. Ovary size was determined by measuring the surface area with Fiji (Schindelin et al., 2012). Ovary size was compared using a two-sample, two-tailed, equal-variance t-test. The dataset and statistics are provided in Table S15.

### EDTP’s temporal rescue fertility assay

To determine at which stage of the life cycle *EDTP* expression is required for normal adult female fertility, a temperature sensitive GAL80^ts^ system was used to control the timing of *EDTP* rescue. Three genotypes were measured: *DJ694;GAL80^ts^/+* (homozygous *DJ694* control, generated by crossing ♂*DJ694* with ♀*DJ694;GAL80^ts^*), *DJ694/+* (heterozygous wild-type control, generated by crossing ♂*DJ694* with ♀*w^1118^*), and *DJ694;GAL80^ts^/UAS-EDTP* (experiment group, generated by crossing ♂*DJ694;UAS-EDTP* with ♀*DJ694;GAL80^ts^)*. All parental crosses were initially established at 18°C. On day 4, parents were transferred to a fresh bottle at 18°C to generate a second replicate for the 18°C developmental conditions. On day 8, parents were transferred to 30°C for one intermediate day to allow GAL80^ts^ to inactivate, then transferred to a fresh bottle at 30°C on day 9 where eggs were laid, and larvae developed at 30°C. On day 11, parents were transferred to another fresh bottle at 30°C to generate a second replicate for the 30°C developmental conditions. This design ensured that all four temperature conditions within a replicate set were derived from the same parental cross, minimizing variation due to parental background. Upon eclosion, virgins were collected under nitrogen and placed in groups of three females with three *w¹¹¹⁸* males per vial, then shifted to their designated adult temperature. *w¹¹¹⁸* males were reared at 25°C and, collected and isolated two days prior to the assay. The same males were kept in each vial throughout the experiment. Eggs were counted under a dissecting microscope every day for 30°C adult groups and every two days for 18°C adult groups. Before counting, flies were transferred to a new vial. The number of eggs laid were recorded for 60 days for 18°C to 18°C, 40 days for 30°C to 18°C, 25 days for 18°C to 30°C, and 20 days for 30°C to 30°C. Four independent biological replicates were performed, each consisting of a separate parental cross generating two replicates for the 18°C developmental conditions and two replicates for the 30°C developmental conditions. Statistical comparisons were made using a two-sample, two-tailed, equal-variance t-test. A partial rescue is achieved when there is a significant difference (p < 0.05) between the rescue genotype and the homozygous control. A full rescue is achieved when there is no significant difference between the rescue genotype and the heterozygous wildtype control. Statistical outcomes from each independent biological replicate are represented by triangles and squares as described in the reproduction assay methods, where triangle direction and symbol colour denote the direction of change and the specific genotype comparison being made, as detailed in the individual figure captions. Dataset and statistics are provided in Table S16 and S17.

### Eye GFP imaging

Desired genotypes were obtained by crossing *GMR-GAL4* males with *Q72*, *Q72;;UAS-lacZ*, *Q72;;UAS-EDTP^D^*, *Q72;;UAS-EDTP^E^*females. Progeny were collected upon eclosion and aged to 7, 14, and 28 days. Animals were decapitated with a razor blade and heads were bisected sagittally so that the head lays flat. GFP fluorescence was visualized directly by epifluorescence with a Zeiss Stemi SV 11 stereomicroscope equipped with a Zeiss AxioCam HR camera and OpenLab imaging software (Improvision, Lexington, MA, USA). All images were captured under identical exposure and illumination conditions.

### Cryosectioning and Immunohistochemistry

Whole flies were embedded in Optimal Cutting Temperature compound (OCT, Tissue-Tek) and 20µm sagittal serial sections were cut with a Leica CM 1850 cryostat. Sections were fixed in 2% glutaraldehyde for 10 minutes at room temperature and washed 5 times 5min with 1x PBS, 0.2% Triton X-100. Sections were then incubated for 30 minutes at room temperature in blocking buffer (1x PBS, 0.2% Triton X-100 and 5% BSA). Primary antibody incubation was done overnight at room temperature with a commercial rabbit anti-GFP antibody (1:1000 in blocking buffer, Fisher cat# A11122). Sections were washed 5 times 5min with 1x PBS, 0.2% Triton X-100 and incubated again for 30 minutes in blocking buffer. Secondary antibody incubation was done overnight at room temperature with a commercial goat anti-rabbit antibody conjugated to Alexa Fluor 488 (1:1000 in blocking buffer, Fisher cat# A11034). Sections were washed 5 times 5min with 1x PBS, 0.2% Triton X-100 and with Fluoromount-G with DAPI (Fisher cat# 501128966). Images were acquired using a Zeiss Axioplan II imaging microscope equipped with a Leica DC500 high-resolution camera and OpenLab imaging software (Improvision, Lexington, MA, USA). All images were captured under identical exposure and illumination conditions.

## Results

### *DJ694* is an *EDTP*-deficient allele

*DJ694* is a GAL4 enhancer trap inserted in the first intron of *EDTP* (Seroude et al., 2002). *DJ694* is predominantly expressed in muscles and the expression is regulated by physiological age (Seroude et al., 2002). Because the insertion lies near another gene, *CG18467*, it was initially unclear whether the reporter reflects *EDTP* or *CG18467* transcription. Real-time PCR analysis indicates that *EDTP* transcripts are present in dissected indirect flight muscles (IFM) and increase with age, whereas *CG18467* transcripts are absent from IFM and show detectable expression only in the brain (Singh et al., 2014). These findings support the conclusion that *DJ694* reports the expression pattern of *EDTP* in muscle. Enhancer-trap insertions can both reveal expression patterns and potentially disrupt gene function. To determine whether *DJ694* affects *EDTP* transcription, we compared *EDTP* mRNA levels between homozygous *DJ694* mutants and w¹¹¹⁸ controls using quantitative RT-PCR. *EDTP* transcript levels were significantly reduced in homozygous *DJ694* (Figure 1A). Although we have not examined the effect of the insertion at the protein level, it is very likely that *DJ694* is an *EDTP*-deficient allele.

**Figure 1.**
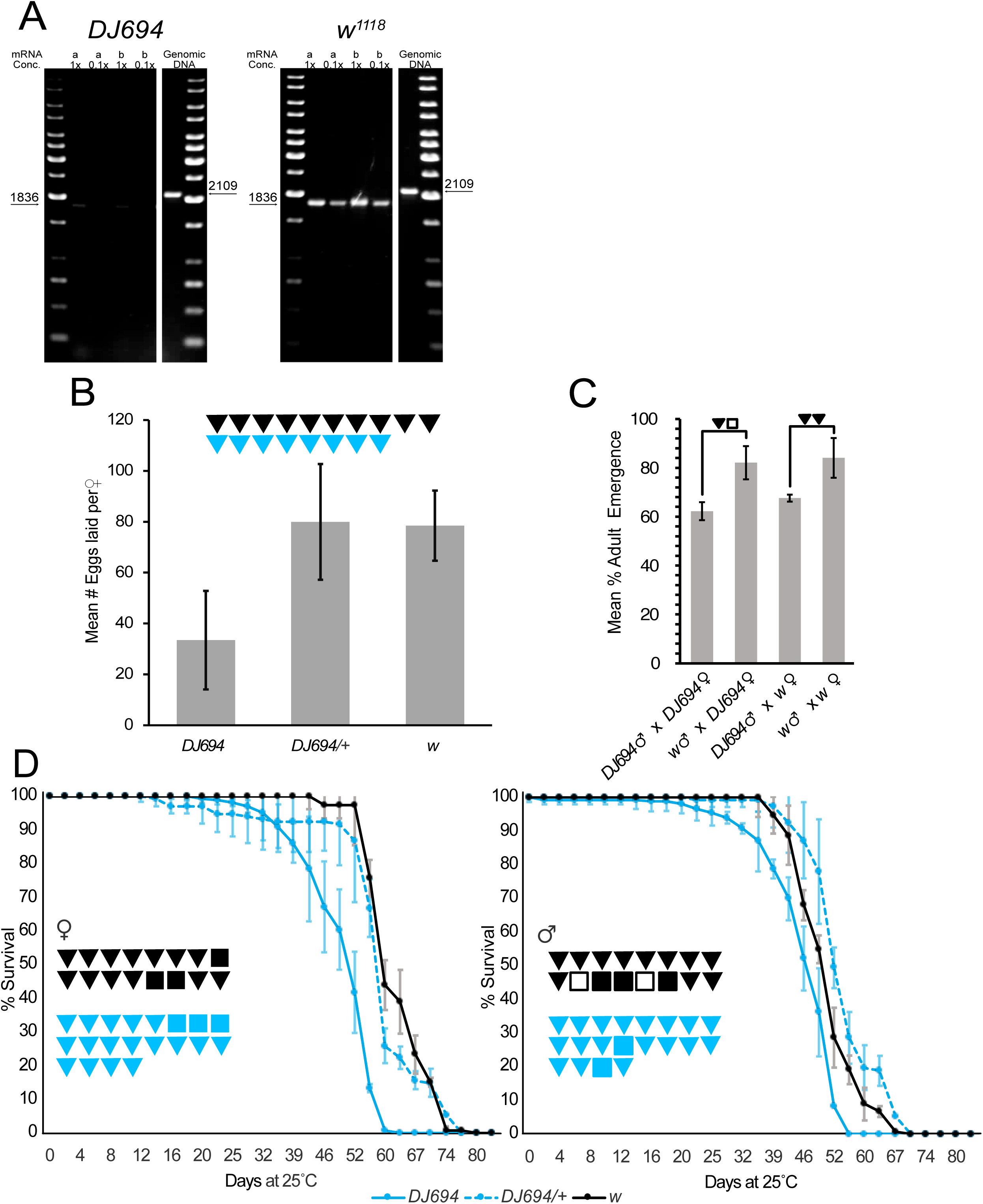
*DJ694* is *EDTP*-deficient and has impaired reproduction and longevity. A) Semi quantitative RT-PCR of *EDTP mRNA* transcripts. In each panel, the left shows the product of the RT-PCR reactions, and the right shows the PCR product that is produced if genomic DNA is present. a and b indicate the two independent RNA extractions (1x = 250 ng, 0.1x = 25 ng). Panel B, C, and D showed one representative replicate. Error bars represent standard deviation. Symbols indicate statistical outcomes from independent biological replicates. Filled symbols indicate p < 0.05 and unfilled symbols indicate p ≥ 0.05. A downward triangle indicates a decrease. A square indicates no change. For panel C and E, Black symbols indicate comparison between homozygous *DJ694* to *w* control. Blue symbols indicate comparison between homozygous *DJ694* to heterozygous *DJ694/+*. (C) Quantification of female fertility shown as the mean cumulative number of eggs laid per female per day from day 29 to day 40 at 25°C. (D) Quantification of male fertility shown as the percentage of eggs that successfully emerge as adults, calculated as the cumulative number of emerged adults divided by the cumulative number of eggs laid over 85 days at 25°C. (E) Survival curves shown separately for females and males.

### Homozygous *DJ694* flies show impaired reproduction

Since null allele is embryonic lethal (Yamaguchi et al., 2005), we first investigated whether the *DJ694* allele also exhibit developmental lethality. We found no evidence of developmental lethality in *DJ694* (Table S1). Since germline clones of the null allele show severe ovarian defects (Yamaguchi et al., 2005), we investigated *DJ694*’s reproductive functions. Fertility was assessed as the mean cumulative number of eggs produced over the complete lifespan at 25°C (85 days) by females mated with wild-type or mutant males (Figure S1A). Upon time segmentation of the data, several time windows display significant decline (Table S18). Subsequent measurements were performed until 40 days and statistical analyses were conducted between 29 and 40 days. *DJ694* females laid significantly fewer eggs than both the *w^1118^* control and the *DJ694/+* heterozygous control (Figure 1C; Table S2&S3). Since *DJ694/+* females did not differ significantly from the *w^1118^* control, the fertility defect is recessive. We also observed that *DJ694* females produces less eggs at 18°C and 29°C indicating that the reduced fertility phenotype is temperature independent (Figure S2; Table S19&S20). To confirm the phenotype is due to the insertion rather than genetics backgrounds, four DJ694 strains with different genetic backgrounds were tested with wild-type males. All four backgrounds displayed the phenotype with similar expressivity (Figure S1B; Table S2&S3) leading to the conclusion that this phenotype is indeed due to the insertion (Figure 1C). Although the reproductive output of *DJ694* females throughout the lifespan (assessed as the percentage of eggs that yielded viable adults) was not different from *w^1118^* females (Figure 1D; Table S2&S4), it is worth noticing that at oldest ages (30d and older) fewer number of adults were obtained (Figure S3). It is obvious that eggs laid by wildtype or *DJ694* female fertilized with *DJ694* male produced significantly less adults than eggs fertilized by wildtype males (Figure 1D; Table S2&S4) indicating that *DJ694* males also have reproductive defects. We scored by visual inspection for 10 minutes the mating of 50 individual wildtype or *DJ694* males with wildtype females at 3 and 28 days old. 89% of the young *DJ694* male mated, while 59% of the wildtype mated. When the same males were tested at 28 days, *DJ694* (52%) still show a higher mating rate than the wildtype (9%). *DJ694* males do not have mating impairments, implying that the reproductive defect either result from poor ejaculation or defective sperm.

### Homozygous *DJ694* flies are short lived

Next, we examined whether the insertion is affecting longevity by comparing the longevity of *DJ694* with wild-type and heterozygous *DJ694/+*. Because longevity is a complex phenotype sensitive to genetic background (Partridge & Gems, 2007), five *DJ694* backgrounds were measured (Figure 1E; Table S9). Both male and female homozygous *DJ694* flies were significantly shorter-lived than *DJ694/+* controls. *DJ694/+* flies were indistinguishable from *w¹¹¹⁸* indicating that the longevity phenotype is recessive. The magnitude of the lifespan reduction ranges from 2–20% in males and 2–29% in females. When 4 backgrounds are measured simultaneously, *DJ694^H^*and *DJ694^L^* backgrounds display a more severe reduction in females than *DJ694^O^* and *DJ694^P^* whereas similar reduction are seen in males (Figure S4; Table S9). Another independent *EDTP* allele with diminished amount of EDTP protein is also short lived. Homozygous *EDTP^MI08496^* adults with reduced EDTP protein level cannot be obtained that are older than 14 days at 29°C (Komlos et al., 2023).

### Homozygous *DJ694* flies show no locomotor or muscle defects

The human ortholog of EDTP is MTMR14 also known as hJumpy because of the jumpy behavior of homozygous *DJ694* flies (Alonso et al., 2004; Tosch et al., 2006). Upon disturbance, young homozygous *DJ694* flies exhibit hyperactive reactions, and difficulties in stopping buzzing their wings after landing. At older age, they also seem to be slower and appear to sporadically display tremor. We investigated whether this behavior can be reflected and quantified with a locomotion assay that measure distance traveled, mean velocity, and the proportion of time spent immobile (Taylor Barwell et al., 2023; Barwell et al., 2021, 2022). *DJ694* did not exhibit any impairments of those parameters in the locomotion assay (Figure 2); age-matched comparisons and statistics are provided in Table S13&S14. Since *MIP/MTMR14* knockout mice show accelerated muscle wasting, we examine muscle morphology, myofibril organization and sarcomere ultrastructure but did not find any defects in *DJ694* even at older age (Figure 3).

**Figure 2.**
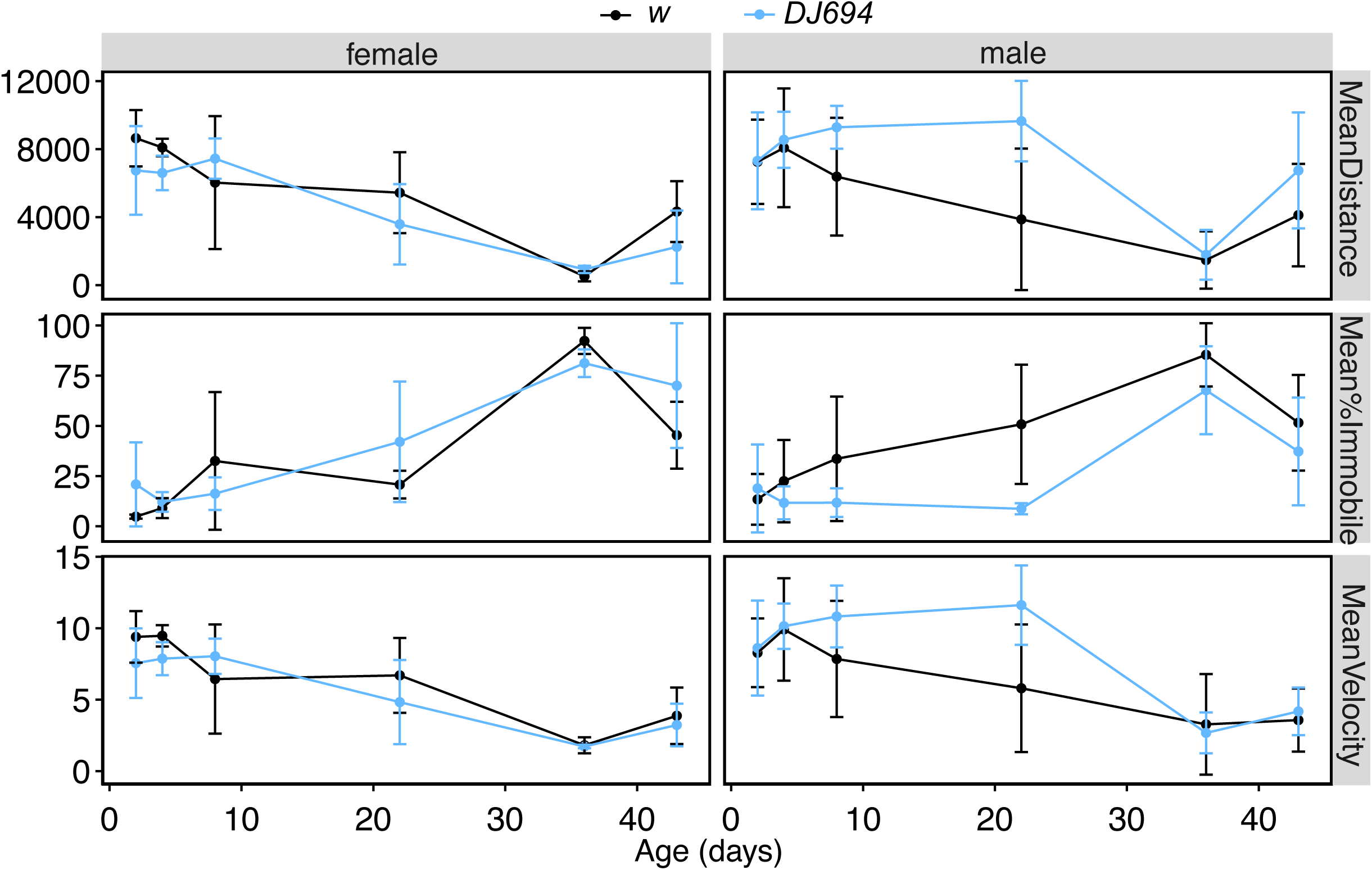
*DJ694* locomotion assay. The y-axis shows the distance traveled in mm for the mean distance, the percent spent immobile in % for the mean % immobile, and velocity in mm/s for the mean velocity. One representative replicate is shown.

**Figure 3.**
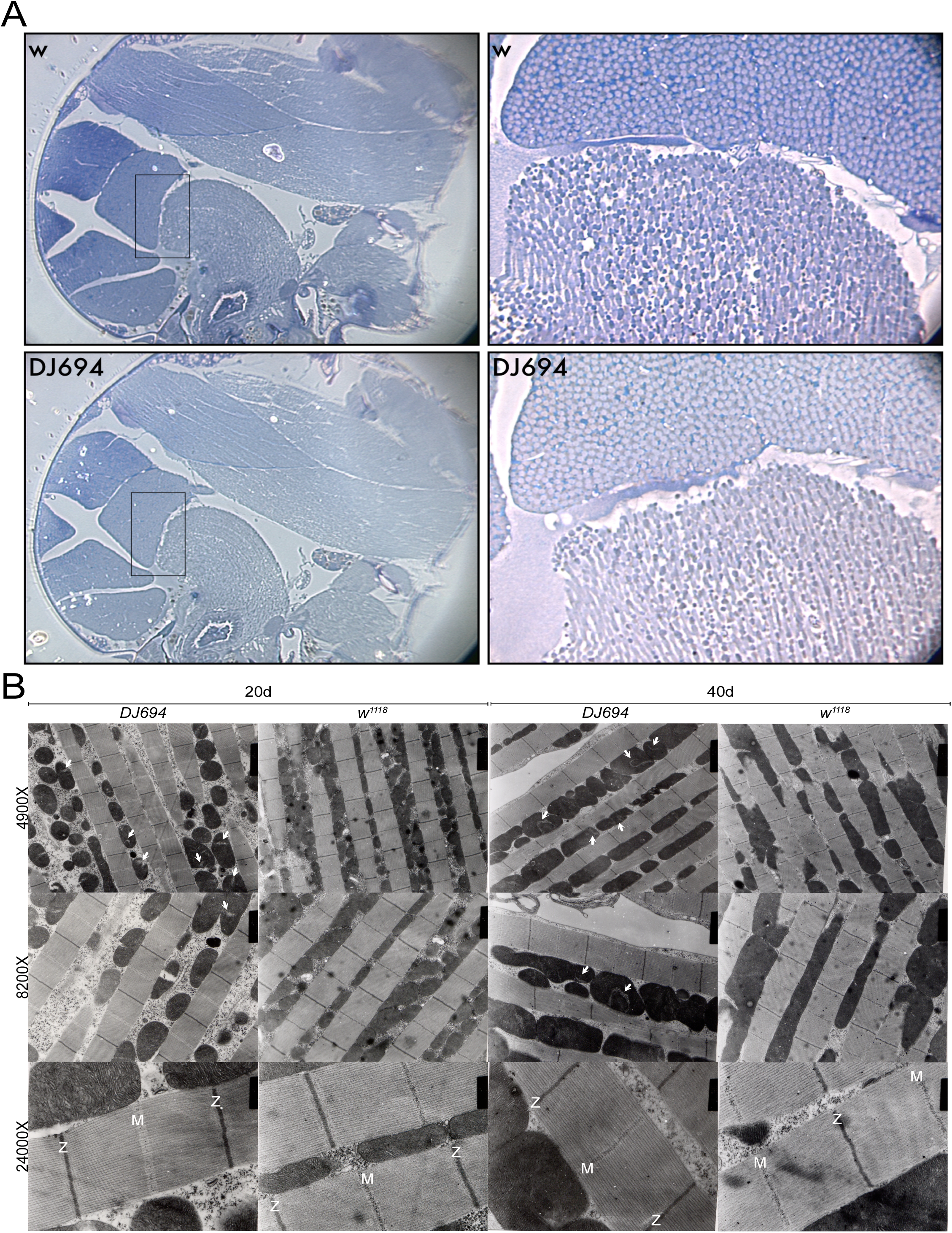
Structure of *DJ694* muscle, myofibrillar and sarcomere. A) Overall structure of thoracic muscles is shown on the left. Higher magnifications of the boxed regions show the structure of the myofibrillar. B) Ultrastructure of myofibrils and sarcomeres. Arrow point out mitochondrial swirl. Z: Z-disc. M: M-band.

### Restoring *EDTP* expression is sufficient to rescue *DJ694*’s Phenotypes

The *DJ694* insertion is responsible for the longevity and reproduction phenotypes but it does not prove that the *EDTP* gene is involved. We performed genetic rescue experiments using UAS-*EDTP* constructs because genomic rescue would require the removal of the *CG18467* gene from the first *EDTP* intron. A cDNA rescue requires choosing a promoter that mimics *EDTP*’s native expression pattern. *DJ694* is well-suited since the enhancer-trap allele itself can act as the GAL4 driver, restoring *EDTP* expression in the tissues where it is normally active. Three independent UAS-*EDTP* insertions (UAS*-EDTP^A^*, UAS*-EDTP^B^*, UAS*-EDTP^E^*) were each introduced into the *DJ694* background, and fertility and longevity were assessed. Female fertility was tested by mating *DJ694* females, *w¹¹¹⁸* females, and rescue females to *w¹¹¹⁸* males. Depending on the replicate, rescue genotypes were *DJ694;UAS-EDTP^A^*, *DJ694;UAS-EDTP^B^*, or *DJ694;UAS-EDTP^E^/+*. Eggs were scored between days 29 and 40. All three UAS*-EDTP* insertions significantly increased egg output of *DJ694* females (Figure 4A). Eggs fertilized by rescued males showed significantly higher mean adult emergence rates than those fertilized by *DJ694* males, indicating that male fertility has been restored (Figure 4B). The genotype used to rescue longevity were *DJ694;UAS-EDTP^A^/+*, *DJ694;UAS-EDTP^B^/+, DJ694;UAS-EDTP^A^/UAS-EDTP^B^*or *DJ694;UAS-EDTP^E^/+*. Almost all measurements (20 out of 22 in males, 21 out of 22 in females) showed that the addition of a UAS construct is sufficient to significantly increase the lifespan of homozygous animals (Figure 4C upward black triangles). Most of the measurements (12 out of 14 in males, 10 out of 14 in females) showed that the addition of a UAS construct result in a longevity that is similar to heterozygous animals (Figure 4C red squares). However, it cannot be excluded that the UAS-EDTP constructs drives *EDTP* expression above normal endogenous levels rather than restoring it precisely. Therefore, we tested whether *EDTP* can increase longevity or fertility in a wild-type background. Experimental animals with a copy of *DJ694* and a UAS transgene were compared to controls missing either *DJ694* or the UAS. *DJ694/+;UAS-EDTP/+* females consistently laid a number of eggs comparable to or lower than the controls (Figure S4; Table S7). We therefore conclude that EDTP does not increase fertility on its own. 10 out of 12 measurements of longevity did not show any change between experimental and control animals (Table S10), showing that *EDTP* expression in a heterozygous background is not sufficient to extend lifespan. The rescue of fertility and longevity with three independent UAS*-EDTP* insertions demonstrates that these phenotypes are solely caused by the disruption of *EDTP*.

**Figure 4.**
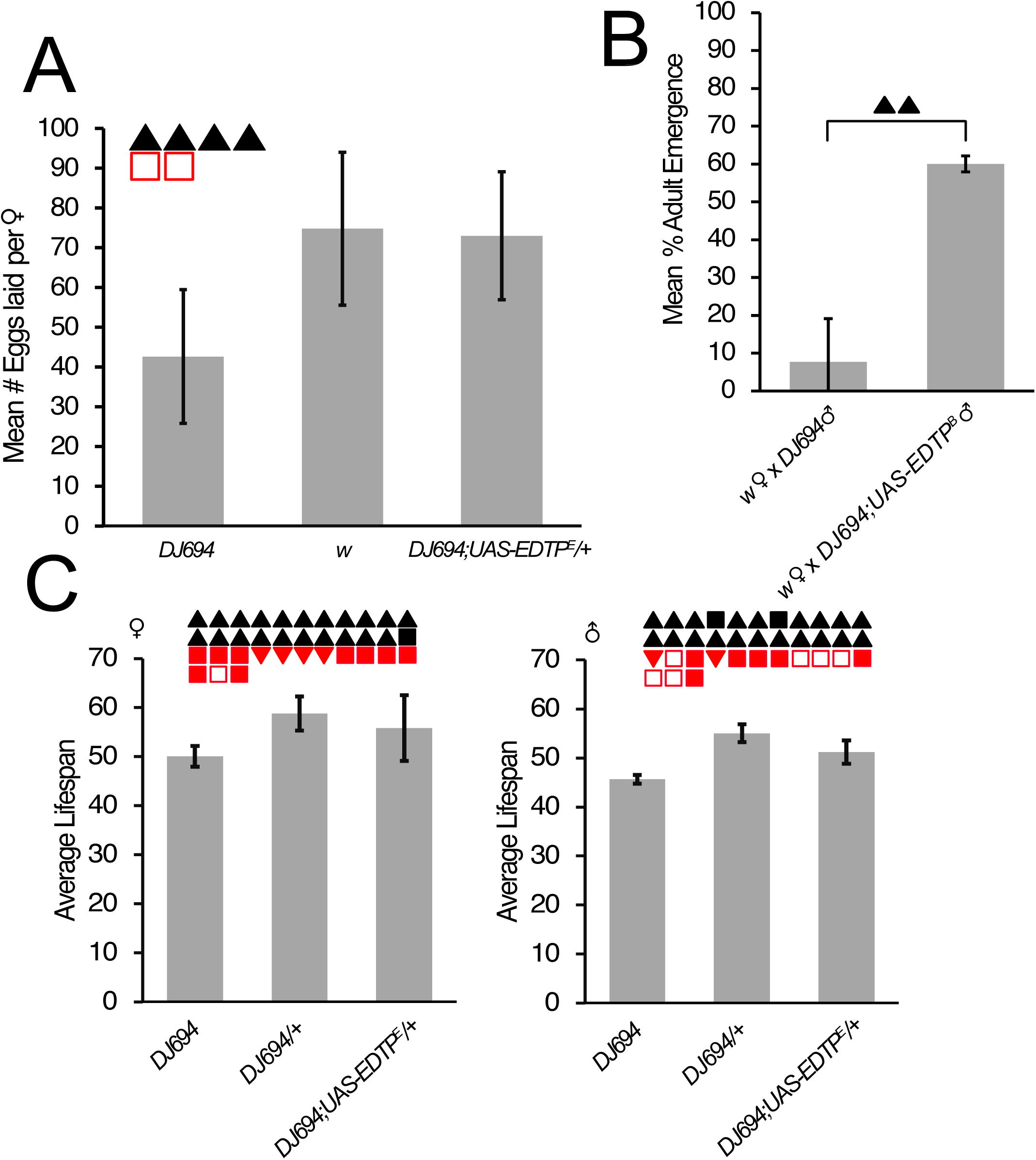
Rescue of *DJ694*’s reproduction and longevity phenotypes with UAS-*EDTP*. Each panel shows one representative replicate. Error bars represent standard deviation. Symbols indicate the statistical outcome for each independent replicate. Filled symbols indicate p < 0.05 and unfilled symbols indicate p ≥ 0.05. An upward triangle indicates an increase. A downward triangle indicates a decrease. A square indicates no change. For panel A and C, black symbols indicate comparison between rescue group to homozygous *DJ694* mutants. Red symbols indicate comparison between the rescue group to w or *DJ694/+* mutants. (A) Quantification of female fertility shown as the mean cumulative number of eggs laid per female from day 29 to day 40. (B) Quantification of male fertility is shown as the emergence percentages, calculated as the total number of adults obtained divided by the total number of eggs laid from day 29 to day 48. (C) Quantification of longevity shown as mean lifespan for the indicated genotypes.

### *DJ694* have normal ovaries at early age

To investigate the cause of the fertility defect in females, the anatomy of the ovaries of *DJ694^H^*, *DJ694^L^*, *DJ694^O^*and *DJ694^P^* backgrounds was examined. We dissected and imaged ovaries from mated and well-fed *DJ694* and *w¹¹¹⁸* females at 5 days old. We verified and confirmed that the fertility is significantly reduced at day 5 (Table S21). No obvious morphological abnormalities were observed (Figure 5A) and the quantification of ovary size revealed no significant difference between genotypes (Figure 5B; Table S15). These results suggest that the reduced fecundity in *DJ694* females is more likely due to a functional impairment than a structural defect.

**Figure 5.**
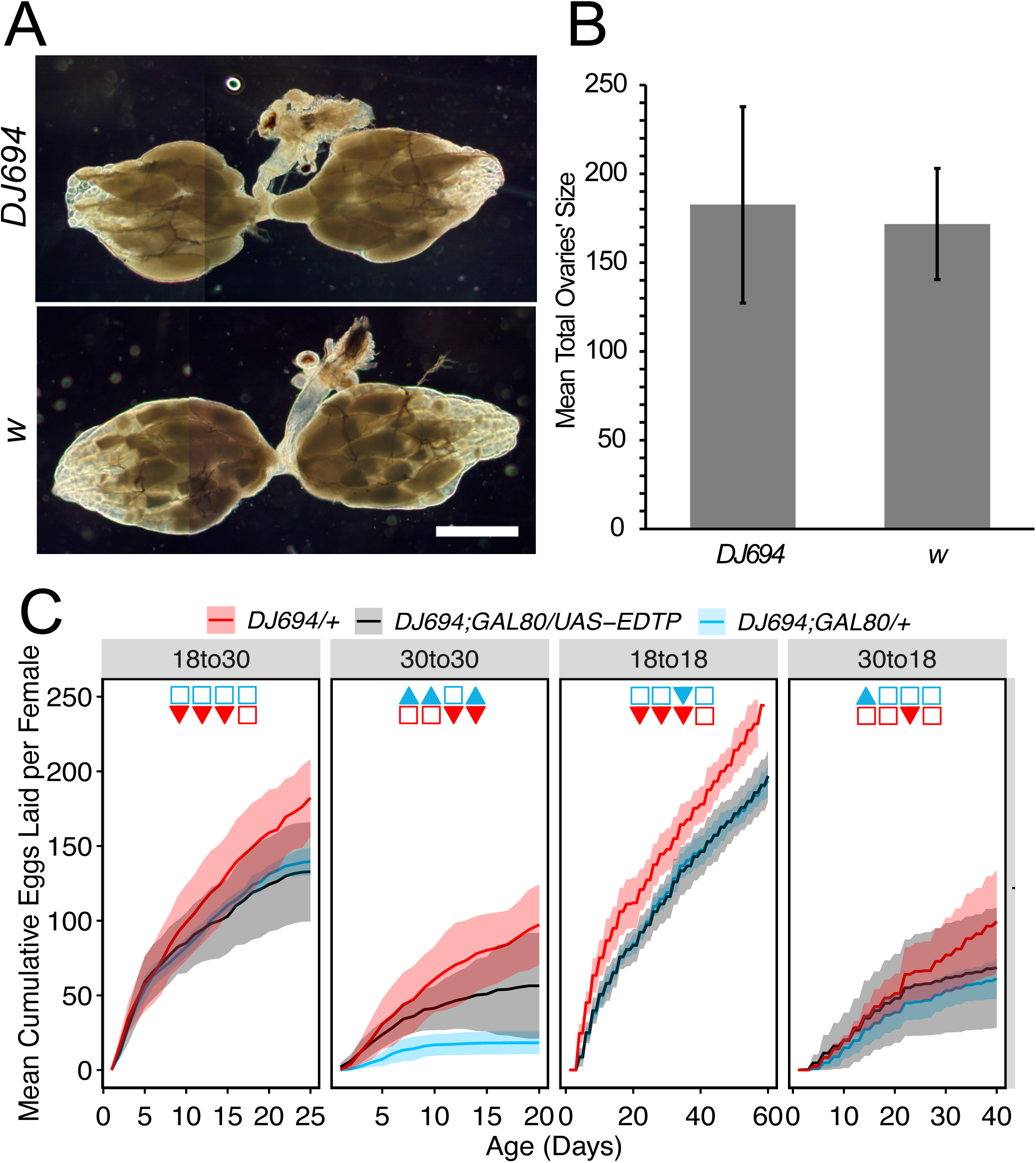
Ovary morphology and temporal rescue of female fertility for *DJ694*. (A) Representative images of ovaries from 5d females (*DJ694* vs. *w*). Scale bar = 5 μm. (B) Quantification of ovary size at day 5. y-axis unit is in μm². Bars represent mean ± SD. One representative replicate is shown. (C) Temporal rescue experiment showing mean cumulative eggs laid per female over time under different temperature conditions. Shaded regions represent standard deviation. ▴: significant increase. ▾: significant decrease. □: no change. Blue and red indicate comparisons with the negative and the positive control respectively. One representative replicate is shown.

### *EDTP* is needed during both development and adulthood to ensure normal fertility

Because *EDTP* is expressed both during embryogenesis and oogenesis, we asked at which stage of the life cycle *EDTP* is required to support normal adult female fertility by using the temperature sensitive GAL4 inhibitor, GAL80^ts^ (McGuire et al., 2003). At 18°C, GAL80 inhibits GAL4 and the transcription of the UAS transgene, preventing EDTP expression. At 30°C, GAL80^ts^ becomes inactive, allowing GAL4 to drive UAS*-EDTP* expression and restore *EDTP* activity. By shifting between these temperatures at different life stages, we can test if *EDTP* expression during development, adulthood, or both is sufficient to rescue female fertility. We previously demonstrated that GAL80^ts^ effectively suppresses the GAL4 activity of *DJ694* at 18°C during both developmental and adult stages (T. Barwell et al., 2023).The fertility of *DJ694;GAL80^ts^/UAS-EDTP* experimental females was compared to the fertility of negative *DJ694;GAL80^ts^/+* and positive *DJ694/+* controls.

Four temperature conditions were tested, where unshifted temperatures provide positive (30°C) and negative (18°C) controls and shifted temperatures as experimental groups (Figure 5C; Table S16&S17). Indeed, when UAS*-EDTP* is never expressed (18°C), experimental females are not significantly different from the homozygous control in all replicates. When UAS*-EDTP* is continuously expressed (30°C), experimental females are significantly different from the homozygous control in 3 out of 4 replicates. When UAS*-EDTP* was expressed only during adulthood (development at 18°C, adults at 30°C), experimental females are not significantly different from the homozygous control in all replicates. Therefore, *EDTP* expression during adulthood is insufficient to rescue female fertility. When UAS*-EDTP* was only expressed during developmental stages (development at 30°C, adults at 18°C), experimental females are not significantly different from the homozygous control in 3 out of 4 replicates. We therefore conclude that *EDTP* expression during development is also insufficient to rescue female fertility.

### *EDTP* decrease aggregates in the eye but not in the muscle

Since an important function of autophagy is the clearance of protein aggregates, we next asked whether EDTP’s regulation of autophagy has a measurable biological consequence in its native tissue. We recently discovered that muscle expression of a *UAS-Htt-Q72-eGFP* construct (*Q72*) encoding the first exon of human Huntingtin with 72 polyglutamine repeats fused to GFP, produces protein aggregates and reduces longevity in a dose-dependent manner (T. Barwell et al., 2023). We first examined whether expressing *EDTP* can alter the abundance of polyQ aggregates at all, using the *Drosophila* eye, a classical system widely used to model neurodegenerative disease and screen for modifiers of aggregate accumulation (Jackson et al., 1998; Warrick et al., 1998). We expressed the *Q72* construct and *EDTP* cDNA in the eye using the *GMR-GAL4* driver. Since the addition of a second UAS promoter in those experimental animals may decrease the expression of the *Q72* construct (titration effect), a *UAS-lacZ* construct was also co-expressed with Q72 to obtain an additional control that has the same number of UAS elements. When *Q72* is expressed in the eye, the eGFP-tagged aggregates are readily detectable as previously reported (Zhang et al., 2010) (Figure 6A). The co-expression of UAS*-EDTP^D^*or UAS*-EDTP^E^* obviously decreased the amount of polyglutamine aggregates whereas the co-expression of UAS-lacZ has no effect. If the reduction of aggregates is the result of autophagy-mediated clearance, it would imply that EDTP is a positive regulator of autophagy in this context, the opposite from its reported role as a negative regulator in the larval fat body and adult muscle (Komlos et al., 2023; Manzéger et al., 2021; Papp et al., 2016). Since *EDTP* is not normally expressed in the eye, we next determined whether *EDTP* has a similar effect on aggregates in its native tissue.

**Figure 6.**
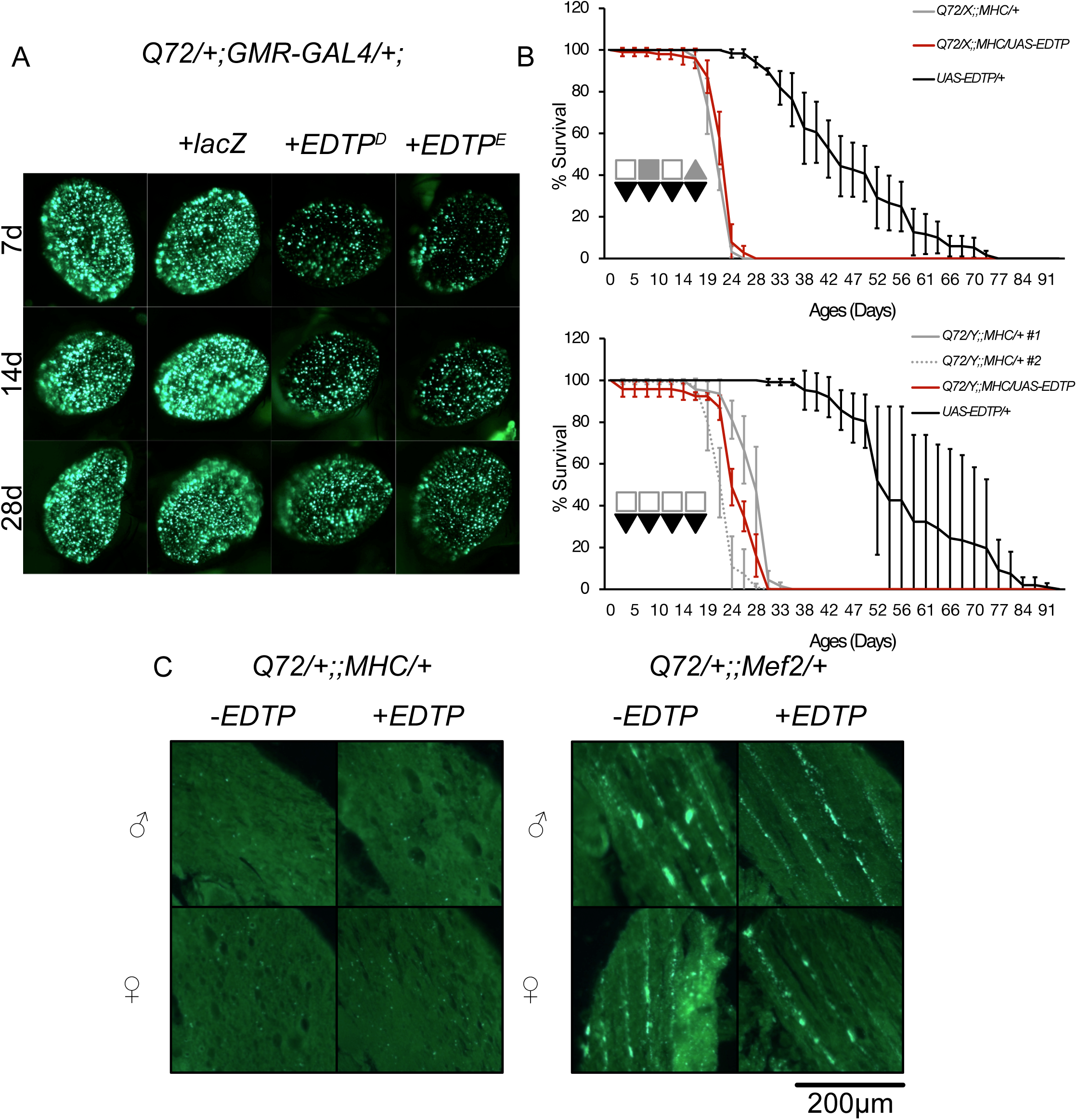
Co-expression of *EDTP* with polyglutamine aggregates. A) Co-expression of *EDTP* with polyglutamine aggregates in the eye. Aggregates are visualized by eGFP fluorescents. The left column is *Q72/+;GMR-Gal4/+;+.* The right columns show the same genotype with the indicated co-expression. B) Lifespan of flies co-expressing *EDTP* in muscles with the MHC-GeneSwitch driver treated with RU486. A representative survival curve from one of four independent biological replicates is shown. Error bars represent standard deviation. Filled symbols indicate p < 0.05 and unfilled symbols indicate p ≥ 0.05. An upward triangle indicates an increase. A downward triangle indicates a decrease. A square indicates no change. Grey and black indicate comparisons with the negative and the positive control respectively. C) Co-expression of *EDTP* with polyglutamine aggregates in the indirect flight muscle. Aggregates are visualized by immunofluorescent with anti-eGFP antibody. *–EDTP* corresponds to the genotypes on top of each panel. *+EDTP* corresponds to the same genotypes with the addition of the *UAS-EDTP^E^*.

We co-expressed UAS*-EDTP^E^* with the polyQ construct in muscle using two muscle-specific GAL4 drivers, *Mef2* and *MHC* Geneswitch, which produce different level of toxicity (T. Barwell et al., 2023). Both drivers are expressed in the muscle at all stages, but at markedly different levels. *Mef2* is highly expressed in the pupae stage; aggregates form during metamorphosis, adopt a striated distribution, and exhibit a severe reduction in lifespan. *MHC*, on the other hand, is expressed at a low level throughout development, but drive enough *Q72* expression on its own in adults to cause randomly distributed aggregates that form after emergence and a moderate decrease in lifespan. With *MHC*, both the aggregate accumulation and the longevity decline can be strongly aggravated by treatment with RU486. The co-expression of *EDTP* with the *MHC* driver did not affect longevity in either sex (4 out of 4 in males, 3 out of 4 in females) (Figure 6B; Table S11). Indeed, the co-expression of *EDTP* with the *MHC* or *Mef2* drivers did not show any effect on both kind of aggregates (Figure 6C). We therefore concluded that EDTP in adult muscles does not alleviate the detrimental effects of *Q72* aggregates on longevity.

## Discussion

As *DJ694* males aged, an increasing proportion failed to fertilize eggs: 96% to 70% produced progeny from 3 to 28 days old, compared with 100% to 95% for *w^1118^*. The cause of the male fertility defect in *DJ694* remains to be fully characterized. The reduced eclosion rate of eggs laid by females housed with *DJ694* males could be due to failure to mate, defective sperm, or impaired ejaculation. Preliminary mating assays did not support the hypothesis that the males are unable to mate, since young and mid-age *DJ694* males did mate more readily than *w¹¹¹⁸* controls within ten minutes. In mice, loss of the EDTP ortholog MTMR14 reduces the contractile force of the vas deferens (the muscular duct that propels sperm during ejaculation), and produces structurally abnormal, less motile sperm with impaired acrosome reaction, which is needed for proper fertilization (Wen et al., 2018). Ejaculation in *Drosophila* depends on contraction of the ejaculatory bulb, an organ wrapped in muscle (Cohen & Wolfner, 2018). Since *EDTP* is expressed specifically in adult muscle, a defect in ejaculatory bulb muscle function offers a plausible mechanism for *DJ694* males mating successfully while failing to deliver sperm. This could be tested by dissecting the female reproductive tract shortly after mating to confirm whether sperm reach the uterus and the sperm storage organs (seminal receptacle and paired spermathecae) (Bloch Qazi et al., 2003), using a fluorescently marked sperm line (Tomaru et al., 2018) or a simpler marker-free protocol developed for a related species (Avanesyan et al., 2017). A sperm quality defect could arise either from abnormal sperm development during spermatogenesis or from a functional deficit in normally formed sperm. Sperm morphology could be examined to determine whether spermatogenesis proceeds normally, and if sperm structure appears normal, motility of mature sperm could be assessed to test for a functional defect.

In germline clones completely lacking *EDTP*, egg chamber development is stopped at stage 4 and never reached stage 5, exactly when *EDTP* is normally first expressed (Yamaguchi et al., 2005). This shows *EDTP* is required for egg chambers to mature past stage 5. *DJ694* is not a complete loss-of-function allele and the amount of *EDTP* mRNA produced is sufficient for oogenesis to pass past stage 5. Hence, no visible change in ovary morphology was observed, but fertility is impaired. Ovarian muscle contraction, which requires juvenile hormone (JH) signaling, generates the mechanical force required for ovulation; disruption of this signaling causes retention of mature eggs and enlarged ovaries starting at day 7 (Luo et al., 2021). Only 1 replicate showed significantly larger ovaries than the wildtype control, but it cannot be excluded dissections were done too early. However, neither impaired oogenesis nor muscle contraction defect can fully account for the phenotype since adult-only expression of *EDTP* failed to rescue fertility, indicating that *EDTP* is likely needed during a different process during development. Because *EDTP* is expressed during embryogenesis (Yamaguchi et al., 2005), when the ovaries begin to form (Gilboa, 2015; Jin & Zhao, 2023), one can speculate that missing *EDTP* during development impacts the establishment of JH signaling which later impacts fertility. The terminal filament (TF) continuously imports lipophilic molecules from the hemolymph and delivers them to the germarium, supporting germ cell and follicle development throughout adult life (Maurya & Spradling, 2026). TF cells differentiate during larval stages from a somatic precursor population that itself traces back to the embryonic gonad. If *EDTP*’s embryonic role affects the size or quality of this founding precursor population, a smaller or less robust TF could result in limiting lipid supply to the germarium therefore reducing egg production.

Phosphoinositide metabolism is involved in both autophagy and insulin signaling, which are both linked to longevity. However, EDTP does not use PIP_3_ or PI(4,5)P_2_ as substrates, the core lipids of insulin/Akt signaling, making a direct role for EDTP in this pathway unlikely. The relationship between autophagy and lifespan depends on the specific gene manipulated and the tissue involved. In the nervous system, the suppression of autophagy by *Atg8* loss of function shortens the lifespan and increases the accumulation of cellular damages (Simonsen et al., 2008). Reciprocally the activation of autophagy by overexpression of *Atg8a* or *Atg1* extends lifespan (Simonsen et al., 2008; Ulgherait et al., 2014). In the adult fat body, Atg1 overexpression can either extend or shorten lifespan depending on dosage (Bjedov et al., 2020). Contradictory effects have also been observed upon experimental alteration of autophagy in muscles. Both *FOXO/4E-BP* overexpression, which increases basal autophagic and lysosomal activity, and *Atg8a* overexpression have been reported to extend lifespan (Demontis & Perrimon, 2010)(Bai et al., 2013). Yet, a more recent study could not replicate the life extension using independent *Atg8a* transgenic lines, and instead found that *Atg1* overexpression in adult muscle induce ubiquitinated protein aggregates and shortens lifespan (Bierlein et al., 2023). The apparently contradictory results are indeed due to the degree of autophagy activation. Activation is beneficial, but excessive autophagy is detrimental. Flies with overactivation of autophagy displays a reduced lifespan and muscle atrophy (Bargiela et al., 2015; Blazquez-Bernal et al., 2021). The *EDTP^MI08496^* allele with severely reduced EDTP protein level showed increased autophagy (decreased Ref2P) and shortened lifespan (Komlos et al., 2023). This study demonstrates that the alteration of *EDTP* transcript in the *DJ694* allele is responsible for the reduction of the longevity, but we only assess the involvement of *EDTP* in autophagy by testing the clearance of polyglutamine aggregates. If EDTP inhibits autophagy in muscle, overexpressing *EDTP* should decrease autophagy, increase aggregates, and further shorten lifespan. Muscle *EDTP* overexpression did not affect the abundance or appearance of polyglutamine aggregates nor it improves or aggravates the effect of the aggregates on longevity. This may suggest that autophagy is not the mechanism underlying the effect on longevity although autophagy and lifespan are both affected by loss of *EDTP* in muscles. An alternative possibility is that autophagy is not important for the muscle. RNAi knockdown of key autophagy genes (*Atg1*, *Atg18*) in muscle did not alter autophagy or lifespan, whereas the same knockdown in adipose tissue nearly abolished autophagosome formation and reduced lifespan (Bierlein et al., 2023). Ectopic *EDTP* expression in the eye decreased the amount of polyQ aggregates, though whether this reflects an effect on aggregate clearance or formation is unknown. Notably, identical effect was reported with the co-expression of the chaperone *Hsp70* (Taylor Barwell et al., 2023), raising the possibility that EDTP is involved in protein folding rather than autophagy. However, if EDTP functions as a molecular chaperone, a reduction of the amount of aggregates would have been expected when *EDTP* is overexpressed in its native tissue.

## Reference

Alonso, A., Sasin, J., Bottini, N., Friedberg, I., Friedberg, I., Osterman, A., Godzik, A., Hunter, T., Dixon, J., & Mustelin, T. (2004). Protein tyrosine phosphatases in the human genome. Cell, 117(6), 699–711. 10.1016/j.cell.2004.05.018

Avanesyan, A., Jaffe, B. D., & Guédot, C. (2017). Isolating Spermathecae and Determining Mating Status of Drosophila suzukii: A Protocol for Tissue Dissection and Its Applications. Insects, 8(1). 10.3390/insects8010032

Bai, H., Kang, P., Hernandez, A. M., & Tatar, M. (2013). Activin Signaling Targeted by Insulin/dFOXO Regulates Aging and Muscle Proteostasis in Drosophila. PLOS Genetics, 9(11), e1003941. 10.1371/journal.pgen.1003941

Bargiela, A., Herreros, E., Fernández Costa, J. M., Vílchez, J., Llamusi, B., & Artero, R. (2015). Increased autophagy and apoptosis contribute to muscle atrophy in a myotonic dystrophy type 1 Drosophila model. Disease Models & Mechanisms, 8, 679–690. 10.1242/dmm.018127

Barwell, T., DeVeale, B., Poirier, L., Zheng, J., Seroude, F., & Seroude, L. (2017). Regulating the UAS/GAL4 system in adult Drosophila with Tet-off GAL80 transgenes. PeerJ, 5, e4167. 10.7717/peerj.4167

Barwell, T., Geld, S., & Seroude, L. (2023). Comparison of GAL80ts and Tet-off GAL80 transgenes. MicroPubl Biol, 2023. 10.17912/micropub.biology.000770

Barwell, T., Raina, S., Page, A., MacCharles, H., & Seroude, L. (2023). Juvenile and adult expression of polyglutamine expanded huntingtin produce distinct aggregate distributions in Drosophila muscle. Human Molecular Genetics, 32(16), 2656–2668. 10.1093/hmg/ddad098

Barwell, T., Raina, S., & Seroude, L. (2021). Versatile method to measure locomotion in adult Drosophila. Genome, 64(2), 139–145. 10.1139/gen-2020-0044%M32552119

Barwell, T., Raina, S., & Seroude, L. (2022). Protocol for recording and analyzing spontaneous locomotion in Drosophila. STAR Protocols, 3(4), 101888. 10.1016/j.xpro.2022.101888

Bierlein, M., Charles, J., Polisuk-Balfour, T., Bretscher, H., Rice, M., Zvonar, J., Pohl, D., Winslow, L., Wasie, B., Deurloo, S., Van Wert, J., Williams, B., Ankney, G., Harmon, Z., Dann, E., Azuz, A., Guzman-Vargas, A., Kuhns, E., Neufeld, T. P., … Zhu, C. C. (2023). Autophagy impairment and lifespan reduction caused by Atg1 RNAi or Atg18 RNAi expression in adult fruit flies (Drosophila melanogaster). Genetics, 225(2). 10.1093/genetics/iyad154

Billes, V., Kovács, T., Hotzi, B., Manzéger, A., Tagscherer, K., Komlós, M., Tarnóci, A., Pádár, Z., Erdős, A., Bjelik, A., Legradi, A., Gulya, K., Gulyás, B., & Vellai, T. (2016). AUTEN-67 (Autophagy Enhancer-67) Hampers the Progression of Neurodegenerative Symptoms in a Drosophila model of Huntington’s Disease. J Huntingtons Dis, 5(2), 133–147. 10.3233/jhd-150180

Bjedov, I., Cochemé, H. M., Foley, A., Wieser, D., Woodling, N. S., Castillo-Quan, J. I., Norvaisas, P., Lujan, C., Regan, J. C., Toivonen, J. M., Murphy, M. P., Thornton, J., Kinghorn, K. J., Neufeld, T. P., Cabreiro, F., & Partridge, L. (2020). Fine-tuning autophagy maximises lifespan and is associated with changes in mitochondrial gene expression in Drosophila. PLoS Genet, 16(11), e1009083. 10.1371/journal.pgen.1009083

Blazquez-Bernal, A., Fernandez-Costa, J. M., Bargiela, A., & Artero, R. (2021). Inhibition of autophagy rescues muscle atrophy in a LGMDD2 Drosophila model. FASEB J, 35(10), e21914. 10.1096/fj.202100539RR

Bloch Qazi, M. C., Heifetz, Y., & Wolfner, M. F. (2003). The developments between gametogenesis and fertilization: ovulation and female sperm storage in drosophila melanogaster. Developmental Biology, 256(2), 195–211. 10.1016/S0012-1606(02)00125-2

Cohen, A. B., & Wolfner, M. F. (2018). Dynamic changes in ejaculatory bulb size during Drosophila melanogaster aging and mating. J Insect Physiol, 107, 152–156. 10.1016/j.jinsphys.2018.04.005

Demontis, F., & Perrimon, N. (2010). FOXO/4E-BP Signaling in Drosophila Muscles Regulates Organism-wide Proteostasis during Aging. Cell, 143, 813–825. https://www.sciencedirect.com/science/article/pii/S0092867410011438?via%3Dihub

Dowling, J. J., Low, S. E., Busta, A. S., & Feldman, E. L. (2010). Zebrafish MTMR14 is required for excitation-contraction coupling, developmental motor function and the regulation of autophagy. Hum Mol Genet, 19(13), 2668–2681. 10.1093/hmg/ddq153

Fleming, A., Bourdenx, M., Fujimaki, M., Karabiyik, C., Krause, G. J., Lopez, A., Martín-Segura, A., Puri, C., Scrivo, A., Skidmore, J., Son, S. M., Stamatakou, E., Wrobel, L., Zhu, Y., Cuervo, A. M., & Rubinsztein, D. C. (2022). The different autophagy degradation pathways and neurodegeneration. Neuron, 110(6), 935–966. 10.1016/j.neuron.2022.01.017

Gilboa, L. (2015). Organizing stem cell units in the Drosophila ovary. Curr Opin Genet Dev, 32, 31–36. 10.1016/j.gde.2015.01.005

Hnia, K., Kretz, C., Amoasii, L., Böhm, J., Liu, X., Messaddeq, N., Qu, C.-K., & Laporte, J. (2012). Primary T-tubule and autophagy defects in the phosphoinositide phosphatase Jumpy/MTMR14 knockout mice muscle. Advances in Biological Regulation, 52(1), 98–107. 10.1016/j.advenzreg.2011.09.007

Jackson, G. R., Salecker, I., Dong, X., Yao, X., Arnheim, N., Faber, P. W., MacDonald, M. E., & Zipursky, S. L. (1998). Polyglutamine-expanded human huntingtin transgenes induce degeneration of Drosophila photoreceptor neurons. Neuron, 21(3), 633–642. 10.1016/s0896-6273(00)80573-5

Jin, J., & Zhao, T. (2023). Niche formation and function in developing tissue: studies from the Drosophila ovary. Cell Commun Signal, 21(1), 23. 10.1186/s12964-022-01035-7

Karess, R. E., & Rubin, G. M. (1984). Analysis of P transposable element functions in Drosophila. Cell, 38(1), 135–146. 10.1016/0092-8674(84)90534-8

Kim, S. A., Taylor, G. S., Torgersen, K. M., & Dixon, J. E. (2002). Myotubularin and MTMR2, phosphatidylinositol 3-phosphatases mutated in myotubular myopathy and type 4B Charcot-Marie-Tooth disease. J Biol Chem, 277(6), 4526–4531. 10.1074/jbc.M111087200

Komlos, M., Szinyakovics, J., Falcsik, G., Sigmond, T., Jezso, B., Vellai, T., & Kovacs, T. (2023). The Small-Molecule Enhancers of Autophagy AUTEN-67 and -99 Delay Ageing in Drosophila Striated Muscle Cells. Int J Mol Sci, 24(9). 10.3390/ijms24098100

Kovács, T., Billes, V., Komlós, M., Hotzi, B., Manzéger, A., Tarnóci, A., Papp, D., Szikszai, F., Szinyákovics, J., Rácz, Á., Noszál, B., Veszelka, S., Walter, F. R., Deli, M. A., Hackler, L., Alfoldi, R., Huzian, O., Puskas, L. G., Liliom, H., … Vellai, T. (2017). The small molecule AUTEN-99 (autophagy enhancer-99) prevents the progression of neurodegenerative symptoms. Scientific Reports, 7(1), 42014. 10.1038/srep42014

Laporte, J., Biancalana, V., Tanner, S. M., Kress, W., Schneider, V., Wallgren-Pettersson, C., Herger, F., Buj-Bello, A., Blondeau, F., Liechti-Gallati, S., & Mandel, J.-L. (2000). MTM1 mutations in X-linked myotubular myopathy. Human Mutation, 15(5), 393–409. 10.1002/(SICI)1098-1004(200005)15:5<393::AID-HUMU1>3.0.CO;2-R

Laporte, J., Hu, L. J., Kretz, C., Mandel, J. L., Kioschis, P., Coy, J. F., Klauck, S. M., Poustka, A., & Dahl, N. (1996). A gene mutated in X-linked myotubular myopathy defines a new putative tyrosine phosphatase family conserved in yeast. Nat Genet, 13(2), 175–182. 10.1038/ng0696-175

Liu, J., Lv, Y., Liu, Q. H., Qu, C. K., & Shen, J. (2014). Deficiency of MTMR14 promotes autophagy and proliferation of mouse embryonic fibroblasts. Mol Cell Biochem, 392(1-2), 31–37. 10.1007/s11010-014-2015-5

Luo, W., Liu, S., Zhang, W., Yang, L., Huang, J., Zhou, S., Feng, Q., Palli, S. R., Wang, J., Roth, S., & Li, S. (2021). Juvenile hormone signaling promotes ovulation and maintains egg shape by inducing expression of extracellular matrix genes. Proceedings of the National Academy of Sciences, 118(39), e2104461118. doi:10.1073/pnas.2104461118

Manzéger, A., Tagscherer, K., Lőrincz, P., Szaker, H., Lukácsovich, T., Pilz, P., Kméczik, R., Csikós, G., Erdélyi, M., Sass, M., Kovács, T., Vellai, T., & Billes, V. A. (2021). Condition-dependent functional shift of two Drosophila Mtmr lipid phosphatases in autophagy control. Autophagy, 17(12), 4010–4028. 10.1080/15548627.2021.1899681

Maurya, B., & Spradling, A. C. (2026). The Drosophila ovarian terminal filament imports lipophilic molecules that support cyst and follicle development within the ovariole. Dev Biol, 538, 9–20. 10.1016/j.ydbio.2026.06.013

McGuire, S. E., Le, P. T., Osborn, A. J., Matsumoto, K., & Davis, R. L. (2003). Spatiotemporal Rescue of Memory Dysfunction in Drosophila. Science, 302(5651), 1765–1768. 10.1126/science.1089035

Papp, D., Kovács, T., Billes, V., Varga, M., Tarnóci, A., Hackler Jr, L., Puskás, L. G., Liliom, H., Tárnok, K., Schlett, K., Borsy, A., Pádár, Z., Kovács, A. L., Hegedűs, K., Juhász, G., Komlós, M., Erdős, A., Gulyás, B., & Vellai, T. (2016). AUTEN-67, an autophagy-enhancing drug candidate with potent antiaging and neuroprotective effects. Autophagy, 12(2), 273–286. 10.1080/15548627.2015.1082023

Partridge, L., & Gems, D. (2007). Benchmarks for ageing studies. Nature, 450(7167), 165–167. 10.1038/450165a

Renfranz, P. J., & Benzer, S. (1989). Monoclonal antibody probes discriminate early and late mutant defects in development of the Drosophila retina. Dev Biol, 136(2), 411–429. 10.1016/0012-1606(89)90267-4

Robertson, H. M., Preston, C. R., Phillis, R. W., Johnson-Schlitz, D. M., Benz, W. K., & Engels, W. R. (1988). A stable genomic source of P element transposase in Drosophila melanogaster. Genetics, 118(3), 461–470. 10.1093/genetics/118.3.461

Romero-Suarez, S., Shen, J., Brotto, L., Hall, T., Mo, C., Valdivia, H. H., Andresen, J., Wacker, M., Nosek, T. M., Qu, C. K., & Brotto, M. (2010). Muscle-specific inositide phosphatase (MIP/MTMR14) is reduced with age and its loss accelerates skeletal muscle aging process by altering calcium homeostasis. Aging (Albany NY), 2(8), 504–513. 10.18632/aging.100190

Seroude, L., Brummel, T., Kapahi, P., & Benzer, S. (2002). Spatio-temporal analysis of gene expression during aging in Drosophila melanogaster. Aging Cell, 1(1), 47–56. 10.1046/j.1474-9728.2002.00007.x

Shen, J., Yu, W. M., Brotto, M., Scherman, J. A., Guo, C., Stoddard, C., Nosek, T. M., Valdivia, H. H., & Qu, C. K. (2009). Deficiency of MIP/MTMR14 phosphatase induces a muscle disorder by disrupting Ca(2+) homeostasis. Nat Cell Biol, 11(6), 769–776. 10.1038/ncb1884

Simonsen, A., Cumming, R. C., Brech, A., Isakson, P., Schubert, D. R., & Finley, K. D. (2008). Promoting basal levels of autophagy in the nervous system enhances longevity and oxidant resistance in adult Drosophila. Autophagy, 4(2), 176–184. 10.4161/auto.5269

Singh, S. H., Ramachandra, N. B., & Nongthomba, U. (2014). Egg-derived tyrosine phosphatase as a potential biomarker for muscle ageing and degeneration in Drosophila melanogaster. J Genet Genomics, 41(4), 221–224. 10.1016/j.jgg.2014.01.008

Spradling, A. C., & Rubin, G. M. (1982). Transposition of cloned P elements into Drosophila germ line chromosomes. Science, 218(4570), 341–347. 10.1126/science.6289435

Taylor, G. S., & Dixon, J. E. (2003). PTEN and myotubularins: families of phosphoinositide phosphatases. Methods Enzymol, 366, 43–56. 10.1016/s0076-6879(03)66004-0

Taylor, G. S., Maehama, T., & Dixon, J. E. (2000). Myotubularin, a protein tyrosine phosphatase mutated in myotubular myopathy, dephosphorylates the lipid second messenger, phosphatidylinositol 3-phosphate. Proc Natl Acad Sci U S A, 97(16), 8910–8915. 10.1073/pnas.160255697

Tomaru, M., Ohsako, T., Watanabe, M., Juni, N., Matsubayashi, H., Sato, H., Takahashi, A., & Yamamoto, M.-T. (2018). Severe Fertility Effects of sheepish Sperm Caused by Failure To Enter Female Sperm Storage Organs in Drosophila melanogaster. G3 Genes|Genomes|Genetics, 8(1), 149–160. 10.1534/g3.117.300171

Tosch, V., Rohde, H. M., Tronchère, H., Zanoteli, E., Monroy, N., Kretz, C., Dondaine, N., Payrastre, B., Mandel, J.-L., & Laporte, J. (2006). A novel PtdIns3P and PtdIns(3,5)P2 phosphatase with an inactivating variant in centronuclear myopathy. Human Molecular Genetics, 15(21), 3098–3106. 10.1093/hmg/ddl250

Ulgherait, M., Rana, A., Rera, M., Graniel, J., & Walker, D. W. (2014). AMPK modulates tissue and organismal aging in a non-cell-autonomous manner. Cell Rep, 8(6), 1767–1780. 10.1016/j.celrep.2014.08.006

Vergne, I., Roberts, E., Elmaoued, R. A., Tosch, V., Delgado, M. A., Proikas-Cezanne, T., Laporte, J., & Deretic, V. (2009). Control of autophagy initiation by phosphoinositide 3-phosphatase Jumpy. Embo j, 28(15), 2244–2258. 10.1038/emboj.2009.159

Warrick, J. M., Paulson, H. L., Gray-Board, G. L., Bui, Q. T., Fischbeck, K. H., Pittman, R. N., & Bonini, N. M. (1998). Expanded polyglutamine protein forms nuclear inclusions and causes neural degeneration in. Cell, 93(6), 939–949. Doi 10.1016/S0092-8674(00)81200-3

Wen, N., Yu, M. F., Liu, J., Cai, C., Liu, Q. H., & Shen, J. (2018). Deficiency of MTMR14 impairs male fertility in Mus musculus. PLoS One, 13(11), e0206224. 10.1371/journal.pone.0206224

Xiao, C. F., Mileva-Seitz, V., Seroude, L., & Robertson, R. M. (2007). Targeting HSP70 to motoneurons protects locomotor activity from hyperthermia in. Developmental Neurobiology, 67(4), 438–455. 10.1002/dneu.20344

Yamaguchi, S., Homma, K.-i., & Natori, S. (1999). A novel egg-derived tyrosine phosphatase, EDTP, that participates in the embryogenesis of Sarcophaga peregrina (flesh fly) [10.1046/j.1432-1327.1999.00143.x]. European Journal of Biochemistry, 259(3), 946–955. 10.1046/j.1432-1327.1999.00143.x

Yamaguchi, S., Katagiri, S., Sekimizu, K., Natori, S., & Homma, K. J. (2005). Involvement of EDTP, an egg-derived tyrosine phosphatase, in the early development of Drosophila melanogaster. J Biochem, 138(6), 721–728. 10.1093/jb/mvi176

Yamamoto, H., Zhang, S., & Mizushima, N. (2023). Autophagy genes in biology and disease. Nat Rev Genet, 24(6), 382–400. 10.1038/s41576-022-00562-w

Zhang, S., Binari, R., Zhou, R., & Perrimon, N. (2010). A genomewide RNA interference screen for modifiers of aggregates formation by mutant Huntingtin in Drosophila. Genetics, 184(4), 1165–1179. 10.1534/genetics.109.112516

