## Supplemental Figures and Tables for "EDTP Loss of Function Impairs Longevity and Reproduction"

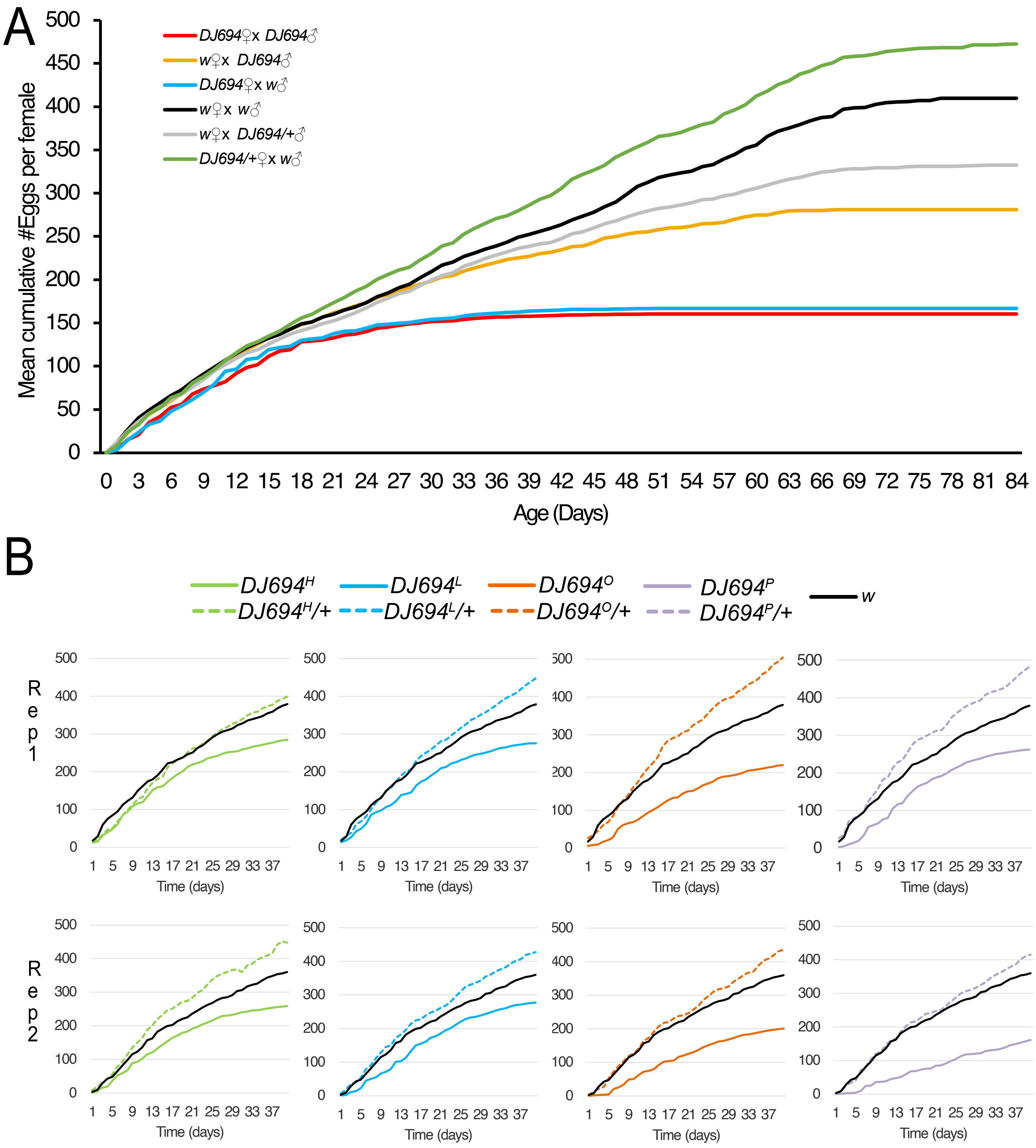


**Figure S1. Female fertility of *DJ694* across the lifespan and across multiple genetic backgrounds** Error bars are not shown for clarity. (A) Cumulative number of eggs laid per female across age. Curves represent the cumulative mean number of eggs laid per female for one replicate. (B) Cumulative number of eggs laid per female from day 1 to day 40. Four independent *DJ694* backgrounds were tested. One representative replicate is shown.

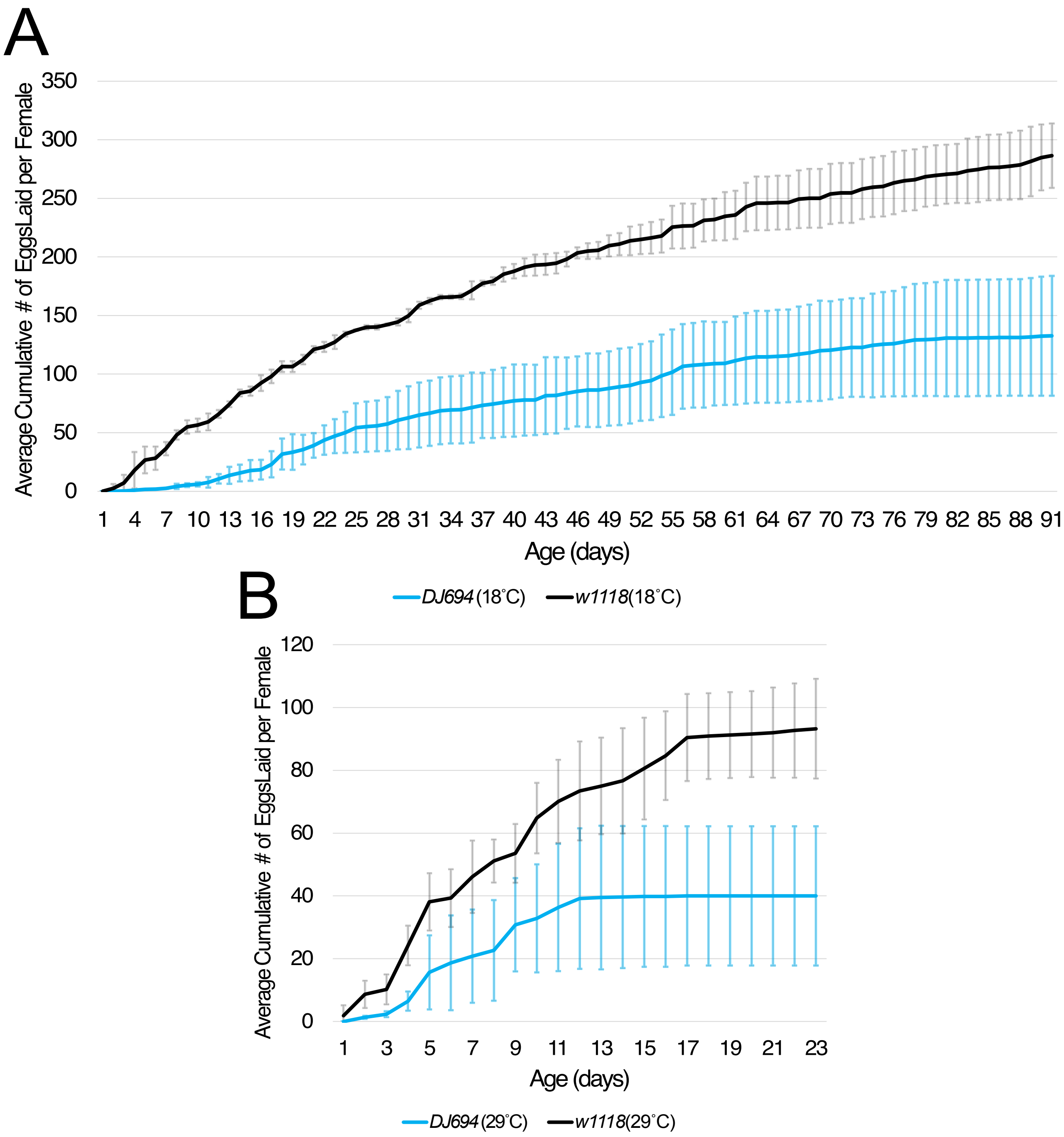


**Figure S2.** ***DJ694*** **female fertility phenotype at 18°C and 29°C** Mean cumulative number of eggs laid per female over time for *DJ694* and $w^{1118}$ females held at (A) 18°C or (B) 29°C. Flies were raised at 25°C and transferred to the indicated temperature upon emergence. All females were mated with an equal number of $w^{1118}$ males. Error bars represent standard deviation.

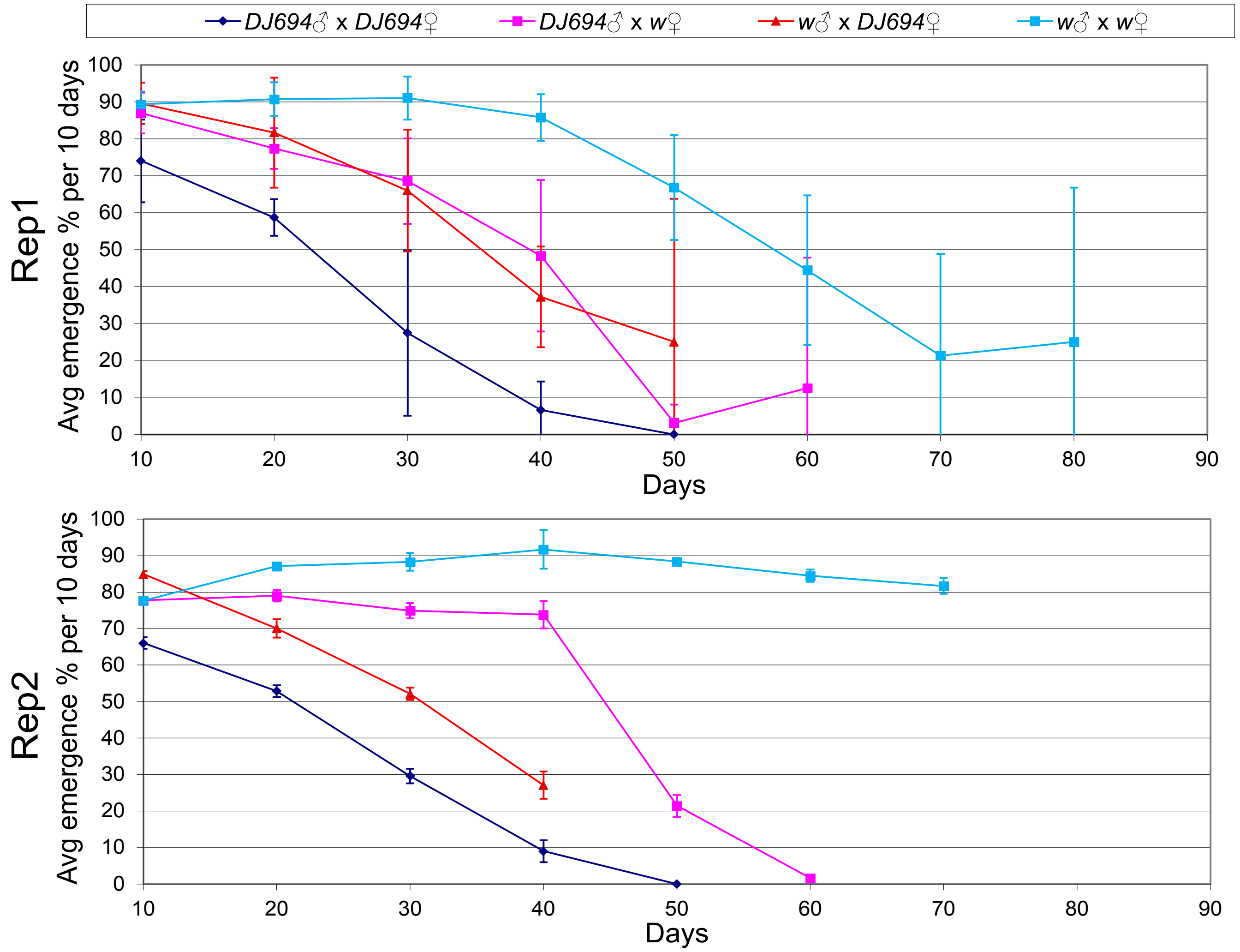


**Figure S3. Average emergence percentage per 10-day interval at 25°C.** For each day, the emergence percentage was calculated as the number of adults emerged divided by the number of eggs laid that day, then averaged across three vials to give a daily mean. These daily means were then averaged across each 10-day interval, and error bars represent the standard deviation of the daily means within each interval. Two independent replicates are shown.

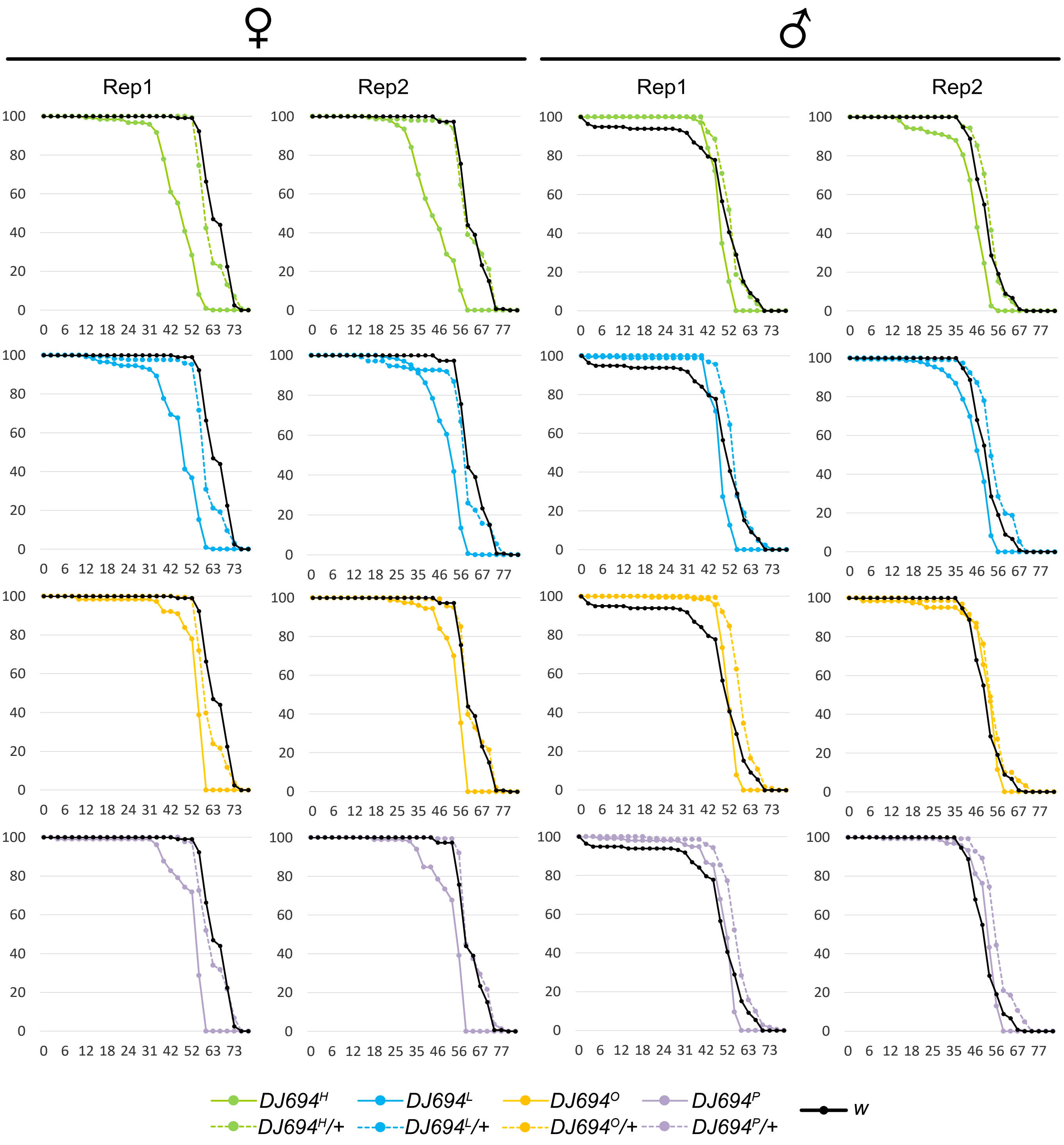


**Figure S4. Survival curves for homozygous *DJ694* and $w^{1118}$ controls shown separately for females and males across two independent replicates** The y-axis represents survival percentage and the x-axis represents age in days. Four independent *DJ694* backgrounds were tested. Error bars are not shown for clarity.

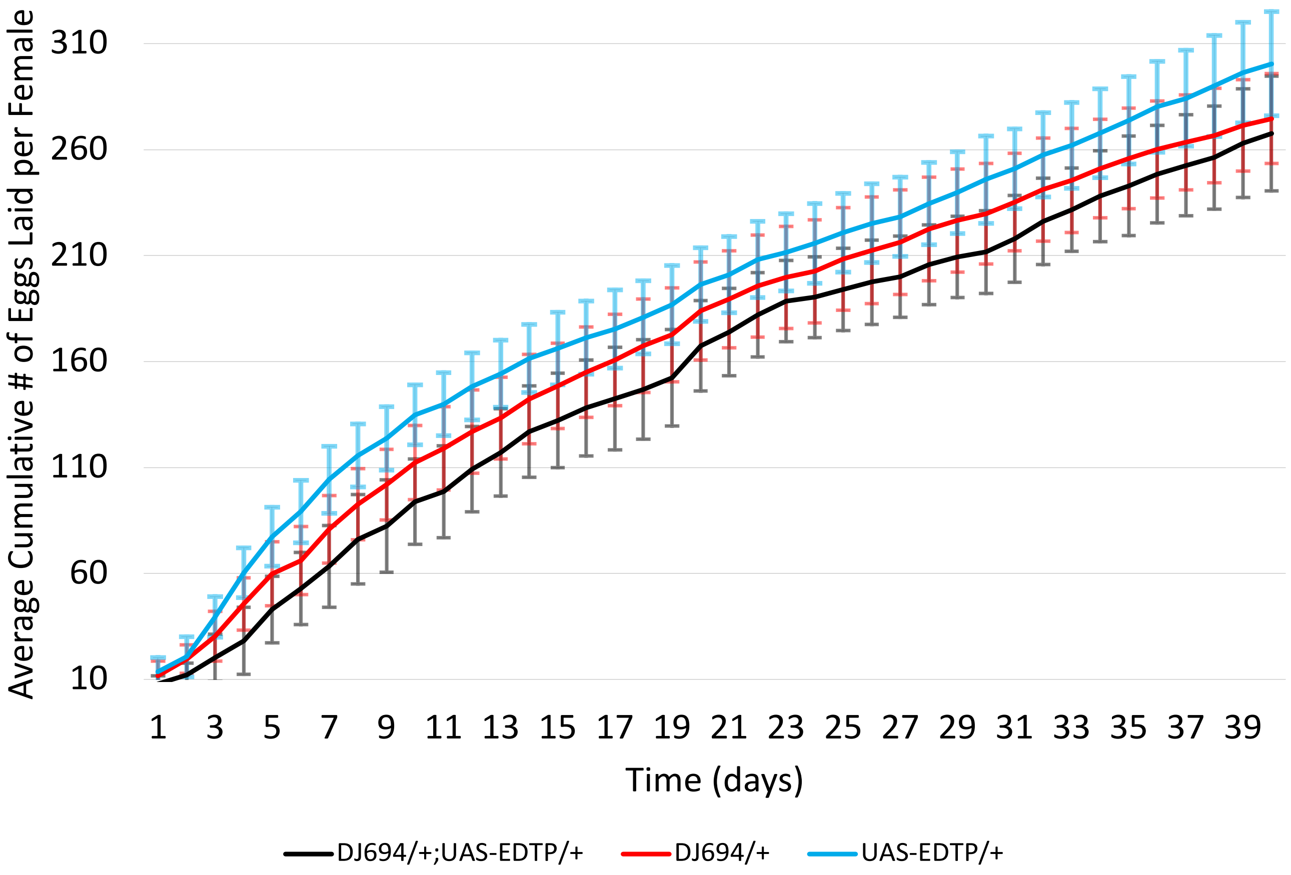


**Figure S5. Expression of *EDTP* in *DJ694/+* does not increase female fertility** Cumulative number of eggs laid per female from day 1 to day 40. Error bars represent standard deviation averaged across vials.

| | | Cross | | Date | | | | | Mean | | | SD | | | P (Homo vs. Wt) | | | P (Homo vs. Hetero) | | |
|---|---|---|---|---|---|---|---|---|---|---|---|---|---|---|---|---|---|---|---|---|
| **Exp** | **Rep** | **♂** | **V♀** | **(YY-MM-DD)** | **# Eggs** | **% L1** | **% P** | **% A** | **L1** | **P** | **A** | **L1** | **P** | **A** | **L1** | **P** | **A** | **L1** | **P** | **A** |
| 1 | A | $DJ694^{H}$ | $DJ694^{H}$ | 24-02-05 | 25 | 76 | 63 | 100 | 69.33 | 74.10 | 96.30 | 11.55 | 17.94 | 6.41 | - | - | - | 0.1776 | 0.6014 | 0.4950 |
| 1 | A | $DJ694^{H}$ | $DJ694^{H}$ | 24-02-05 | 25 | 76 | 95 | 89 | | | | | | | | | | | | |
| 1 | A | $DJ694^{H}$ | $DJ694^{H}$ | 24-02-05 | 25 | 56 | 64 | 100 | | | | | | | | | | | | |
| 1 | B | $DJ694^{H}$ | $DJ694^{H}$ | 24-02-06 | 25 | 76 | 69 | 100 | 70.00 | 84.78 | 93.55 | 5.16 | 11.23 | 5.12 | 0.0069 | 0.4384 | 0.5725 | 0.3388 | 0.9883 | 0.1402 |
| 1 | B | $DJ694^{H}$ | $DJ694^{H}$ | 24-02-06 | 25 | 64 | 88 | 93 | | | | | | | | | | | | |
| 1 | B | $DJ694^{H}$ | $DJ694^{H}$ | 24-02-06 | 25 | 68 | 94 | 94 | | | | | | | | | | | | |
| 1 | B | $DJ694^{H}$ | $DJ694^{H}$ | 24-02-06 | 25 | 72 | 89 | 88 | | | | | | | | | | | | |
| 1 | C | $DJ694^{H}$ | $DJ694^{H}$ | 24-02-07 | 25 | 72 | 78 | 100 | 63.00 | 84.58 | 92.30 | 6.83 | 6.73 | 8.95 | 0.0359 | 0.0050 | 0.1359 | 0.0033 | 0.5694 | 0.4984 |
| 1 | C | $DJ694^{H}$ | $DJ694^{H}$ | 24-02-07 | 25 | 64 | 81 | 100 | | | | | | | | | | | | |
| 1 | C | $DJ694^{H}$ | $DJ694^{H}$ | 24-02-07 | 25 | 60 | 93 | 86 | | | | | | | | | | | | |
| 1 | C | $DJ694^{H}$ | $DJ694^{H}$ | 24-02-07 | 25 | 56 | 86 | 83 | | | | | | | | | | | | |
| 1 | A | $DJ694^{H}$ | $w^{1118}$ | 24-02-05 | 25 | 88 | 86 | 100 | 85 | 82 | 100 | 4.31 | 6.08 | 0.00 | - | - | - | - | - | - |
| 1 | A | $DJ694^{H}$ | $w^{1118}$ | 24-02-05 | 11 | 82 | 78 | 100 | | | | | | | | | | | | |
| 1 | B | $DJ694^{H}$ | $w^{1118}$ | 24-02-06 | 25 | 64 | 94 | 100 | 76.00 | 84.88 | 98.55 | 10.33 | 6.81 | 2.90 | - | - | - | - | - | - |
| 1 | B | $DJ694^{H}$ | $w^{1118}$ | 24-02-06 | 25 | 80 | 85 | 94 | | | | | | | | | | | | |
| 1 | B | $DJ694^{H}$ | $w^{1118}$ | 24-02-06 | 25 | 72 | 83 | 100 | | | | | | | | | | | | |
| 1 | B | $DJ694^{H}$ | $w^{1118}$ | 24-02-06 | 25 | 88 | 77 | 100 | | | | | | | | | | | | |
| 1 | C | $DJ694^{H}$ | $w^{1118}$ | 24-02-07 | 25 | 84 | 91 | 90 | 83.00 | 88.35 | 96.00 | 5.03 | 10.59 | 5.05 | - | - | - | - | - | - |
| 1 | C | $DJ694^{H}$ | $w^{1118}$ | 24-02-07 | 25 | 76 | 95 | 95 | | | | | | | | | | | | |
| 1 | C | $DJ694^{H}$ | $w^{1118}$ | 24-02-07 | 25 | 88 | 73 | 100 | | | | | | | | | | | | |
| 1 | C | $DJ694^{H}$ | $w^{1118}$ | 24-02-07 | 25 | 84 | 95 | 100 | | | | | | | | | | | | |
| 1 | A | $DJ694^{L}$ | $DJ694^{L}$ | 24-02-05 | 25 | 56 | 64 | 89 | 69.33 | 86.53 | 86.80 | 14.05 | 19.40 | 9.43 | - | - | - | 0.2954 | 0.8134 | 0.0583 |
| 1 | A | $DJ694^{L}$ | $DJ694^{L}$ | 24-02-05 | 25 | 84 | 95 | 95 | | | | | | | | | | | | |
| 1 | A | $DJ694^{L}$ | $DJ694^{L}$ | 24-02-05 | 25 | 68 | 100 | 77 | | | | | | | | | | | | |
| 1 | B | $DJ694^{L}$ | $DJ694^{L}$ | 24-02-06 | 25 | 44 | 100 | 91 | 55.00 | 87.28 | 97.75 | 12.81 | 12.30 | 4.50 | 0.0042 | 0.2998 | 0.5768 | 0.0347 | 0.8769 | 0.5243 |
| 1 | B | $DJ694^{L}$ | $DJ694^{L}$ | 24-02-06 | 25 | 44 | 91 | 100 | | | | | | | | | | | | |
| 1 | B | $DJ694^{L}$ | $DJ694^{L}$ | 24-02-06 | 25 | 64 | 88 | 100 | | | | | | | | | | | | |
| 1 | B | $DJ694^{L}$ | $DJ694^{L}$ | 24-02-06 | 25 | 68 | 71 | 100 | | | | | | | | | | | | |

Table S1. *DJ694* 's developmental lethality dataset and statistic results

| | | Cross | | Date | | | | | Mean | | | SD | | | P (Homo vs. Wt) | | | P (Homo vs. Hetero) | | |
|---|---|---|---|---|---|---|---|---|---|---|---|---|---|---|---|---|---|---|---|---|
| **Exp** | **Rep** | **♂** | **V♀** | **(YY-MM-DD)** | **# Eggs** | **% L1** | **% P** | **% A** | **L1** | **P** | **A** | **L1** | **P** | **A** | **L1** | **P** | **A** | **L1** | **P** | **A** |
| 1 | C | $DJ694^{L}$ | $DJ694^{L}$ | 24-02-07 | 25 | 52 | 85 | 91 | 60.00 | 86.25 | 95.98 | 7.30 | 6.47 | 4.71 | 0.0215 | 0.0034 | 0.1384 | 0.0145 | 0.1395 | 0.1384 |
| 1 | C | $DJ694^{L}$ | $DJ694^{L}$ | 24-02-07 | 25 | 64 | 88 | 93 | | | | | | | | | | | | |
| 1 | C | $DJ694^{L}$ | $DJ694^{L}$ | 24-02-07 | 25 | 56 | 79 | 100 | | | | | | | | | | | | |
| 1 | C | $DJ694^{L}$ | $DJ694^{L}$ | 24-02-07 | 25 | 68 | 94 | 100 | | | | | | | | | | | | |
| 1 | A | $DJ694^{L}$ | $w^{1118}$ | 24-02-05 | 25 | 68 | 88 | 100 | 80.00 | 84.03 | 98.63 | 10.33 | 6.28 | 2.75 | - | - | - | - | - | - |
| 1 | A | $DJ694^{L}$ | $w^{1118}$ | 24-02-05 | 25 | 92 | 78 | 95 | | | | | | | | | | | | |
| 1 | A | $DJ694^{L}$ | $w^{1118}$ | 24-02-05 | 25 | 84 | 91 | 100 | | | | | | | | | | | | |
| 1 | A | $DJ694^{L}$ | $w^{1118}$ | 24-02-05 | 25 | 76 | 79 | 100 | | | | | | | | | | | | |
| 1 | B | $DJ694^{L}$ | $w^{1118}$ | 24-02-06 | 25 | 76 | 90 | 100 | 76.00 | 88.28 | 95.13 | 8.64 | 1.34 | 6.33 | - | - | - | - | - | - |
| 1 | B | $DJ694^{L}$ | $w^{1118}$ | 24-02-06 | 25 | 72 | 89 | 94 | | | | | | | | | | | | |
| 1 | B | $DJ694^{L}$ | $w^{1118}$ | 24-02-06 | 25 | 68 | 88 | 87 | | | | | | | | | | | | |
| 1 | B | $DJ694^{L}$ | $w^{1118}$ | 24-02-06 | 25 | 88 | 86 | 100 | | | | | | | | | | | | |
| 1 | C | $DJ694^{L}$ | $w^{1118}$ | 24-02-07 | 25 | 84 | 91 | 100 | 77.00 | 76.15 | 100 | 6.83 | 9.94 | 0.00 | - | - | - | - | - | - |
| 1 | C | $DJ694^{L}$ | $w^{1118}$ | 24-02-07 | 25 | 80 | 75 | 100 | | | | | | | | | | | | |
| 1 | C | $DJ694^{L}$ | $w^{1118}$ | 24-02-07 | 25 | 68 | 71 | 100 | | | | | | | | | | | | |
| 1 | C | $DJ694^{L}$ | $w^{1118}$ | 24-02-07 | 25 | 76 | 69 | 100 | | | | | | | | | | | | |
| 1 | A | $DJ694^{O}$ | $DJ694^{O}$ | 24-02-05 | 25 | 56 | 57 | 75 | 56 | 54 | 66 | 0.00 | 5.09 | 12.59 | - | - | - | 0.0413 | 0.0012 | 0.0145 |
| 1 | A | $DJ694^{O}$ | $DJ694^{O}$ | 24-02-05 | 25 | 56 | 50 | 57 | | | | | | | | | | | | |
| 1 | B | $DJ694^{O}$ | $DJ694^{O}$ | 24-02-06 | 25 | 72 | 61 | 100 | 59.00 | 79.58 | 97.93 | 9.45 | 13.27 | 4.15 | 0.0028 | 0.9844 | 0.5323 | 0.4110 | 0.2873 | 0.4150 |
| 1 | B | $DJ694^{O}$ | $DJ694^{O}$ | 24-02-06 | 25 | 52 | 92 | 100 | | | | | | | | | | | | |
| 1 | B | $DJ694^{O}$ | $DJ694^{O}$ | 24-02-06 | 25 | 52 | 85 | 100 | | | | | | | | | | | | |
| 1 | B | $DJ694^{O}$ | $DJ694^{O}$ | 24-02-06 | 25 | 60 | 80 | 92 | | | | | | | | | | | | |
| 1 | C | $DJ694^{O}$ | $DJ694^{O}$ | 24-02-07 | 25 | 36 | 78 | 100 | 43.00 | 83.70 | 92.75 | 8.25 | 4.58 | 9.50 | 0.0017 | 0.0035 | 0.1778 | 0.0544 | 0.2062 | 0.1778 |
| 1 | C | $DJ694^{O}$ | $DJ694^{O}$ | 24-02-07 | 25 | 48 | 83 | 80 | | | | | | | | | | | | |
| 1 | C | $DJ694^{O}$ | $DJ694^{O}$ | 24-02-07 | 25 | 52 | 85 | 91 | | | | | | | | | | | | |
| 1 | C | $DJ694^{O}$ | $DJ694^{O}$ | 24-02-07 | 25 | 36 | 89 | 100 | | | | | | | | | | | | |
| 1 | A | $DJ694^{O}$ | $w^{1118}$ | 24-02-05 | 25 | 80 | 95 | 100 | 77.33 | 93.23 | 100 | 8.33 | 2.40 | 0.00 | - | - | - | - | - | - |
| 1 | A | $DJ694^{O}$ | $w^{1118}$ | 24-02-05 | 25 | 84 | 91 | 100 | | | | | | | | | | | | |
| 1 | A | $DJ694^{O}$ | $w^{1118}$ | 24-02-05 | 25 | 68 | 94 | 100 | | | | | | | | | | | | |

Table S1. *DJ694*'s developmental lethality dataset and statistic results

| | | Cross | | Date | | | | | Mean | | | SD | | | P (Homo vs. Wt) | | | P (Homo vs. Hetero) | | |
|---|---|---|---|---|---|---|---|---|---|---|---|---|---|---|---|---|---|---|---|---|
| Exp | Rep | ♂ | V♀ | (YY-MM-DD) | # Eggs | % L1 | % P | % A | L1 | P | A | L1 | P | A | L1 | P | A | L1 | P | A |
| 1 | B | $DJ694^{O}$ | $w^{1118}$ | 24-02-06 | 25 | 76 | 100 | 84 | 67.00 | 90.45 | 94.18 | 15.45 | 13.07 | 7.50 | - | - | - | - | - | - |
| 1 | B | $DJ694^{O}$ | $w^{1118}$ | 24-02-06 | 25 | 76 | 90 | 100 | | | | | | | | | | | | |
| 1 | B | $DJ694^{O}$ | $w^{1118}$ | 24-02-06 | 25 | 72 | 72 | 92 | | | | | | | | | | | | |
| 1 | B | $DJ694^{O}$ | $w^{1118}$ | 24-02-06 | 25 | 44 | 100 | 100 | | | | | | | | | | | | |
| 1 | C | $DJ694^{O}$ | $w^{1118}$ | 24-02-07 | 25 | 88 | 86 | 100 | 66.00 | 75.70 | 100 | 17.44 | 10.32 | 0.00 | - | - | - | - | - | - |
| 1 | C | $DJ694^{O}$ | $w^{1118}$ | 24-02-07 | 25 | 52 | 77 | 100 | | | | | | | | | | | | |
| 1 | C | $DJ694^{O}$ | $w^{1118}$ | 24-02-07 | 25 | 72 | 78 | 100 | | | | | | | | | | | | |
| 1 | C | $DJ694^{O}$ | $w^{1118}$ | 24-02-07 | 25 | 52 | 62 | 100 | | | | | | | | | | | | |
| 1 | A | $w^{1118}$ | $w^{1118}$ | 24-02-05 | 16 | 94 | 93 | 100 | - | - | - | - | - | - | - | - | - | - | - | - |
| 1 | B | $w^{1118}$ | $w^{1118}$ | 24-02-06 | 25 | 96 | 79 | 90 | 88.00 | 79.43 | 95.73 | 7.30 | 6.34 | 5.19 | - | - | - | - | - | - |
| 1 | B | $w^{1118}$ | $w^{1118}$ | 24-02-06 | 25 | 80 | 80 | 100 | | | | | | | | | | | | |
| 1 | B | $w^{1118}$ | $w^{1118}$ | 24-02-06 | 25 | 92 | 87 | 100 | | | | | | | | | | | | |
| 1 | B | $w^{1118}$ | $w^{1118}$ | 24-02-06 | 25 | 84 | 72 | 93 | | | | | | | | | | | | |
| 1 | C | $w^{1118}$ | $w^{1118}$ | 24-02-07 | 25 | 88 | 59 | 100 | 81.00 | 61.15 | 100 | 11.49 | 8.52 | 0.00 | - | - | - | - | - | - |
| 1 | C | $w^{1118}$ | $w^{1118}$ | 24-02-07 | 25 | 88 | 50 | 100 | | | | | | | | | | | | |
| 1 | C | $w^{1118}$ | $w^{1118}$ | 24-02-07 | 25 | 84 | 67 | 100 | | | | | | | | | | | | |
| 1 | C | $w^{1118}$ | $w^{1118}$ | 24-02-07 | 25 | 64 | 69 | 100 | | | | | | | | | | | | |
| 2 | A | $DJ694^{H}$ | $DJ694^{H}$ | 24-02-26 | 25 | 84 | 91 | 90 | 84.00 | 87.10 | 91.55 | 3.27 | 11.86 | 5.82 | 0.1140 | 0.2273 | 0.9137 | 0.6202 | 0.9150 | 0.3369 |
| 2 | A | $DJ694^{H}$ | $DJ694^{H}$ | 24-02-26 | 25 | 84 | 72 | 87 | | | | | | | | | | | | |
| 2 | A | $DJ694^{H}$ | $DJ694^{H}$ | 24-02-26 | 25 | 80 | 100 | 90 | | | | | | | | | | | | |
| 2 | A | $DJ694^{H}$ | $DJ694^{H}$ | 24-02-26 | 25 | 88 | 86 | 100 | | | | | | | | | | | | |
| 2 | B | $DJ694^{H}$ | $DJ694^{H}$ | 24-02-27 | 25 | 84 | 95 | 90 | 86.00 | 94.45 | 93.88 | 5.16 | 5.42 | 4.66 | 0.4772 | 0.0205 | 0.1210 | 0.0170 | 0.2699 | 0.0710 |
| 2 | B | $DJ694^{H}$ | $DJ694^{H}$ | 24-02-27 | 25 | 88 | 96 | 91 | | | | | | | | | | | | |
| 2 | B | $DJ694^{H}$ | $DJ694^{H}$ | 24-02-27 | 25 | 80 | 100 | 95 | | | | | | | | | | | | |
| 2 | B | $DJ694^{H}$ | $DJ694^{H}$ | 24-02-27 | 25 | 92 | 87 | 100 | | | | | | | | | | | | |
| 2 | A | $DJ694^{H}$ | $w^{1118}$ | 24-02-26 | 25 | 84 | 100 | 100 | 86.00 | 86.25 | 100 | 6.93 | 9.63 | 0.00 | - | - | - | - | - | - |
| 2 | A | $DJ694^{H}$ | $w^{1118}$ | 24-02-26 | 25 | 80 | 80 | 100 | | | | | | | | | | | | |
| 2 | A | $DJ694^{H}$ | $w^{1118}$ | 24-02-26 | 25 | 96 | 79 | 100 | | | | | | | | | | | | |
| 2 | A | $DJ694^{H}$ | $w^{1118}$ | 24-02-26 | 25 | 84 | 86 | 100 | | | | | | | | | | | | |

Table S1. *DJ694*'s developmental lethality dataset and statistic results

| | | Cross | | Date | | | | | Mean | | | SD | | | P (Homo vs. Wt) | | | P (Homo vs. Hetero) | | |
|---|---|---|---|---|---|---|---|---|---|---|---|---|---|---|---|---|---|---|---|---|
| Exp | Rep | ♂ | V♀ | (YY-MM-DD) | # Eggs | % L1 | % P | % A | L1 | P | A | L1 | P | A | L1 | P | A | L1 | P | A |
| 2 | B | $DJ694^{H}$ | $w^{1118}$ | 24-02-27 | 25 | 72 | 89 | 94 | 76.00 | 90.73 | 98.45 | 3.27 | 2.86 | 3.10 | - | - | - | - | - | - |
| 2 | B | $DJ694^{H}$ | $w^{1118}$ | 24-02-27 | 25 | 76 | 90 | 100 | | | | | | | | | | | | |
| 2 | B | $DJ694^{H}$ | $w^{1118}$ | 24-02-27 | 25 | 80 | 95 | 100 | | | | | | | | | | | | |
| 2 | B | $DJ694^{H}$ | $w^{1118}$ | 24-02-27 | 25 | 76 | 90 | 100 | | | | | | | | | | | | |
| 2 | A | $DJ694^{L}$ | $DJ694^{L}$ | 24-02-26 | 25 | 88 | 64 | 100 | 87.00 | 74.60 | 93.38 | 5.03 | 9.88 | 9.93 | 0.3822 | 0.9267 | 0.3365 | 0.6932 | 0.0127 | 0.0065 |
| 2 | A | $DJ694^{L}$ | $DJ694^{L}$ | 24-02-26 | 25 | 80 | 70 | 100 | | | | | | | | | | | | |
| 2 | A | $DJ694^{L}$ | $DJ694^{L}$ | 24-02-26 | 25 | 92 | 78 | 95 | | | | | | | | | | | | |
| 2 | A | $DJ694^{L}$ | $DJ694^{L}$ | 24-02-26 | 25 | 88 | 86 | 79 | | | | | | | | | | | | |
| 2 | B | $DJ694^{L}$ | $DJ694^{L}$ | 24-02-27 | 25 | 60 | 80 | 92 | 62.00 | 92.38 | 96.48 | 10.07 | 9.62 | 4.20 | 0.0037 | 0.0544 | 0.1622 | 0.0361 | 0.6846 | 0.0490 |
| 2 | B | $DJ694^{L}$ | $DJ694^{L}$ | 24-02-27 | 25 | 52 | 100 | 100 | | | | | | | | | | | | |
| 2 | B | $DJ694^{L}$ | $DJ694^{L}$ | 24-02-27 | 25 | 76 | 90 | 94 | | | | | | | | | | | | |
| 2 | B | $DJ694^{L}$ | $DJ694^{L}$ | 24-02-27 | 25 | 60 | 100 | 100 | | | | | | | | | | | | |
| 2 | A | $DJ694^{L}$ | $w^{1118}$ | 24-02-26 | 25 | 88 | 91 | 95 | 89.00 | 92.25 | 97.50 | 8.25 | 1.89 | 2.89 | - | - | - | - | - | - |
| 2 | A | $DJ694^{L}$ | $w^{1118}$ | 24-02-26 | 25 | 100 | 92 | 100 | | | | | | | | | | | | |
| 2 | A | $DJ694^{L}$ | $w^{1118}$ | 24-02-26 | 25 | 80 | 95 | 100 | | | | | | | | | | | | |
| 2 | A | $DJ694^{L}$ | $w^{1118}$ | 24-02-26 | 25 | 88 | 91 | 95 | | | | | | | | | | | | |
| 2 | B | $DJ694^{L}$ | $w^{1118}$ | 24-02-27 | 25 | 68 | 82 | 100 | 83.00 | 90.03 | 100 | 11.94 | 5.37 | 0.00 | - | - | - | - | - | - |
| 2 | B | $DJ694^{L}$ | $w^{1118}$ | 24-02-27 | 25 | 96 | 92 | 100 | | | | | | | | | | | | |
| 2 | B | $DJ694^{L}$ | $w^{1118}$ | 24-02-27 | 25 | 88 | 91 | 100 | | | | | | | | | | | | |
| 2 | B | $DJ694^{L}$ | $w^{1118}$ | 24-02-27 | 25 | 80 | 95 | 100 | | | | | | | | | | | | |
| 2 | A | $DJ694^{O}$ | $DJ694^{O}$ | 24-02-26 | 25 | 68 | 47 | 100 | 67.00 | 47.08 | 100 | 8.25 | 8.29 | 0.00 | 0.0042 | 0.0099 | 0.0021 | 0.4697 | 5E-05 | 0.0015 |
| 2 | A | $DJ694^{O}$ | $DJ694^{O}$ | 24-02-26 | 25 | 68 | 47 | 100 | | | | | | | | | | | | |
| 2 | A | $DJ694^{O}$ | $DJ694^{O}$ | 24-02-26 | 25 | 56 | 57 | 100 | | | | | | | | | | | | |
| 2 | A | $DJ694^{O}$ | $DJ694^{O}$ | 24-02-26 | 25 | 76 | 37 | 100 | | | | | | | | | | | | |
| 2 | B | $DJ694^{O}$ | $DJ694^{O}$ | 24-02-27 | 25 | 48 | 92 | 100 | 59.00 | 77.95 | 97.75 | 10.52 | 12.40 | 4.50 | 0.0026 | 0.6837 | 0.0073 | 0.5945 | 0.0495 | 0.0941 |
| 2 | B | $DJ694^{O}$ | $DJ694^{O}$ | 24-02-27 | 25 | 52 | 85 | 91 | | | | | | | | | | | | |
| 2 | B | $DJ694^{O}$ | $DJ694^{O}$ | 24-02-27 | 25 | 68 | 71 | 100 | | | | | | | | | | | | |
| 2 | B | $DJ694^{O}$ | $DJ694^{O}$ | 24-02-27 | 25 | 68 | 65 | 100 | | | | | | | | | | | | |
| 2 | A | $DJ694^{O}$ | $w^{1118}$ | 24-02-26 | 25 | 64 | 100 | 100 | 74.00 | 96.35 | 100 | 16.17 | 4.96 | 0.00 | - | - | - | - | - | - |
| 2 | A | $DJ694^{O}$ | $w^{1118}$ | 24-02-26 | 25 | 76 | 90 | 100 | | | | | | | | | | | | |
| 2 | A | $DJ694^{O}$ | $w^{1118}$ | 24-02-26 | 25 | 96 | 96 | 100 | | | | | | | | | | | | |
| 2 | A | $DJ694^{O}$ | $w^{1118}$ | 24-02-26 | 25 | 60 | 100 | 100 | | | | | | | | | | | | |

| | | Cross | | Date | | | | | Mean | | | SD | | | P (Homo vs. Wt) | | | P (Homo vs. Hetero) | | |
|---|---|---|---|---|---|---|---|---|---|---|---|---|---|---|---|---|---|---|---|---|
| Exp | Rep | ♂ | V♀ | (YY-MM-DD) | # Eggs | % L1 | % P | % A | L1 | P | A | L1 | P | A | L1 | P | A | L1 | P | A |
| 2 | B | $DJ694^{O}$ | $w^{1118}$ | 24-02-27 | 25 | 72 | 89 | 94 | 65.00 | 94.85 | 96.90 | 18.58 | 5.98 | 3.58 | - | - | - | - | - | - |
| 2 | B | $DJ694^{O}$ | $w^{1118}$ | 24-02-27 | 25 | 40 | 100 | 100 | | | | | | | | | | | | |
| 2 | B | $DJ694^{O}$ | $w^{1118}$ | 24-02-27 | 25 | 64 | 100 | 94 | | | | | | | | | | | | |
| 2 | B | $DJ694^{O}$ | $w^{1118}$ | 24-02-27 | 25 | 84 | 91 | 100 | | | | | | | | | | | | |
| 2 | A | $w^{1118}$ | $w^{1118}$ | 24-02-26 | 25 | 100 | 80 | 95 | 91.00 | 75.38 | 98.75 | 6.83 | 12.79 | 2.50 | - | - | - | - | - | - |
| 2 | A | $w^{1118}$ | $w^{1118}$ | 24-02-26 | 25 | 84 | 57 | 100 | | | | | | | | | | | | |
| 2 | A | $w^{1118}$ | $w^{1118}$ | 24-02-26 | 25 | 92 | 87 | 100 | | | | | | | | | | | | |
| 2 | A | $w^{1118}$ | $w^{1118}$ | 24-02-26 | 25 | 88 | 77 | 100 | | | | | | | | | | | | |
| 2 | B | $w^{1118}$ | $w^{1118}$ | 24-02-27 | 25 | 96 | 67 | 100 | 89.00 | 74.30 | 100 | 6.00 | 11.72 | 0.00 | - | - | - | - | - | - |
| 2 | B | $w^{1118}$ | $w^{1118}$ | 24-02-27 | 25 | 84 | 86 | 100 | | | | | | | | | | | | |
| 2 | B | $w^{1118}$ | $w^{1118}$ | 24-02-27 | 25 | 84 | 62 | 100 | | | | | | | | | | | | |
| 2 | B | $w^{1118}$ | $w^{1118}$ | 24-02-27 | 25 | 92 | 83 | 100 | | | | | | | | | | | | |
| 3 | A | $DJ694^{H}$ | $DJ694^{H}$ | 24-03-07 | 25 | 80 | 85 | 88 | 82 | 88 | 89 | 2.83 | 3.89 | 0.85 | 0.0432 | 0.1691 | 0.0001 | 0.6741 | 0.2262 | 0.0715 |
| 3 | A | $DJ694^{H}$ | $DJ694^{H}$ | 24-03-07 | 25 | 84 | 91 | 90 | | | | | | | | | | | | |
| 3 | A | $DJ694^{H}$ | $w^{1118}$ | 24-03-07 | 25 | 84 | 100 | 100 | 80.00 | 94.73 | 97.63 | 5.66 | 6.11 | 4.75 | - | - | - | - | - | - |
| 3 | A | $DJ694^{H}$ | $w^{1118}$ | 24-03-07 | 25 | 84 | 100 | 91 | | | | | | | | | | | | |
| 3 | A | $DJ694^{H}$ | $w^{1118}$ | 24-03-07 | 25 | 80 | 90 | 100 | | | | | | | | | | | | |
| 3 | A | $DJ694^{H}$ | $w^{1118}$ | 24-03-07 | 25 | 72 | 89 | 100 | | | | | | | | | | | | |
| 3 | A | $DJ694^{L}$ | $DJ694^{L}$ | 24-03-07 | 25 | 36 | 89 | 88 | 36.00 | 89.63 | 95.83 | 4.00 | 10.02 | 7.22 | 8E-05 | 0.1202 | 0.3739 | 4E-06 | 0.7867 | 0.8597 |
| 3 | A | $DJ694^{L}$ | $DJ694^{L}$ | 24-03-07 | 25 | 40 | 80 | 100 | | | | | | | | | | | | |
| 3 | A | $DJ694^{L}$ | $DJ694^{L}$ | 24-03-07 | 25 | 32 | 100 | 100 | | | | | | | | | | | | |
| 3 | A | $DJ694^{L}$ | $w^{1118}$ | 24-03-07 | 25 | 84 | 100 | 100 | 85.00 | 87.28 | 94.45 | 2.00 | 11.31 | 11.10 | - | - | - | - | - | - |
| 3 | A | $DJ694^{L}$ | $w^{1118}$ | 24-03-07 | 25 | 84 | 91 | 100 | | | | | | | | | | | | |
| 3 | A | $DJ694^{L}$ | $w^{1118}$ | 24-03-07 | 25 | 84 | 86 | 78 | | | | | | | | | | | | |
| 3 | A | $DJ694^{L}$ | $w^{1118}$ | 24-03-07 | 25 | 88 | 73 | 100 | | | | | | | | | | | | |
| 3 | A | $DJ694^{O}$ | $DJ694^{O}$ | 24-03-07 | 25 | 36 | 56 | 100 | 31 | 61 | 100 | 7.78 | 7.85 | 0.00 | 0.0012 | 0.2730 | 1 | 0.0244 | 0.0127 | 0.3631 |
| 3 | A | $DJ694^{O}$ | $DJ694^{O}$ | 24-03-07 | 12 | 25 | 67 | 100 | | | | | | | | | | | | |
| 3 | A | $DJ694^{O}$ | $w^{1118}$ | 24-03-07 | 25 | 48 | 92 | 100 | 68.00 | 85.83 | 95.43 | 13.47 | 6.18 | 5.95 | - | - | - | - | - | - |
| 3 | A | $DJ694^{O}$ | $w^{1118}$ | 24-03-07 | 25 | 72 | 78 | 100 | | | | | | | | | | | | |
| 3 | A | $DJ694^{O}$ | $w^{1118}$ | 24-03-07 | 25 | 76 | 84 | 88 | | | | | | | | | | | | |
| 3 | A | $DJ694^{O}$ | $w^{1118}$ | 24-03-07 | 25 | 76 | 90 | 94 | | | | | | | | | | | | |

| | | Cross | | Date | | | | | Mean | | | SD | | | P (Homo vs. Wt) | | | P (Homo vs. Hetero) | | |
|---|---|---|---|---|---|---|---|---|---|---|---|---|---|---|---|---|---|---|---|---|
| Exp | Rep | ♂ | V♀ | (YY-MM-DD) | # Eggs | % L1 | % P | % A | L1 | P | A | L1 | P | A | L1 | P | A | L1 | P | A |
| 3 | A | $w^{1118}$ | $w^{1118}$ | 24-03-07 | 25 | 92 | 61 | 100 | 94.67 | 73.07 | 100 | 4.62 | 10.57 | 0.00 | - | - | - | - | - | - |
| 3 | A | $w^{1118}$ | $w^{1118}$ | 24-03-07 | 25 | 100 | 80 | 100 | | | | | | | | | | | | |
| 3 | A | $w^{1118}$ | $w^{1118}$ | 24-03-07 | 25 | 92 | 78 | 100 | | | | | | | | | | | | |
| 4 | A | $DJ694^{H}$ | $DJ694^{H}$ | 24-03-28 | 25 | 84 | 100 | 91 | 83.00 | 91.33 | 96.50 | 5.03 | 10.11 | 4.53 | 0.3464 | 0.0563 | 0.1731 | 0.2483 | 0.5991 | 0.3609 |
| 4 | A | $DJ694^{H}$ | $DJ694^{H}$ | 24-03-28 | 25 | 88 | 100 | 96 | | | | | | | | | | | | |
| 4 | A | $DJ694^{H}$ | $DJ694^{H}$ | 24-03-28 | 25 | 84 | 81 | 100 | | | | | | | | | | | | |
| 4 | A | $DJ694^{H}$ | $DJ694^{H}$ | 24-03-28 | 25 | 76 | 84 | 100 | | | | | | | | | | | | |
| 4 | B | $DJ694^{H}$ | $DJ694^{H}$ | 24-03-29 | 25 | 96 | 100 | 92 | 83.00 | 93.60 | 97.93 | 11.02 | 8.01 | 4.15 | 0.1534 | 0.1009 | 0.3559 | 0.6394 | 0.9091 | 0.7188 |
| 4 | B | $DJ694^{H}$ | $DJ694^{H}$ | 24-03-29 | 25 | 76 | 100 | 100 | | | | | | | | | | | | |
| 4 | B | $DJ694^{H}$ | $DJ694^{H}$ | 24-03-29 | 25 | 88 | 91 | 100 | | | | | | | | | | | | |
| 4 | B | $DJ694^{H}$ | $DJ694^{H}$ | 24-03-29 | 25 | 72 | 83 | 100 | | | | | | | | | | | | |
| 4 | C | $DJ694^{H}$ | $DJ694^{H}$ | 24-03-30 | 25 | 60 | 87 | 100 | 71.00 | 89.75 | 100 | 11.02 | 3.88 | 0.00 | 0.0728 | 0.0073 | 1 | 0.1738 | 0.1903 | 0.3559 |
| 4 | C | $DJ694^{H}$ | $DJ694^{H}$ | 24-03-30 | 25 | 64 | 88 | 100 | | | | | | | | | | | | |
| 4 | C | $DJ694^{H}$ | $DJ694^{H}$ | 24-03-30 | 25 | 84 | 95 | 100 | | | | | | | | | | | | |
| 4 | C | $DJ694^{H}$ | $DJ694^{H}$ | 24-03-30 | 25 | 76 | 90 | 100 | | | | | | | | | | | | |
| 4 | D | $DJ694^{H}$ | $DJ694^{H}$ | 24-04-01 | 25 | 88 | 91 | 100 | 76.00 | 85.08 | 95.53 | 11.78 | 4.53 | 5.63 | 0.3997 | 0.7558 | 0.9358 | 0.3835 | 0.0385 | 0.3766 |
| 4 | D | $DJ694^{H}$ | $DJ694^{H}$ | 24-04-01 | 25 | 76 | 84 | 94 | | | | | | | | | | | | |
| 4 | D | $DJ694^{H}$ | $DJ694^{H}$ | 24-04-01 | 25 | 60 | 80 | 100 | | | | | | | | | | | | |
| 4 | D | $DJ694^{H}$ | $DJ694^{H}$ | 24-04-01 | 25 | 80 | 85 | 88 | | | | | | | | | | | | |
| 4 | A | $DJ694^{H}$ | $w^{1118}$ | 24-03-28 | 25 | 80 | 85 | 100 | 78 | 87 | 100 | 3.54 | 2.76 | 0.00 | - | - | - | - | - | - |
| 4 | A | $DJ694^{H}$ | $w^{1118}$ | 24-03-28 | 12 | 75 | 89 | 100 | | | | | | | | | | | | |
| 4 | B | $DJ694^{H}$ | $w^{1118}$ | 24-03-29 | 25 | 80 | 95 | 100 | 86.00 | 93.10 | 98.83 | 5.16 | 2.52 | 2.35 | - | - | - | - | - | - |
| 4 | B | $DJ694^{H}$ | $w^{1118}$ | 24-03-29 | 25 | 84 | 91 | 100 | | | | | | | | | | | | |
| 4 | B | $DJ694^{H}$ | $w^{1118}$ | 24-03-29 | 25 | 92 | 91 | 100 | | | | | | | | | | | | |
| 4 | B | $DJ694^{H}$ | $w^{1118}$ | 24-03-29 | 25 | 88 | 96 | 95 | | | | | | | | | | | | |
| 4 | C | $DJ694^{H}$ | $w^{1118}$ | 24-03-30 | 25 | 72 | 83 | 100 | 81.00 | 82.55 | 98.75 | 6.83 | 8.95 | 2.50 | - | - | - | - | - | - |
| 4 | C | $DJ694^{H}$ | $w^{1118}$ | 24-03-30 | 25 | 80 | 70 | 100 | | | | | | | | | | | | |
| 4 | C | $DJ694^{H}$ | $w^{1118}$ | 24-03-30 | 25 | 88 | 91 | 95 | | | | | | | | | | | | |
| 4 | C | $DJ694^{H}$ | $w^{1118}$ | 24-03-30 | 25 | 84 | 86 | 100 | | | | | | | | | | | | |

| | | Cross | | Date | | | | | Mean | | | SD | | | P (Homo vs. Wt) | | | P (Homo vs. Hetero) | | |
|---|---|---|---|---|---|---|---|---|---|---|---|---|---|---|---|---|---|---|---|---|
| Exp | Rep | ♂ | V♀ | (YY-MM-DD) | # Eggs | % L1 | % P | % A | L1 | P | A | L1 | P | A | L1 | P | A | L1 | P | A |
| 4 | D | *DJ694*$^{H}$ | $w^{1118}$ | 24-04-01 | 25 | 72 | 89 | 100 | 85.00 | 91.78 | 98.55 | 15.10 | 2.29 | 2.90 | - | - | - | - | - | - |
| 4 | D | *DJ694*$^{H}$ | $w^{1118}$ | 24-04-01 | 25 | 100 | 92 | 100 | | | | | | | | | | | | |
| 4 | D | *DJ694*$^{H}$ | $w^{1118}$ | 24-04-01 | 25 | 72 | 95 | 94 | | | | | | | | | | | | |
| 4 | D | *DJ694*$^{H}$ | $w^{1118}$ | 24-04-01 | 25 | 96 | 92 | 100 | | | | | | | | | | | | |
| 4 | A | *DJ694*$^{L}$ | *DJ694*$^{L}$ | 24-03-28 | 25 | 76 | 90 | 94 | 79.00 | 86.28 | 94.40 | 3.83 | 8.38 | 4.53 | 0.0656 | 0.1282 | 0.0484 | 0.3169 | 0.9511 | 0.0484 |
| 4 | A | *DJ694*$^{L}$ | *DJ694*$^{L}$ | 24-03-28 | 25 | 84 | 86 | 89 | | | | | | | | | | | | |
| 4 | A | *DJ694*$^{L}$ | *DJ694*$^{L}$ | 24-03-28 | 25 | 76 | 95 | 95 | | | | | | | | | | | | |
| 4 | A | *DJ694*$^{L}$ | *DJ694*$^{L}$ | 24-03-28 | 25 | 80 | 75 | 100 | | | | | | | | | | | | |
| 4 | B | *DJ694*$^{L}$ | *DJ694*$^{L}$ | 24-03-29 | 25 | 92 | 96 | 96 | 85.33 | 98.57 | 96.77 | 8.33 | 2.48 | 2.82 | 0.1583 | 0.0220 | 0.0638 | 0.3868 | 0.2019 | 0.8952 |
| 4 | B | *DJ694*$^{L}$ | *DJ694*$^{L}$ | 24-03-29 | 25 | 76 | 100 | 95 | | | | | | | | | | | | |
| 4 | B | *DJ694*$^{L}$ | *DJ694*$^{L}$ | 24-03-29 | 25 | 88 | 100 | 100 | | | | | | | | | | | | |
| 4 | C | *DJ694*$^{L}$ | *DJ694*$^{L}$ | 24-03-30 | 25 | 76 | 84 | 100 | 77.00 | 91.68 | 97.10 | 8.25 | 9.64 | 3.51 | 0.1755 | 0.0158 | 0.1498 | - | - | - |
| 4 | C | *DJ694*$^{L}$ | *DJ694*$^{L}$ | 24-03-30 | 25 | 88 | 100 | 96 | | | | | | | | | | | | |
| 4 | C | *DJ694*$^{L}$ | *DJ694*$^{L}$ | 24-03-30 | 25 | 68 | 82 | 93 | | | | | | | | | | | | |
| 4 | C | *DJ694*$^{L}$ | *DJ694*$^{L}$ | 24-03-30 | 25 | 76 | 100 | 100 | | | | | | | | | | | | |
| 4 | D | *DJ694*$^{L}$ | *DJ694*$^{L}$ | 24-04-01 | 25 | 68 | 94 | 100 | 73.00 | 93.43 | 95.70 | 6.00 | 4.86 | 5.57 | 0.1272 | 0.1216 | 0.9702 | - | - | - |
| 4 | D | *DJ694*$^{L}$ | *DJ694*$^{L}$ | 24-04-01 | 25 | 68 | 100 | 100 | | | | | | | | | | | | |
| 4 | D | *DJ694*$^{L}$ | *DJ694*$^{L}$ | 24-04-01 | 25 | 76 | 90 | 88 | | | | | | | | | | | | |
| 4 | D | *DJ694*$^{L}$ | *DJ694*$^{L}$ | 24-04-01 | 25 | 80 | 90 | 95 | | | | | | | | | | | | |
| 4 | A | *DJ694*$^{L}$ | $w^{1118}$ | 24-03-28 | 25 | 76 | 90 | 17 | 71.30 | 86.88 | 14.00 | 13.58 | 16.81 | 6.06 | - | - | - | - | - | - |
| 4 | A | *DJ694*$^{L}$ | $w^{1118}$ | 24-03-28 | 25 | 88 | 96 | 21 | | | | | | | | | | | | |
| 4 | A | *DJ694*$^{L}$ | $w^{1118}$ | 24-03-28 | 25 | 64 | 63 | 10 | | | | | | | | | | | | |
| 4 | A | *DJ694*$^{L}$ | $w^{1118}$ | 24-03-28 | 14 | 57 | 100 | 8 | | | | | | | | | | | | |
| 4 | B | *DJ694*$^{L}$ | $w^{1118}$ | 24-03-29 | 25 | 76 | 95 | 18 | 77.00 | 88.45 | 16.25 | 13.22 | 11.47 | 1.26 | - | - | - | - | - | - |
| 4 | B | *DJ694*$^{L}$ | $w^{1118}$ | 24-03-29 | 25 | 92 | 74 | 16 | | | | | | | | | | | | |
| 4 | B | *DJ694*$^{L}$ | $w^{1118}$ | 24-03-29 | 25 | 60 | 100 | 15 | | | | | | | | | | | | |
| 4 | B | *DJ694*$^{L}$ | $w^{1118}$ | 24-03-29 | 25 | 80 | 85 | 16 | | | | | | | | | | | | |
| 4 | A | *DJ694*$^{O}$ | *DJ694*$^{O}$ | 24-03-28 | 25 | 40 | 80 | 100 | 55.00 | 65.95 | 94.20 | 11.49 | 24.86 | 7.84 | 0.0026 | 0.4813 | 0.1896 | 0.3961 | 0.3734 | 0.7319 |
| 4 | A | *DJ694*$^{O}$ | *DJ694*$^{O}$ | 24-03-28 | 25 | 64 | 44 | 100 | | | | | | | | | | | | |
| 4 | A | *DJ694*$^{O}$ | *DJ694*$^{O}$ | 24-03-28 | 25 | 52 | 46 | 83 | | | | | | | | | | | | |
| 4 | A | *DJ694*$^{O}$ | *DJ694*$^{O}$ | 24-03-28 | 25 | 64 | 94 | 93 | | | | | | | | | | | | |

Table S1. *DJ694*'s developmental lethality dataset and statistic results

| Exp | Rep | Cross ♂ | Cross V♀ | Date (YY-MM-DD) | # Eggs | % L1 | % P | % A | Mean L1 | Mean P | Mean A | SD L1 | SD P | SD A | P (Homo vs. Wt) L1 | P (Homo vs. Wt) P | P (Homo vs. Wt) A | P (Homo vs. Hetero) L1 | P (Homo vs. Hetero) P | P (Homo vs. Hetero) A |
|---|---|---|---|---|---|---|---|---|---|---|---|---|---|---|---|---|---|---|---|---|
| 4 | B | $DJ694^{O}$ | $DJ694^{O}$ | 24-03-29 | 25 | 28 | 72 | 80 | 46.00 | 73.38 | 92.23 | 16.81 | 10.77 | 9.69 | 0.0016 | 0.2139 | 0.1595 | 0.1239 | 0.1542 | 0.1595 |
| 4 | B | $DJ694^{O}$ | $DJ694^{O}$ | 24-03-29 | 25 | 36 | 89 | 100 | | | | | | | | | | | | |
| 4 | B | $DJ694^{O}$ | $DJ694^{O}$ | 24-03-29 | 25 | 64 | 69 | 100 | | | | | | | | | | | | |
| 4 | B | $DJ694^{O}$ | $DJ694^{O}$ | 24-03-29 | 25 | 56 | 64 | 89 | | | | | | | | | | | | |
| 4 | C | $DJ694^{O}$ | $DJ694^{O}$ | 24-03-30 | 25 | 36 | 89 | 100 | 43.00 | 83.28 | 97.23 | 9.45 | 12.69 | 5.55 | 0.0005 | 0.1218 | 0.3559 | 0.0014 | 0.6936 | 0.6841 |
| 4 | C | $DJ694^{O}$ | $DJ694^{O}$ | 24-03-30 | 25 | 56 | 64 | 89 | | | | | | | | | | | | |
| 4 | C | $DJ694^{O}$ | $DJ694^{O}$ | 24-03-30 | 25 | 36 | 89 | 100 | | | | | | | | | | | | |
| 4 | C | $DJ694^{O}$ | $DJ694^{O}$ | 24-03-30 | 25 | 44 | 91 | 100 | | | | | | | | | | | | |
| 4 | D | $DJ694^{O}$ | $DJ694^{O}$ | 24-04-01 | 25 | 32 | 63 | 100 | - | - | - | - | - | - | - | - | - | - | - | - |
| 4 | A | $DJ694^{O}$ | $w^{1118}$ | 24-03-28 | 25 | 56 | 79 | 100 | 63.00 | 79.75 | 95.85 | 13.22 | 14.36 | 4.79 | - | - | - | - | - | - |
| 4 | A | $DJ694^{O}$ | $w^{1118}$ | 24-03-28 | 25 | 48 | 100 | 92 | | | | | | | | | | | | |
| 4 | A | $DJ694^{O}$ | $w^{1118}$ | 24-03-28 | 25 | 72 | 67 | 92 | | | | | | | | | | | | |
| 4 | A | $DJ694^{O}$ | $w^{1118}$ | 24-03-28 | 25 | 76 | 74 | 100 | | | | | | | | | | | | |
| 4 | B | $DJ694^{O}$ | $w^{1118}$ | 24-03-29 | 25 | 60 | 73 | 100 | 63.00 | 85.88 | 100 | 8.87 | 10.92 | 0.00 | - | - | - | - | - | - |
| 4 | B | $DJ694^{O}$ | $w^{1118}$ | 24-03-29 | 25 | 56 | 86 | 100 | | | | | | | | | | | | |
| 4 | B | $DJ694^{O}$ | $w^{1118}$ | 24-03-29 | 25 | 60 | 100 | 100 | | | | | | | | | | | | |
| 4 | B | $DJ694^{O}$ | $w^{1118}$ | 24-03-29 | 25 | 76 | 84 | 100 | | | | | | | | | | | | |
| 4 | C | $DJ694^{O}$ | $w^{1118}$ | 24-03-30 | 25 | 72 | 83 | 100 | 73.00 | 86.55 | 94.65 | 5.03 | 9.48 | 10.70 | - | - | - | - | - | - |
| 4 | C | $DJ694^{O}$ | $w^{1118}$ | 24-03-30 | 25 | 80 | 85 | 100 | | | | | | | | | | | | |
| 4 | C | $DJ694^{O}$ | $w^{1118}$ | 24-03-30 | 25 | 68 | 100 | 100 | | | | | | | | | | | | |
| 4 | C | $DJ694^{O}$ | $w^{1118}$ | 24-03-30 | 25 | 72 | 78 | 79 | | | | | | | | | | | | |
| 4 | D | $DJ694^{O}$ | $w^{1118}$ | 24-04-01 | 25 | 60 | 93 | 100 | 64.00 | 92.60 | 100 | 8.64 | 5.51 | 0.00 | - | - | - | - | - | - |
| 4 | D | $DJ694^{O}$ | $w^{1118}$ | 24-04-01 | 25 | 76 | 90 | 100 | | | | | | | | | | | | |
| 4 | D | $DJ694^{O}$ | $w^{1118}$ | 24-04-01 | 25 | 64 | 88 | 100 | | | | | | | | | | | | |
| 4 | D | $DJ694^{O}$ | $w^{1118}$ | 24-04-01 | 25 | 56 | 100 | 100 | | | | | | | | | | | | |
| 4 | A | $w^{1118}$ | $w^{1118}$ | 24-03-28 | 25 | 84 | 86 | 100 | 87.00 | 75.80 | 100 | 6.00 | 8.42 | 0.00 | - | - | - | - | - | - |
| 4 | A | $w^{1118}$ | $w^{1118}$ | 24-03-28 | 25 | 96 | 79 | 100 | | | | | | | | | | | | |
| 4 | A | $w^{1118}$ | $w^{1118}$ | 24-03-28 | 25 | 84 | 72 | 100 | | | | | | | | | | | | |
| 4 | A | $w^{1118}$ | $w^{1118}$ | 24-03-28 | 25 | 84 | 67 | 100 | | | | | | | | | | | | |

| | | Cross | | Date | | | | | Mean | | | SD | | | P (Homo vs. Wt) | | | P (Homo vs. Hetero) | | |
|---|---|---|---|---|---|---|---|---|---|---|---|---|---|---|---|---|---|---|---|---|
| Exp | Rep | ♂ | V♀ | (YY-MM-DD) | # Eggs | % L1 | % P | % A | L1 | P | A | L1 | P | A | L1 | P | A | L1 | P | A |
| 4 | B | $w^{1118}$ | $w^{1118}$ | 24-03-29 | 25 | 92 | 91 | 100 | 92.00 | 82.68 | 100 | 0.00 | 7.94 | 0.00 | - | - | - | - | - | - |
| 4 | B | $w^{1118}$ | $w^{1118}$ | 24-03-29 | 25 | 92 | 87 | 100 | | | | | | | | | | | | |
| 4 | B | $w^{1118}$ | $w^{1118}$ | 24-03-29 | 25 | 92 | 74 | 100 | | | | | | | | | | | | |
| 4 | B | $w^{1118}$ | $w^{1118}$ | 24-03-29 | 25 | 92 | 78 | 100 | | | | | | | | | | | | |
| 4 | C | $w^{1118}$ | $w^{1118}$ | 24-03-30 | 25 | 76 | 69 | 100 | 86.00 | 68.88 | 100 | 8.33 | 9.73 | 0.00 | - | - | - | - | - | - |
| 4 | C | $w^{1118}$ | $w^{1118}$ | 24-03-30 | 25 | 96 | 58 | 100 | | | | | | | | | | | | |
| 4 | C | $w^{1118}$ | $w^{1118}$ | 24-03-30 | 25 | 88 | 82 | 100 | | | | | | | | | | | | |
| 4 | C | $w^{1118}$ | $w^{1118}$ | 24-03-30 | 25 | 84 | 67 | 100 | | | | | | | | | | | | |
| 4 | D | $w^{1118}$ | $w^{1118}$ | 24-04-01 | 25 | 84 | 86 | 95 | 83.00 | 86.33 | 95.85 | 10.00 | 6.20 | 5.31 | - | - | - | - | - | - |
| 4 | D | $w^{1118}$ | $w^{1118}$ | 24-04-01 | 25 | 72 | 78 | 100 | | | | | | | | | | | | |
| 4 | D | $w^{1118}$ | $w^{1118}$ | 24-04-01 | 25 | 96 | 92 | 100 | | | | | | | | | | | | |
| 4 | D | $w^{1118}$ | $w^{1118}$ | 24-04-01 | 25 | 80 | 90 | 89 | | | | | | | | | | | | |
| 5 | A | $DJ694^{H}$ | $DJ694^{H}$ | 24-04-17 | 25 | 44 | 73 | 100 | 50.00 | 71.10 | 89.60 | 5.16 | 21.40 | 12.49 | 0.0002 | 0.6601 | 0.3230 | 0.0053 | 0.1640 | 0.2919 |
| 5 | A | $DJ694^{H}$ | $DJ694^{H}$ | 24-04-17 | 25 | 56 | 100 | 100 | | | | | | | | | | | | |
| 5 | A | $DJ694^{H}$ | $DJ694^{H}$ | 24-04-17 | 25 | 52 | 62 | 75 | | | | | | | | | | | | |
| 5 | A | $DJ694^{H}$ | $DJ694^{H}$ | 24-04-17 | 25 | 48 | 50 | 83 | | | | | | | | | | | | |
| 5 | B | $DJ694^{H}$ | $DJ694^{H}$ | 24-04-18 | 25 | 68 | 82 | 93 | 66.00 | 91.00 | 96.85 | 7.66 | 5.77 | 3.70 | 0.1723 | 0.0000 | 0.7122 | 0.6269 | 0.1751 | 0.5317 |
| 5 | B | $DJ694^{H}$ | $DJ694^{H}$ | 24-04-18 | 25 | 76 | 95 | 95 | | | | | | | | | | | | |
| 5 | B | $DJ694^{H}$ | $DJ694^{H}$ | 24-04-18 | 25 | 60 | 93 | 100 | | | | | | | | | | | | |
| 5 | B | $DJ694^{H}$ | $DJ694^{H}$ | 24-04-18 | 25 | 60 | 93 | 100 | | | | | | | | | | | | |
| 5 | C | $DJ694^{H}$ | $DJ694^{H}$ | 24-04-19 | 25 | 84 | 81 | 88 | 73.00 | 86.58 | 88.53 | 8.87 | 3.81 | 8.40 | 0.0519 | 0.0004 | 0.3377 | 0.0265 | 0.1031 | 0.1526 |
| 5 | C | $DJ694^{H}$ | $DJ694^{H}$ | 24-04-19 | 25 | 76 | 90 | 100 | | | | | | | | | | | | |
| 5 | C | $DJ694^{H}$ | $DJ694^{H}$ | 24-04-19 | 25 | 64 | 88 | 86 | | | | | | | | | | | | |
| 5 | C | $DJ694^{H}$ | $DJ694^{H}$ | 24-04-19 | 25 | 68 | 88 | 80 | | | | | | | | | | | | |
| 5 | D | $DJ694^{H}$ | $DJ694^{H}$ | 24-04-20 | 25 | 68 | 94 | 94 | 69.00 | 88.53 | 95.68 | 11.49 | 3.96 | 5.38 | 0.0115 | 0.0071 | 0.3048 | 0.5569 | 0.3139 | 0.2323 |
| 5 | D | $DJ694^{H}$ | $DJ694^{H}$ | 24-04-20 | 25 | 56 | 86 | 100 | | | | | | | | | | | | |
| 5 | D | $DJ694^{H}$ | $DJ694^{H}$ | 24-04-20 | 25 | 68 | 88 | 100 | | | | | | | | | | | | |
| 5 | D | $DJ694^{H}$ | $DJ694^{H}$ | 24-04-20 | 25 | 84 | 86 | 89 | | | | | | | | | | | | |
| 5 | A | $DJ694^{H}$ | $w^{1118}$ | 24-04-17 | 25 | 80 | 90 | 95 | 75.00 | 88.38 | 97.08 | 10.52 | 4.13 | 3.39 | - | - | - | - | - | - |
| 5 | A | $DJ694^{H}$ | $w^{1118}$ | 24-04-17 | 25 | 76 | 84 | 94 | | | | | | | | | | | | |
| 5 | A | $DJ694^{H}$ | $w^{1118}$ | 24-04-17 | 25 | 60 | 93 | 100 | | | | | | | | | | | | |
| 5 | A | $DJ694^{H}$ | $w^{1118}$ | 24-04-17 | 25 | 84 | 86 | 100 | | | | | | | | | | | | |

Table S1. *DJ694* 's developmental lethality dataset and statistic results

| | | Cross | | Date | | | | | Mean | | | SD | | | P (Homo vs. Wt) | | | P (Homo vs. Hetero) | | |
|---|---|---|---|---|---|---|---|---|---|---|---|---|---|---|---|---|---|---|---|---|
| Exp | Rep | ♂ | V♀ | (YY-MM-DD) | # Eggs | % L1 | % P | % A | L1 | P | A | L1 | P | A | L1 | P | A | L1 | P | A |
| 5 | B | $DJ694^{H}$ | $w^{1118}$ | 24-04-18 | 25 | 68 | 94 | 94 | 63.00 | 77.85 | 98.45 | 8.87 | 16.10 | 3.10 | - | - | - | - | - | - |
| 5 | B | $DJ694^{H}$ | $w^{1118}$ | 24-04-18 | 25 | 60 | 67 | 100 | | | | | | | | | | | | |
| 5 | B | $DJ694^{H}$ | $w^{1118}$ | 24-04-18 | 25 | 52 | 62 | 100 | | | | | | | | | | | | |
| 5 | B | $DJ694^{H}$ | $w^{1118}$ | 24-04-18 | 25 | 72 | 89 | 100 | | | | | | | | | | | | |
| 5 | C | $DJ694^{H}$ | $w^{1118}$ | 24-04-19 | 25 | 60 | 93 | 100 | 58.00 | 93.25 | 96.28 | 5.16 | 5.81 | 4.36 | - | - | - | - | - | - |
| 5 | C | $DJ694^{H}$ | $w^{1118}$ | 24-04-19 | 25 | 52 | 100 | 100 | | | | | | | | | | | | |
| 5 | C | $DJ694^{H}$ | $w^{1118}$ | 24-04-19 | 25 | 56 | 86 | 92 | | | | | | | | | | | | |
| 5 | C | $DJ694^{H}$ | $w^{1118}$ | 24-04-19 | 25 | 64 | 94 | 93 | | | | | | | | | | | | |
| 5 | D | $DJ694^{H}$ | $w^{1118}$ | 24-04-20 | 25 | 72 | 89 | 100 | 73.33 | 92.80 | 100 | 2.31 | 6.24 | 0.00 | - | - | - | - | - | - |
| 5 | D | $DJ694^{H}$ | $w^{1118}$ | 24-04-20 | 25 | 72 | 100 | 100 | | | | | | | | | | | | |
| 5 | D | $DJ694^{H}$ | $w^{1118}$ | 24-04-20 | 25 | 76 | 90 | 100 | | | | | | | | | | | | |
| 5 | A | $DJ694^{L}$ | $DJ694^{L}$ | 24-04-17 | 25 | 24 | 83 | 80 | 29.00 | 95.85 | 86.25 | 8.87 | 8.30 | 11.09 | 0.0001 | 0.0091 | 0.1275 | 0.0076 | 0.1498 | 0.3944 |
| 5 | A | $DJ694^{L}$ | $DJ694^{L}$ | 24-04-17 | 25 | 20 | 100 | 100 | | | | | | | | | | | | |
| 5 | A | $DJ694^{L}$ | $DJ694^{L}$ | 24-04-17 | 25 | 40 | 100 | 90 | | | | | | | | | | | | |
| 5 | A | $DJ694^{L}$ | $DJ694^{L}$ | 24-04-17 | 25 | 32 | 100 | 75 | | | | | | | | | | | | |
| 5 | B | $DJ694^{L}$ | $DJ694^{L}$ | 24-04-18 | 25 | 64 | 88 | 93 | 55.00 | 91.18 | 95.73 | 10.52 | 7.27 | 5.08 | 0.0235 | 0.0001 | 0.5272 | 0.5620 | 0.2009 | 0.6361 |
| 5 | B | $DJ694^{L}$ | $DJ694^{L}$ | 24-04-18 | 25 | 44 | 100 | 100 | | | | | | | | | | | | |
| 5 | B | $DJ694^{L}$ | $DJ694^{L}$ | 24-04-18 | 25 | 64 | 94 | 100 | | | | | | | | | | | | |
| 5 | B | $DJ694^{L}$ | $DJ694^{L}$ | 24-04-18 | 25 | 48 | 83 | 90 | | | | | | | | | | | | |
| 5 | C | $DJ694^{L}$ | $DJ694^{L}$ | 24-04-19 | 25 | 48 | 83 | 90 | 43.00 | 86.10 | 97.50 | 3.83 | 5.28 | 5.00 | 0.1042 | 0.0019 | 0.4632 | 0.1434 | 0.1375 | 0.2809 |
| 5 | C | $DJ694^{L}$ | $DJ694^{L}$ | 24-04-19 | 25 | 40 | 90 | 100 | | | | | | | | | | | | |
| 5 | C | $DJ694^{L}$ | $DJ694^{L}$ | 24-04-19 | 25 | 40 | 80 | 100 | | | | | | | | | | | | |
| 5 | C | $DJ694^{L}$ | $DJ694^{L}$ | 24-04-19 | 25 | 44 | 91 | 100 | | | | | | | | | | | | |
| 5 | D | $DJ694^{L}$ | $DJ694^{L}$ | 24-04-20 | 25 | 28 | 72 | 100 | 37.00 | 90.63 | 94.38 | 10.52 | 13.44 | 6.93 | 0.0001 | 0.3293 | 0.2563 | 0.0696 | 0.6092 | 0.5702 |
| 5 | D | $DJ694^{L}$ | $DJ694^{L}$ | 24-04-20 | 25 | 44 | 91 | 100 | | | | | | | | | | | | |
| 5 | D | $DJ694^{L}$ | $DJ694^{L}$ | 24-04-20 | 25 | 28 | 100 | 86 | | | | | | | | | | | | |
| 5 | D | $DJ694^{L}$ | $DJ694^{L}$ | 24-04-20 | 25 | 48 | 100 | 92 | | | | | | | | | | | | |
| 5 | A | $DJ694^{L}$ | $w^{1118}$ | 24-04-17 | 25 | 84 | 91 | 90 | 74 | 80 | 95 | 14.14 | 15.34 | 7.42 | - | - | - | - | - | - |
| 5 | A | $DJ694^{L}$ | $w^{1118}$ | 24-04-17 | 25 | 64 | 69 | 100 | | | | | | | | | | | | |

Table S1. *DJ694*'s developmental lethality dataset and statistic results

| | | Cross | | Date | | | | | Mean | | | SD | | | P (Homo vs. Wt) | | | P (Homo vs. Hetero) | | |
|---|---|---|---|---|---|---|---|---|---|---|---|---|---|---|---|---|---|---|---|---|
| Exp | Rep | ♂ | V♀ | (YY-MM-DD) | # Eggs | % L1 | % P | % A | L1 | P | A | L1 | P | A | L1 | P | A | L1 | P | A |
| 5 | B | $DJ694^{L}$ | $w^{1118}$ | 24-04-18 | 25 | 60 | 67 | 90 | 47.00 | 79.68 | 97.50 | 23.86 | 14.26 | 5.00 | - | - | - | - | - | - |
| 5 | B | $DJ694^{L}$ | $w^{1118}$ | 24-04-18 | 25 | 12 | 100 | 100 | | | | | | | | | | | | |
| 5 | B | $DJ694^{L}$ | $w^{1118}$ | 24-04-18 | 25 | 64 | 75 | 100 | | | | | | | | | | | | |
| 5 | B | $DJ694^{L}$ | $w^{1118}$ | 24-04-18 | 25 | 52 | 77 | 100 | | | | | | | | | | | | |
| 5 | C | $DJ694^{L}$ | $w^{1118}$ | 24-04-19 | 25 | 72 | 83 | 87 | 56.00 | 94.20 | 93.13 | 14.97 | 7.84 | 5.44 | - | - | - | - | - | - |
| 5 | C | $DJ694^{L}$ | $w^{1118}$ | 24-04-19 | 25 | 60 | 93 | 93 | | | | | | | | | | | | |
| 5 | C | $DJ694^{L}$ | $w^{1118}$ | 24-04-19 | 25 | 36 | 100 | 100 | | | | | | | | | | | | |
| 5 | C | $DJ694^{L}$ | $w^{1118}$ | 24-04-19 | 25 | 56 | 100 | 93 | | | | | | | | | | | | |
| 5 | D | $DJ694^{L}$ | $w^{1118}$ | 24-04-20 | 25 | 52 | 92 | 92 | 52.00 | 95.10 | 97.23 | 4.00 | 4.25 | 4.79 | - | - | - | - | - | - |
| 5 | D | $DJ694^{L}$ | $w^{1118}$ | 24-04-20 | 25 | 48 | 100 | 100 | | | | | | | | | | | | |
| 5 | D | $DJ694^{L}$ | $w^{1118}$ | 24-04-20 | 25 | 56 | 93 | 100 | | | | | | | | | | | | |
| 5 | A | $DJ694^{O}$ | $DJ694^{O}$ | 24-04-17 | 25 | 12 | 100 | 100 | - | - | - | - | - | - | - | - | - | - | - | - |
| 5 | B | $DJ694^{O}$ | $DJ694^{O}$ | 24-04-18 | 25 | 28 | 86 | 100 | 30 | 93 | 94 | 2.83 | 10.04 | 8.84 | 0.0012 | 0.0019 | 0.4450 | 0.7396 | 0.9212 | 0.7016 |
| 5 | B | $DJ694^{O}$ | $DJ694^{O}$ | 24-04-18 | 25 | 32 | 100 | 88 | | | | | | | | | | | | |
| 5 | C | $DJ694^{O}$ | $DJ694^{O}$ | 24-04-19 | 25 | 32 | 88 | 100 | 33.00 | 90.23 | 97.23 | 6.83 | 7.06 | 5.55 | 0.0187 | 0.0325 | 0.5152 | 0.0399 | 0.1091 | 0.3559 |
| 5 | C | $DJ694^{O}$ | $DJ694^{O}$ | 24-04-19 | 25 | 40 | 90 | 89 | | | | | | | | | | | | |
| 5 | C | $DJ694^{O}$ | $DJ694^{O}$ | 24-04-19 | 25 | 36 | 100 | 100 | | | | | | | | | | | | |
| 5 | C | $DJ694^{O}$ | $DJ694^{O}$ | 24-04-19 | 25 | 24 | 83 | 100 | | | | | | | | | | | | |
| 5 | D | $DJ694^{O}$ | $DJ694^{O}$ | 24-04-20 | 25 | 36 | 67 | 83 | - | - | - | - | - | - | - | - | - | - | - | - |
| 5 | A | $DJ694^{O}$ | $w^{1118}$ | 24-04-17 | 25 | 20 | 60 | 100 | 18.67 | 86.67 | 100 | 2.31 | 23.09 | 0.00 | - | - | - | - | - | - |
| 5 | A | $DJ694^{O}$ | $w^{1118}$ | 24-04-17 | 25 | 16 | 100 | 100 | | | | | | | | | | | | |
| 5 | A | $DJ694^{O}$ | $w^{1118}$ | 24-04-17 | 25 | 20 | 100 | 100 | | | | | | | | | | | | |
| 5 | B | $DJ694^{O}$ | $w^{1118}$ | 24-04-18 | 25 | 24 | 83 | 100 | 32.00 | 87.80 | 96.45 | 7.30 | 9.54 | 7.10 | - | - | - | - | - | - |
| 5 | B | $DJ694^{O}$ | $w^{1118}$ | 24-04-18 | 25 | 40 | 90 | 100 | | | | | | | | | | | | |
| 5 | B | $DJ694^{O}$ | $w^{1118}$ | 24-04-18 | 25 | 36 | 78 | 100 | | | | | | | | | | | | |
| 5 | B | $DJ694^{O}$ | $w^{1118}$ | 24-04-18 | 25 | 28 | 100 | 86 | | | | | | | | | | | | |
| 5 | C | $DJ694^{O}$ | $w^{1118}$ | 24-04-19 | 25 | 48 | 92 | 100 | 47.00 | 97.93 | 100 | 8.25 | 4.15 | 0.00 | - | - | - | - | - | - |
| 5 | C | $DJ694^{O}$ | $w^{1118}$ | 24-04-19 | 25 | 56 | 100 | 100 | | | | | | | | | | | | |
| 5 | C | $DJ694^{O}$ | $w^{1118}$ | 24-04-19 | 25 | 48 | 100 | 100 | | | | | | | | | | | | |
| 5 | C | $DJ694^{O}$ | $w^{1118}$ | 24-04-19 | 25 | 36 | 100 | 100 | | | | | | | | | | | | |

| | | Cross | | Date | | | | | Mean | | | SD | | | P (Homo vs. Wt) | | | P (Homo vs. Hetero) | | |
|---|---|---|---|---|---|---|---|---|---|---|---|---|---|---|---|---|---|---|---|---|
| Exp | Rep | ♂ | V♀ | (YY-MM-DD) | # Eggs | % L1 | % P | % A | L1 | P | A | L1 | P | A | L1 | P | A | L1 | P | A |
| 5 | D | *DJ694*$^{O}$ | $w^{1118}$ | 24-04-20 | 25 | 48 | 83 | 100 | 45.00 | 94.20 | 98.23 | 11.49 | 7.84 | 3.55 | - | - | - | - | - | - |
| 5 | D | *DJ694*$^{O}$ | $w^{1118}$ | 24-04-20 | 25 | 36 | 100 | 100 | | | | | | | | | | | | |
| 5 | D | *DJ694*$^{O}$ | $w^{1118}$ | 24-04-20 | 25 | 36 | 100 | 100 | | | | | | | | | | | | |
| 5 | D | *DJ694*$^{O}$ | $w^{1118}$ | 24-04-20 | 25 | 60 | 93 | 93 | | | | | | | | | | | | |
| 5 | A | $w^{1118}$ | $w^{1118}$ | 24-04-17 | 25 | 72 | 72 | 92 | 80.00 | 76.25 | 96.65 | 5.66 | 6.18 | 3.94 | - | - | - | - | - | - |
| 5 | A | $w^{1118}$ | $w^{1118}$ | 24-04-17 | 25 | 84 | 76 | 100 | | | | | | | | | | | | |
| 5 | A | $w^{1118}$ | $w^{1118}$ | 24-04-17 | 25 | 84 | 72 | 100 | | | | | | | | | | | | |
| 5 | A | $w^{1118}$ | $w^{1118}$ | 24-04-17 | 25 | 80 | 85 | 94 | | | | | | | | | | | | |
| 5 | B | $w^{1118}$ | $w^{1118}$ | 24-04-18 | 25 | 68 | 59 | 100 | 74.00 | 56.73 | 97.93 | 6.93 | 3.28 | 4.15 | - | - | - | - | - | - |
| 5 | B | $w^{1118}$ | $w^{1118}$ | 24-04-18 | 25 | 68 | 53 | 100 | | | | | | | | | | | | |
| 5 | B | $w^{1118}$ | $w^{1118}$ | 24-04-18 | 25 | 80 | 55 | 100 | | | | | | | | | | | | |
| 5 | B | $w^{1118}$ | $w^{1118}$ | 24-04-18 | 25 | 80 | 60 | 92 | | | | | | | | | | | | |
| 5 | C | $w^{1118}$ | $w^{1118}$ | 24-04-19 | 25 | 40 | 100 | 90 | 55.00 | 100 | 94.18 | 11.94 | 0.00 | 6.86 | - | - | - | - | - | - |
| 5 | C | $w^{1118}$ | $w^{1118}$ | 24-04-19 | 25 | 60 | 100 | 87 | | | | | | | | | | | | |
| 5 | C | $w^{1118}$ | $w^{1118}$ | 24-04-19 | 25 | 68 | 100 | 100 | | | | | | | | | | | | |
| 5 | C | $w^{1118}$ | $w^{1118}$ | 24-04-19 | 25 | 52 | 100 | 100 | | | | | | | | | | | | |
| 5 | D | $w^{1118}$ | $w^{1118}$ | 24-04-20 | 25 | 88 | 96 | 100 | 92.00 | 97.88 | 98.93 | 5.66 | 2.46 | 2.15 | - | - | - | - | - | - |
| 5 | D | $w^{1118}$ | $w^{1118}$ | 24-04-20 | 25 | 88 | 100 | 100 | | | | | | | | | | | | |
| 5 | D | $w^{1118}$ | $w^{1118}$ | 24-04-20 | 25 | 100 | 96 | 100 | | | | | | | | | | | | |
| 5 | D | $w^{1118}$ | $w^{1118}$ | 24-04-20 | 25 | 92 | 100 | 96 | | | | | | | | | | | | |
| 6 | A | *DJ694*$^{L}$ | *DJ694*$^{L}$ | 23-01-26 | 25 | 80 | 95 | 90 | 67 | 85 | 95 | 18.81 | 14.14 | 7.42 | 0.1718 | 0.9455 | 0.5549 | 0.3527 | 0.2899 | 0.3830 |
| 6 | A | *DJ694*$^{L}$ | *DJ694*$^{L}$ | 23-01-26 | 15 | 53 | 75 | 100 | | | | | | | | | | | | |
| 6 | B | *DJ694*$^{L}$ | *DJ694*$^{L}$ | 23-01-27 | 25 | 76 | 100 | 79 | 80.00 | 92.78 | 89.23 | 4.62 | 6.06 | 9.70 | 0.2283 | 0.4172 | 0.1082 | 0.7049 | 0.4028 | 0.1082 |
| 6 | B | *DJ694*$^{L}$ | *DJ694*$^{L}$ | 23-01-27 | 25 | 76 | 95 | 83 | | | | | | | | | | | | |
| 6 | B | *DJ694*$^{L}$ | *DJ694*$^{L}$ | 23-01-27 | 25 | 84 | 86 | 95 | | | | | | | | | | | | |
| 6 | B | *DJ694*$^{L}$ | *DJ694*$^{L}$ | 23-01-27 | 25 | 84 | 91 | 100 | | | | | | | | | | | | |
| 6 | A | *DJ694*$^{L}$ | $w^{1118}$ | 23-01-26 | 25 | 72 | 95 | 100 | 76.00 | 94.88 | 98.55 | 4.62 | 7.08 | 2.90 | - | - | - | - | - | - |
| 6 | A | *DJ694*$^{L}$ | $w^{1118}$ | 23-01-26 | 25 | 80 | 85 | 94 | | | | | | | | | | | | |
| 6 | A | *DJ694*$^{L}$ | $w^{1118}$ | 23-01-26 | 25 | 80 | 100 | 100 | | | | | | | | | | | | |
| 6 | A | *DJ694*$^{L}$ | $w^{1118}$ | 23-01-26 | 25 | 72 | 100 | 100 | | | | | | | | | | | | |

Table S1. *DJ694*'s developmental lethality dataset and statistic results 

| | | Cross | | Date | | | | | Mean | | | SD | | | P (Homo vs. Wt) | | | P (Homo vs. Hetero) | | |
|---|---|---|---|---|---|---|---|---|---|---|---|---|---|---|---|---|---|---|---|---|
| Exp | Rep | ♂ | V♀ | (YY-MM-DD) | # Eggs | % L1 | % P | % A | L1 | P | A | L1 | P | A | L1 | P | A | L1 | P | A |
| 6 | B | $DJ694^{L}$ | $w^{1118}$ | 23-01-27 | 25 | 80 | 90 | 100 | 79.00 | 96.25 | 98.70 | 2.00 | 4.79 | 2.60 | - | - | - | - | - | - |
| 6 | B | $DJ694^{L}$ | $w^{1118}$ | 23-01-27 | 25 | 80 | 95 | 95 | | | | | | | | | | | | |
| 6 | B | $DJ694^{L}$ | $w^{1118}$ | 23-01-27 | 25 | 80 | 100 | 100 | | | | | | | | | | | | |
| 6 | B | $DJ694^{L}$ | $w^{1118}$ | 23-01-27 | 25 | 76 | 100 | 100 | | | | | | | | | | | | |
| 6 | A | $w^{1118}$ | $w^{1118}$ | 23-01-26 | 25 | 84 | 81 | 100 | 84.00 | 85.55 | 97.75 | 8.64 | 5.91 | 4.50 | - | - | - | - | - | - |
| 6 | A | $w^{1118}$ | $w^{1118}$ | 23-01-26 | 25 | 76 | 90 | 100 | | | | | | | | | | | | |
| 6 | A | $w^{1118}$ | $w^{1118}$ | 23-01-26 | 25 | 96 | 92 | 91 | | | | | | | | | | | | |
| 6 | A | $w^{1118}$ | $w^{1118}$ | 23-01-26 | 25 | 80 | 80 | 100 | | | | | | | | | | | | |
| 6 | B | $w^{1118}$ | $w^{1118}$ | 23-01-27 | 25 | 92 | 87 | 100 | 86.00 | 89.38 | 98.70 | 7.66 | 4.92 | 2.60 | - | - | - | - | - | - |
| 6 | B | $w^{1118}$ | $w^{1118}$ | 23-01-27 | 25 | 84 | 91 | 95 | | | | | | | | | | | | |
| 6 | B | $w^{1118}$ | $w^{1118}$ | 23-01-27 | 25 | 92 | 96 | 100 | | | | | | | | | | | | |
| 6 | B | $w^{1118}$ | $w^{1118}$ | 23-01-27 | 25 | 76 | 84 | 100 | | | | | | | | | | | | |
| 7 | A | $DJ694^{L}$ | $DJ694^{L}$ | 23-02-08 | 25 | 48 | 83 | 100 | 44.00 | 79.60 | 100 | 4.62 | 8.87 | 0.00 | 7E-06 | 0.1149 | 0.1726 | 3E-05 | 0.0107 | 0.1783 |
| 7 | A | $DJ694^{L}$ | $DJ694^{L}$ | 23-02-08 | 25 | 48 | 75 | 100 | | | | | | | | | | | | |
| 7 | A | $DJ694^{L}$ | $DJ694^{L}$ | 23-02-08 | 25 | 40 | 90 | 100 | | | | | | | | | | | | |
| 7 | A | $DJ694^{L}$ | $DJ694^{L}$ | 23-02-08 | 25 | 40 | 70 | 100 | | | | | | | | | | | | |
| 7 | B | $DJ694^{L}$ | $DJ694^{L}$ | 23-02-09 | 25 | 80 | 80 | 88 | 71.00 | 87.68 | 86.60 | 6.00 | 11.23 | 19.38 | 0.0008 | 0.3830 | 0.2934 | 0.0300 | 0.4576 | 0.2161 |
| 7 | B | $DJ694^{L}$ | $DJ694^{L}$ | 23-02-09 | 25 | 68 | 77 | 100 | | | | | | | | | | | | |
| 7 | B | $DJ694^{L}$ | $DJ694^{L}$ | 23-02-09 | 25 | 68 | 100 | 59 | | | | | | | | | | | | |
| 7 | B | $DJ694^{L}$ | $DJ694^{L}$ | 23-02-09 | 25 | 68 | 94 | 100 | | | | | | | | | | | | |
| 7 | A | $DJ694^{L}$ | $w^{1118}$ | 23-02-08 | 25 | 88 | 100 | 100 | 84.00 | 97.75 | 96.20 | 5.66 | 4.50 | 4.99 | - | - | - | - | - | - |
| 7 | A | $DJ694^{L}$ | $w^{1118}$ | 23-02-08 | 25 | 76 | 100 | 90 | | | | | | | | | | | | |
| 7 | A | $DJ694^{L}$ | $w^{1118}$ | 23-02-08 | 25 | 84 | 100 | 95 | | | | | | | | | | | | |
| 7 | A | $DJ694^{L}$ | $w^{1118}$ | 23-02-08 | 25 | 88 | 91 | 100 | | | | | | | | | | | | |
| 7 | B | $DJ694^{L}$ | $w^{1118}$ | 23-02-09 | 25 | 76 | 100 | 100 | 83.00 | 93.10 | 100 | 6.00 | 7.80 | 0.00 | - | - | - | - | - | - |
| 7 | B | $DJ694^{L}$ | $w^{1118}$ | 23-02-09 | 25 | 88 | 96 | 100 | | | | | | | | | | | | |
| 7 | B | $DJ694^{L}$ | $w^{1118}$ | 23-02-09 | 25 | 80 | 95 | 100 | | | | | | | | | | | | |
| 7 | B | $DJ694^{L}$ | $w^{1118}$ | 23-02-09 | 25 | 88 | 82 | 100 | | | | | | | | | | | | |
| 7 | A | $w^{1118}$ | $w^{1118}$ | 23-02-08 | 25 | 92 | 96 | 100 | 87.00 | 89.55 | 96.13 | 3.83 | 6.16 | 5.01 | - | - | - | - | - | - |
| 7 | A | $w^{1118}$ | $w^{1118}$ | 23-02-08 | 25 | 84 | 81 | 100 | | | | | | | | | | | | |
| 7 | A | $w^{1118}$ | $w^{1118}$ | 23-02-08 | 25 | 84 | 91 | 90 | | | | | | | | | | | | |
| 7 | A | $w^{1118}$ | $w^{1118}$ | 23-02-08 | 25 | 88 | 91 | 95 | | | | | | | | | | | | |

Table S1. *DJ694* 's developmental lethality dataset and statistic results



| Exp | Rep | Cross ♂ | V♀ | Date (YY-MM-DD) | # Eggs | % L1 | % P | % A | Mean L1 | Mean P | Mean A | SD L1 | SD P | SD A | P (Homo vs. Wt) L1 | P (Homo vs. Wt) P | P (Homo vs. Wt) A | P (Homo vs. Hetero) L1 | P (Homo vs. Hetero) P | P (Homo vs. Hetero) A |
|---|---|---|---|---|---|---|---|---|---|---|---|---|---|---|---|---|---|---|---|---|
| 7 | B | $w^{1118}$ | $w^{1118}$ | 23-02-09 | 25 | 96 | 88 | 100 | 93.00 | 93.55 | 97.85 | 3.83 | 5.46 | 2.49 | - | - | - | - | - | - |
| 7 | B | $w^{1118}$ | $w^{1118}$ | 23-02-09 | 25 | 88 | 91 | 100 | | | | | | | | | | | | |
| 7 | B | $w^{1118}$ | $w^{1118}$ | 23-02-09 | 25 | 96 | 100 | 96 | | | | | | | | | | | | |
| 7 | B | $w^{1118}$ | $w^{1118}$ | 23-02-09 | 25 | 92 | 96 | 96 | | | | | | | | | | | | |

Column "Exp" show the number of independent experiments done.
Column "Rep" show the technical replicates within each independent experiment.
Survival was assessed at three successive developmental transitions: egg to L1 larvae, L1 larvae to pupae, and pupae to adults. Counts were taken 24 hours after egg transfer for egg hatching, 7–9 days later for pupal development, and a further 5–7 days thereafter for adult emergence.
Column "% L1" is the percentage of eggs that hatched.
Column "% P" is the percentage of 1d larvae that pupaed.
Column "% A" is the percentage of pupae that become adults.
Column "Mean" and "SD" are the average and standard deviation calculated within each technical replicates.
Two sample unpaired two tail T-Tests were performed to compare the development lethality between homozygous *DJ694* (HOMO) to the wildtype control (Wt), and to the heterozygous *DJ694/+* (Hetero).

| | | Date | | Avg Cumulative #Eggs/♀ | | Avg Emergence% | |
|---|---|---|---|---|---|---|---|
| Cross | Rep | YY-MM-DD | n♀ | Mean | sd | Mean | sd |
| *DJ694* ♀ x *DJ694* ♂ | 1 | 04-03-16 | 15 | 9.40 | 10.38 | 62.23 | 3.66 |
| | 2 | 05-02-20 | 15 | 33.23 | 10.51 | 50.94 | 20.93 |
| $w^{1118/CS10}$ ♀ x *DJ694* ♂ | 1 | 04-03-16 | 15 | 39.08 | 15.82 | 67.59 | 6.80 |
| | 2 | 05-02-20 | 15 | 68.48 | 4.99 | 64.24 | 3.72 |
| *DJ694* ♀ x $w^{1118/CS10}$ ♂ | 1 | 04-03-16 | 15 | 13.57 | 7.76 | 82.11 | 1.41 |
| | 2 | 05-02-20 | 15 | 28.38 | 13.19 | 72.66 | 6.98 |
| $w^{1118/CS10}$ ♀ x $w^{1118}$ ♂ | 1 | 04-03-16 | 15 | 60.93 | 26.14 | 84.06 | 8.10 |
| | 2 | 05-02-20 | 15 | 61.07 | 0.90 | 84.49 | 4.53 |
| *DJ694/+* ♀ x $w^{1118/CS10}$ ♂ | 1 | 04-03-16 | 15 | 78.47 | 19.16 | 86.37 | 11.47 |
| $w^{1118/CS10}$ ♀ x *DJ694/+* ♂ | 1 | 04-03-16 | 15 | 54.76 | 17.54 | 88.86 | 2.65 |
| $DJ694;UAS\text{-}EDTP^{A}$ ♀ x $w^{1118}$ ♂ | 3 | 04-11-30 | 15 | 45.54 | 16.36 | 54.17 | 22.07 |
| $DJ694;UAS\text{-}EDTP^{B}$ ♀ x $w^{1118}$ ♂ | 3 | 04-11-30 | 15 | 73.79 | 30.97 | 81.39 | 2.09 |
| *DJ694* ♀ x $w^{1118}$ ♂ | 3 | 04-11-30 | 20 | 18.87 | 9.59 | 15.06 | 11.38 |
| $w^{1118}$ ♀ x $DJ694;UAS\text{-}EDTP^{A}$ ♂ | 3 | 04-11-30 | 20 | 55.78 | 30.63 | 48.75 | 23.00 |
| $w^{1118}$ ♀ x $DJ694;UAS\text{-}EDTP^{B}$ ♂ | 3 | 04-11-30 | 20 | 90.29 | 51.06 | 60.05 | 12.47 |
| $w^{1118}$ ♀ x *DJ694* ♂ | 3 | 04-11-30 | 15 | 40.70 | 30.78 | 7.69 | 7.05 |
| $DJ694^{H}$ ♀ x $w^{1118}$ ♂ | 4 | 24-07-09 | 24 | 33.40 | 19.36 | - | - |
| | 5 | 24-07-10 | 24 | 28.67 | 17.76 | - | - |
| $DJ694^{H}/+$ ♀ x $w^{1118}$ ♂ | 4 | 24-07-09 | 24 | 79.98 | 22.76 | - | - |
| $DJ694^{L}$ ♀ x $w^{1118}$ ♂ | 4 | 24-07-09 | 24 | 29.79 | 26.46 | - | - |
| | 5 | 24-07-10 | 24 | 42.58 | 16.82 | - | - |
| $DJ694^{L}/+$ ♀ x $w^{1118}$ ♂ | 4 | 24-07-09 | 24 | 105.08 | 11.65 | - | - |
| | 5 | 24-07-10 | 24 | 91.65 | 15.89 | - | - |
| $DJ694^{O}$ ♀ x $w^{1118}$ ♂ | 4 | 24-07-09 | 24 | 30.75 | 19.52 | - | - |
| | 5 | 24-07-10 | 12 | 35.92 | 7.52 | - | - |
| $DJ694^{O}/+$ ♀ x $w^{1118}$ ♂ | 4 | 24-07-09 | 24 | 113.21 | 10.55 | - | - |
| | 5 | 24-07-10 | 24 | 114.43 | 11.14 | - | - |
| $DJ694^{P}$ ♀ x $w^{1118}$ ♂ | 4 | 24-07-09 | 9 | 30.69 | 16.23 | - | - |
| | 5 | 24-07-10 | 12 | 41.63 | 17.04 | - | - |
| $DJ694^{P}/+$ ♀ x $w^{1118}$ ♂ | 4 | 24-07-09 | 24 | 98.67 | 6.12 | - | - |
| | 5 | 24-07-10 | 24 | 105.98 | 11.28 | - | - |
| $w^{1118}$ ♀ x $w^{1118}$ ♂ | 4 | 24-07-09 | 18 | 78.44 | 13.81 | - | - |
| | 5 | 24-07-10 | 21 | 78.00 | 17.83 | - | - |

| | | | | | | | |
|---|---|---|---|---|---|---|---|
| *DJ694*$^{L}$;*UAS-EDTP*$^{E}$/+ ♀ x *w*$^{1118}$ ♂ | 4 | 24-07-09 | 18 | 81.69 | 17.53 | - | - |
| | 5 | 24-07-10 | 21 | 76.67 | 14.51 | - | - |

Columns indicate the cross, replicate number, date of experiment, and the initial number of females analyzed (n♀). The value of n♀ represents the number of females at the start of the assay; some females died during the experiment. “Avg Cumulative #Eggs/♀” represents the average cumulative number of eggs laid per female from day 29 to day 40 for all replicates. “Avg Emergence %” represents the average percentage of eggs that successfully became adults. For Replicates 1 and 2, emergence percentage was calculated as the total number of adults emerged divided by the total number of eggs laid across 85d. For Replicate 3, emergence percentage was calculated as the total number of adults eclosed divided by the total number of eggs laid from day 29 to day 48. Mean and standard deviation (sd) are shown for each measurement. All experiments are done in 25˚C.

Table S3. Statistical analysis of fertility differences among *DJ694*, controls, and rescue strains.



| | | DJ vs. W | | | DJ/+ vs. W | | | DJ vs. DJ/+ | | | Rescue's Geno | Res vs. DJ | | | Res vs. W | Res vs. DJ/+ |
|---|---|---|---|---|---|---|---|---|---|---|---|---|---|---|---|---|
| Rep | ♀GAL4 | %↓ | Δ | P | %↓ | Δ | P | %↓ | Δ | P | | %↑ | Δ | P | P | P |
| 1 | *DJ694* | -77.73 | ↓ | 0.0396 | 28.79 | ↑ | 0.4017 | -82.71 | ↓ | 0.0055 | - | - | - | - | - | - |
| 2 | *DJ694* | -53.53 | ↓ | 0.0128 | - | - | - | - | - | - | - | - | - | - | - | - |
| 3 | *DJ694* | - | - | - | - | - | - | - | - | - | *DJ694;UAS-EDTP*$^{A}$ | 141.32 | ↑ | 0.0407 | - | - |
| | | - | - | - | - | - | - | - | - | - | *DJ694;UAS-EDTP*$^{B}$ | 291.02 | ↑ | 0.0186 | - | - |
| 4 | *DJ694*$^{H}$ | -51.47 | ↓ | 0.0037 | 16.23 | ↑ | 0.3292 | -58.24 | ↓ | 0.0006 | - | - | - | - | - | - |
| | *DJ694*$^{L}$ | -56.71 | ↓ | 0.0059 | 52.71 | ↑ | 0.0009 | -71.65 | ↓ | 3.53E-06 | *DJ694;UAS-EDTP*$^{E}$/+ | 134.83 | ↑ | 0.0089 | 0.9254 | 0.0040 |
| | *DJ694*$^{O}$ | -55.31 | ↓ | 0.0023 | 64.52 | ↑ | 0.0001 | -72.84 | ↓ | 5.02E-08 | - | - | - | - | - | - |
| | *DJ694*$^{P}$ | -55.40 | ↓ | 0.0013 | 43.38 | ↑ | 0.0462 | -68.90 | ↓ | 0.0001 | - | - | - | - | - | - |
| 5 | *DJ694*$^{H}$ | -61.64 | ↓ | 0.0002 | - | - | - | - | - | - | - | - | - | - | - | - |
| | *DJ694*$^{L}$ | -43.02 | ↓ | 0.0031 | 22.64 | ↑ | 0.0757 | -53.53 | ↓ | 3.27E-05 | *DJ694;UAS-EDTP*$^{E}$/+ | 71.33 | ↑ | 0.0024 | 0.8446 | 0.0348 |
| | *DJ694*$^{O}$ | -51.94 | ↓ | 0.0034 | 53.12 | ↑ | 0.0004 | -68.61 | ↓ | 5.71E-07 | - | - | - | - | - | - |
| | *DJ694*$^{P}$ | -44.30 | ↓ | 0.0157 | 41.81 | ↑ | 0.0024 | -60.72 | ↓ | 3.26E-05 | - | - | - | - | - | - |

Fertility differences were assessed by comparisons of the cumulative number of eggs laid per female between days 29 and 40. Percent decline or increase represent either the reduction or increase in fertility, calculated from the average cumulative egg counts. Specifically, %↓ indicates the decline in fertility of homozygous *DJ694* relative to w (DJ vs. W), the decline in fertility of homozygous *DJ694* relative to *DJ694/+* (DJ vs. DJ/+), or the decline in fertility of *DJ694/+* relative to w (DJ/+ vs. W), as indicated by each comparison. Percent increase (%↑) represents the increase in fertility of rescue genotypes relative to homozygous *DJ694*, which reflects the extent to which the *DJ694* fertility phenotype is rescued (Res vs. DJ). The "Rescue's Geno" column indicates the genotype used for fertility rescue in each comparison. P-values were calculated independently for each biological replicate using two-tailed, two-sample t-tests assuming equal variance. Column "Δ" under each comparison indicates whether a noticeable difference between the two groups is observed. A directional arrow (↓ or ↑) indicates that the mean of the first genotype is lower/higher than the mean value of the second. Comparisons involving rescue genotypes are shown relative to both *DJ694* (Res vs. DJ), wildtype (Res vs. W), and *DJ694/+* (Res vs. DJ/+). All females are mated with *w* males.

Table S4. Statistical comparison of emergence percentages between genetic crosses 

| Reps | Age Range | Crosses | *DJ694* ♂ x *DJ694* ♀ | | *DJ694* ♂ x *w* ♀ | | *w* ♂ x *DJ694* ♀ | | *w* ♂ x *w* ♀ | | *DJ694/+* ♂ x *w* ♀ | | *w* ♂ x *DJ694/+* ♀ | |
|---|---|---|---|---|---|---|---|---|---|---|---|---|---|---|
| | | | P-value | Δ | P-value | Δ | P-value | Δ | P-value | Δ | P-value | Δ | P-value | Δ |
| 1 | 1-85d | *DJ694* ♂ x *DJ694* ♀ | | | | | | | | | | | | |
| | | *DJ694* ♂ x *w* ♀ | 0.2956 | X | | | | | | | | | | |
| | | *w* ♂ x *DJ694* ♀ | 0.0009 | ↓ | 0.0223 | ↓ | | | | | | | | |
| | | *w* ♂ x *w* ♀ | 0.0131 | ↓ | 0.0542 | ↓ | 0.7019 | X | | | | | | |
| | | *DJ694/+* ♂ x *w* ♀ | 0.0005 | ↓ | 0.0072 | ↓ | 0.0175 | ↓ | 0.3838 | X | | | | |
| | | *w* ♂ x *DJ694/+* ♀ | 0.0255 | ↓ | 0.0712 | ↓ | 0.5574 | X | 0.7896 | X | 0.7322 | X | | |
| 2 | 1-85d | *DJ694* ♂ x *DJ694* ♀ | | | | | | | | | | | | |
| | | *DJ694* ♂ x *w* ♀ | 0.3396 | X | | | | | | | | | | |
| | | *w* ♂ x *DJ694* ♀ | 0.1634 | X | 0.1390 | X | | | | | | | | |
| | | *w* ♂ x *w* ♀ | 0.0533 | ↓ | 0.0039 | ↓ | 0.0695 | ↓ | | | | | | |

| Reps | Age Range | Crosses | *w* ♂ x *DJ694;UAS-EDTP*$^{A}$ ♀ | | *w* ♂ x *DJ694;UAS-EDTP*$^{B}$ ♀ | | *w* ♂ x *DJ694* ♀ | | *DJ694;UAS-EDTP*$^{A}$ ♂ x *w* ♀ | | *DJ694;UAS-EDTP*$^{B}$ ♂ x *w* ♀ | | *DJ694* ♂ x *w* ♀ | |
|---|---|---|---|---|---|---|---|---|---|---|---|---|---|---|
| | | | P-value | Δ | P-value | Δ | P-value | Δ | P-value | Δ | P-value | Δ | P-value | Δ |
| 3 | 29-48d | *w* ♂ x *DJ694;UAS-EDTP*$^{A}$ ♀ | | | | | | | | | | | | |
| | | *w* ♂ x *DJ694;UAS-EDTP*$^{B}$ ♀ | 0.1006 | - | | | | | | | | | | |
| | | *w* ♂ x *DJ694* ♀ | 0.0192 | ↑ | 0.0001 | ↑ | | | | | | | | |
| | | *DJ694;UAS-EDTP*$^{A}$ ♂ x *w* ♀ | 0.7828 | - | 0.0706 | - | 0.0344 | ↓ | | | | | | |
| | | *DJ694;UAS-EDTP*$^{B}$ ♂ x *w* ♀ | 0.6205 | - | 0.0507 | - | 0.0013 | ↓ | 0.4352 | - | | | | |
| | | *DJ694* ♂ x *w* ♀ | 0.0097 | ↑ | 1.23E-05 | ↑ | 0.6095 | X | 0.0180 | ↑ | 0.0004 | ↑ | | |

Statistical comparisons of emergence percentages were performed between two genetic crosses for each biological replicate, as indicated in the table. Emergence percentage was calculated as the proportion of eggs that successfully developed into adults. P-values are reported for each comparison and were calculated independently for each replicate using two-tailed, two-sample t-tests assuming equal variance. Column "Δ" under each comparison indicates whether a noticeable difference between the two groups is observed. Directional arrows (↓ or ↑) indicates that the mean value of the genotype in the horizontal is lower/higher than the mean value of the genotype in the vertical. "X" indicates the difference is not significant.

Table S5. Female fertility data (eggs laid per female per day) for the indicated genetic crosses 

| Cross | Rep | Start Date (YYMMDD) | Vial | Days 1 | 2 | 3 | 4 | 5 | 6 | 7 | 8 | 9 | 10 | 11 | 12 | 13 | 14 | 15 | 16 | 17 | 18 | 19 | 20 |
|---|---|---|---|---|---|---|---|---|---|---|---|---|---|---|---|---|---|---|---|---|---|---|---|
| *DJ694* ♀ x *DJ694* ♂ | 1 | 04-03-16 | 1 | 1.8 | 6.6 | 6.0 | 14.4 | 7.2 | 11.2 | 7.8 | 16.6 | 12.4 | 0.6 | 3.0 | 14.2 | 3.4 | 3.8 | 9.0 | 13.0 | 3.4 | 10.8 | 1.2 | 2.6 |
| | | | 2 | 1.2 | 14.0 | 3.4 | 20.4 | 4.2 | 15.0 | 1.0 | 11.4 | 1.0 | 7.4 | 6.0 | 12.0 | 12.0 | 4.4 | 17.0 | 4.6 | 1.2 | 14.8 | 1.0 | 0.8 |
| | | | 3 | 8.8 | 12.2 | 8.4 | 9.2 | 7.0 | 5.8 | 2.6 | 8.0 | 3.6 | 4.0 | 3.4 | 3.8 | 4.2 | 2.2 | 2.2 | 0.6 | 0.2 | 2.4 | 0.4 | 0.4 |
| | 2 | 05-02-20 | 1 | 7.2 | 8.4 | 13.4 | 5.8 | 5.8 | 11.6 | 9.2 | 2.6 | 8.0 | 6.6 | 10.2 | 11.4 | 9.6 | 15.0 | 6.4 | 14.4 | 6.8 | 4.0 | 2.8 | 5.2 |
| | | | 2 | 7.0 | 12.6 | 12.4 | 12.0 | 7.2 | 9.6 | 15.0 | 6.6 | 4.2 | 10.8 | 3.2 | 14.0 | 14.6 | 5.8 | 6.6 | 9.0 | 3.8 | 14.2 | 4.0 | 11.5 |
| | | | 3 | 4.4 | 10.0 | 13.0 | 8.0 | 8.0 | 8.6 | 10.8 | 10.4 | 3.8 | 10.0 | 6.6 | 11.8 | 6.8 | 13.8 | 2.4 | 13.2 | 5.6 | 12.2 | 3.6 | 2.0 |
| $w^{1118/CS10}$ ♀ x *DJ694* ♂ | 1 | 04-03-16 | 1 | 7.8 | 9.2 | 11.2 | 10.2 | 6.6 | 5.0 | 7.4 | 9.8 | 9.4 | 5.0 | 10.0 | 7.4 | 5.2 | 6.4 | 9.2 | 2.2 | 6.0 | 5.8 | 2.8 | 6.8 |
| | | | 2 | 13.4 | 17.6 | 13.4 | 10.8 | 9.0 | 11.4 | 8.4 | 11.6 | 10.2 | 9.0 | 13.0 | 8.8 | 8.2 | 6.2 | 7.2 | 6.2 | 5.8 | 6.0 | 5.6 | 6.8 |
| | | | 3 | 8.4 | 12.2 | 11.2 | 8.2 | 6.8 | 7.2 | 8.8 | 7.0 | 9.2 | 8.8 | 7.8 | 7.6 | 6.0 | 5.6 | 5.6 | 5.4 | 4.2 | 4.0 | 1.6 | 2.2 |
| | 2 | 05-02-20 | 1 | 12.2 | 15.0 | 7.0 | 12.6 | 7.8 | 11.8 | 8.8 | 9.0 | 11.4 | 13.2 | 10.0 | 10.2 | 10.6 | 12.0 | 7.8 | 8.8 | 9.0 | 7.0 | 7.2 | 10.2 |
| | | | 2 | 18.6 | 8.6 | 13.6 | 11.8 | 9.0 | 8.0 | 11.2 | 8.2 | 14.8 | 8.6 | 13.8 | 13.5 | 10.0 | 11.3 | 11.0 | 9.0 | 8.3 | 10.0 | 9.3 | 9.0 |
| | | | 3 | 13.8 | 12.8 | 10.8 | 12.8 | 4.6 | 14.0 | 8.4 | 7.6 | 10.8 | 12.2 | 10.8 | 11.8 | 10.2 | 5.2 | 11.0 | 3.4 | 4.4 | 13.8 | 9.0 | 9.0 |
| *DJ694* ♀ x $w^{1118/CS10}$ ♂ | 1 | 04-03-16 | 1 | 1.4 | 13.4 | 3.4 | 14.8 | 1.6 | 8.6 | 11.8 | 2.5 | 1.0 | 17.5 | 9.0 | 1.5 | 13.8 | 1.8 | 5.0 | 1.5 | 0.8 | 1.5 | 2.3 | 0.3 |
| | | | 2 | 7.2 | 12.0 | 13.6 | 7.0 | 6.0 | 11.2 | 0.8 | 7.0 | 12.0 | 8.6 | 15.4 | 2.6 | 11.2 | 1.4 | 10.8 | 4.4 | 2.6 | 6.4 | 1.0 | 3.0 |
| | | | 3 | 1.0 | 8.8 | 10.4 | 5.4 | 3.6 | 14.6 | 6.8 | 11.8 | 11.6 | 2.4 | 20.2 | 2.6 | 9.6 | 1.4 | 13.6 | 0.8 | 1.0 | 13.6 | 1.0 | 0.6 |
| | 2 | 05-02-20 | 1 | 9.4 | 12.0 | 10.6 | 8.0 | 9.8 | 9.0 | 5.6 | 4.8 | 15.8 | 3.2 | 7.0 | 16.4 | 1.8 | 7.6 | 13.2 | 5.0 | 4.6 | 11.0 | 1.2 | 3.6 |
| | | | 2 | 6.8 | 8.2 | 13.0 | 7.6 | 10.8 | 13.4 | 7.0 | 9.2 | 6.4 | 11.6 | 1.6 | 10.6 | 12.6 | 12.6 | 2.2 | 11.2 | 2.4 | 19.4 | 4.0 | 3.4 |
| | | | 3 | 2.4 | 18.2 | 4.8 | 3.8 | 11.2 | 10.2 | 4.8 | 3.2 | 13.4 | 3.8 | 9.0 | 2.0 | 14.6 | 5.4 | 10.8 | 4.0 | 2.8 | 13.5 | 2.3 | 2.0 |
| $w^{1118/CS10}$ ♀ x $w^{1118}$ ♂ | 1 | 04-03-16 | 1 | 15.6 | 12.0 | 11.2 | 8.2 | 7.4 | 7.0 | 7.0 | 8.8 | 7.4 | 6.8 | 7.2 | 7.6 | 4.6 | 3.8 | 5.6 | 2.6 | 5.2 | 3.4 | 2.6 | 4.2 |
| | | | 2 | 10.2 | 15.6 | 13.0 | 7.8 | 8.4 | 10.4 | 5.2 | 9.2 | 9.2 | 8.0 | 8.0 | 7.8 | 8.8 | 5.4 | 4.4 | 7.0 | 5.4 | 9.0 | 1.4 | 7.0 |
| | | | 3 | 9.4 | 17.2 | 16.6 | 11.8 | 8.0 | 7.8 | 8.2 | 11.2 | 9.4 | 8.8 | 9.2 | 7.8 | 6.4 | 7.8 | 5.8 | 4.0 | 6.8 | 5.6 | 3.6 | 5.8 |
| | 2 | 05-02-20 | 1 | 8.8 | 8.8 | 9.6 | 5.8 | 4.6 | 13.6 | 11.0 | 5.6 | 7.2 | 11.8 | 8.8 | 9.0 | 9.8 | 9.2 | 4.0 | 13.6 | 7.0 | 7.0 | 10.0 | 8.4 |
| | | | 2 | 14.0 | 15.4 | 6.2 | 11.6 | 8.2 | 7.6 | 14.0 | 7.8 | 7.0 | 10.0 | 10.0 | 7.6 | 10.4 | 9.2 | 9.0 | 9.6 | 8.4 | 9.0 | 4.8 | 11.2 |
| | | | 3 | 18.2 | 13.6 | 12.0 | 6.4 | 6.6 | 10.4 | 12.0 | 7.2 | 7.8 | 7.0 | 9.8 | 10.4 | 7.8 | 9.0 | 5.6 | 8.8 | 5.6 | 9.4 | 7.4 | 10.8 |
| *DJ694/+* ♀ x $w^{1118/CS10}$ ♂ | 1 | 04-03-16 | 1 | 8.2 | 10.8 | 10.8 | 8.6 | 7.2 | 11.2 | 3.6 | 13.8 | 5.0 | 8.0 | 10.0 | 7.2 | 5.2 | 4.0 | 5.0 | 6.6 | 5.2 | 8.0 | 3.2 | 7.6 |
| | | | 2 | 7.4 | 20.0 | 7.8 | 13.4 | 6.0 | 12.6 | 3.2 | 15.6 | 6.2 | 11.0 | 11.2 | 10.8 | 9.0 | 6.2 | 8.6 | 2.6 | 12.8 | 8.2 | 5.5 | 7.3 |
| | | | 3 | 9.6 | 16.4 | 6.8 | 17.2 | 4.8 | 12.0 | 7.8 | 8.2 | 10.4 | 6.4 | 8.6 | 8.6 | 10.2 | 3.4 | 5.8 | 7.6 | 6.4 | 5.4 | 4.6 | 5.4 |
| $w^{1118/CS10}$ ♀ x *DJ694/+* ♂ | 1 | 04-03-16 | 1 | 12.8 | 13.8 | 11.0 | 9.2 | 7.6 | 7.6 | 5.4 | 8.6 | 7.8 | 11.4 | 6.6 | 7.4 | 6.8 | 4.6 | 7.4 | 5.6 | 5.4 | 5.2 | 2.8 | 4.2 |
| | | | 2 | 12.8 | 14.2 | 9.8 | 14.4 | 7.2 | 9.0 | 7.2 | 7.8 | 8.2 | 9.6 | 7.8 | 7.8 | 5.4 | 3.4 | 4.4 | 6.8 | 4.6 | 2.4 | 2.0 | 5.2 |
| | | | 3 | 9.2 | 11.4 | 13.2 | 5.2 | 7.2 | 7.2 | 8.4 | 7.6 | 9.2 | 9.0 | 8.2 | 6.6 | 6.4 | 2.8 | 7.8 | 5.8 | 6.6 | 4.8 | 5.6 | 3.8 |
| *DJ694* Virgins | 1 | 04-03-16 | 1 | 0.0 | 0.0 | 0.0 | 0.0 | 0.0 | 0.0 | 2.2 | 3.6 | 0.2 | 5.2 | 9.6 | 10.2 | 26.0 | 2.6 | 2.0 | 14.6 | 2.8 | 3.6 | 0.2 | 4.4 |
| | | | 2 | 0.0 | 0.0 | 0.0 | 0.0 | 0.0 | 0.0 | 0.0 | 4.2 | 9.2 | 10.2 | 13.4 | 14.4 | 8.0 | 1.4 | 14.4 | 0.6 | 4.0 | 7.0 | 2.0 | 8.2 |
| | | | 3 | 0.0 | 0.0 | 0.0 | 0.0 | 0.0 | 0.0 | 0.0 | 4.4 | 4.0 | 3.4 | 7.6 | 13.0 | 6.4 | 5.2 | 14.0 | 6.4 | 0.6 | 5.6 | 0.0 | 0.0 |
| *DJ694/+* Virgins | 1 | 04-03-16 | 1 | 0.0 | 0.0 | 0.0 | 0.0 | 0.0 | 0.0 | 0.0 | 3.0 | 0.0 | 1.8 | 7.2 | 8.8 | 8.0 | 0.0 | 0.5 | 24.0 | 1.5 | 0.0 | 0.0 | 10.5 |
| | | | 2 | 0.0 | 0.0 | 0.0 | 0.0 | 0.0 | 0.0 | 3.5 | 0.0 | 2.0 | 1.8 | 27.5 | 10.3 | 3.0 | 16.8 | 15.3 | 0.0 | 8.0 | 2.5 | 5.5 | 0.3 |
| | | | 3 | 0.0 | 0.0 | 0.0 | 0.0 | 0.0 | 6.6 | 3.8 | 0.0 | 10.8 | 5.8 | 16.4 | 5.8 | 8.6 | 2.8 | 9.8 | 0.2 | 16.4 | 3.0 | 0.0 | 2.8 |
| $w^{1118/CS10}$ Virgins | 1 | 04-03-16 | 1 | 0.0 | 0.0 | 0.0 | 1.0 | 0.0 | 1.4 | 5.6 | 5.8 | 7.2 | 10.6 | 7.6 | 9.2 | 3.4 | 1.6 | 6.8 | 2.2 | 5.0 | 2.4 | 0.6 | 1.8 |
| | | | 2 | 0.0 | 0.0 | 0.0 | 0.0 | 0.0 | 2.4 | 3.0 | 8.8 | 6.6 | 8.4 | 7.8 | 8.8 | 5.2 | 0.2 | 8.0 | 4.8 | 4.4 | 3.8 | 3.0 | 6.0 |
| | | | 3 | 0.0 | 0.0 | 0.0 | 0.0 | 0.0 | 0.0 | 0.0 | 2.2 | 1.0 | 9.2 | 9.8 | 0.0 | 14.8 | 0.6 | 6.4 | 3.2 | 4.6 | 4.6 | 3.4 | 1.8 |

| Cross | Rep | Start Date (YYMMDD) | Vial | Days 21 | 22 | 23 | 24 | 25 | 26 | 27 | 28 | 29 | 30 | 31 | 32 | 33 | 34 | 35 | 36 | 37 | 38 | 39 | 40 |
|---|---|---|---|---|---|---|---|---|---|---|---|---|---|---|---|---|---|---|---|---|---|---|---|
| *DJ694* ♀ x *DJ694* ♂ | 1 | 04-03-16 | 1 | 6.4 | 2.8 | 1.6 | 2.0 | 0.2 | 0.6 | 1.4 | 0.6 | 0.8 | 1.2 | 0.2 | 0.4 | 0.6 | 0.6 | 1.4 | 0.8 | 0.2 | 0.0 | 0.0 | 0.0 |
| | | | 2 | 0.8 | 6.6 | 1.6 | 6.4 | 11.4 | 1.8 | 4.0 | 3.2 | 1.4 | 4.6 | 0.4 | 1.0 | 4.6 | 2.0 | 1.6 | 1.4 | 0.6 | 1.8 | 0.6 | 1.0 |
| | | | 3 | 0.0 | 0.4 | 0.0 | 0.0 | 0.3 | 1.0 | 1.0 | 1.0 | 1.0 | 0.0 | x | x | x | x | x | x | x | x | x | x |
| | 2 | 05-02-20 | 1 | 8.4 | 3.4 | 6.0 | 4.2 | 4.4 | 5.0 | 3.0 | 4.8 | 2.8 | 8.2 | 2.4 | 3.4 | 1.0 | 1.0 | 1.6 | 2.8 | 1.5 | 1.0 | 0.5 | 0.3 |
| | | | 2 | 0.8 | 8.8 | 1.0 | 18.5 | 1.5 | 10.5 | 3.8 | 10.5 | 3.5 | 3.5 | 2.3 | 8.3 | 3.7 | 4.3 | 4.7 | 10.0 | 0.0 | 2.7 | 1.5 | 1.0 |
| | | | 3 | 1.8 | 5.2 | 4.8 | 2.6 | 6.2 | 5.0 | 3.2 | 2.8 | 2.8 | 4.2 | 5.2 | 2.0 | 2.0 | 1.5 | 4.5 | 2.5 | 1.0 | 1.0 | 0.8 | 0.5 |
| $w^{1118/CS10}$ ♀ x *DJ694* ♂ | 1 | 04-03-16 | 1 | 5.4 | 3.2 | 4.0 | 3.6 | 7.6 | 2.8 | 4.0 | 4.0 | 3.2 | 2.4 | 4.0 | 0.5 | 6.5 | 4.0 | 4.0 | 2.8 | 2.8 | 1.0 | 0.7 | 5.3 |
| | | | 2 | 7.2 | 5.4 | 6.4 | 5.4 | 7.0 | 5.0 | 6.4 | 5.0 | 4.8 | 7.0 | 6.0 | 3.4 | 6.6 | 4.2 | 4.2 | 4.8 | 5.6 | 2.8 | 2.8 | 3.6 |
| | | | 3 | 2.2 | 2.8 | 3.2 | 2.0 | 3.0 | 3.2 | 2.2 | 2.6 | 2.4 | 2.2 | 4.0 | 1.3 | 2.3 | 2.3 | 1.5 | 2.0 | 1.5 | 2.0 | 1.3 | 1.8 |
| | 2 | 05-02-20 | 1 | 9.2 | 6.6 | 4.4 | 8.4 | 11.4 | 9.0 | 10.4 | 7.8 | 6.8 | 9.4 | 6.4 | 4.0 | 8.0 | 4.4 | 3.0 | 9.4 | 2.4 | 4.0 | 2.8 | 5.0 |
| | | | 2 | 9.8 | 10.0 | 9.5 | 9.0 | 9.8 | 5.0 | 15.3 | 7.5 | 6.5 | 9.8 | 10.8 | 2.8 | 9.5 | 6.8 | 7.5 | 5.0 | 4.0 | 7.3 | 1.8 | 2.8 |
| | | | 3 | 10.8 | 8.4 | 9.0 | 7.0 | 8.8 | 5.4 | 12.2 | 5.6 | 10.8 | 6.4 | 7.2 | 5.4 | 6.4 | 5.0 | 6.4 | 4.2 | 4.2 | 4.4 | 3.4 | 1.8 |
| *DJ694* ♀ x $w^{1118/CS10}$ ♂ | 1 | 04-03-16 | 1 | 1.5 | 1.8 | 0.5 | 0.5 | 0.8 | 0.5 | 0.8 | 0.3 | 1.0 | 1.3 | 0.7 | 0.7 | 1.0 | 0.3 | 1.0 | 0.0 | 0.0 | 0.0 | 0.0 | 0.0 |
| | | | 2 | 4.0 | 1.4 | 1.0 | 6.4 | 1.2 | 1.4 | 1.8 | 1.6 | 0.6 | 1.6 | 1.4 | 0.6 | 2.6 | 0.4 | 0.8 | 1.0 | 2.2 | 0.8 | 0.6 | 0.6 |
| | | | 3 | 8.8 | 4.8 | 0.2 | 3.2 | 9.0 | 0.6 | 0.4 | 0.8 | 4.8 | 1.4 | 0.8 | 0.8 | 3.4 | 4.4 | 0.4 | 0.8 | 0.4 | 1.2 | 2.6 | 0.5 |
| | 2 | 05-02-20 | 1 | 11.0 | 1.8 | 2.8 | 7.8 | 5.2 | 2.6 | 9.2 | 1.4 | 3.8 | 5.2 | 3.0 | 2.6 | 0.8 | 1.6 | 0.8 | 0.4 | 0.2 | 0.0 | 0.0 | 0.0 |
| | | | 2 | 10.4 | 4.8 | 5.6 | 11.6 | 2.6 | 1.0 | 9.6 | 3.2 | 2.4 | 4.2 | 5.6 | 1.8 | 1.0 | 3.6 | 2.6 | 0.4 | 0.4 | 0.8 | 0.2 | 0.4 |
| | | | 3 | 14.3 | 2.5 | 3.3 | 8.8 | 4.8 | 8.7 | 3.3 | 2.7 | 3.7 | 3.7 | 14.3 | 0.3 | 7.7 | 2.0 | 5.0 | 0.3 | 5.7 | 0.3 | 0.3 | 0.0 |
| $w^{1118/CS10}$ ♀ x $w^{1118}$ ♂ | 1 | 04-03-16 | 1 | 5.2 | 2.8 | 3.0 | 4.2 | 6.4 | 3.2 | 4.2 | 6.2 | 1.8 | 5.0 | 3.8 | 0.4 | 3.6 | 2.0 | 2.6 | 2.8 | 3.6 | 2.6 | 1.4 | 1.6 |
| | | | 2 | 1.4 | 6.2 | 3.8 | 5.6 | 6.8 | 3.4 | 5.4 | 2.8 | 9.8 | 7.8 | 10.0 | 5.3 | 8.3 | 3.5 | 3.0 | 4.8 | 3.0 | 6.3 | 6.0 | 3.8 |
| | | | 3 | 3.2 | 5.0 | 3.8 | 5.6 | 7.2 | 6.3 | 8.0 | 4.0 | 10.7 | 7.7 | 9.3 | 5.3 | 8.0 | 6.3 | 7.0 | 3.7 | 7.0 | 7.3 | 3.3 | 4.7 |
| | 2 | 05-02-20 | 1 | 9.2 | 8.6 | 5.6 | 9.0 | 5.4 | 8.8 | 8.8 | 2.4 | 9.0 | 7.8 | 3.6 | 6.4 | 5.2 | 4.2 | 8.8 | 1.8 | 7.8 | 2.4 | 2.0 | 2.0 |
| | | | 2 | 11.0 | 5.4 | 6.0 | 8.0 | 10.6 | 6.8 | 5.4 | 15.0 | 6.4 | 7.8 | 6.8 | 7.2 | 6.2 | 7.4 | 2.4 | 2.0 | 7.0 | 3.8 | 4.0 | 1.0 |
| | | | 3 | 8.4 | 5.2 | 9.6 | 9.0 | 6.0 | 6.0 | 6.2 | 10.4 | 7.0 | 6.8 | 7.0 | 4.8 | 8.0 | 3.4 | 8.4 | 2.2 | 7.8 | 2.4 | 1.6 | 0.8 |
| *DJ694/+* ♀ x $w^{1118/CS10}$ ♂ | 1 | 04-03-16 | 1 | 4.8 | 5.8 | 8.8 | 3.6 | 7.8 | 3.4 | 4.8 | 3.4 | 6.8 | 5.2 | 5.6 | 5.4 | 7.0 | 7.0 | 4.8 | 5.4 | 0.8 | 5.0 | 6.6 | 3.0 |
| | | | 2 | 12.0 | 6.0 | 9.3 | 5.5 | 10.0 | 8.5 | 4.8 | 3.8 | 11.8 | 10.5 | 13.5 | 2.5 | 15.0 | 8.0 | 5.0 | 4.8 | 5.5 | 5.3 | 5.7 | 12.3 |
| | | | 3 | 3.4 | 5.4 | 4.8 | 7.0 | 7.0 | 5.0 | 5.2 | 3.2 | 7.2 | 6.0 | 7.4 | 2.8 | 8.0 | 5.4 | 7.3 | 6.3 | 4.0 | 5.8 | 8.0 | 5.0 |
| $w^{1118/CS10}$ ♀ x *DJ694/+* ♂ | 1 | 04-03-16 | 1 | 4.2 | 2.8 | 6.4 | 6.8 | 9.2 | 6.4 | 4.8 | 4.0 | 8.5 | 9.5 | 8.3 | 5.5 | 7.3 | 4.8 | 6.3 | 6.0 | 3.8 | 7.5 | 4.0 | 3.5 |
| | | | 2 | 3.4 | 4.6 | 2.4 | 6.2 | 4.2 | 3.6 | 4.4 | 0.8 | 4.6 | 2.4 | 3.2 | 1.4 | 11.3 | 3.3 | 5.3 | 3.0 | 3.7 | 1.7 | 0.7 | 1.3 |
| | | | 3 | 2.6 | 5.6 | 6.0 | 3.8 | 6.4 | 4.0 | 6.6 | 3.6 | 7.0 | 6.0 | 4.6 | 2.6 | 4.8 | 3.2 | 3.0 | 3.4 | 3.4 | 2.6 | 3.5 | 3.5 |
| *DJ694* Virgins | 1 | 04-03-16 | 1 | 0.6 | 4.8 | 2.6 | 1.4 | 10.4 | 0.6 | 4.0 | 0.8 | 0.6 | 6.8 | 1.2 | 0.6 | 0.6 | 1.2 | 10.8 | 1.4 | 1.2 | 0.8 | 1.6 | 0.6 |
| | | | 2 | 0.8 | 5.6 | 1.4 | 2.4 | 6.8 | 0.4 | 6.8 | 4.8 | 3.4 | 0.6 | 0.2 | 0.8 | 1.6 | 3.4 | 4.6 | 3.4 | 0.6 | 0.2 | 0.2 | 0.2 |
| | | | 3 | 3.2 | 10.6 | 4.0 | 3.5 | 7.5 | 5.0 | 5.8 | 0.3 | 5.8 | 0.3 | 8.8 | 2.0 | 7.0 | 4.0 | 4.5 | 2.0 | 0.0 | 0.8 | 6.5 | 0.8 |
| *DJ694/+* Virgins | 1 | 04-03-16 | 1 | 6.0 | 0.0 | 22.0 | 0.0 | 23.0 | 2.0 | 6.0 | 6.0 | 9.0 | 1.0 | 0.0 | 0.0 | 18.0 | 6.0 | 5.0 | 0.0 | 0.0 | 21.0 | 2.0 | 1.0 |
| | | | 2 | 8.8 | 2.5 | 1.3 | 9.5 | 9.3 | 8.8 | 1.5 | 3.5 | 7.3 | 6.5 | 7.3 | 4.0 | 4.8 | 7.0 | 1.8 | 4.3 | 1.8 | 2.8 | 6.8 | 1.5 |
| | | | 3 | 2.4 | 7.6 | 0.4 | 3.8 | 4.4 | 3.8 | 4.4 | 4.2 | 3.2 | 7.8 | 5.0 | 3.2 | 6.8 | 3.6 | 5.2 | 4.0 | 2.8 | 2.8 | 1.0 | 6.2 |
| $w^{1118/CS10}$ Virgins | 1 | 04-03-16 | 1 | 0.8 | 3.6 | 1.6 | 6.0 | 9.0 | 6.4 | 3.2 | 4.0 | 6.2 | 1.6 | 3.4 | 3.8 | 6.0 | 4.4 | 7.6 | 2.8 | 5.4 | 1.6 | 0.8 | 4.8 |
| | | | 2 | 1.2 | 4.4 | 3.0 | 4.6 | 6.6 | 1.8 | 3.0 | 2.8 | 5.6 | 3.4 | 2.8 | 3.8 | 3.4 | 2.4 | 6.0 | 2.2 | 3.2 | 3.2 | 2.8 | 2.4 |
| | | | 3 | 0.4 | 0.8 | 1.0 | 8.8 | 7.6 | 4.6 | 2.4 | 2.4 | 4.8 | 0.8 | 6.5 | 3.8 | 12.5 | 5.0 | 3.5 | 4.3 | 4.3 | 1.3 | 3.3 | 1.8 |

| Cross | Rep | Start Date (YYMMDD) | Vial | 41 | 42 | 43 | 44 | 45 | 46 | 47 | 48 | 49 | 50 | 51 | 52 | 53 | 54 | 55 | 56 | 57 | 58 | 59 | 60 |
|---|---|---|---|---|---|---|---|---|---|---|---|---|---|---|---|---|---|---|---|---|---|---|---|
| *DJ694*♀ x *DJ694*♂ | 1 | 04-03-16 | 1 | 0.2 | 0.0 | 0.0 | 0.0 | 0.0 | 0.0 | 0.0 | 0.0 | 0.0 | 0.0 | 0.0 | 0.0 | 0.0 | 0.0 | 0.0 | 0.0 | x | x | x | x |
| | | | 2 | 1.2 | 0.8 | 0.8 | 0.8 | 0.8 | 0.3 | 0.5 | 0.3 | 0.3 | 0.3 | 0.0 | 0.0 | 0.0 | 0.0 | 0.0 | 0.0 | 0.0 | 0.0 | 0.0 | x |
| | | | 3 | x | x | x | x | x | x | x | x | x | x | x | x | x | x | x | x | x | x | x | x |
| | 2 | 05-02-20 | 1 | 1.3 | 0.3 | 0.7 | 0.3 | 0.0 | 0.5 | 2.0 | 0.5 | 0.0 | 0.0 | 0.5 | 0.5 | 0.5 | 0.0 | 0.0 | x | x | x | x | x |
| | | | 2 | 0.5 | 0.5 | 0.0 | 1.0 | 0.0 | 0.5 | 1.0 | 0.0 | 0.5 | 0.0 | 0.0 | 0.0 | 0.0 | 0.0 | 0.0 | x | x | x | x | x |
| | | | 3 | 0.3 | 0.5 | 0.5 | 0.0 | 0.3 | 1.0 | 2.3 | 0.5 | 0.5 | 0.3 | 0.0 | 0.0 | 1.0 | 0.0 | 0.0 | 0.0 | 0.0 | 0.0 | x | x |
| $w^{1118/CS10}$♀ x *DJ694* ♂ | 1 | 04-03-16 | 1 | 1.0 | 1.0 | 4.0 | 0.0 | 7.7 | 12.0 | 3.7 | 7.0 | 3.7 | 3.3 | 5.5 | 4.0 | 1.0 | 4.0 | 2.5 | 1.5 | 0.0 | 7.0 | 6.0 | 4.5 |
| | | | 2 | 1.4 | 2.4 | 5.2 | 2.0 | 4.2 | 1.4 | 0.4 | 1.0 | 0.6 | 0.0 | 1.0 | 1.0 | 0.3 | 0.8 | 3.0 | 0.0 | 2.5 | 3.0 | 2.0 | 1.5 |
| | | | 3 | 2.5 | 5.0 | 1.8 | 0.0 | 0.0 | 1.8 | 1.7 | 0.0 | 1.3 | 0.0 | 0.0 | 1.7 | 0.0 | 0.0 | 2.5 | 0.0 | 1.5 | 0.5 | 0.0 | 0.0 |
| | 2 | 05-02-20 | 1 | 4.2 | 2.0 | 0.2 | 4.4 | 3.6 | 2.0 | 1.8 | 3.6 | 1.4 | 3.2 | 0.4 | 1.4 | 0.8 | 0.0 | 2.4 | 0.2 | 0.0 | 4.0 | 3.2 | 1.4 |
| | | | 2 | 4.3 | 3.5 | 6.8 | 8.0 | 2.5 | 8.8 | 3.5 | 2.3 | 1.7 | 0.0 | 0.0 | 10.3 | 7.7 | 3.7 | 13.7 | 4.7 | 0.3 | 2.3 | 7.0 | 1.0 |
| | | | 3 | 3.8 | 1.6 | 1.6 | 3.4 | 3.4 | 2.6 | 4.6 | 2.0 | 3.4 | 3.4 | 2.2 | 3.4 | 5.2 | 1.6 | 6.4 | 4.2 | 0.8 | 2.4 | 3.6 | 3.2 |
| *DJ694*♀ x $w^{1118/CS10}$ ♂ | 1 | 04-03-16 | 1 | 0.0 | 0.0 | 0.0 | 0.0 | 0.0 | 0.0 | 0.0 | x | x | x | x | x | x | x | x | x | x | x | x | x |
| | | | 2 | 0.2 | 0.2 | 0.0 | 0.0 | 0.0 | 0.0 | 0.0 | 0.0 | 0.0 | 0.0 | 0.0 | 0.0 | 0.0 | x | x | x | x | x | x | x |
| | | | 3 | 0.7 | 2.0 | 1.3 | 0.0 | 0.0 | 0.0 | 1.5 | 0.0 | 1.0 | 1.0 | 0.5 | 0.0 | 0.0 | 0.0 | 0.0 | 0.0 | x | x | x | x |
| | 2 | 05-02-20 | 1 | 0.0 | 0.0 | 0.0 | 0.0 | 0.0 | 0.0 | 0.0 | 0.0 | 0.0 | 0.0 | 0.0 | 0.0 | 0.0 | 0.0 | x | x | x | x | x | x |
| | | | 2 | 0.0 | 0.0 | 0.0 | 0.0 | 0.0 | 0.0 | 0.0 | 0.0 | 0.0 | 0.0 | 0.0 | 0.0 | 0.0 | 0.0 | 0.0 | 0.0 | 0.0 | 0.0 | x | x |
| | | | 3 | 0.0 | 0.0 | 0.0 | 0.0 | 0.0 | 0.0 | 0.0 | 0.0 | 0.0 | 0.0 | 0.0 | x | x | x | x | x | x | x | x | x |
| $w^{1118/CS10}$♀ x $w^{1118}$♂ | 1 | 04-03-16 | 1 | 2.2 | 2.6 | 3.0 | 0.8 | 1.4 | 1.0 | 0.8 | 0.2 | 0.3 | 0.5 | 0.0 | 0.0 | 0.0 | 0.0 | 0.3 | 0.0 | 0.0 | 2.5 | 0.0 | 0.5 |
| | | | 2 | 4.0 | 3.5 | 3.5 | 5.8 | 5.3 | 7.0 | 3.3 | 1.8 | 5.0 | 1.5 | 1.5 | 0.5 | 1.5 | 1.5 | 4.5 | 1.3 | 3.7 | 5.7 | 11.0 | 5.0 |
| | | | 3 | 5.0 | 5.7 | 10.3 | 7.0 | 6.7 | 11.5 | 12.0 | 24.0 | 23.0 | 12.0 | 15.0 | 9.0 | 5.0 | 5.0 | 11.0 | 5.0 | 16.0 | 8.0 | 10.0 | 6.0 |
| | 2 | 05-02-20 | 1 | 6.8 | 2.4 | 0.8 | 6.4 | 4.0 | 4.2 | 8.2 | 5.0 | 7.8 | 7.6 | 4.8 | 7.0 | 7.8 | 5.0 | 4.0 | 6.0 | 1.0 | 4.0 | 7.0 | 6.7 |
| | | | 2 | 6.0 | 3.0 | 2.6 | 2.4 | 3.6 | 6.4 | 5.2 | 4.0 | 4.4 | 7.8 | 8.6 | 6.8 | 5.8 | 3.8 | 5.2 | 3.4 | 1.8 | 4.4 | 2.6 | 3.8 |
| | | | 3 | 5.8 | 5.8 | 2.2 | 7.0 | 6.0 | 6.0 | 8.2 | 6.0 | 7.8 | 8.8 | 8.6 | 4.6 | 5.4 | 4.8 | 4.2 | 3.2 | 0.8 | 6.0 | 3.0 | 3.8 |
| *DJ694/+*♀ x $w^{1118/CS10}$♂ | 1 | 04-03-16 | 1 | 2.2 | 5.0 | 7.2 | 2.8 | 2.8 | 5.2 | 5.2 | 4.2 | 4.6 | 1.0 | 5.8 | 1.8 | 1.4 | 4.8 | 3.6 | 2.4 | 3.8 | 3.8 | 3.2 | 9.8 |
| | | | 2 | 3.7 | 9.3 | 12.7 | 6.0 | 6.3 | 9.0 | 8.0 | 9.3 | 4.7 | 7.7 | 6.3 | 0.3 | 3.3 | 6.0 | 2.3 | 3.3 | 14.7 | 6.3 | 7.0 | 13.3 |
| | | | 3 | 7.7 | 9.7 | 12.7 | 7.7 | 5.3 | 2.7 | 14.3 | 5.0 | 8.3 | 7.0 | 8.7 | 3.3 | 3.0 | 4.0 | 7.0 | 2.7 | 9.3 | 4.7 | 8.0 | 6.3 |
| $w^{1118/CS10}$♀ x *DJ694/+*♂ | 1 | 04-03-16 | 1 | 1.5 | 7.0 | 9.5 | 1.8 | 6.5 | 7.5 | 6.3 | 6.8 | 5.5 | 5.5 | 4.3 | 4.8 | 3.5 | 2.8 | 4.3 | 2.5 | 4.5 | 2.5 | 5.7 | 5.7 |
| | | | 2 | 1.3 | 1.7 | 1.3 | 2.3 | 1.7 | 1.7 | 0.3 | 0.3 | 1.3 | 0.7 | 0.3 | 0.0 | 0.3 | 0.3 | 0.5 | 0.0 | 0.0 | 0.0 | 0.0 | 0.0 |
| | | | 3 | 2.0 | 4.8 | 6.0 | 2.3 | 6.0 | 5.3 | 3.3 | 6.0 | 5.0 | 3.3 | 4.0 | 0.5 | 3.5 | 3.5 | 5.8 | 1.0 | 5.5 | 3.8 | 6.0 | 3.3 |
| *DJ694* Virgins | 1 | 04-03-16 | 1 | 3.8 | 1.8 | 7.2 | 3.0 | 3.8 | 4.3 | 3.5 | 1.0 | 5.8 | 2.3 | 2.3 | 4.0 | 2.0 | 5.3 | 5.7 | 2.3 | 2.7 | 4.3 | 4.0 | 3.0 |
| | | | 2 | 2.0 | 2.0 | 7.4 | 0.8 | 0.0 | 0.8 | 0.4 | 1.4 | 3.2 | 3.2 | 3.2 | 0.2 | 1.0 | 1.0 | 0.0 | 0.8 | 1.7 | 3.7 | 7.0 | 4.5 |
| | | | 3 | 1.8 | 2.3 | 8.0 | 3.0 | 4.5 | 1.3 | 1.8 | 0.5 | 2.5 | 2.3 | 2.3 | 0.3 | 3.7 | 2.0 | 3.0 | 5.5 | 6.5 | 10.5 | 9.0 | 5.5 |
| *DJ694/+* Virgins | 1 | 04-03-16 | 1 | 0.0 | 11.0 | 6.0 | 8.0 | 14.0 | 12.0 | 4.0 | 12.0 | 0.0 | 10.0 | 4.0 | 5.0 | 8.0 | 5.0 | 3.0 | 6.0 | 2.0 | 14.0 | 13.0 | 8.0 |
| | | | 2 | 3.8 | 6.3 | 6.3 | 4.5 | 3.0 | 9.5 | 6.0 | 4.0 | 1.3 | 4.8 | 4.0 | 2.0 | 6.3 | 4.3 | 2.8 | 4.0 | 6.0 | 5.0 | 6.8 | 6.3 |
| | | | 3 | 3.6 | 4.0 | 5.4 | 5.2 | 3.0 | 5.0 | 4.4 | 5.4 | 1.8 | 4.4 | 3.6 | 4.2 | 4.2 | 3.6 | 5.0 | 3.4 | 4.4 | 5.4 | 5.6 | 4.8 |
| $w^{1118/CS10}$ Virgins | 1 | 04-03-16 | 1 | 3.4 | 4.2 | 4.0 | 0.8 | 2.6 | 6.8 | 1.6 | 8.0 | 2.2 | 2.0 | 0.8 | 0.8 | 1.6 | 3.4 | 4.6 | 2.3 | 4.8 | 4.5 | 4.0 | 5.5 |
| | | | 2 | 3.0 | 4.8 | 7.4 | 2.4 | 2.8 | 2.8 | 1.2 | 3.6 | 4.0 | 0.5 | 2.8 | 0.3 | 4.0 | 1.7 | 4.0 | 0.0 | 0.5 | 2.0 | 2.0 | 1.5 |
| | | | 3 | 1.8 | 7.0 | 5.5 | 6.0 | 4.0 | 5.5 | 2.3 | 7.8 | 3.5 | 2.8 | 1.8 | 2.8 | 4.3 | 4.8 | 2.8 | 4.0 | 4.3 | 4.5 | 6.3 | 7.0 |

| Cross | Rep | Start Date (YYMMDD) | Vial | 61 | 62 | 63 | 64 | 65 | 66 | 67 | 68 | 69 | 70 | 71 | 72 | 73 | 74 | 75 | 76 | 77 | 78 | 79 | 80 |
|---|---|---|---|---|---|---|---|---|---|---|---|---|---|---|---|---|---|---|---|---|---|---|---|
| | | | | Days | | | | | | | | | | | | | | | | | | | |
| *DJ694* ♀ x *DJ694* ♂ | 1 | 04-03-16 | 1 | x | x | x | x | x | x | x | x | x | x | x | x | x | x | x | x | x | x | x | x |
| | | | 2 | x | x | x | x | x | x | x | x | x | x | x | x | x | x | x | x | x | x | x | x |
| | | | 3 | x | x | x | x | x | x | x | x | x | x | x | x | x | x | x | x | x | x | x | x |
| | 2 | 05-02-20 | 1 | x | x | x | x | x | x | x | x | x | x | x | x | x | x | x | x | x | x | x | x |
| | | | 2 | x | x | x | x | x | x | x | x | x | x | x | x | x | x | x | x | x | x | x | x |
| | | | 3 | x | x | x | x | x | x | x | x | x | x | x | x | x | x | x | x | x | x | x | x |
| $w^{1118/CS10}$ ♀ x *DJ694* ♂ | 1 | 04-03-16 | 1 | 0.0 | 8.0 | 6.0 | 1.0 | 0.0 | 0.0 | 0.0 | 0.0 | x | x | x | x | x | x | x | x | x | x | x | x |
| | | | 2 | 0.0 | 0.0 | 0.0 | 0.0 | 0.0 | 0.0 | 0.0 | 0.0 | 0.0 | 0.0 | 0.0 | 0.0 | x | x | x | x | x | x | x | x |
| | | | 3 | 0.0 | 1.0 | 0.0 | 0.0 | 0.0 | 0.5 | 2.0 | 0.5 | 0.0 | 0.0 | 0.0 | 0.0 | 0.0 | 0.0 | 0.0 | 0.0 | 0.0 | 0.0 | 0.0 | 0.0 |
| | 2 | 05-02-20 | 1 | 2.0 | 1.6 | 2.8 | 1.6 | 1.3 | 0.3 | 0.0 | 0.0 | 0.5 | 6.0 | 4.0 | 2.5 | 1.0 | 4.0 | 0.0 | 0.0 | 4.0 | 0.0 | 1.0 | 0.0 |
| | | | 2 | 3.3 | 9.7 | 6.0 | 1.3 | 8.7 | 0.3 | 1.0 | 0.0 | 0.3 | 3.0 | 1.3 | 0.3 | 2.3 | 1.3 | 0.3 | 1.5 | 1.5 | 0.0 | 0.0 | 0.0 |
| | | | 3 | 2.2 | 2.2 | 4.4 | 4.8 | 1.4 | 2.2 | 0.2 | 0.2 | 2.2 | 1.6 | 1.0 | 0.8 | 1.0 | 0.4 | 1.0 | 0.2 | 0.0 | 0.2 | 0.0 | 0.0 |
| *DJ694* ♀ x $w^{1118/CS10}$ ♂ | 1 | 04-03-16 | 1 | x | x | x | x | x | x | x | x | x | x | x | x | x | x | x | x | x | x | x | x |
| | | | 2 | x | x | x | x | x | x | x | x | x | x | x | x | x | x | x | x | x | x | x | x |
| | | | 3 | x | x | x | x | x | x | x | x | x | x | x | x | x | x | x | x | x | x | x | x |
| | 2 | 05-02-20 | 1 | x | x | x | x | x | x | x | x | x | x | x | x | x | x | x | x | x | x | x | x |
| | | | 2 | x | x | x | x | x | x | x | x | x | x | x | x | x | x | x | x | x | x | x | x |
| | | | 3 | x | x | x | x | x | x | x | x | x | x | x | x | x | x | x | x | x | x | x | x |
| $w^{1118/CS10}$ ♀ x $w^{1118}$ ♂ | 1 | 04-03-16 | 1 | 0.0 | 0.0 | 0.0 | 0.0 | 0.0 | 0.0 | 0.0 | 0.0 | 0.0 | 0.0 | 0.0 | 0.0 | 0.0 | 0.0 | 0.0 | 0.0 | 7.0 | 0.0 | 0.0 | 0.0 |
| | | | 2 | 6.5 | 8.0 | 4.0 | 1.0 | 0.0 | 0.0 | x | x | x | x | x | x | x | x | x | x | x | x | x | x |
| | | | 3 | 23.0 | 10.0 | 7.0 | 11.0 | 14.0 | 10.0 | 4.0 | 25.0 | 5.0 | 2.0 | 11.0 | 6.0 | 2.0 | 1.0 | 4.0 | 0.0 | 1.0 | 0.0 | 0.0 | 0.0 |
| | 2 | 05-02-20 | 1 | 4.3 | 4.7 | 5.3 | 4.3 | 2.5 | 1.5 | 0.5 | 1.5 | 1.0 | 0.5 | 1.0 | 1.0 | 0.0 | 0.0 | 0.0 | 0.0 | 0.0 | 0.0 | 0.0 | 0.0 |
| | | | 2 | 3.4 | 2.2 | 3.8 | 1.3 | 0.3 | 0.3 | 1.0 | 0.3 | 0.0 | 0.0 | 0.0 | 0.0 | 0.0 | 0.0 | 0.0 | 0.0 | 0.0 | 0.0 | 0.0 | 0.0 |
| | | | 3 | 4.0 | 3.4 | 2.6 | 1.8 | 0.8 | 1.2 | 1.8 | 0.6 | 0.8 | 0.6 | 0.0 | 0.0 | 0.5 | 0.3 | 0.0 | 0.0 | 0.0 | 0.0 | 0.0 | 0.0 |
| *DJ694/+* ♀ x $w^{1118/CS10}$ ♂ | 1 | 04-03-16 | 1 | 2.8 | 7.5 | 3.0 | 8.3 | 2.8 | 7.5 | 5.0 | 6.3 | 3.0 | 0.8 | 3.0 | 5.3 | 2.0 | 1.8 | 0.5 | 1.3 | 0.3 | 0.0 | 0.5 | 0.3 |
| | | | 2 | 4.7 | 3.7 | 3.7 | 9.0 | 4.7 | 3.7 | 1.0 | 8.7 | 1.0 | 0.3 | 0.7 | 0.7 | 1.5 | 0.5 | 3.0 | 0.0 | 0.0 | 0.0 | 0.0 | 0.0 |
| | | | 3 | 9.7 | 12.0 | 6.7 | 5.3 | 3.7 | 7.0 | 3.7 | 4.3 | 0.3 | 0.3 | 2.3 | 3.0 | 0.3 | 1.0 | 0.3 | 0.3 | 0.0 | 0.0 | 0.0 | 9.0 |
| $w^{1118/CS10}$ ♀ x *DJ694/+* ♂ | 1 | 04-03-16 | 1 | 4.3 | 6.0 | 5.3 | 1.0 | 8.3 | 6.3 | 3.3 | 2.7 | 0.7 | 0.3 | 2.3 | 0.0 | 0.0 | 3.0 | 0.5 | 0.0 | 1.0 | 0.0 | 0.0 | 0.0 |
| | | | 2 | 0.0 | 0.0 | 0.0 | 0.0 | 0.0 | 0.0 | 0.0 | 0.0 | x | x | x | x | x | x | x | x | x | x | x | x |
| | | | 3 | 6.3 | 4.7 | 4.3 | 5.7 | 1.3 | 2.3 | 0.0 | 3.0 | 1.7 | 0.0 | 1.0 | 0.3 | 0.3 | 0.3 | 0.0 | 0.0 | 0.0 | 0.0 | 0.7 | 1.0 |
| *DJ694* Virgins | 1 | 04-03-16 | 1 | 5.0 | 3.0 | 1.0 | 2.0 | 6.5 | 3.0 | 4.5 | 6.5 | 1.5 | 5.5 | 1.5 | 2.5 | 2.0 | 1.5 | 1.5 | 2.5 | 2.0 | 1.5 | 0.0 | 0.0 |
| | | | 2 | 3.0 | 5.0 | 1.0 | 2.0 | 1.0 | 0.0 | x | x | x | x | x | x | x | x | x | x | x | x | x | x |
| | | | 3 | 9.5 | 6.0 | 1.0 | 4.0 | 6.0 | 7.0 | 5.0 | 12.0 | 5.0 | 6.0 | 3.0 | 0.0 | 6.0 | 1.0 | 0.0 | 1.0 | 0.0 | 1.0 | 0.0 | 1.0 |
| *DJ694/+* Virgins | 1 | 04-03-16 | 1 | 13.0 | 15.0 | 8.0 | 9.0 | 13.0 | 11.0 | 11.0 | 9.0 | 4.0 | 6.0 | 6.0 | 12.0 | 6.0 | 9.0 | 7.0 | 5.0 | 0.0 | 10.0 | 10.0 | 6.0 |
| | | | 2 | 8.3 | 8.3 | 5.0 | 4.3 | 3.5 | 6.8 | 4.8 | 5.3 | 2.8 | 2.5 | 5.0 | 3.8 | 2.7 | 3.3 | 4.3 | 3.3 | 1.7 | 2.0 | 3.0 | 0.3 |
| | | | 3 | 8.6 | 8.0 | 5.2 | 7.6 | 5.0 | 8.2 | 6.2 | 6.4 | 6.8 | 3.0 | 6.3 | 2.0 | 5.0 | 4.8 | 4.8 | 2.8 | 3.3 | 1.8 | 2.8 | 1.0 |
| $w^{1118/CS10}$ Virgins | 1 | 04-03-16 | 1 | 4.0 | 3.8 | 2.5 | 3.3 | 1.8 | 4.3 | 2.0 | 2.3 | 0.0 | 0.8 | 1.8 | 0.0 | 0.3 | 1.0 | 0.0 | 0.0 | 0.0 | 0.0 | 0.0 | 0.0 |
| | | | 2 | 1.5 | 2.5 | 1.0 | 3.5 | 2.0 | 0.0 | 0.0 | 3.0 | 0.0 | 0.0 | 0.0 | 0.0 | 0.0 | x | x | x | x | x | x | x |
| | | | 3 | 9.0 | 8.3 | 5.3 | 8.3 | 8.0 | 6.3 | 4.7 | 4.3 | 3.3 | 1.3 | 5.0 | 2.7 | 3.7 | 2.7 | 2.3 | 1.7 | 1.0 | 0.7 | 1.3 | 0.7 |

| Cross | Rep | Start Date (YYMMDD) | Vial | Days 81 | 82 | 83 | 84 | 85 |
|---|---|---|---|---|---|---|---|---|
| *DJ694* ♀ x *DJ694* ♂ | 1 | 04-03-16 | 1 | x | x | x | x | x |
| | | | 2 | x | x | x | x | x |
| | | | 3 | x | x | x | x | x |
| | 2 | 05-02-20 | 1 | x | x | x | x | x |
| | | | 2 | x | x | x | x | x |
| | | | 3 | x | x | x | x | x |
| $w^{1118/CS10}$ ♀ x *DJ694* ♂ | 1 | 04-03-16 | 1 | x | x | x | x | x |
| | | | 2 | x | x | x | x | x |
| | | | 3 | 0.0 | 0.0 | 0.0 | 0.0 | x |
| | 2 | 05-02-20 | 1 | 0.0 | 0.0 | 0.0 | 0.0 | 0.0 |
| | | | 2 | 0.0 | 0.0 | 0.0 | 0.0 | 0.0 |
| | | | 3 | 0.0 | x | x | x | x |
| *DJ694* ♀ x $w^{1118/CS10}$ ♂ | 1 | 04-03-16 | 1 | x | x | x | x | x |
| | | | 2 | x | x | x | x | x |
| | | | 3 | x | x | x | x | x |
| | 2 | 05-02-20 | 1 | x | x | x | x | x |
| | | | 2 | x | x | x | x | x |
| | | | 3 | x | x | x | x | x |
| $w^{1118/CS10}$ ♀ x $w^{1118}$ ♂ | 1 | 04-03-16 | 1 | 0.0 | 0.0 | 0.0 | 0.0 | x |
| | | | 2 | x | x | x | x | x |
| | | | 3 | 0.0 | 0.0 | 0.0 | 0.0 | x |
| | 2 | 05-02-20 | 1 | 0.0 | 0.0 | 0.0 | 0.0 | x |
| | | | 2 | x | x | x | x | x |
| | | | 3 | 0.0 | 0.0 | 0.0 | 0.0 | x |
| *DJ694/+* ♀ x $w^{1118/CS10}$ ♂ | 1 | 04-03-16 | 1 | 0.0 | 0.0 | 0.0 | 0.0 | 0.0 |
| | | | 2 | 0.0 | 0.0 | x | x | x |
| | | | 3 | 1.0 | 0.0 | 2.0 | 1.0 | 0.0 |
| $w^{1118/CS10}$ ♀ x *DJ694/+* ♂ | 1 | 04-03-16 | 1 | 2.0 | 0.0 | 0.0 | 0.0 | 0.0 |
| | | | 2 | x | x | x | x | x |
| | | | 3 | 0.0 | 0.3 | 0.0 | 0.0 | 0.0 |
| *DJ694* Virgins | 1 | 04-03-16 | 1 | 2.0 | 0.5 | 1.5 | 0.5 | 0.0 |
| | | | 2 | x | x | x | x | x |
| | | | 3 | 1.0 | 1.0 | 1.0 | 1.0 | 0.0 |
| *DJ694/+* Virgins | 1 | 04-03-16 | 1 | 3.0 | 1.0 | 0.0 | 0.0 | 0.0 |
| | | | 2 | 1.0 | 0.7 | 1.0 | 0.3 | 0.0 |
| | | | 3 | 1.0 | 0.0 | 0.0 | 0.0 | 0.0 |
| $w^{1118/CS10}$ Virgins | 1 | 04-03-16 | 1 | 0.0 | 0.0 | 0.0 | 0.0 | 0.0 |
| | | | 2 | x | x | x | x | x |
| | | | 3 | 0.3 | 0.5 | 1.5 | 0.0 | 0.0 |

| Cross | Rep | Start Date (YYMMDD) | Vial | Days 29 | 30 | 31 | 32 | 33 | 34 | 35 | 36 | 37 | 38 | 39 | 40 | 41 | 42 | 43 | 44 | 45 | 46 | 47 | 48 |
|---|---|---|---|---|---|---|---|---|---|---|---|---|---|---|---|---|---|---|---|---|---|---|---|
| *$w^{1118}$* ♀ x *DJ694;UAS-EDTP$^{A}$* ♂ | 3 | 04-11-30 | 1 | 11.3 | 7.3 | 7.3 | 7.7 | 4.7 | 3.3 | 2.0 | 2.0 | 3.3 | 3.3 | 4.7 | 4.5 | 4.5 | 2.0 | 2.0 | 0.5 | 0.0 | 0.0 | 0.0 | 0.0 |
| | | | 2 | 3.7 | 7.0 | 5.0 | 3.0 | 5.3 | 4.0 | 5.7 | 5.7 | 5.3 | 5.3 | 5.0 | 6.0 | 10.0 | 4.5 | 8.0 | 7.5 | 16.0 | 12.0 | 8.0 | 2.0 |
| | | | 3 | 2.0 | 2.4 | 2.8 | 0.8 | 1.4 | 0.2 | 1.6 | 0.2 | 0.6 | 0.6 | 0.6 | 0.4 | 0.0 | 0.0 | 0.0 | 0.0 | 0.0 | 0.0 | 0.0 | 0.0 |
| | | | 4 | 11.5 | 10.0 | 10.5 | 0.0 | 0.0 | 4.0 | 13.0 | 10.0 | 12.0 | 8.0 | 8.0 | 0.0 | 1.0 | 0.0 | 0.0 | 2.0 | 0.0 | 0.0 | 0.0 | 0.0 |
| *DJ694;UAS-EDTP$^{A}$* ♀ x *$w^{1118}$* ♂ | 3 | 04-11-30 | 1 | 3.4 | 9.2 | 5.6 | 4.6 | 5.0 | 1.6 | 5.8 | 3.0 | 4.4 | 6.8 | 4.4 | 1.8 | 1.4 | 6.6 | 8.6 | 2.4 | 2.0 | 3.4 | 2.6 | 2.2 |
| | | | 2 | 8.4 | 3.2 | 5.4 | 4.6 | 3.2 | 2.6 | 2.6 | 2.8 | 5.6 | 5.2 | 5.8 | 5.0 | 6.3 | 4.3 | 6.7 | 1.7 | 2.0 | 0.7 | 0.0 | 0.0 |
| | | | 3 | 3.0 | 3.0 | 6.0 | 3.7 | 2.5 | 1.5 | 4.5 | 1.5 | 0.5 | 0.5 | 0.0 | 0.0 | 0.0 | 0.0 | 0.0 | 0.0 | 0.0 | 0.0 | 0.0 | 0.0 |
| *$w^{1118}$* ♀ x *DJ694* ♂ | 3 | 04-11-30 | 1 | 2.4 | 4.5 | 1.0 | 1.5 | 0.0 | 0.3 | 0.0 | 0.3 | 0.0 | 0.5 | 0.8 | 0.0 | 0.0 | 0.0 | 0.0 | 0.0 | 0.0 | 0.0 | 0.0 | 0.0 |
| | | | 2 | 4.2 | 2.6 | 2.7 | 2.0 | 2.3 | 1.7 | 1.7 | 0.0 | 0.0 | 0.0 | 0.0 | 0.0 | 0.0 | 0.0 | 0.0 | 0.0 | 0.0 | 0.0 | 0.0 | 0.0 |
| | | | 3 | 14.3 | 6.8 | 6.0 | 4.3 | 3.8 | 2.3 | 4.8 | 6.5 | 4.5 | 5.8 | 4.0 | 5.8 | 3.8 | 4.0 | 5.3 | 3.8 | 2.5 | 6.3 | 2.5 | 0.3 |
| | | | 4 | 5.8 | 8.0 | 5.4 | 5.6 | 3.2 | 5.4 | 7.0 | 5.0 | 4.0 | 7.6 | 4.4 | 4.6 | 3.2 | 0.8 | 1.2 | 3.6 | 2.6 | 2.0 | 4.0 | 1.0 |
| *$w^{1118}$* ♀ x *DJ694;UAS-EDTP$^{B}$* ♂ | 3 | 04-11-30 | 1 | 20.5 | 12.5 | 11.0 | 12.0 | 8.0 | 11.0 | 8.0 | 12.0 | 15.5 | 16.5 | 8.5 | 25.5 | 11.5 | 8.0 | 18.5 | 10.5 | 10.5 | 5.0 | 12.5 | 4.0 |
| | | | 2 | 5.4 | 5.0 | 7.4 | 4.6 | 3.4 | 2.6 | 4.4 | 3.6 | 5.2 | 7.8 | 4.0 | 4.0 | 6.5 | 1.3 | 5.5 | 3.3 | 3.0 | 3.3 | 0.7 | 0.3 |
| | | | 3 | 9.0 | 8.0 | 7.0 | 4.5 | 8.0 | 8.0 | 11.0 | 5.5 | 9.5 | 10.0 | 6.0 | 7.5 | 19.0 | 9.0 | 15.0 | 13.0 | 5.0 | 0.0 | 3.0 | 2.0 |
| | | | 4 | 5.5 | 7.3 | 4.5 | 1.8 | 4.3 | 4.0 | 3.0 | 3.8 | 5.5 | 3.8 | 3.0 | 2.5 | 3.8 | 2.0 | 2.0 | 1.8 | 2.3 | 0.0 | 2.8 | 1.0 |
| *DJ694;UAS-EDTP$^{B}$* ♀ x *$w^{1118}$* ♂ | 3 | 04-11-30 | 1 | 3.6 | 3.2 | 8.8 | 2.0 | 5.6 | 3.5 | 4.8 | 5.3 | 5.3 | 3.8 | 3.0 | 5.7 | 7.0 | 8.0 | 9.0 | 3.0 | 4.0 | 2.0 | 1.0 | 6.5 |
| | | | 2 | 9.0 | 7.5 | 11.5 | 7.5 | 2.5 | 10.0 | 14.5 | 10.0 | 12.5 | 8.5 | 9.0 | 7.0 | 8.5 | 12.0 | 13.5 | 4.5 | 1.5 | 8.0 | 1.5 | 6.5 |
| | | | 3 | 7.8 | 4.3 | 7.0 | 3.8 | 2.8 | 6.0 | 3.7 | 5.3 | 1.3 | 5.7 | 4.3 | 5.7 | 12.7 | 11.5 | 9.0 | 7.0 | 2.5 | 4.0 | 1.0 | 4.5 |
| *DJ694* ♀ x *$w^{1118}$* ♂ | 3 | 04-11-30 | 1 | 1.8 | 0.0 | 2.3 | 1.3 | 0.5 | 0.3 | 0.5 | 0.0 | 0.0 | 0.0 | 0.0 | 0.0 | 0.0 | 0.0 | x | x | x | x | x | x |
| | | | 2 | 3.2 | 1.4 | 4.4 | 1.4 | 2.0 | 1.2 | 0.8 | 0.4 | 1.6 | 0.6 | 1.0 | 1.4 | 0.6 | 0.2 | 0.2 | 0.2 | 0.2 | 0.2 | 0.8 | 0.4 |
| | | | 3 | 1.8 | 2.0 | 1.8 | 2.0 | 2.8 | 3.5 | 1.5 | 2.3 | 2.0 | 1.7 | 4.0 | 4.7 | 1.7 | 1.0 | 1.7 | 1.7 | 2.0 | 0.7 | 1.0 | 2.3 |
| | | | 4 | 3.7 | 1.0 | 0.3 | 4.7 | 1.3 | 2.7 | 2.3 | 0.3 | 1.0 | 1.7 | 0.7 | 0.0 | 0.0 | 0.0 | 1.5 | 1.5 | 0.5 | 0.0 | 1.0 | 0.5 |

| | | Start Date | | Days | | | | | | | | | | | | | | | | | | | |
|---|---|---|---|---|---|---|---|---|---|---|---|---|---|---|---|---|---|---|---|---|---|---|---|
| **Cross** | **Rep** | **(YYMMDD)** | **Vial** | **1** | **2** | **3** | **4** | **5** | **6** | **7** | **8** | **9** | **10** | **11** | **12** | **13** | **14** | **15** | **16** | **17** | **18** | **19** | **20** |
| *DJ694*$^{H}$ ♀ x *w*$^{1118}$ ♂ | 4 | 24-07-09 | 1 | 15.0 | 1.0 | 13.0 | 4.3 | 28.0 | 4.7 | 22.3 | 28.0 | 1.0 | 17.0 | 0.5 | 30.5 | 18.0 | 11.0 | 0.5 | 11.5 | 15.0 | 9.0 | 18.5 | 16.5 |
| | | | 2 | 20.7 | 3.7 | 10.3 | 12.7 | 1.7 | 11.0 | 17.3 | 9.7 | 9.7 | 5.3 | 2.3 | 14.3 | 14.3 | 4.7 | 3.3 | 13.3 | 14.0 | 3.7 | 13.0 | 3.3 |
| | | | 3 | 7.0 | 0.3 | 5.0 | 19.0 | 9.0 | 8.0 | 35.0 | 13.0 | 0.7 | 19.7 | 7.3 | 15.0 | 12.7 | 1.0 | 5.7 | 14.3 | 16.7 | 13.3 | 7.0 | 6.7 |
| | | | 4 | 11.0 | 0.7 | 32.0 | 1.0 | 2.7 | 20.0 | 13.3 | 9.3 | 11.7 | 4.3 | 0.7 | 7.3 | 22.3 | 6.7 | 6.3 | 5.0 | 2.0 | 22.0 | 4.3 | 14.7 |
| | | | 5 | 4.0 | 3.3 | 8.7 | 5.0 | 12.7 | 16.0 | 25.0 | 15.0 | 14.7 | 6.3 | 0.7 | 31.0 | 7.3 | 18.3 | 1.0 | 13.7 | 6.7 | 15.0 | 3.3 | 6.3 |
| | | | 6 | 11.7 | 7.7 | 26.3 | 13.0 | 6.3 | 8.3 | 29.0 | 1.0 | 17.0 | 1.0 | 12.0 | 8.0 | 23.0 | 1.0 | 0.5 | 16.5 | 13.5 | 2.0 | 2.5 | 10.0 |
| | | | 7 | 19.0 | 4.3 | 34.3 | 4.3 | 2.0 | 19.0 | 21.7 | 0.7 | 29.0 | 5.3 | 1.3 | 2.0 | 26.0 | 1.0 | 1.0 | 13.3 | 10.3 | 5.3 | 1.7 | 24.3 |
| | | | 8 | 9.7 | 1.0 | 1.3 | 8.7 | 5.7 | 20.0 | 22.3 | 1.0 | 24.0 | 7.0 | 0.7 | 16.7 | 7.3 | 13.3 | 2.0 | 6.0 | 11.0 | 3.3 | 13.0 | 9.0 |
| | 5 | 24-07-10 | 1 | 1.0 | 7.0 | 6.7 | 6.7 | 4.3 | 26.7 | 5.3 | 11.3 | 23.0 | 2.7 | 28.0 | 4.3 | 4.7 | 9.3 | 4.0 | 4.7 | 19.7 | 8.3 | 8.0 | 2.3 |
| | | | 2 | 0.3 | 5.3 | 11.3 | 0.3 | 11.3 | 35.0 | 13.7 | 10.3 | 27.7 | 0.7 | 1.0 | 10.7 | 9.7 | 10.7 | 11.0 | 11.0 | 12.7 | 16.7 | 6.3 | 7.0 |
| | | | 3 | 0.0 | 0.7 | 10.0 | 1.3 | 9.3 | 29.7 | 1.0 | 13.3 | 32.3 | 2.3 | 1.0 | 29.7 | 1.0 | 8.3 | 15.3 | 16.0 | 18.7 | 1.7 | 1.7 | 3.3 |
| | | | 4 | 0.0 | 2.7 | 7.3 | 0.7 | 25.3 | 1.0 | 9.3 | 2.3 | 24.7 | 8.3 | 0.7 | 7.3 | 12.0 | 10.3 | 13.0 | 12.7 | 10.3 | 4.7 | 5.3 | 2.0 |
| | | | 5 | 1.0 | 13.7 | 9.7 | 8.0 | 30.3 | 12.3 | 4.0 | 7.3 | 25.0 | 6.3 | 22.3 | 8.3 | 1.0 | 22.3 | 13.5 | 13.5 | 11.5 | 1.0 | 1.0 | 12.0 |
| | | | 6 | 0.0 | 4.7 | 9.7 | 0.3 | 9.0 | 6.3 | 0.3 | 7.3 | 3.0 | 3.0 | 10.7 | 1.0 | 0.3 | 17.0 | 1.3 | 1.0 | 10.7 | 0.3 | 7.7 | 1.3 |
| | | | 7 | 0.0 | 3.3 | 16.7 | 1.3 | 30.0 | 1.7 | 18.0 | 1.0 | 25.3 | 0.7 | 1.0 | 34.5 | 1.0 | 1.0 | 20.0 | 15.0 | 0.5 | 18.0 | 12.0 | 1.0 |
| | | | 8 | 0.0 | 0.0 | 12.3 | 8.7 | 18.3 | 14.3 | 3.0 | 13.0 | 12.0 | 7.7 | 11.7 | 8.7 | 14.3 | 11.0 | 10.0 | 9.0 | 1.0 | 3.0 | 23.5 | 0.5 |

| Cross | Rep | Start Date (YYMMDD) | Vial | Days 21 | 22 | 23 | 24 | 25 | 26 | 27 | 28 | 29 | 30 | 31 | 32 | 33 | 34 | 35 | 36 | 37 | 38 | 39 | 40 |
|---|---|---|---|---|---|---|---|---|---|---|---|---|---|---|---|---|---|---|---|---|---|---|---|
| *DJ694*$^{H}$ ♀ x *w*$^{1118}$ ♂ | 4 | 24-07-09 | 1 | 4.5 | 1.0 | 14.0 | 7.5 | 4.0 | 4.5 | 15.0 | 3.5 | 0.5 | 1.0 | 9.5 | 2.5 | 13.0 | 7.0 | 5.5 | 0.5 | 1.0 | 1.0 | 13.0 | 2.0 |
| | | | 2 | 7.3 | 5.0 | 2.7 | 2.7 | 3.0 | 3.0 | 4.0 | 1.3 | 0.3 | 1.0 | 7.3 | 0.3 | 0.7 | 1.0 | 2.3 | 1.7 | 6.0 | 2.0 | 0.7 | 2.7 |
| | | | 3 | 4.7 | 2.0 | 11.7 | 0.7 | 4.7 | 3.7 | 4.3 | 2.7 | 1.7 | 0.7 | 0.7 | 0.3 | 0.0 | 1.3 | 1.0 | 1.3 | 0.7 | 1.3 | 1.3 | 0.0 |
| | | | 4 | 10.3 | 1.3 | 2.0 | 13.0 | 3.7 | 3.3 | 4.3 | 5.3 | 3.3 | 1.7 | 8.7 | 5.0 | 5.3 | 1.7 | 3.0 | 6.3 | 3.3 | 6.7 | 0.7 | 1.3 |
| | | | 5 | 5.0 | 1.3 | 8.3 | 2.0 | 1.7 | 2.0 | 5.0 | 6.7 | 2.0 | 3.7 | 6.7 | 4.7 | 1.7 | 2.7 | 3.0 | 7.7 | 5.5 | 7.5 | 0.5 | 5.5 |
| | | | 6 | 11.5 | 2.5 | 2.0 | 3.0 | 3.0 | 3.5 | 1.0 | 0.0 | 0.0 | 0.0 | 0.5 | 2.0 | 0.0 | 0.0 | 0.0 | 0.5 | 0.0 | 0.0 | 0.0 | 0.0 |
| | | | 7 | 8.0 | 9.3 | 1.7 | 11.3 | 5.0 | 3.3 | 3.7 | 3.3 | 0.7 | 2.0 | 7.0 | 1.0 | 6.3 | 3.3 | 6.0 | 3.0 | 5.0 | 6.0 | 2.0 | 0.0 |
| | | | 8 | 3.3 | 2.0 | 1.0 | 20.3 | 1.7 | 1.7 | 9.7 | 7.3 | 0.7 | 0.0 | 0.7 | 8.3 | 4.0 | 1.0 | 3.0 | 1.0 | 1.3 | 4.7 | 3.0 | 3.3 |
| | 5 | 24-07-10 | 1 | 2.0 | 5.0 | 7.3 | 6.0 | 6.3 | 10.0 | 10.0 | 0.7 | 1.3 | 6.7 | 1.0 | 3.3 | 6.7 | 3.0 | 2.7 | 2.7 | 2.7 | 3.0 | 4.0 | 2.7 |
| | | | 2 | 5.3 | 7.7 | 9.3 | 5.7 | 6.0 | 4.0 | 8.0 | 1.0 | 0.3 | 1.0 | 0.3 | 0.7 | 0.3 | 2.0 | 0.3 | 1.7 | 0.3 | 0.3 | 0.0 | 0.3 |
| | | | 3 | 18.0 | 2.7 | 10.3 | 7.0 | 6.7 | 11.0 | 14.7 | 3.3 | 2.7 | 4.0 | 14.0 | 2.7 | 1.7 | 2.3 | 4.0 | 11.0 | 2.7 | 4.0 | 3.7 | 3.3 |
| | | | 4 | 7.3 | 4.0 | 7.7 | 5.0 | 5.0 | 6.0 | 3.3 | 1.3 | 2.7 | 1.7 | 1.3 | 3.0 | 2.7 | 0.3 | 4.3 | 2.3 | 1.3 | 2.3 | 2.0 | 0.7 |
| | | | 5 | 3.0 | 3.0 | 16.5 | 9.0 | 9.5 | 10.0 | 2.0 | 0.5 | 1.0 | 6.0 | 4.5 | 4.5 | 4.0 | 0.5 | 2.5 | 2.5 | 1.5 | 1.5 | 0.5 | 1.0 |
| | | | 6 | 1.7 | 10.3 | 3.3 | 0.7 | 1.0 | 10.7 | 2.0 | 3.0 | 3.7 | 4.0 | 2.7 | 0.7 | 0.0 | 0.3 | 0.7 | 0.0 | 0.3 | 0.0 | 0.0 | 0.0 |
| | | | 7 | 17.0 | 3.5 | 1.5 | 4.0 | 5.0 | 2.5 | 1.0 | 0.0 | 4.0 | 2.0 | 0.0 | 0.0 | 4.0 | 0.0 | 0.0 | 0.0 | 0.0 | 1.0 | 1.0 | 0.0 |
| | | | 8 | 17.5 | 0.5 | 5.0 | 7.0 | 6.5 | 11.0 | 2.5 | 3.5 | 1.5 | 3.5 | 9.0 | 2.0 | 5.0 | 0.5 | 0.5 | 7.5 | 5.5 | 3.5 | 2.0 | 6.5 |

| Cross | Rep | Start Date (YYMMDD) | Vial | Days | | | | | | | | | | | | | | | | | | | |
|---|---|---|---|---|---|---|---|---|---|---|---|---|---|---|---|---|---|---|---|---|---|---|---|
| | | | | 1 | 2 | 3 | 4 | 5 | 6 | 7 | 8 | 9 | 10 | 11 | 12 | 13 | 14 | 15 | 16 | 17 | 18 | 19 | 20 |
| *$DJ694^{H}$/+* ♀ x *$w^{1118}$* ♂ | 4 | 24-07-09 | 1 | 8.0 | 5.3 | 1.0 | 20.7 | 0.7 | 7.0 | 18.3 | 14.3 | 10.7 | 8.0 | 11.0 | 20.0 | 9.0 | 3.3 | 12.7 | 17.7 | 4.7 | 6.0 | 3.3 | 16.7 |
| | | | 2 | 9.0 | 4.0 | 27.5 | 1.0 | 1.0 | 24.5 | 16.5 | 12.5 | 13.5 | 1.0 | 1.5 | 41.0 | 18.0 | 7.5 | 6.0 | 20.0 | 14.0 | 4.0 | 13.0 | 13.0 |
| | | | 3 | 14.7 | 13.7 | 15.0 | 10.3 | 4.3 | 13.0 | 10.7 | 23.7 | 10.0 | 15.7 | 16.7 | 16.3 | 16.3 | 6.0 | 9.7 | 28.3 | 9.3 | 9.0 | 13.3 | 8.3 |
| | | | 4 | 14.7 | 1.0 | 2.0 | 16.3 | 1.0 | 28.7 | 19.0 | 13.0 | 9.3 | 21.7 | 4.7 | 28.7 | 6.3 | 18.7 | 2.3 | 29.3 | 9.3 | 1.0 | 1.3 | 19.3 |
| | | | 5 | 16.3 | 18.7 | 6.7 | 11.0 | 6.3 | 18.0 | 9.3 | 15.0 | 18.7 | 15.7 | 3.0 | 20.0 | 22.7 | 15.0 | 6.7 | 31.3 | 12.7 | 10.7 | 13.0 | 6.3 |
| | | | 6 | 16.7 | 2.0 | 23.0 | 13.0 | 1.0 | 25.3 | 10.7 | 2.7 | 18.3 | 16.0 | 4.3 | 20.3 | 15.3 | 13.7 | 4.0 | 18.3 | 14.3 | 10.7 | 12.7 | 15.0 |
| | | | 7 | 7.0 | 21.3 | 4.7 | 16.7 | 2.3 | 16.3 | 21.3 | 22.0 | 12.3 | 24.0 | 0.0 | 19.7 | 13.0 | 3.3 | 5.0 | 26.0 | 21.3 | 14.3 | 8.3 | 15.7 |
| | | | 8 | 16.3 | 11.7 | 9.3 | 20.3 | 14.3 | 9.7 | 17.7 | 16.0 | 15.3 | 12.7 | 5.3 | 21.3 | 12.0 | 13.7 | 1.0 | 27.0 | 3.7 | 6.3 | 1.0 | 7.3 |
| | 5 | 24-07-10 | 1 | 10.0 | 17.3 | 13.7 | 5.7 | 15.3 | 22.0 | 30.3 | 5.0 | 27.3 | 27.0 | 19.7 | 14.7 | 18.7 | 13.0 | 10.0 | 15.0 | 11.5 | 5.0 | 22.0 | 2.0 |
| | | | 2 | 10.0 | 11.7 | 10.3 | 1.3 | 21.3 | 21.7 | 12.7 | 25.7 | 20.3 | 20.0 | 9.7 | 35.7 | 5.7 | 16.7 | 14.0 | 14.0 | 3.7 | 7.0 | 16.0 | 5.5 |
| | | | 3 | 7.7 | 8.0 | 14.7 | 1.0 | 10.0 | 32.0 | 25.3 | 20.0 | 15.3 | 11.7 | 14.3 | 19.0 | 9.0 | 7.7 | 13.3 | 19.5 | 7.0 | 1.0 | 14.0 | 1.0 |
| | | | 4 | 15.3 | 13.0 | 13.3 | 8.7 | 23.0 | 31.0 | 8.0 | 21.7 | 29.7 | 21.0 | 17.0 | 24.7 | 21.0 | 18.3 | 15.0 | 15.0 | 3.3 | 15.7 | 11.3 | 1.3 |
| | | | 5 | 10.7 | 8.3 | 6.0 | 17.0 | 14.7 | 17.3 | 12.0 | 21.3 | 17.0 | 17.3 | 23.0 | 14.7 | 19.7 | 17.0 | 14.5 | 14.5 | 9.5 | 1.0 | 12.5 | 1.0 |
| | | | 6 | 0.0 | 13.7 | 4.0 | 11.0 | 7.0 | 20.7 | 7.0 | 29.3 | 20.7 | 12.0 | 14.7 | 7.0 | 32.0 | 6.7 | 17.3 | 16.7 | 1.3 | 12.3 | 1.0 | 0.7 |
| | | | 7 | 5.3 | 5.3 | 25.3 | 14.0 | 9.3 | 21.3 | 15.3 | 24.3 | 20.0 | 1.0 | 1.3 | 45.0 | 1.0 | 36.0 | 5.0 | 8.3 | 7.0 | 21.3 | 9.3 | 1.0 |
| | | | 8 | 4.0 | 13.0 | 6.0 | 24.3 | 14.3 | 14.3 | 5.7 | 22.7 | 15.3 | 10.0 | 15.7 | 25.0 | 1.0 | 31.3 | 13.7 | 6.0 | 1.0 | 12.3 | 5.0 | 1.0 |
| *$DJ694^{L}$* ♀ x *$w^{1118}$* ♂ | 4 | 24-07-09 | 1 | 11.0 | 4.0 | 10.0 | 24.0 | 7.7 | 5.0 | 19.3 | 9.3 | 2.7 | 8.3 | 10.0 | 9.0 | 16.3 | 3.7 | 4.3 | 21.5 | 21.5 | 7.5 | 13.0 | 12.5 |
| | | | 2 | 13.0 | 0.5 | 1.5 | 7.5 | 11.5 | 27.0 | 44.0 | 4.0 | 10.5 | 7.5 | 13.0 | 11.0 | 33.0 | 1.0 | 1.0 | 22.5 | 22.0 | 5.0 | 20.0 | 7.0 |
| | | | 3 | 14.3 | 18.3 | 3.7 | 17.3 | 14.0 | 4.7 | 24.0 | 13.7 | 1.0 | 12.3 | 3.7 | 7.3 | 14.0 | 4.7 | 8.0 | 7.0 | 5.7 | 5.0 | 9.7 | 14.3 |
| | | | 4 | 15.7 | 3.0 | 11.7 | 33.0 | 1.3 | 18.7 | 24.7 | 1.0 | 7.0 | 8.3 | 1.7 | 18.3 | 18.3 | 2.0 | 4.7 | 29.0 | 17.0 | 6.0 | 4.0 | 17.0 |
| | | | 5 | 15.3 | 1.3 | 19.0 | 14.3 | 16.0 | 6.0 | 21.0 | 6.7 | 3.0 | 7.0 | 9.3 | 20.0 | 8.0 | 4.7 | 4.3 | 17.3 | 3.0 | 9.7 | 12.3 | 13.0 |
| | | | 6 | 7.0 | 0.7 | 11.0 | 1.7 | 0.7 | 13.0 | 16.3 | 1.0 | 4.0 | 9.3 | 0.0 | 1.7 | 4.0 | 2.3 | 3.0 | 2.0 | 2.0 | 0.7 | 0.3 | 0.7 |
| | | | 7 | 21.7 | 4.0 | 1.3 | 19.7 | 1.7 | 13.7 | 23.0 | 11.0 | 1.0 | 9.3 | 1.0 | 9.3 | 20.7 | 2.0 | 1.3 | 33.3 | 9.3 | 0.7 | 15.0 | 2.3 |
| | | | 8 | 12.7 | 5.3 | 10.7 | 8.7 | 5.0 | 17.0 | 12.7 | 9.7 | 9.3 | 8.3 | 3.0 | 10.0 | 11.3 | 3.7 | 4.3 | 14.0 | 3.7 | 10.0 | 8.3 | 3.0 |
| *$DJ694^{L}$* ♀ x *$w^{1118}$* ♂ | 5 | 24-07-10 | 1 | 0.0 | 10.3 | 10.0 | 2.3 | 18.3 | 12.7 | 12.3 | 23.0 | 4.0 | 10.3 | 7.7 | 21.0 | 1.0 | 3.3 | 18.7 | 16.7 | 3.7 | 5.7 | 23.7 | 3.0 |
| | | | 2 | 0.0 | 1.0 | 25.0 | 0.7 | 2.0 | 28.0 | 1.7 | 0.7 | 13.3 | 1.7 | 13.3 | 20.0 | 1.0 | 4.3 | 13.3 | 16.0 | 2.7 | 3.7 | 14.3 | 5.7 |
| | | | 3 | 0.0 | 2.7 | 13.0 | 3.0 | 12.0 | 17.0 | 8.3 | 4.3 | 23.3 | 4.7 | 10.0 | 15.7 | 0.3 | 9.7 | 11.0 | 8.0 | 6.7 | 18.3 | 0.7 | 8.0 |
| | | | 4 | 0.0 | 0.7 | 0.7 | 2.0 | 1.0 | 31.7 | 2.0 | 0.7 | 43.3 | 0.3 | 1.0 | 5.0 | 1.0 | 25.7 | 23.3 | 15.0 | 3.3 | 1.3 | 9.7 | 6.7 |
| | | | 5 | 0.0 | 0.3 | 1.3 | 2.0 | 13.3 | 4.0 | 12.3 | 1.0 | 2.7 | 7.7 | 12.3 | 12.7 | 6.7 | 3.3 | 17.7 | 19.3 | 12.0 | 4.3 | 12.0 | 3.0 |
| | | | 6 | 0.0 | 0.7 | 6.0 | 1.0 | 7.7 | 22.0 | 2.7 | 2.0 | 0.7 | 4.0 | 14.0 | 31.3 | 2.7 | 10.7 | 22.3 | 22.3 | 10.7 | 3.0 | 17.3 | 3.3 |
| | | | 7 | 0.0 | 0.0 | 4.0 | 3.3 | 7.3 | 29.7 | 4.3 | 1.7 | 6.0 | 5.7 | 8.3 | 36.3 | 1.0 | 6.3 | 18.3 | 18.0 | 5.3 | 0.7 | 17.0 | 3.7 |
| | | | 8 | 0.0 | 0.0 | 10.3 | 6.0 | 7.0 | 34.7 | 1.3 | 0.7 | 3.3 | 1.3 | 6.0 | 23.0 | 1.3 | 20.7 | 24.0 | 29.3 | 1.0 | 6.7 | 1.0 | 4.0 |
| | 6 | 24-09-28 | 1 | 11.0 | 9.3 | 7.3 | 15.7 | 26.0 | 3.0 | 14.3 | 19.3 | 8.3 | 10.3 | 4.0 | 6.7 | 3.7 | 11.3 | 10.3 | 3.0 | 9.7 | 9.7 | 6.0 | 17.3 |
| | | | 2 | 13.7 | 10.7 | 1.3 | 24.3 | 11.0 | 7.0 | 12.3 | 12.3 | 5.0 | 15.3 | 2.0 | 9.3 | 7.3 | 11.3 | 6.0 | 7.0 | 6.7 | 6.0 | 3.0 | 9.7 |
| | | | 3 | 9.7 | 5.3 | 21.0 | 11.7 | 14.0 | 6.0 | 9.3 | 12.7 | 12.0 | 3.3 | 6.0 | 11.0 | 9.7 | 7.3 | 4.0 | 3.0 | 6.3 | 8.0 | 3.7 | 4.0 |
| | | | 4 | 15.0 | 6.3 | 23.0 | 5.7 | 19.3 | 14.7 | 6.3 | 15.7 | 4.3 | 17.3 | 6.0 | 3.7 | 6.7 | 11.7 | 3.7 | 3.3 | 1.0 | 16.3 | 4.0 | 8.3 |
| | | | 5 | 6.3 | 4.7 | 14.3 | 9.7 | 9.7 | 8.0 | 8.7 | 15.7 | 13.3 | 9.0 | 4.3 | 6.3 | 7.0 | 13.3 | 3.0 | 12.7 | 1.0 | 8.0 | 9.0 | 11.3 |
| | | | 6 | 8.7 | 8.3 | 11.3 | 9.3 | 17.7 | 6.0 | 13.3 | 13.7 | 9.7 | 12.3 | 3.3 | 3.3 | 7.3 | 5.0 | 5.0 | 1.7 | 11.3 | 7.0 | 1.7 | 14.7 |
| | | | 7 | 11.3 | 9.3 | 10.0 | 13.0 | 17.3 | 9.3 | 5.0 | 9.0 | 11.0 | 8.3 | 8.7 | 4.3 | 7.7 | 12.0 | 7.3 | 6.3 | 3.7 | 9.0 | 11.7 | 4.3 |
| | | | 8 | 17.0 | 11.0 | 29.0 | 1.3 | 26.0 | 2.7 | 13.7 | 20.3 | 1.0 | 18.3 | 16.7 | 13.0 | 1.3 | 9.7 | 6.3 | 11.0 | 8.7 | 5.0 | 4.7 | 16.3 |

| Cross | Rep | Start Date (YYMMDD) | Vial | Days 21 | 22 | 23 | 24 | 25 | 26 | 27 | 28 | 29 | 30 | 31 | 32 | 33 | 34 | 35 | 36 | 37 | 38 | 39 | 40 |
|---|---|---|---|---|---|---|---|---|---|---|---|---|---|---|---|---|---|---|---|---|---|---|---|
| *DJ694^H/+* ♀ x *w^1118* ♂ | 4 | 24-07-09 | 1 | 9.7 | 0.3 | 1.0 | 16.0 | 8.0 | 6.7 | 10.7 | 4.3 | 6.7 | 4.3 | 7.7 | 4.3 | 3.3 | 1.0 | 11.3 | 5.0 | 7.0 | 1.3 | 6.7 | 8.3 |
| | | | 2 | 11.0 | 1.0 | 17.0 | 6.5 | 12.5 | 13.5 | 7.0 | 8.5 | 7.0 | 1.0 | 15.0 | 15.0 | 2.5 | 5.0 | 14.5 | 8.0 | 5.0 | 10.0 | 12.5 | 6.0 |
| | | | 3 | 10.3 | 4.3 | 11.3 | 6.3 | 11.0 | 10.7 | 6.3 | 7.0 | 4.3 | 8.7 | 7.0 | 6.3 | 4.0 | 6.3 | 3.3 | 6.0 | 6.5 | 5.5 | 5.5 | 5.0 |
| | | | 4 | 10.7 | 3.7 | 2.0 | 9.0 | 13.3 | 12.3 | 5.0 | 8.0 | 4.3 | 1.0 | 1.0 | 11.0 | 7.7 | 3.7 | 6.7 | 1.3 | 3.7 | 14.0 | 6.7 | 2.3 |
| | | | 5 | 7.0 | 2.5 | 7.5 | 14.0 | 9.0 | 8.0 | 11.0 | 1.0 | 11.0 | 7.0 | 8.0 | 15.0 | 1.0 | 1.0 | 1.0 | 0.0 | 1.0 | 1.0 | 0.0 | 0.0 |
| | | | 6 | 8.0 | 2.0 | 18.7 | 4.7 | 12.3 | 5.7 | 9.0 | 5.7 | 10.3 | 4.7 | 9.7 | 4.3 | 13.0 | 3.7 | 5.7 | 1.7 | 6.7 | 13.7 | 1.3 | 8.0 |
| | | | 7 | 8.3 | 2.0 | 14.7 | 5.3 | 12.0 | 11.5 | 8.5 | 11.5 | 15.5 | 6.5 | 1.0 | 18.0 | 5.0 | 1.0 | 11.5 | 17.0 | 0.5 | 26.0 | 2.0 | 9.5 |
| | | | 8 | 15.0 | 11.7 | 4.7 | 4.3 | 14.3 | 13.3 | 7.3 | 8.3 | 4.0 | 3.3 | 16.7 | 1.7 | 17.0 | 1.0 | 15.7 | 1.0 | 8.3 | 8.3 | 10.0 | 10.3 |
| | 5 | 24-07-10 | 1 | 1.0 | 18.0 | 4.5 | 14.0 | 13.5 | 15.0 | 3.5 | 1.0 | 1.0 | 1.0 | 4.0 | 26.0 | 4.5 | 19.0 | 4.0 | 17.5 | 5.5 | 9.5 | 12.5 | 4.5 |
| | | | 2 | 1.0 | 9.0 | 1.0 | 16.0 | 17.0 | 11.0 | 1.0 | 0.0 | 2.0 | 1.0 | 2.0 | 1.0 | 2.0 | 1.0 | 2.0 | 3.0 | 0.0 | x | x | x |
| | | | 3 | 1.0 | 37.0 | 1.0 | 18.5 | 19.0 | 7.5 | 8.5 | 1.0 | 18.0 | 2.0 | 1.5 | 11.5 | 1.0 | 22.5 | 4.0 | 10.0 | 10.0 | 18.0 | 2.0 | 8.0 |
| | | | 4 | 19.3 | 5.0 | 14.3 | 14.0 | 13.0 | 9.7 | 7.0 | 3.0 | 21.0 | 1.0 | x | x | x | x | x | x | x | x | x | x |
| | | | 5 | 24.5 | 5.5 | 17.5 | 12.5 | 10.0 | 12.5 | 10.0 | 9.5 | 1.0 | 1.0 | 1.0 | 33.0 | 14.0 | 7.0 | 21.0 | 27.0 | 0.0 | 20.0 | 11.0 | x |
| | | | 6 | 6.3 | 9.3 | 1.7 | 20.7 | 20.7 | 8.0 | 14.0 | 9.0 | 1.3 | 1.0 | 1.0 | 15.7 | 1.3 | 8.0 | 1.0 | 17.3 | 11.7 | 12.0 | 10.7 | 10.7 |
| | | | 7 | 22.0 | 11.0 | 14.0 | 22.0 | 10.0 | 19.0 | 2.0 | 9.0 | 1.0 | 1.0 | 1.0 | 36.0 | 2.0 | 19.0 | 0.0 | x | x | x | x | x |
| | | | 8 | 1.0 | 22.3 | 10.0 | 12.0 | 12.3 | 8.3 | 8.0 | 13.5 | 1.0 | 4.0 | 4.0 | 13.0 | 9.0 | 10.0 | 9.5 | 8.5 | 7.5 | 10.5 | 11.5 | 17.5 |
| *DJ694^L* ♀ x *w^1118* ♂ | 4 | 24-07-09 | 1 | 11.0 | 1.0 | 10.5 | 6.5 | 5.0 | 5.5 | 12.5 | 10.5 | 2.0 | 10.0 | 2.5 | 10.0 | 2.0 | 1.0 | 5.5 | 2.0 | 1.5 | 6.0 | 0.0 | 1.0 |
| | | | 2 | 13.5 | 3.5 | 23.5 | 3.5 | 9.0 | 5.0 | 10.0 | 7.0 | 5.5 | 3.0 | 7.5 | 16.0 | 12.0 | 1.5 | 15.5 | 4.0 | 5.5 | 2.5 | 2.5 | 1.0 |
| | | | 3 | 8.3 | 3.0 | 8.0 | 2.7 | 4.7 | 3.3 | 3.0 | 1.7 | 3.0 | 2.3 | 3.3 | 1.7 | 4.0 | 1.0 | 3.3 | 1.0 | 0.0 | 0.7 | 0.0 | 0.0 |
| | | | 4 | 26.7 | 5.7 | 18.0 | 5.7 | 12.7 | 11.0 | 8.0 | 4.0 | 2.0 | 7.0 | 8.7 | 3.7 | 9.0 | 5.7 | 1.0 | 7.0 | 2.3 | 3.3 | 3.0 | 0.7 |
| | | | 5 | 8.3 | 3.7 | 6.7 | 5.0 | 5.3 | 1.3 | 7.7 | 6.0 | 1.3 | 0.7 | 2.0 | 2.0 | 2.0 | 0.3 | 0.7 | 2.0 | 3.0 | 4.0 | 2.3 | 1.0 |
| | | | 6 | 0.0 | 0.0 | 0.0 | 0.3 | 0.0 | 0.0 | 0.0 | 0.0 | 0.0 | 0.0 | 0.0 | 0.0 | 0.0 | 0.0 | 0.0 | 0.0 | 0.0 | 0.0 | 0.0 | 0.0 |
| | | | 7 | 10.3 | 6.7 | 5.3 | 4.3 | 6.3 | 5.0 | 4.3 | 3.0 | 1.7 | 5.0 | 5.0 | 4.0 | 1.7 | 2.0 | 2.3 | 0.7 | 0.3 | 0.0 | 0.7 | 0.0 |
| | | | 8 | 7.7 | 0.5 | 1.5 | 2.0 | 5.0 | 0.0 | 3.5 | 1.5 | 0.0 | 0.0 | 0.0 | 0.0 | 0.0 | 0.0 | 0.0 | 0.0 | 0.0 | 0.0 | 0.0 | 0.0 |
| *DJ694^L* ♀ x *w^1118* ♂ | 5 | 24-07-10 | 1 | 3.3 | 14.3 | 2.7 | 10.3 | 6.3 | 11.3 | 6.3 | 1.3 | 12.0 | 7.3 | 8.3 | 7.3 | 2.0 | 0.3 | 14.7 | 4.3 | 1.7 | 6.0 | 2.3 | 2.7 |
| | | | 2 | 1.0 | 8.7 | 6.3 | 5.3 | 4.0 | 6.7 | 2.0 | 1.7 | 1.3 | 3.0 | 1.0 | 4.0 | 0.3 | 1.0 | 2.3 | 1.3 | 0.0 | 1.3 | 1.0 | 0.7 |
| | | | 3 | 8.3 | 5.7 | 14.3 | 5.0 | 3.0 | 6.0 | 6.3 | 1.7 | 2.7 | 5.0 | 11.3 | 3.0 | 4.0 | 5.0 | 4.0 | 4.7 | 0.7 | 3.3 | 3.0 | 5.0 |
| | | | 4 | 3.3 | 26.0 | 4.7 | 3.7 | 8.3 | 11.3 | 12.0 | 2.3 | 0.7 | 0.3 | 3.0 | 6.0 | 0.7 | 2.3 | 5.0 | 5.0 | 0.7 | 1.3 | 2.0 | 1.0 |
| | | | 5 | 16.3 | 6.7 | 2.3 | 17.7 | 18.0 | 1.7 | 0.0 | 1.3 | 1.3 | 1.0 | 5.0 | 9.3 | 0.3 | 1.3 | 3.3 | 0.3 | 2.0 | 6.3 | 1.0 | 0.3 |
| | | | 6 | 5.7 | 1.3 | 10.7 | 6.7 | 8.7 | 4.3 | 6.3 | 0.7 | 4.3 | 2.3 | 3.3 | 2.7 | 11.0 | 1.3 | 6.0 | 1.3 | 1.3 | 3.3 | 1.7 | 1.0 |
| | | | 7 | 12.7 | 14.7 | 3.3 | 11.7 | 11.7 | 10.7 | 1.7 | 1.0 | 7.3 | 5.7 | 6.0 | 3.3 | 3.7 | 8.3 | 3.3 | 6.0 | 2.7 | 0.0 | 0.0 | 0.0 |
| | | | 8 | 11.7 | 1.3 | 2.7 | 18.7 | 16.7 | 9.7 | 5.7 | 2.7 | 7.3 | 5.3 | 6.7 | 3.0 | 9.7 | 6.3 | 5.0 | 7.3 | 1.0 | 3.0 | 2.0 | 0.3 |
| | 6 | 24-09-28 | 1 | 2.3 | 10.0 | 1.3 | 5.0 | 3.0 | 10.3 | 3.7 | 8.7 | 3.0 | 5.3 | 3.0 | 5.7 | 10.0 | 2.0 | 6.3 | 4.0 | 1.7 | 1.3 | 7.0 | 1.7 |
| | | | 2 | 4.7 | 9.3 | 1.0 | 5.3 | 7.3 | 1.3 | 5.0 | 1.7 | 3.7 | 7.7 | 3.7 | 2.7 | 4.7 | 8.3 | 6.0 | 3.3 | 3.0 | 3.7 | 5.0 | 6.7 |
| | | | 3 | 9.7 | 7.3 | 1.3 | 3.3 | 1.0 | 9.7 | 4.3 | 7.0 | 2.3 | 1.7 | 10.7 | 6.7 | 6.3 | 4.7 | 7.7 | 5.3 | 1.3 | 1.3 | 6.7 | 7.0 |
| | | | 4 | 9.7 | 10.0 | 0.7 | 4.0 | 3.3 | 10.0 | 2.0 | 5.0 | 2.7 | 3.7 | 7.3 | 5.7 | 5.0 | 6.7 | 6.7 | 5.0 | 6.0 | 3.7 | 4.0 | 4.3 |
| | | | 5 | 1.3 | 4.3 | 4.7 | 1.0 | 6.7 | 5.0 | 3.3 | 6.0 | 7.3 | 1.7 | 4.7 | 5.0 | 8.0 | 7.3 | 10.7 | 5.7 | 3.3 | 1.7 | 6.7 | 1.7 |
| | | | 6 | 2.0 | 3.0 | 3.3 | 2.3 | 12.0 | 3.0 | 7.0 | 6.7 | 2.7 | 7.0 | 5.7 | 3.0 | 6.3 | 6.0 | 4.7 | 6.7 | 2.7 | 6.3 | 4.3 | 3.3 |
| | | | 7 | 12.3 | 2.0 | 6.7 | 3.0 | 6.3 | 3.3 | 4.3 | 5.0 | 5.7 | 4.3 | 3.7 | 6.0 | 7.3 | 6.3 | 3.3 | 5.7 | 2.0 | 2.0 | 5.0 | 0.3 |
| | | | 8 | 3.3 | 8.3 | 5.5 | 0.7 | 6.3 | 4.3 | 1.7 | 8.3 | 5.7 | 1.0 | 2.3 | 11.3 | 3.7 | 4.7 | 5.7 | 1.0 | 3.7 | 0.3 | 0.7 | 1.3 |

| Cross | Rep | Start Date (YYMMDD) | Vial | Days 1 | 2 | 3 | 4 | 5 | 6 | 7 | 8 | 9 | 10 | 11 | 12 | 13 | 14 | 15 | 16 | 17 | 18 | 19 | 20 |
|---|---|---|---|---|---|---|---|---|---|---|---|---|---|---|---|---|---|---|---|---|---|---|---|
| *DJ694*$^{L}$ ♀ x *w*$^{1118}$ ♂ | 7 | 24-09-28 | 1 | 28.0 | 6.7 | 9.3 | 23.7 | 18.0 | 1.0 | 25.7 | 7.0 | 11.3 | 11.3 | 12.0 | 6.7 | 2.7 | 15.0 | 3.0 | 7.3 | 6.7 | 5.7 | 7.0 | 11.7 |
| | | | 2 | 20.7 | 0.7 | 13.0 | 24.3 | 3.7 | 3.3 | 22.3 | 10.0 | 15.0 | 9.3 | 9.3 | 8.0 | 8.3 | 11.3 | 3.3 | 5.0 | 3.7 | 2.7 | 2.7 | 16.7 |
| | | | 3 | 18.7 | 5.7 | 8.7 | 28.0 | 7.3 | 18.3 | 14.7 | 5.0 | 12.3 | 3.7 | 5.7 | 12.0 | 11.0 | 5.0 | 9.7 | 12.7 | 1.3 | 6.3 | 4.3 | 9.3 |
| | | | 4 | 4.7 | 15.3 | 7.3 | 17.7 | 1.0 | 1.0 | 27.3 | 6.0 | 13.7 | 10.0 | 10.7 | 6.0 | 1.0 | 13.3 | 2.7 | 11.0 | 2.0 | 8.0 | 7.7 | 9.7 |
| | | | 5 | 9.0 | 10.7 | 8.7 | 1.3 | 17.3 | 6.0 | 9.3 | 11.7 | 5.3 | 10.0 | 9.7 | 6.7 | 6.7 | 4.0 | 9.7 | 1.7 | 6.7 | 6.0 | 4.7 | 12.7 |
| | | | 6 | 6.7 | 7.7 | 1.0 | 20.3 | 1.0 | 7.7 | 23.7 | 15.3 | 3.7 | 10.7 | 7.3 | 6.7 | 7.3 | 5.0 | 4.3 | 6.3 | 10.7 | 7.0 | 8.7 | 6.3 |
| | | | 7 | 0.7 | 5.3 | 0.7 | 13.0 | 15.0 | 1.3 | 17.3 | 5.0 | 9.0 | 10.3 | 1.0 | 15.3 | 3.3 | 3.3 | 12.3 | 6.7 | 2.7 | 1.7 | 2.7 | 14.7 |
| | | | 8 | 5.3 | 11.7 | 4.7 | 26.0 | 22.7 | 4.7 | 14.0 | 9.3 | 13.7 | 6.3 | 1.0 | 8.3 | 10.3 | 3.0 | 10.3 | 5.7 | 8.7 | 2.0 | 1.7 | 13.0 |
| *DJ694*$^{L}$/+ ♀ x *w*$^{1118}$ ♂ | 4 | 24-07-09 | 1 | 17.0 | 4.7 | 17.0 | 20.7 | 10.7 | 18.3 | 11.3 | 26.0 | 15.7 | 13.0 | 3.0 | 21.3 | 13.3 | 8.3 | 8.0 | 27.7 | 5.7 | 3.3 | 7.7 | 12.0 |
| | | | 2 | 12.0 | 6.0 | 1.0 | 24.3 | 12.7 | 4.0 | 19.3 | 7.3 | 11.3 | 14.3 | 12.7 | 17.7 | 9.0 | 13.3 | 1.0 | 18.3 | 10.7 | 10.0 | 4.7 | 19.3 |
| | | | 3 | 22.0 | 1.0 | 0.7 | 39.3 | 1.3 | 15.0 | 30.0 | 19.3 | 18.7 | 19.7 | 16.0 | 1.3 | 30.3 | 7.3 | 1.0 | 25.7 | 19.7 | 18.0 | 6.3 | 12.0 |
| | | | 4 | 19.0 | 18.7 | 1.0 | 30.0 | 1.3 | 10.0 | 28.7 | 13.0 | 25.3 | 21.3 | 13.3 | 13.0 | 19.7 | 14.0 | 10.3 | 22.7 | 16.7 | 1.0 | 6.0 | 22.0 |
| | | | 5 | 23.0 | 14.0 | 16.3 | 15.0 | 5.7 | 15.3 | 17.3 | 13.0 | 9.0 | 20.3 | 5.3 | 22.0 | 9.3 | 4.7 | 6.3 | 18.0 | 6.7 | 8.3 | 12.0 | 1.0 |
| | | | 6 | 22.7 | 22.3 | 8.7 | 16.0 | 7.7 | 9.7 | 20.0 | 19.0 | 5.7 | 20.0 | 9.3 | 16.0 | 17.7 | 4.7 | 9.0 | 30.0 | 8.0 | 5.0 | 13.7 | 10.3 |
| | | | 7 | 28.3 | 13.3 | 7.0 | 24.3 | 4.3 | 20.0 | 16.0 | 17.3 | 11.3 | 12.7 | 3.3 | 15.7 | 17.3 | 15.0 | 1.0 | 19.7 | 15.0 | 12.3 | 9.0 | 7.0 |
| | | | 8 | 19.0 | 6.7 | 10.0 | 11.3 | 15.7 | 19.0 | 8.7 | 19.5 | 21.0 | 14.0 | 0.5 | 25.0 | 7.0 | 22.0 | 6.0 | 21.0 | 20.0 | 11.5 | 7.5 | 1.5 |
| | 5 | 24-07-10 | 1 | 5.0 | 11.3 | 19.7 | 0.7 | 23.0 | 13.0 | 13.7 | 24.7 | 25.3 | 15.0 | 1.0 | 28.0 | 7.3 | 25.7 | 10.0 | 2.3 | 1.0 | 1.0 | 14.0 | 4.5 |
| | | | 2 | 8.0 | 22.7 | 5.7 | 1.0 | 27.7 | 10.3 | 10.3 | 9.0 | 28.0 | 19.3 | 5.3 | 25.7 | 18.3 | 1.0 | 19.7 | 14.0 | 2.3 | 21.7 | 15.7 | 9.0 |
| | | | 3 | 1.3 | 13.0 | 20.7 | 6.0 | 20.7 | 17.3 | 11.3 | 10.3 | 24.0 | 17.0 | 10.0 | 24.3 | 5.0 | 20.7 | 18.0 | 15.0 | 5.0 | 6.7 | 1.0 | 17.7 |
| | | | 4 | 0.0 | 7.3 | 14.0 | 1.0 | 1.3 | 31.3 | 7.7 | 29.0 | 11.7 | 22.0 | 3.3 | 15.3 | 2.0 | 1.0 | 26.3 | 21.3 | 2.0 | 10.3 | 3.3 | 3.0 |
| | | | 5 | 11.3 | 0.7 | 13.7 | 1.0 | 1.3 | 46.3 | 9.3 | 25.7 | 8.7 | 11.3 | 3.0 | 14.3 | 10.7 | 4.7 | 29.0 | 24.7 | 4.3 | 1.0 | 9.0 | 8.3 |
| | | | 6 | 12.0 | 11.3 | 19.0 | 10.3 | 20.7 | 19.7 | 20.3 | 13.0 | 27.7 | 17.3 | 7.3 | 28.0 | 1.0 | 30.7 | 15.0 | 4.3 | 1.0 | 11.7 | 8.7 | 3.0 |
| | | | 7 | 5.3 | 19.0 | 16.7 | 4.3 | 16.7 | 18.0 | 10.3 | 24.0 | 18.0 | 10.0 | 5.0 | 21.7 | 12.3 | 12.7 | 13.3 | 3.0 | 10.3 | 14.7 | 11.7 | 1.3 |
| | | | 8 | 8.0 | 13.3 | 16.7 | 10.7 | 5.0 | 40.7 | 19.0 | 18.3 | 6.7 | 14.7 | 8.0 | 18.0 | 24.3 | 11.0 | 13.3 | 0.7 | 4.0 | 20.7 | 6.0 | 1.0 |
| *DJ694*$^{L}$/+ ♀ x *w*$^{1118}$ ♂ | 6 | 24-09-28 | 1 | 11.0 | 9.3 | 7.3 | 15.7 | 26.0 | 3.0 | 14.3 | 19.3 | 8.3 | 10.3 | 4.0 | 6.7 | 3.7 | 11.3 | 10.3 | 3.0 | 9.7 | 9.7 | 6.0 | 17.3 |
| | | | 2 | 13.7 | 10.7 | 1.3 | 24.3 | 11.0 | 7.0 | 12.3 | 12.3 | 5.0 | 15.3 | 2.0 | 9.3 | 7.3 | 11.3 | 6.0 | 7.0 | 6.7 | 6.0 | 3.0 | 9.7 |
| | | | 3 | 9.7 | 5.3 | 21.0 | 11.7 | 14.0 | 6.0 | 9.3 | 12.7 | 12.0 | 3.3 | 6.0 | 11.0 | 9.7 | 7.3 | 4.0 | 3.0 | 6.3 | 8.0 | 3.7 | 4.0 |
| | | | 4 | 15.0 | 6.3 | 23.0 | 5.7 | 19.3 | 14.7 | 6.3 | 15.7 | 4.3 | 17.3 | 6.0 | 3.7 | 6.7 | 11.7 | 3.7 | 3.3 | 1.0 | 16.3 | 4.0 | 8.3 |
| | | | 5 | 6.3 | 4.7 | 14.3 | 9.7 | 9.7 | 8.0 | 8.7 | 15.7 | 13.3 | 9.0 | 4.3 | 6.3 | 7.0 | 13.3 | 3.0 | 12.7 | 1.0 | 8.0 | 9.0 | 11.3 |
| | | | 6 | 8.7 | 8.3 | 11.3 | 9.3 | 17.7 | 6.0 | 13.3 | 13.7 | 9.7 | 12.3 | 3.3 | 3.3 | 7.3 | 5.0 | 5.0 | 1.7 | 11.3 | 7.0 | 1.7 | 14.7 |
| | | | 7 | 11.3 | 9.3 | 10.0 | 13.0 | 17.3 | 9.3 | 5.0 | 9.0 | 11.0 | 8.3 | 8.7 | 4.3 | 7.7 | 12.0 | 7.3 | 6.3 | 3.7 | 9.0 | 11.7 | 4.3 |
| | | | 8 | 17.0 | 11.0 | 29.0 | 1.3 | 26.0 | 2.7 | 13.7 | 20.3 | 1.0 | 18.3 | 16.7 | 13.0 | 1.3 | 9.7 | 6.3 | 11.0 | 8.7 | 5.0 | 4.7 | 16.3 |
| | 7 | 24-09-28 | 1 | 28.0 | 6.7 | 9.3 | 23.7 | 18.0 | 1.0 | 25.7 | 7.0 | 11.3 | 11.3 | 12.0 | 6.7 | 2.7 | 15.0 | 3.0 | 7.3 | 6.7 | 5.7 | 7.0 | 11.7 |
| | | | 2 | 20.7 | 0.7 | 13.0 | 24.3 | 3.7 | 3.3 | 22.3 | 10.0 | 15.0 | 9.3 | 9.3 | 8.0 | 8.3 | 11.3 | 3.3 | 5.0 | 3.7 | 2.7 | 2.7 | 16.7 |
| | | | 3 | 18.7 | 5.7 | 8.7 | 28.0 | 7.3 | 18.3 | 14.7 | 5.0 | 12.3 | 3.7 | 5.7 | 12.0 | 11.0 | 5.0 | 9.7 | 12.7 | 1.3 | 6.3 | 4.3 | 9.3 |
| | | | 4 | 4.7 | 15.3 | 7.3 | 17.7 | 1.0 | 1.0 | 27.3 | 6.0 | 13.7 | 10.0 | 10.7 | 6.0 | 1.0 | 13.3 | 2.7 | 11.0 | 2.0 | 8.0 | 7.7 | 9.7 |
| | | | 5 | 9.0 | 10.7 | 8.7 | 1.3 | 17.3 | 6.0 | 9.3 | 11.7 | 5.3 | 10.0 | 9.7 | 6.7 | 6.7 | 4.0 | 9.7 | 1.7 | 6.7 | 6.0 | 4.7 | 12.7 |
| | | | 6 | 6.7 | 7.7 | 1.0 | 20.3 | 1.0 | 7.7 | 23.7 | 15.3 | 3.7 | 10.7 | 7.3 | 6.7 | 7.3 | 5.0 | 4.3 | 6.3 | 10.7 | 7.0 | 8.7 | 6.3 |
| | | | 7 | 0.7 | 5.3 | 0.7 | 13.0 | 15.0 | 1.3 | 17.3 | 5.0 | 9.0 | 10.3 | 1.0 | 15.3 | 3.3 | 3.3 | 12.3 | 6.7 | 2.7 | 1.7 | 2.7 | 14.7 |
| | | | 8 | 5.3 | 11.7 | 4.7 | 26.0 | 22.7 | 4.7 | 14.0 | 9.3 | 13.7 | 6.3 | 1.0 | 8.3 | 10.3 | 3.0 | 10.3 | 5.7 | 8.7 | 2.0 | 1.7 | 13.0 |

| Cross | Rep | Start Date (YYMMDD) | Vial | Days 21 | 22 | 23 | 24 | 25 | 26 | 27 | 28 | 29 | 30 | 31 | 32 | 33 | 34 | 35 | 36 | 37 | 38 | 39 | 40 |
|---|---|---|---|---|---|---|---|---|---|---|---|---|---|---|---|---|---|---|---|---|---|---|---|
| *$DJ694^L$* ♀ x *$w^{1118}$* ♂ | 7 | 24-09-28 | 1 | 2.0 | 10.0 | 1.7 | 2.3 | 6.7 | 2.0 | 2.7 | 1.7 | 1.7 | 1.3 | 7.3 | 4.3 | 1.3 | 3.3 | 1.0 | 3.3 | 2.3 | 6.0 | 3.3 | 3.0 |
| | | | 2 | 3.0 | 7.7 | 5.3 | 3.3 | 8.3 | 0.7 | 5.3 | 9.7 | 3.7 | 3.3 | 3.3 | 5.0 | 0.7 | 1.7 | 2.0 | 1.0 | 1.0 | 0.3 | 3.3 | 0.7 |
| | | | 3 | 7.7 | 2.0 | 7.0 | 5.3 | 6.0 | 6.3 | 4.0 | 4.3 | 7.3 | 1.0 | 4.0 | 10.0 | 1.7 | 6.3 | 3.3 | 4.0 | 2.0 | 8.0 | 3.7 | 4.0 |
| | | | 4 | 6.7 | 4.0 | 8.3 | 3.7 | 4.3 | 5.0 | 3.0 | 7.0 | 1.7 | 0.7 | 6.7 | 4.7 | 2.7 | 4.7 | 5.3 | 5.0 | 4.0 | 4.0 | 3.0 | 3.7 |
| | | | 5 | 1.0 | 6.7 | 1.3 | 1.7 | 5.0 | 0.7 | 3.7 | 8.7 | 5.3 | 4.0 | 5.7 | 4.0 | 4.3 | 8.3 | 4.3 | 3.3 | 7.7 | 7.0 | 7.3 | 2.0 |
| | | | 6 | 11.7 | 2.7 | 10.0 | 2.3 | 4.3 | 2.3 | 3.7 | 6.0 | 3.3 | 4.0 | 11.7 | 5.3 | 1.7 | 5.3 | 3.7 | 4.0 | 3.7 | 2.7 | 6.3 | 4.3 |
| | | | 7 | 2.7 | 7.3 | 3.3 | 1.7 | 6.7 | 2.0 | 3.3 | 7.0 | 5.3 | 4.3 | 3.7 | 5.0 | 2.0 | 9.0 | 1.3 | 6.7 | 3.0 | 1.0 | 5.7 | 4.3 |
| | | | 8 | 8.0 | 5.0 | 3.7 | 1.0 | 5.3 | 1.0 | 3.0 | 6.3 | 3.7 | 1.0 | 5.3 | 8.0 | 3.7 | 4.3 | 5.3 | 4.3 | 7.0 | 1.3 | 4.7 | 2.3 |
| *$DJ694^L$/+* ♀ x *$w^{1118}$* ♂ | 4 | 24-07-09 | 1 | 14.7 | 5.7 | 1.0 | 12.7 | 10.0 | 11.7 | 11.7 | 8.3 | 7.7 | 14.0 | 11.3 | 7.3 | 13.3 | 3.3 | 5.0 | 6.3 | 9.7 | 13.3 | 1.0 | 10.5 |
| | | | 2 | 9.3 | 7.7 | 13.0 | 8.3 | 7.7 | 7.7 | 15.3 | 6.3 | 8.0 | 4.3 | 7.0 | 3.7 | 12.0 | 7.3 | 11.3 | 5.3 | 6.7 | 7.7 | 12.7 | 9.7 |
| | | | 3 | 14.0 | 2.3 | 11.7 | 8.7 | 5.0 | 5.7 | 15.0 | 6.3 | 4.3 | 3.0 | 7.3 | 4.7 | 20.0 | 4.0 | 11.7 | 7.0 | 14.0 | 5.5 | 14.0 | 12.0 |
| | | | 4 | 10.0 | 5.3 | 13.7 | 4.0 | 16.3 | 13.3 | 6.0 | 8.3 | 7.3 | 4.7 | 5.0 | 13.0 | 13.5 | 7.5 | 1.0 | 6.0 | 22.5 | 15.0 | 7.0 | 16.0 |
| | | | 5 | 9.3 | 3.3 | 16.3 | 4.7 | 11.0 | 8.3 | 8.0 | 1.0 | 15.3 | 3.3 | 11.3 | 2.0 | 5.0 | 10.7 | 10.0 | 0.7 | 2.0 | 8.3 | 9.7 | 7.3 |
| | | | 6 | 10.7 | 1.7 | 10.3 | 9.0 | 12.0 | 12.0 | 7.7 | 6.7 | 17.3 | 2.7 | 5.3 | 6.0 | 12.0 | 14.5 | 6.5 | 7.5 | 17.5 | 9.0 | 7.0 | 13.5 |
| | | | 7 | 4.3 | 1.0 | 22.3 | 8.0 | 9.0 | 9.0 | 8.3 | 6.0 | 6.0 | 3.3 | 20.3 | 7.3 | 8.7 | 16.0 | 4.7 | 1.0 | 6.7 | 18.7 | 3.3 | 2.7 |
| | | | 8 | 19.5 | 1.0 | 1.0 | 26.5 | 12.5 | 7.5 | 17.5 | 3.5 | 1.0 | 18.0 | 8.0 | 18.0 | 4.5 | 6.5 | 6.0 | 13.5 | 6.5 | 12.5 | 11.5 | 7.0 |
| | 5 | 24-07-10 | 1 | 11.5 | 1.5 | 3.5 | 21.0 | 21.0 | 20.0 | 4.0 | 1.5 | 1.0 | 7.5 | 5.0 | 10.0 | 14.5 | 6.5 | 13.5 | 7.5 | 1.0 | 15.0 | 5.0 | 8.5 |
| | | | 2 | 0.7 | 1.0 | 21.7 | 13.0 | 11.7 | 11.7 | 4.7 | 6.0 | 0.7 | 18.0 | 5.0 | 11.0 | 1.0 | 11.3 | 14.0 | 8.3 | 10.0 | 12.0 | 6.0 | 6.3 |
| | | | 3 | 16.7 | 8.0 | 11.3 | 14.3 | 14.0 | 10.7 | 5.3 | 11.7 | 9.0 | 8.0 | 3.0 | 7.7 | 1.3 | 4.3 | 10.7 | 11.3 | 7.7 | 6.0 | 1.0 | 1.3 |
| | | | 4 | 1.3 | 27.3 | 6.3 | 18.0 | 16.7 | 15.0 | 6.3 | 2.7 | 9.3 | 11.3 | 7.0 | 12.7 | 1.0 | 15.0 | 2.3 | 12.7 | 3.0 | 13.3 | 1.3 | 2.7 |
| | | | 5 | 1.3 | 1.0 | 17.7 | 18.3 | 12.3 | 10.3 | 6.0 | 4.7 | 1.3 | 5.0 | 7.0 | 12.3 | 9.3 | 2.7 | 4.7 | 10.3 | 5.7 | 12.7 | 8.7 | 1.3 |
| | | | 6 | 6.7 | 9.0 | 10.0 | 13.0 | 11.7 | 8.7 | 6.7 | 1.7 | 17.3 | 15.7 | 1.3 | 9.0 | 6.3 | 5.0 | 19.3 | 8.0 | 1.0 | 27.0 | 1.3 | 10.7 |
| | | | 7 | 12.0 | 6.0 | 2.5 | 14.5 | 15.0 | 23.0 | 5.0 | 5.5 | 13.0 | 10.0 | 13.0 | 9.0 | 6.0 | 6.0 | 4.0 | 1.0 | 6.0 | 5.0 | 5.0 | 1.0 |
| | | | 8 | 14.3 | 0.7 | 18.7 | 12.5 | 17.5 | 15.0 | 11.0 | 6.0 | 3.5 | 17.5 | 9.5 | 5.5 | 4.5 | 3.0 | 13.0 | 11.5 | 0.5 | 10.5 | 9.5 | 1.0 |
| *$DJ694^L$/+* ♀ x *$w^{1118}$* ♂ | 6 | 24-09-28 | 1 | 2.3 | 10.0 | 1.3 | 5.0 | 3.0 | 10.3 | 3.7 | 8.7 | 3.0 | 5.3 | 3.0 | 5.7 | 10.0 | 2.0 | 6.3 | 4.0 | 1.7 | 1.3 | 7.0 | 1.7 |
| | | | 2 | 4.7 | 9.3 | 1.0 | 5.3 | 7.3 | 1.3 | 5.0 | 1.7 | 3.7 | 7.7 | 3.7 | 2.7 | 4.7 | 8.3 | 6.0 | 3.3 | 3.0 | 3.7 | 5.0 | 6.7 |
| | | | 3 | 9.7 | 7.3 | 1.3 | 3.3 | 1.0 | 9.7 | 4.3 | 7.0 | 2.3 | 1.7 | 10.7 | 6.7 | 6.3 | 4.7 | 7.7 | 5.3 | 1.3 | 1.3 | 6.7 | 7.0 |
| | | | 4 | 9.7 | 10.0 | 0.7 | 4.0 | 3.3 | 10.0 | 2.0 | 5.0 | 2.7 | 3.7 | 7.3 | 5.7 | 5.0 | 6.7 | 6.7 | 5.0 | 6.0 | 3.7 | 4.0 | 4.3 |
| | | | 5 | 1.3 | 4.3 | 4.7 | 1.0 | 6.7 | 5.0 | 3.3 | 6.0 | 7.3 | 1.7 | 4.7 | 5.0 | 8.0 | 7.3 | 10.7 | 5.7 | 3.3 | 1.7 | 6.7 | 1.7 |
| | | | 6 | 2.0 | 3.0 | 3.3 | 2.3 | 12.0 | 3.0 | 7.0 | 6.7 | 2.7 | 7.0 | 5.7 | 3.0 | 6.3 | 6.0 | 4.7 | 6.7 | 2.7 | 6.3 | 4.3 | 3.3 |
| | | | 7 | 12.3 | 2.0 | 6.7 | 3.0 | 6.3 | 3.3 | 4.3 | 5.0 | 5.7 | 4.3 | 3.7 | 6.0 | 7.3 | 6.3 | 3.3 | 5.7 | 2.0 | 2.0 | 5.0 | 0.3 |
| | | | 8 | 3.3 | 8.3 | 5.5 | 0.7 | 6.3 | 4.3 | 1.7 | 8.3 | 5.7 | 1.0 | 2.3 | 11.3 | 3.7 | 4.7 | 5.7 | 1.0 | 3.7 | 0.3 | 0.7 | 1.3 |
| | 7 | 24-09-28 | 1 | 2.0 | 10.0 | 1.7 | 2.3 | 6.7 | 2.0 | 2.7 | 1.7 | 1.7 | 1.3 | 7.3 | 4.3 | 1.3 | 3.3 | 1.0 | 3.3 | 2.3 | 6.0 | 3.3 | 3.0 |
| | | | 2 | 3.0 | 7.7 | 5.3 | 3.3 | 8.3 | 0.7 | 5.3 | 9.7 | 3.7 | 3.3 | 3.3 | 5.0 | 0.7 | 1.7 | 2.0 | 1.0 | 1.0 | 0.3 | 3.3 | 0.7 |
| | | | 3 | 7.7 | 2.0 | 7.0 | 5.3 | 6.0 | 6.3 | 4.0 | 4.3 | 7.3 | 1.0 | 4.0 | 10.0 | 1.7 | 6.3 | 3.3 | 4.0 | 2.0 | 8.0 | 3.7 | 4.0 |
| | | | 4 | 6.7 | 4.0 | 8.3 | 3.7 | 4.3 | 5.0 | 3.0 | 7.0 | 1.7 | 0.7 | 6.7 | 4.7 | 2.7 | 4.7 | 5.3 | 5.0 | 4.0 | 4.0 | 3.0 | 3.7 |
| | | | 5 | 1.0 | 6.7 | 1.3 | 1.7 | 5.0 | 0.7 | 3.7 | 8.7 | 5.3 | 4.0 | 5.7 | 4.0 | 4.3 | 8.3 | 4.3 | 3.3 | 7.7 | 7.0 | 7.3 | 2.0 |
| | | | 6 | 11.7 | 2.7 | 10.0 | 2.3 | 4.3 | 2.3 | 3.7 | 6.0 | 3.3 | 4.0 | 11.7 | 5.3 | 1.7 | 5.3 | 3.7 | 4.0 | 3.7 | 2.7 | 6.3 | 4.3 |
| | | | 7 | 2.7 | 7.3 | 3.3 | 1.7 | 6.7 | 2.0 | 3.3 | 7.0 | 5.3 | 4.3 | 3.7 | 5.0 | 2.0 | 9.0 | 1.3 | 6.7 | 3.0 | 1.0 | 5.7 | 4.3 |
| | | | 8 | 8.0 | 5.0 | 3.7 | 1.0 | 5.3 | 1.0 | 3.0 | 6.3 | 3.7 | 1.0 | 5.3 | 8.0 | 3.7 | 4.3 | 5.3 | 4.3 | 7.0 | 1.3 | 4.7 | 2.3 |

| Cross | Rep | Start Date (YYMMDD) | Vial | Days 1 | 2 | 3 | 4 | 5 | 6 | 7 | 8 | 9 | 10 | 11 | 12 | 13 | 14 | 15 | 16 | 17 | 18 | 19 | 20 |
|---|---|---|---|---|---|---|---|---|---|---|---|---|---|---|---|---|---|---|---|---|---|---|---|
| *$DJ694^{O}$ ♀ x $w^{1118}$ ♂* | 4 | 24-07-09 | 1 | 2.0 | 0.7 | 0.3 | 6.0 | 11.3 | 4.7 | 37.3 | 4.0 | 18.3 | 3.0 | 3.0 | 1.0 | 11.0 | 13.0 | 1.0 | 11.0 | 20.0 | 10.0 | 1.0 | 21.0 |
| | | | 2 | 2.0 | 1.3 | 0.7 | 2.7 | 5.0 | 3.7 | 16.7 | 23.3 | 4.0 | 2.3 | 4.0 | 7.3 | 23.3 | 3.7 | 1.0 | 16.7 | 13.3 | 0.7 | 1.7 | 3.7 |
| | | | 3 | 7.7 | 2.0 | 0.7 | 5.7 | 7.0 | 9.3 | 17.0 | 5.0 | 1.3 | 6.0 | 0.7 | 16.3 | 7.0 | 1.3 | 10.0 | 10.0 | 2.7 | 1.7 | 0.7 | 10.0 |
| | | | 4 | 3.3 | 1.0 | 2.0 | 15.7 | 5.7 | 15.7 | 25.3 | 5.3 | 0.7 | 1.0 | 1.0 | 7.7 | 18.7 | 11.7 | 8.3 | 10.0 | 5.0 | 12.0 | 2.0 | 9.7 |
| | | | 5 | 14.7 | 12.0 | 1.3 | 27.3 | 0.7 | 7.7 | 6.7 | 1.7 | 0.7 | 1.3 | 3.3 | 21.0 | 1.0 | 8.5 | 2.5 | 6.0 | 9.5 | 3.5 | 0.0 | 1.5 |
| | | | 6 | 1.0 | 0.7 | 0.3 | 0.3 | 0.7 | 2.0 | 20.0 | 4.7 | 0.3 | 1.0 | 7.7 | 4.7 | 2.0 | 2.7 | 23.7 | 7.3 | 13.0 | 2.3 | 0.7 | 24.0 |
| | | | 7 | 11.3 | 1.0 | 0.7 | 2.3 | 0.7 | 5.7 | 35.7 | 3.3 | 6.7 | 0.7 | 29.3 | 2.0 | 8.7 | 1.0 | 11.0 | 3.3 | 1.7 | 1.7 | 8.0 | 9.0 |
| | | | 8 | 4.3 | 0.3 | 2.0 | 2.7 | 1.3 | 12.3 | 21.3 | 28.7 | 11.0 | 4.0 | 2.7 | 16.3 | 1.7 | 17.0 | 3.0 | 9.7 | 3.3 | 17.7 | 2.3 | 2.0 |
| | 5 | 24-07-10 | 1 | 0.0 | 0.7 | 1.0 | 1.0 | 1.0 | 19.0 | 4.7 | 23.3 | 10.0 | 1.0 | 19.3 | 4.0 | 2.7 | 11.3 | 13.0 | 13.0 | 0.7 | 4.3 | 21.3 | 2.0 |
| | | | 2 | 0.0 | 6.0 | 0.7 | 1.0 | 0.3 | 16.0 | 11.3 | 1.3 | 22.0 | 1.7 | 18.7 | 10.7 | 1.0 | 0.7 | 14.7 | 13.3 | 0.7 | 1.3 | 16.0 | 1.7 |
| | | | 3 | 0.0 | 0.7 | 0.7 | 0.7 | 0.3 | 31.0 | 1.0 | 0.7 | 16.7 | 0.3 | 11.7 | 1.7 | 5.3 | 5.7 | 10.3 | 8.7 | 3.7 | 0.7 | 4.0 | 12.0 |
| | | | 4 | 0.0 | 0.3 | 1.0 | 1.0 | 0.7 | 2.7 | 2.0 | 6.7 | 11.7 | 1.3 | 1.0 | 20.3 | 0.7 | 1.3 | 12.0 | 7.7 | 1.0 | 0.7 | 1.3 | 1.7 |
| *$DJ694^{O}$ /+ ♀ x $w^{1118}$ ♂* | 4 | 24-07-09 | 1 | 27.7 | 3.7 | 0.7 | 27.3 | 18.7 | 4.0 | 33.0 | 11.3 | 6.0 | 19.3 | 17.3 | 12.0 | 0.7 | 25.0 | 7.0 | 33.3 | 5.0 | 6.3 | 8.0 | 20.0 |
| | | | 2 | 28.0 | 16.7 | 1.3 | 19.3 | 0.5 | 29.5 | 20.5 | 16.5 | 24.5 | 1.0 | 39.5 | 0.5 | 28.5 | 4.5 | 17.0 | 28.0 | 13.0 | 10.0 | 1.0 | 22.5 |
| | | | 3 | 13.3 | 0.7 | 0.7 | 27.0 | 12.7 | 5.3 | 27.0 | 15.7 | 1.0 | 26.7 | 24.0 | 3.7 | 15.0 | 26.0 | 8.3 | 41.0 | 10.0 | 4.0 | 1.0 | 23.0 |
| | | | 4 | 21.7 | 0.3 | 14.0 | 10.3 | 1.0 | 24.7 | 19.7 | 16.7 | 12.0 | 41.3 | 0.0 | 40.7 | 8.7 | 19.0 | 9.7 | 35.7 | 11.7 | 6.3 | 6.0 | 3.7 |
| | | | 5 | 39.3 | 1.3 | 25.0 | 6.7 | 27.3 | 1.3 | 23.3 | 25.3 | 4.7 | 31.0 | 0.7 | 45.3 | 12.7 | 1.0 | 34.0 | 36.3 | 11.7 | 9.3 | 1.3 | 1.0 |
| | | | 6 | 25.0 | 10.0 | 1.0 | 18.7 | 5.0 | 29.7 | 13.3 | 12.7 | 22.7 | 12.0 | 27.0 | 25.3 | 6.3 | 26.3 | 10.0 | 15.0 | 17.3 | 5.3 | 12.3 | 6.0 |
| | | | 7 | 32.0 | 24.3 | 15.3 | 4.3 | 1.0 | 16.7 | 28.0 | 18.3 | 14.7 | 12.0 | 0.5 | 39.5 | 11.0 | 1.5 | 32.0 | 20.0 | 20.0 | 7.0 | 7.5 | 7.0 |
| | | | 8 | 24.3 | 10.0 | 28.0 | 4.3 | 0.7 | 15.7 | 29.3 | 18.7 | 25.7 | 23.7 | 6.3 | 2.3 | 36.3 | 12.3 | 11.0 | 23.0 | 23.0 | 6.7 | 0.7 | 5.7 |
| *$DJ694^{O}$ /+ ♀ x $w^{1118}$ ♂* | 5 | 24-07-10 | 1 | 0.0 | 5.3 | 23.0 | 1.0 | 29.0 | 17.7 | 28.7 | 0.7 | 24.0 | 0.7 | 1.0 | 42.7 | 1.0 | 33.3 | 14.3 | 8.3 | 1.0 | 1.0 | 17.0 | 1.3 |
| | | | 2 | 3.3 | 14.7 | 1.0 | 3.0 | 26.7 | 10.7 | 19.7 | 15.0 | 5.0 | 15.3 | 21.3 | 25.0 | 8.0 | 1.0 | 21.3 | 12.0 | 1.0 | 12.3 | 4.0 | 9.0 |
| | | | 3 | 0.0 | 4.0 | 11.7 | 5.0 | 22.3 | 22.0 | 21.3 | 17.0 | 8.0 | 14.7 | 23.0 | 22.0 | 7.3 | 7.7 | 13.7 | 10.0 | 5.0 | 11.0 | 6.3 | 1.3 |
| | | | 4 | 2.3 | 11.7 | 10.3 | 7.7 | 13.3 | 26.7 | 10.0 | 11.0 | 25.3 | 4.3 | 1.0 | 0.7 | 5.3 | 24.7 | 12.0 | 14.7 | 2.3 | 7.0 | 9.0 | 3.0 |
| | | | 5 | 4.3 | 18.7 | 1.3 | 1.0 | 30.3 | 7.7 | 28.0 | 4.3 | 29.3 | 3.7 | 1.3 | 42.0 | 25.3 | 4.0 | 15.0 | 10.7 | 2.7 | 9.7 | 4.3 | 1.3 |
| | | | 6 | 0.0 | 11.0 | 1.5 | 2.5 | 58.5 | 30.5 | 2.0 | 27.0 | 3.5 | 0.5 | 42.0 | 18.0 | 1.0 | 33.0 | 15.5 | 10.0 | 1.5 | 17.0 | 1.0 | 1.5 |
| | | | 7 | 0.0 | 21.7 | 8.7 | 16.7 | 4.0 | 20.7 | 8.3 | 21.7 | 14.7 | 13.0 | 19.0 | 5.0 | 12.5 | 22.0 | 20.5 | 7.5 | 2.5 | 19.0 | 7.0 | 1.5 |
| *$DJ694^{P}$ ♀ x $w^{1118}$ ♂* | 4 | 24-07-09 | 1 | 2.5 | 1.0 | 0.5 | 4.0 | 11.0 | 13.0 | 26.0 | 2.0 | 1.5 | 27.0 | 1.0 | 24.0 | 28.0 | 1.0 | 26.0 | 12.0 | 24.0 | 10.0 | 7.0 | 19.0 |
| | | | 2 | 7.3 | 1.3 | 8.3 | 4.3 | 9.3 | 9.7 | 31.7 | 1.3 | 4.0 | 20.3 | 0.3 | 18.7 | 13.3 | 5.7 | 18.7 | 14.3 | 9.7 | 1.0 | 8.7 | 14.3 |
| | | | 3 | 5.7 | 5.0 | 8.3 | 5.0 | 5.3 | 8.0 | 34.3 | 2.0 | 1.0 | 1.0 | 6.7 | 21.0 | 14.7 | 4.0 | 18.0 | 13.3 | 10.0 | 2.3 | 6.7 | 7.3 |
| | | | 4 | 2.3 | 1.0 | 3.7 | 0.7 | 1.0 | 2.7 | 11.0 | 8.7 | 5.7 | 0.7 | 2.0 | 26.7 | 2.7 | 0.7 | 24.0 | 6.7 | 7.0 | 2.7 | 4.7 | 11.0 |
| | | | 5 | 3.0 | 0.7 | 5.3 | 3.0 | 4.3 | 37.7 | 1.7 | 6.7 | 11.7 | 3.3 | 4.3 | 24.0 | 13.3 | 2.7 | 8.7 | 20.0 | 10.3 | 6.3 | 2.7 | 7.7 |
| | | | 6 | 1.0 | 1.0 | 6.3 | 1.3 | 0.0 | 1.7 | 34.7 | 0.7 | 4.0 | 21.3 | 4.3 | 26.7 | 5.7 | 1.0 | 20.3 | 11.0 | 13.0 | 5.3 | 0.0 | 9.0 |
| | | | 7 | 1.3 | 0.7 | 1.3 | 7.7 | 2.0 | 13.0 | 31.3 | 7.7 | 4.0 | 6.3 | 4.3 | 32.7 | 1.0 | 1.3 | 29.7 | 11.3 | 16.3 | 5.3 | 1.3 | 3.3 |
| | | | 8 | 0.7 | 0.3 | 11.0 | 9.0 | 6.3 | 21.7 | 18.3 | 7.7 | 9.0 | 4.7 | 7.7 | 26.3 | 11.7 | 6.3 | 4.0 | 13.0 | 6.0 | 15.0 | 5.7 | 5.7 |
| | 5 | 24-07-10 | 1 | 0.0 | 0.7 | 1.3 | 0.7 | 0.7 | 13.7 | 13.3 | 0.3 | 16.7 | 0.3 | 5.3 | 1.3 | 8.3 | 7.3 | 13.3 | 15.3 | 1.0 | 1.3 | 0.7 | 1.0 |
| | | | 2 | 0.0 | 0.7 | 1.7 | 0.7 | 3.3 | 6.3 | 10.0 | 0.7 | 18.0 | 0.3 | 2.0 | 0.7 | 0.3 | 0.7 | 6.7 | 6.7 | 0.0 | 0.7 | 1.0 | 0.3 |
| | | | 3 | 0.0 | 3.0 | 1.0 | 0.0 | 0.3 | 1.0 | 20.7 | 0.7 | 6.7 | 0.3 | 0.3 | 6.0 | 3.0 | 1.3 | 6.0 | 6.0 | 0.0 | 9.3 | 2.0 | 0.0 |
| | | | 4 | 0.0 | 0.0 | 1.7 | 1.0 | 1.0 | 0.7 | 15.7 | 1.3 | 0.7 | 0.7 | 1.0 | 18.3 | 0.3 | 0.3 | 7.7 | 8.3 | 1.0 | 5.7 | 9.7 | 0.0 |

| | | Start Date | | Days | | | | | | | | | | | | | | | | | | | |
|---|---|---|---|---|---|---|---|---|---|---|---|---|---|---|---|---|---|---|---|---|---|---|---|
| Cross | Rep | (YYMMDD) | Vial | 21 | 22 | 23 | 24 | 25 | 26 | 27 | 28 | 29 | 30 | 31 | 32 | 33 | 34 | 35 | 36 | 37 | 38 | 39 | 40 |
| $DJ694^{O}$ ♀ x $w^{1118}$ ♂ | 4 | 24-07-09 | 1 | 7.0 | 0.0 | 0.0 | 13.0 | 11.0 | 12.0 | 13.0 | 0.0 | 3.0 | 5.0 | 1.0 | 0.0 | 1.0 | 2.0 | 4.0 | 3.0 | 3.0 | 2.0 | 0.0 | 0.0 |
| | | | 2 | 11.3 | 2.3 | 1.3 | 13.0 | 1.3 | 1.7 | 5.3 | 3.7 | 1.7 | 4.0 | 1.5 | 7.0 | 12.0 | 0.5 | 1.0 | 1.0 | 0.5 | 2.0 | 4.0 | 4.0 |
| | | | 3 | 1.3 | 0.0 | 1.3 | 3.7 | 4.0 | 0.0 | 1.5 | 1.0 | 1.0 | 0.0 | 0.0 | 0.0 | 0.0 | 0.0 | 0.0 | 0.0 | 0.0 | 0.0 | 0.0 | 0.0 |
| | | | 4 | 4.0 | 1.0 | 27.0 | 6.0 | 2.7 | 2.7 | 7.3 | 6.3 | 2.0 | 1.0 | 3.3 | 7.3 | 7.0 | 3.5 | 2.0 | 5.5 | 0.0 | 5.0 | 5.0 | 3.0 |
| | | | 5 | 2.0 | 0.0 | 1.0 | 2.5 | 1.5 | 0.0 | 4.0 | 3.0 | 1.0 | 1.0 | 1.0 | 4.0 | 0.0 | 0.0 | 0.0 | 0.0 | 0.0 | 0.0 | 1.0 | 0.0 |
| | | | 6 | 2.7 | 3.3 | 21.7 | 9.3 | 3.3 | 3.0 | 18.3 | 8.0 | 0.7 | 1.7 | 7.3 | 3.7 | 8.7 | 3.3 | 1.7 | 1.3 | 2.7 | 2.3 | 3.7 | 2.0 |
| | | | 7 | 3.3 | 1.0 | 1.0 | 2.0 | 9.7 | 10.0 | 4.3 | 2.0 | 0.3 | 1.0 | 1.7 | 5.0 | 7.7 | 0.7 | 1.0 | 1.0 | 5.7 | 4.0 | 1.3 | 0.3 |
| | | | 8 | 11.7 | 1.3 | 4.0 | 11.0 | 8.0 | 7.7 | 16.0 | 12.0 | 3.5 | 1.0 | 6.0 | 7.5 | 7.5 | 1.5 | 9.0 | 3.0 | 8.5 | 2.5 | 9.5 | 1.0 |
| | 5 | 24-07-10 | 1 | 1.7 | 3.3 | 3.7 | 13.0 | 7.0 | 7.3 | 7.0 | 0.7 | 4.3 | 1.0 | 4.7 | 8.0 | 1.3 | 1.7 | 5.0 | 6.0 | 4.7 | 1.0 | 1.7 | 1.7 |
| | | | 2 | 2.3 | 2.7 | 3.0 | 8.0 | 8.0 | 4.0 | 7.3 | 2.0 | 3.0 | 1.0 | 12.7 | 3.7 | 0.3 | 0.3 | 1.0 | 1.3 | 1.3 | 2.3 | 3.3 | 0.0 |
| | | | 3 | 15.0 | 7.3 | 13.0 | 10.3 | 5.0 | 4.3 | 5.3 | 2.3 | 5.0 | 13.3 | 2.7 | 1.3 | 2.0 | 8.0 | 2.3 | 0.3 | 2.0 | 2.0 | 3.7 | 1.0 |
| | | | 4 | 1.7 | 5.3 | 7.3 | 3.3 | 2.7 | 4.0 | 3.3 | 3.3 | 3.0 | 2.3 | 1.3 | 2.7 | 1.3 | 3.3 | 3.3 | 4.0 | 3.0 | 1.7 | 1.7 | 1.0 |
| $DJ694^{O}$ /+ ♀ x $w^{1118}$ ♂ | 4 | 24-07-09 | 1 | 5.3 | 15.7 | 1.0 | 15.3 | 13.7 | 12.3 | 11.3 | 4.0 | 3.0 | 5.3 | 17.0 | 9.7 | 7.0 | 3.7 | 16.0 | 9.3 | 5.3 | 6.7 | 7.7 | 14.0 |
| | | | 2 | 4.5 | 18.5 | 1.0 | 1.0 | 19.0 | 18.5 | 13.5 | 8.5 | 15.5 | 4.5 | 1.0 | 15.5 | 15.0 | 1.0 | 11.5 | 13.5 | 8.0 | 9.5 | 14.0 | 9.0 |
| | | | 3 | 9.5 | 1.0 | 1.0 | 1.0 | 26.0 | 18.0 | 16.5 | 6.0 | 9.0 | 1.0 | 14.0 | 2.5 | 15.5 | 7.5 | 11.5 | 13.5 | 1.0 | 30.5 | 5.5 | 19.5 |
| | | | 4 | 3.3 | 3.7 | 25.7 | 10.0 | 7.0 | 7.0 | 6.0 | 16.7 | 2.7 | 1.0 | 11.0 | 8.7 | 15.3 | 1.0 | 12.7 | 22.0 | 9.7 | 7.3 | 15.0 | 6.3 |
| | | | 5 | 1.0 | 12.3 | 23.0 | 4.3 | 13.0 | 4.0 | 16.3 | 11.3 | 1.0 | 1.7 | 12.7 | 3.0 | 24.3 | 4.0 | 1.0 | 22.0 | 10.7 | 9.7 | 7.0 | 7.3 |
| | | | 6 | 1.3 | 18.7 | 5.3 | 3.3 | 19.7 | 20.0 | 14.3 | 4.0 | 4.0 | 5.0 | 13.7 | 4.0 | 6.0 | 21.0 | 4.3 | 8.7 | 5.7 | 20.0 | 2.7 | 12.7 |
| | | | 7 | 1.0 | 26.5 | 6.0 | 1.0 | 19.5 | 19.0 | 9.5 | 18.5 | 1.5 | 6.5 | 27.5 | 2.0 | 0.0 | 13.0 | 1.0 | 12.0 | 8.5 | 30.0 | 4.0 | 19.0 |
| | | | 8 | 1.0 | 15.3 | 4.7 | 22.3 | 10.3 | 10.3 | 11.3 | 3.7 | 7.3 | 1.0 | 12.7 | 7.7 | 13.0 | 1.0 | 5.0 | 14.0 | 8.0 | 19.7 | 7.3 | 5.7 |
| $DJ694^{O}$ /+ ♀ x $w^{1118}$ ♂ | 5 | 24-07-10 | 1 | 7.0 | 17.3 | 1.0 | 18.3 | 20.0 | 18.3 | 6.0 | 3.3 | 13.3 | 9.0 | 1.0 | 17.3 | 2.3 | 7.0 | 2.0 | 29.3 | 1.3 | 1.3 | 16.3 | 8.0 |
| | | | 2 | 6.0 | 5.3 | 19.0 | 16.0 | 15.7 | 6.7 | 8.3 | 1.0 | 6.0 | 1.0 | 30.0 | 1.3 | 3.7 | 10.7 | 19.7 | 16.3 | 6.0 | 9.3 | 2.0 | 3.7 |
| | | | 3 | 0.7 | 10.0 | 15.0 | 14.7 | 6.0 | 11.0 | 6.7 | 2.3 | 0.7 | 18.3 | 1.0 | 17.5 | 0.5 | 1.0 | 32.5 | 6.5 | 9.5 | 1.0 | 12.0 | 2.0 |
| | | | 4 | 7.0 | 1.3 | 32.0 | 7.0 | 7.0 | 17.0 | 9.0 | 1.0 | 1.3 | 31.7 | 1.0 | 1.0 | 7.7 | 4.0 | 31.3 | 0.3 | 6.7 | 1.3 | 19.0 | 3.0 |
| | | | 5 | 23.0 | 4.7 | 1.0 | 8.3 | 6.7 | 12.7 | 9.7 | 1.0 | 1.0 | 16.0 | 24.7 | 7.0 | 1.0 | 1.0 | 15.0 | 17.0 | 1.0 | 25.7 | 9.3 | 2.0 |
| | | | 6 | 1.0 | 12.5 | 11.0 | 31.5 | 9.0 | 13.5 | 8.5 | 1.0 | 14.0 | 9.0 | 2.0 | 9.0 | 30.5 | 1.0 | 1.0 | 18.0 | 16.0 | 21.0 | 13.5 | 1.0 |
| | | | 7 | 1.0 | 23.0 | 2.0 | 27.0 | 11.0 | 14.0 | 0.5 | 9.5 | 3.0 | 15.5 | 1.0 | 22.5 | 3.0 | 2.0 | 20.0 | 17.5 | 0.5 | 12.0 | 11.0 | 7.5 |
| $DJ694^{P}$ ♀ x $w^{1118}$ ♂ | 4 | 24-07-09 | 1 | 2.0 | 0.0 | 15.0 | 17.0 | 15.0 | 3.0 | 4.0 | 0.0 | 0.0 | 2.0 | 4.0 | 0.0 | 3.0 | 0.0 | 1.0 | 0.0 | 0.0 | 1.0 | 0.0 | 0.0 |
| | | | 2 | 10.7 | 0.7 | 5.3 | 8.3 | 2.0 | 13.3 | 8.0 | 6.0 | 12.3 | 6.0 | 7.3 | 5.0 | 5.7 | 2.3 | 3.7 | 2.7 | 4.0 | 4.3 | 3.3 | 2.0 |
| | | | 3 | 6.0 | 10.7 | 6.7 | 18.0 | 8.3 | 8.7 | 8.3 | 6.7 | 2.3 | 1.7 | 11.7 | 8.3 | 7.0 | 2.3 | 2.7 | 1.0 | 2.3 | 1.3 | 3.0 | 1.7 |
| | | | 4 | 1.7 | 1.0 | 3.3 | 3.7 | 9.3 | 9.7 | 2.7 | 0.3 | 0.7 | 4.0 | 1.0 | 0.3 | 2.0 | 1.7 | 1.3 | 1.0 | 1.3 | 1.7 | 0.3 | 1.3 |
| | | | 5 | 1.7 | 3.3 | 3.0 | 9.3 | 2.7 | 3.0 | 10.0 | 3.0 | 4.7 | 3.0 | 5.7 | 1.3 | 1.3 | 1.3 | 0.7 | 3.7 | 1.3 | 0.7 | 4.3 | 1.3 |
| | | | 6 | 7.5 | 3.0 | 2.0 | 7.0 | 8.5 | 5.0 | 8.5 | 8.0 | 0.0 | 0.5 | 3.5 | 2.0 | 2.5 | 0.5 | 2.5 | 1.0 | 0.0 | 3.0 | 0.0 | 0.0 |
| | | | 7 | 3.3 | 2.0 | 17.3 | 3.0 | 4.5 | 4.5 | 7.0 | 2.0 | 4.0 | 6.0 | 1.5 | 6.5 | 7.0 | 1.0 | 1.5 | 2.5 | 1.5 | 1.0 | 2.0 | 0.5 |
| | | | 8 | 6.3 | 11.0 | 1.3 | 2.0 | 2.0 | 5.3 | 11.7 | 3.7 | 3.0 | 2.5 | 5.0 | 4.0 | 0.5 | 4.0 | 4.0 | 3.0 | 2.5 | 3.0 | 1.5 | 1.0 |
| | 5 | 24-07-10 | 1 | 7.3 | 1.3 | 2.3 | 15.0 | 14.7 | 3.0 | 3.3 | 1.0 | 0.3 | 1.0 | 2.0 | 0.5 | 2.0 | 12.0 | 6.0 | 3.0 | 3.0 | 3.5 | 2.5 | 0.5 |
| | | | 2 | 15.0 | 0.0 | 3.0 | 2.3 | 2.7 | 18.0 | 2.3 | 0.7 | 2.7 | 0.3 | 0.7 | 1.0 | 0.7 | 1.0 | 2.0 | 8.0 | 1.3 | 7.0 | 9.7 | 9.7 |
| | | | 3 | 10.0 | 2.5 | 9.0 | 5.0 | 6.0 | 6.0 | 10.5 | 2.0 | 1.0 | 3.5 | 16.5 | 8.0 | 1.0 | 0.5 | 6.0 | 8.0 | 9.0 | 3.5 | 4.5 | 2.0 |
| | | | 4 | 0.3 | 0.3 | 8.7 | 5.0 | 2.0 | 5.0 | 6.3 | 0.7 | 0.7 | 0.3 | 6.3 | 0.3 | 0.7 | 0.3 | 3.7 | 6.7 | 0.3 | 1.0 | 1.3 | 1.0 |

| Cross | Rep | Start Date (YYMMDD) | Vial | Days 1 | 2 | 3 | 4 | 5 | 6 | 7 | 8 | 9 | 10 | 11 | 12 | 13 | 14 | 15 | 16 | 17 | 18 | 19 | 20 |
|---|---|---|---|---|---|---|---|---|---|---|---|---|---|---|---|---|---|---|---|---|---|---|---|
| *$DJ694^{P}/+$* ♀ x *$w^{1118}$* ♂ | 4 | 24-07-09 | 1 | 19.3 | 6.3 | 45.3 | 3.7 | 1.0 | 22.7 | 33.3 | 23.3 | 8.3 | 17.0 | 7.7 | 21.0 | 18.7 | 13.7 | 13.0 | 15.0 | 17.7 | 6.3 | 10.7 | 7.0 |
| | | | 2 | 19.7 | 5.3 | 25.7 | 19.0 | 8.7 | 12.0 | 18.3 | 14.3 | 21.7 | 34.0 | 6.7 | 32.7 | 7.3 | 3.3 | 27.7 | 25.3 | 9.0 | 9.0 | 2.3 | 1.7 |
| | | | 3 | 40.7 | 16.3 | 36.3 | 2.0 | 7.0 | 0.7 | 32.0 | 10.0 | 24.0 | 19.7 | 1.0 | 21.7 | 20.0 | 1.0 | 21.0 | 20.7 | 10.3 | 1.0 | 7.3 | 11.7 |
| | 5 | 24-07-10 | 1 | 0.0 | 3.7 | 15.3 | 13.0 | 11.3 | 18.3 | 16.7 | 27.7 | 19.3 | 5.7 | 27.7 | 6.3 | 1.3 | 30.0 | 6.7 | 6.7 | 3.0 | 10.0 | 12.3 | 1.7 |
| | | | 2 | 6.7 | 1.0 | 5.7 | 11.0 | 19.0 | 16.7 | 20.3 | 10.7 | 9.3 | 9.7 | 21.3 | 22.7 | 3.3 | 20.0 | 18.0 | 9.0 | 9.3 | 5.0 | 22.0 | 1.0 |
| | | | 3 | 0.0 | 10.0 | 31.0 | 1.0 | 17.0 | 6.3 | 18.3 | 18.7 | 27.0 | 14.3 | 21.7 | 12.7 | 1.0 | 18.0 | 18.7 | 15.3 | 1.0 | 7.3 | 17.3 | 1.3 |
| | | | 4 | 0.0 | 19.7 | 14.7 | 1.0 | 1.3 | 47.0 | 1.7 | 18.0 | 11.7 | 0.3 | 23.3 | 1.0 | 0.7 | 38.7 | 11.0 | 10.3 | 1.0 | 15.7 | 1.3 | 1.3 |
| | | | 5 | 0.0 | 8.0 | 14.3 | 1.0 | 1.0 | 49.3 | 3.7 | 27.7 | 13.0 | 4.3 | 9.0 | 33.7 | 1.0 | 13.0 | 17.3 | 14.3 | 1.3 | 21.0 | 5.3 | 9.3 |
| | | | 6 | 1.7 | 9.0 | 18.7 | 3.0 | 4.0 | 43.0 | 3.3 | 28.7 | 24.0 | 13.7 | 17.7 | 4.7 | 15.3 | 17.7 | 10.3 | 9.3 | 1.0 | 1.0 | 23.7 | 1.7 |
| | | | 7 | 0.0 | 1.3 | 20.3 | 26.3 | 1.0 | 27.7 | 11.7 | 23.3 | 4.7 | 0.7 | 11.3 | 25.0 | 5.7 | 15.7 | 22.0 | 19.0 | 5.0 | 12.0 | 1.0 | 1.3 |
| *$w^{1118}$* ♀ x *$w^{1118}$* ♂ | 4 | 24-07-09 | 1 | 10.3 | 0.7 | 28.7 | 8.3 | 7.0 | 13.0 | 16.0 | 2.3 | 2.5 | 32.5 | 8.5 | 12.0 | 2.0 | 35.5 | 15.5 | 25.0 | 2.0 | 1.0 | 8.5 | 28.0 |
| | | | 2 | 27.5 | 1.0 | 47.5 | 21.0 | 13.0 | 5.0 | 24.5 | 23.5 | 1.0 | 26.5 | 17.5 | 20.5 | 6.0 | 4.0 | 19.0 | 17.5 | 1.0 | 12.5 | 7.0 | 1.5 |
| | | | 3 | 14.3 | 22.7 | 30.7 | 16.0 | 11.3 | 10.0 | 17.3 | 8.3 | 6.7 | 20.3 | 15.3 | 2.0 | 10.0 | 20.0 | 11.7 | 16.0 | 6.7 | 3.3 | 5.7 | 10.3 |
| | | | 4 | 19.7 | 1.0 | 45.3 | 18.3 | 9.3 | 12.0 | 19.7 | 13.0 | 9.7 | 23.0 | 5.3 | 19.0 | 8.3 | 2.0 | 17.3 | 16.7 | 3.7 | 4.3 | 3.7 | 5.0 |
| | | | 5 | 21.7 | 15.7 | 34.3 | 17.0 | 6.3 | 14.7 | 12.7 | 10.7 | 17.3 | 17.0 | 12.3 | 7.7 | 11.3 | 2.7 | 19.0 | 11.3 | 9.0 | 3.0 | 3.3 | 4.7 |
| | | | 6 | 16.3 | 8.3 | 24.3 | 10.0 | 16.7 | 7.3 | 7.3 | 11.7 | 6.3 | 11.0 | 4.7 | 13.7 | 2.7 | 10.7 | 7.0 | 17.3 | 3.0 | 5.5 | 2.0 | 8.0 |
| | | | 7 | 18.3 | 34.3 | 12.3 | 16.0 | 9.3 | 10.7 | 14.7 | 9.0 | 13.7 | 11.0 | 12.0 | 14.7 | 3.0 | 9.7 | 10.3 | 10.7 | 1.7 | 13.7 | 9.0 | 2.7 |
| | | | 8 | 12.3 | 6.0 | 29.0 | 16.3 | 2.7 | 7.7 | 23.0 | 9.3 | 11.7 | 1.7 | 23.7 | 9.3 | 1.0 | 1.0 | 24.0 | 14.0 | 0.7 | 10.3 | 2.0 | 11.0 |
| $w^{1118}$ ♀ x $w^{1118}$ ♂ | 5 | 24-07-10 | 1 | 0.0 | 9.0 | 11.5 | 10.0 | 1.5 | 20.5 | 15.0 | 14.5 | 16.5 | 2.5 | 25.5 | 19.0 | 4.0 | 18.0 | 10.0 | 11.5 | 5.0 | 12.0 | 6.0 | 1.0 |
| | | | 2 | 0.0 | 1.7 | 25.3 | 14.3 | 3.3 | 21.3 | 18.3 | 17.7 | 20.0 | 2.0 | 14.7 | 29.7 | 1.7 | 27.0 | 7.0 | 2.3 | 4.7 | 20.3 | 1.0 | 7.3 |
| | | | 3 | 9.3 | 12.7 | 16.7 | 14.7 | 3.7 | 19.0 | 18.7 | 19.0 | 19.0 | 14.0 | 14.0 | 19.7 | 5.0 | 19.7 | 7.3 | 6.7 | 0.7 | 8.7 | 17.3 | 1.0 |
| | | | 4 | 0.0 | 1.0 | 17.0 | 22.7 | 4.3 | 18.0 | 17.3 | 16.3 | 17.7 | 10.7 | 3.3 | 23.7 | 8.0 | 19.0 | 13.7 | 11.0 | 2.3 | 4.7 | 2.3 | 7.3 |
| | | | 5 | 3.5 | 1.0 | 16.0 | 25.5 | 1.5 | 16.5 | 15.0 | 19.0 | 20.5 | 23.0 | 10.5 | 25.0 | 9.5 | 20.5 | 16.0 | 10.0 | 3.0 | 11.0 | 9.0 | 11.5 |
| | | | 6 | 8.0 | 4.7 | 20.7 | 7.7 | 11.3 | 12.7 | 15.0 | 18.7 | 17.7 | 7.3 | 10.7 | 13.0 | 8.3 | 16.3 | 7.0 | 7.0 | 6.0 | 10.3 | 8.0 | 2.7 |
| | | | 7 | 2.0 | 0.7 | 29.3 | 7.7 | 10.3 | 9.3 | 13.7 | 12.3 | 9.0 | 1.0 | 1.0 | 25.0 | 6.0 | 11.3 | 7.3 | 9.0 | 3.0 | 6.0 | 10.3 | 1.0 |
| | | | 8 | 6.0 | 14.7 | 17.7 | 12.7 | 6.3 | 17.3 | 20.3 | 13.3 | 20.0 | 1.0 | 25.0 | 12.3 | 1.3 | 21.7 | 6.7 | 6.7 | 1.0 | 13.3 | 5.7 | 1.3 |
| *$DJ694^{L}$;UAS-EDTP$^{E}/+$* ♀ x *$w^{1118}$* ♂ | 4 | 24-07-09 | 1 | 4.7 | 1.7 | 16.3 | 1.0 | 0.7 | 29.3 | 24.7 | 16.0 | 0.7 | 24.7 | 2.3 | 2.7 | 17.3 | 28.7 | 1.0 | 17.3 | 10.3 | 1.0 | 5.0 | 32.3 |
| | | | 2 | 7.7 | 0.7 | 15.3 | 1.0 | 2.7 | 12.3 | 18.0 | 47.0 | 1.0 | 12.0 | 4.0 | 30.3 | 19.7 | 1.0 | 4.7 | 41.7 | 7.3 | 13.3 | 1.7 | 6.7 |
| | | | 3 | 21.7 | 14.0 | 22.0 | 13.0 | 7.0 | 16.7 | 28.0 | 37.0 | 10.0 | 32.0 | 4.0 | 34.5 | 1.0 | 2.0 | 20.5 | 46.0 | 22.5 | 8.0 | 0.5 | 24.5 |
| | | | 4 | 23.0 | 5.7 | 14.0 | 1.7 | 25.7 | 6.7 | 14.0 | 4.0 | 11.3 | 13.0 | 15.0 | 12.0 | 16.5 | 0.0 | 15.5 | 10.0 | 2.0 | 13.5 | 0.5 | 14.0 |
| | | | 5 | 15.0 | 1.7 | 20.7 | 0.3 | 7.3 | 12.3 | 15.3 | 12.3 | 9.7 | 37.0 | 1.0 | 3.0 | 4.3 | 4.0 | 10.3 | 5.7 | 0.0 | 10.0 | 2.0 | 1.0 |
| | | | 6 | 21.0 | 1.0 | 21.7 | 5.0 | 1.0 | 33.3 | 10.3 | 17.3 | 1.0 | 19.3 | 6.7 | 4.3 | 37.0 | 2.0 | 18.0 | 33.3 | 3.7 | 1.3 | 21.7 | 1.0 |
| | | | 7 | 11.3 | 5.3 | 4.7 | 14.3 | 0.7 | 18.3 | 32.0 | 7.0 | 5.3 | 18.3 | 4.7 | 34.7 | 20.3 | 1.3 | 21.3 | 40.3 | 23.3 | 1.7 | 6.7 | 21.0 |
| | | | 8 | 17.7 | 0.7 | 25.0 | 4.7 | 19.0 | 0.7 | 13.3 | 8.0 | 25.7 | 13.7 | 12.0 | 4.3 | 7.7 | 0.7 | 47.3 | 10.3 | 5.3 | 1.3 | 16.0 | 6.0 |
| | 5 | 24-07-10 | 1 | 0.0 | 26.0 | 6.0 | 21.3 | 0.7 | 37.7 | 4.7 | 22.0 | 20.0 | 7.3 | 29.0 | 11.7 | 3.7 | 16.0 | 17.0 | 7.3 | 5.7 | 13.7 | 13.3 | 2.0 |
| | | | 2 | 0.0 | 1.0 | 3.3 | 26.3 | 0.3 | 13.3 | 11.7 | 13.7 | 39.7 | 2.3 | 3.7 | 5.0 | 5.7 | 60.7 | 3.3 | 1.7 | 2.3 | 8.0 | 5.3 | 2.0 |
| | | | 3 | 0.0 | 5.3 | 11.3 | 4.7 | 25.7 | 7.7 | 23.3 | 32.3 | 17.7 | 4.0 | 11.7 | 9.7 | 11.0 | 43.3 | 5.0 | 6.3 | 3.3 | 4.3 | 23.7 | 10.7 |
| | | | 4 | 0.0 | 1.3 | 1.3 | 1.3 | 33.0 | 18.7 | 1.3 | 43.7 | 3.7 | 14.0 | 16.0 | 31.7 | 0.7 | 3.7 | 17.7 | 22.3 | 2.0 | 11.0 | 20.7 | 1.3 |
| | | | 5 | 0.0 | 10.7 | 3.7 | 2.7 | 30.0 | 7.7 | 13.3 | 1.0 | 35.0 | 1.0 | 12.0 | 1.0 | 1.0 | 23.3 | 18.3 | 12.0 | 1.3 | 3.7 | 17.7 | 1.3 |
| | | | 6 | 0.0 | 0.7 | 4.0 | 6.0 | 58.0 | 4.0 | 43.0 | 5.0 | 33.3 | 5.7 | 29.0 | 3.3 | 0.7 | 35.7 | 6.7 | 11.7 | 1.7 | 29.3 | 11.0 | 1.7 |
| | | | 7 | 0.0 | 2.7 | 9.3 | 2.7 | 18.0 | 33.7 | 19.7 | 4.0 | 9.3 | 1.0 | 19.7 | 19.3 | 2.7 | 34.7 | 11.0 | 8.7 | 1.3 | 3.7 | 9.3 | 5.0 |
| | | | 8 | 0.3 | 3.0 | 23.3 | 9.7 | 18.3 | 17.7 | 17.0 | 17.7 | 6.0 | 23.7 | 11.0 | 18.7 | 6.0 | 30.7 | 19.0 | 18.0 | 6.7 | 20.7 | 10.7 | 14.0 |

Table S5. Female fertility data (eggs laid per female per day) for the indicated genetic crosses 

| Cross | Rep | Start Date (YYMMDD) | Vial | Days 21 | 22 | 23 | 24 | 25 | 26 | 27 | 28 | 29 | 30 | 31 | 32 | 33 | 34 | 35 | 36 | 37 | 38 | 39 | 40 |
|---|---|---|---|---|---|---|---|---|---|---|---|---|---|---|---|---|---|---|---|---|---|---|---|
| *DJ694$^{P}$/+* ♀ x *w$^{1118}$* ♂ | 4 | 24-07-09 | 1 | 10.0 | 1.3 | 10.3 | 15.0 | 14.7 | 6.7 | 9.7 | 7.7 | 11.3 | 11.7 | 8.3 | 4.7 | 4.7 | 12.0 | 4.7 | 6.3 | 12.3 | 7.3 | 3.7 | 14.3 |
| | | | 2 | 7.3 | 3.0 | 28.7 | 8.0 | 6.7 | 9.3 | 4.3 | 5.7 | 5.0 | 0.7 | 15.0 | 7.0 | 5.3 | 1.0 | 5.7 | 11.0 | 8.3 | 25.3 | 12.3 | 6.3 |
| | | | 3 | 1.0 | 1.0 | 1.0 | 35.0 | 13.3 | 13.0 | 14.0 | 4.7 | 1.3 | 1.0 | 21.0 | 6.0 | 0.7 | 1.0 | 2.7 | 19.7 | 12.0 | 5.7 | 15.3 | 5.3 |
| | 5 | 24-07-10 | 1 | 12.7 | 9.7 | 6.7 | 9.3 | 9.0 | 14.0 | 1.7 | 5.0 | 1.3 | 10.7 | 3.0 | 26.7 | 1.0 | 5.3 | 2.0 | 13.3 | 2.7 | 18.3 | 9.7 | 1.7 |
| | | | 2 | 1.3 | 1.0 | 14.3 | 16.0 | 15.7 | 18.0 | 2.3 | 3.0 | 15.7 | 6.0 | 13.3 | 13.0 | 3.7 | 2.3 | 15.0 | 4.0 | 6.7 | 3.0 | 12.5 | 1.0 |
| | | | 3 | 23.3 | 0.7 | 5.3 | 10.0 | 8.7 | 13.0 | 10.7 | 8.3 | 4.3 | 1.3 | 33.7 | 3.0 | 10.7 | 3.7 | 6.3 | 11.3 | 9.3 | 15.3 | 13.0 | 4.7 |
| | | | 4 | 1.7 | 0.7 | 28.0 | 13.3 | 10.0 | 9.0 | 8.3 | 2.0 | 1.0 | 5.7 | 3.3 | 2.7 | 38.0 | 0.7 | 2.0 | 14.7 | 1.3 | 16.0 | 5.3 | 1.0 |
| | | | 5 | 1.3 | 0.7 | 1.3 | 18.0 | 16.7 | 10.0 | 1.3 | 1.0 | 7.3 | 8.7 | 1.0 | 29.3 | 1.7 | 1.3 | 15.3 | 4.3 | 3.3 | 19.3 | 5.3 | 11.0 |
| | | | 6 | 1.0 | 0.7 | 18.3 | 5.3 | 5.3 | 13.7 | 10.0 | 1.0 | 1.0 | 22.0 | 6.7 | 14.3 | 0.7 | 1.0 | 12.7 | 0.3 | 23.7 | 12.3 | 12.7 | 8.7 |
| | | | 7 | 1.7 | 1.3 | 17.0 | 23.7 | 9.7 | 12.3 | 10.0 | 4.3 | 19.0 | 1.0 | 2.0 | 1.0 | 5.3 | 34.0 | 5.7 | 10.7 | 3.7 | 24.0 | 3.0 | 8.3 |
| *w$^{1118}$* ♀ x *w$^{1118}$* ♂ | 4 | 24-07-09 | 1 | 3.0 | 26.0 | 5.5 | 19.5 | 12.5 | 10.0 | 7.5 | 10.5 | 1.0 | 12.5 | 6.0 | 15.0 | 2.5 | 9.5 | 0.5 | 13.0 | 5.0 | 12.5 | 14.5 | 11.0 |
| | | | 2 | 1.0 | 20.5 | 5.0 | 15.0 | 14.5 | 10.5 | 7.0 | 3.5 | 11.0 | 9.5 | 7.5 | 6.0 | 1.0 | 1.5 | 7.5 | 7.0 | 0.5 | 10.0 | 2.5 | 1.0 |
| | | | 3 | 2.7 | 15.3 | 9.3 | 8.0 | 12.0 | 11.7 | 3.3 | 0.7 | 8.7 | 9.3 | 5.7 | 5.3 | 9.0 | 2.0 | 2.3 | 4.7 | 9.7 | 9.7 | 12.3 | 3.3 |
| | | | 4 | 8.0 | 4.3 | 5.7 | 6.7 | 8.0 | 3.3 | 5.3 | 6.0 | 5.7 | 7.7 | 1.0 | 0.7 | 4.7 | 2.0 | 5.0 | 8.0 | 0.0 | 1.0 | 6.7 | 1.0 |
| | | | 5 | 6.0 | 17.7 | 1.3 | 4.7 | 13.0 | 12.7 | 7.7 | 3.3 | 4.0 | 7.7 | 5.7 | 5.0 | 5.0 | 10.0 | 6.3 | 1.0 | 9.0 | 5.0 | 11.7 | 6.3 |
| | | | 6 | 4.5 | 6.0 | 11.0 | 5.0 | 10.5 | 5.0 | 12.0 | 2.5 | 0.0 | 5.5 | 6.5 | 4.0 | 0.0 | 1.5 | 3.0 | 4.0 | 3.5 | 3.5 | 4.0 | 1.0 |
| | | | 7 | 10.7 | 6.3 | 11.0 | 6.7 | 8.0 | 8.0 | 8.3 | 3.3 | 1.7 | 7.3 | 4.0 | 13.0 | 4.7 | 3.0 | 7.3 | 9.7 | 4.3 | 18.0 | 1.0 | 4.0 |
| | | | 8 | 0.7 | 1.0 | 8.0 | 17.0 | 8.7 | 2.7 | 3.3 | 1.7 | 3.7 | 9.7 | 2.3 | 11.7 | 1.7 | 1.3 | 1.0 | 12.7 | 3.0 | 13.3 | 1.7 | 4.0 |
| w$^{1118}$ ♀ x w$^{1118}$ ♂ | 5 | 24-07-10 | 1 | 17.5 | 5.5 | 13.5 | 7.0 | 7.0 | 7.5 | 5.0 | 1.0 | 2.0 | 27.0 | 1.0 | 21.0 | 4.0 | 1.0 | 16.0 | 14.0 | 3.0 | 1.0 | 1.0 | 5.0 |
| | | | 2 | 10.7 | 9.0 | 10.0 | 8.0 | 7.7 | 14.7 | 6.7 | 1.3 | 8.7 | 12.7 | 8.3 | 13.0 | 1.0 | 3.0 | 9.0 | 5.7 | 9.7 | 6.7 | 1.3 | 3.3 |
| | | | 3 | 0.7 | 10.7 | 5.3 | 9.0 | 3.7 | 4.0 | 8.0 | 8.0 | 0.7 | 13.0 | 0.7 | 2.7 | 12.3 | 0.3 | 10.3 | 5.0 | 3.5 | 0.5 | 1.0 | 9.5 |
| | | | 4 | 16.7 | 12.0 | 6.3 | 6.7 | 6.3 | 13.3 | 7.3 | 3.7 | 7.3 | 4.7 | 4.0 | 19.3 | 1.3 | 14.3 | 6.7 | 7.3 | 7.3 | 12.3 | 3.7 | 4.3 |
| | | | 5 | 16.0 | 9.0 | 12.0 | 7.5 | 7.5 | 8.5 | 10.0 | 3.5 | 8.0 | 14.5 | 2.5 | 8.0 | 11.0 | 1.0 | 9.5 | 15.0 | 3.5 | 14.5 | 6.0 | 3.5 |
| | | | 6 | 12.3 | 1.3 | 14.0 | 10.0 | 1.3 | 9.7 | 5.7 | 3.0 | 8.7 | 8.0 | 2.0 | 8.7 | 2.3 | 0.3 | 8.7 | 9.0 | 2.3 | 4.0 | 3.7 | 2.3 |
| | | | 7 | 6.0 | 13.7 | 6.7 | 9.0 | 9.3 | 7.3 | 2.0 | 1.3 | 0.3 | 16.7 | 0.7 | 10.7 | 1.0 | 4.7 | 8.3 | 5.3 | 5.3 | 4.0 | 2.0 | 1.0 |
| | | | 8 | 13.7 | 5.3 | 6.3 | 5.7 | 5.7 | 5.0 | 1.3 | 1.3 | 8.0 | 5.3 | 4.7 | 2.7 | 6.3 | 2.7 | 10.0 | 3.3 | 1.7 | 1.0 | 2.0 | 2.7 |
| *DJ694$^{L}$;UAS-EDTP$^{E}$/+* ♀ x *w$^{1118}$* ♂ | 4 | 24-07-09 | 1 | 0.7 | 0.3 | 1.3 | 9.7 | 7.7 | 4.7 | 18.7 | 11.7 | 4.0 | 1.0 | 3.0 | 1.7 | 10.7 | 1.0 | 4.7 | 13.3 | 6.7 | 4.7 | 4.3 | 1.3 |
| | | | 2 | 1.7 | 6.0 | 11.3 | 11.0 | 8.0 | 8.0 | 9.0 | 0.3 | 6.0 | 4.3 | 1.7 | 21.7 | 2.0 | 5.0 | 2.7 | 6.0 | 6.7 | 9.3 | 1.0 | 5.3 |
| | | | 3 | 1.0 | 7.5 | 29.5 | 4.0 | 18.0 | 5.0 | 21.5 | 6.0 | 4.5 | 10.0 | 12.5 | 18.0 | 7.0 | 1.5 | 7.0 | 5.5 | 10.5 | 7.0 | 10.0 | 2.0 |
| | | | 4 | 3.5 | 7.0 | 2.0 | 11.0 | 5.5 | 6.0 | 11.5 | 1.0 | 1.0 | 1.0 | 11.5 | 5.5 | 6.0 | 1.0 | 7.5 | 3.0 | 0.5 | 0.0 | 0.0 | 2.5 |
| | | | 5 | 0.3 | 3.3 | 5.7 | 5.3 | 6.3 | 6.3 | 3.3 | 2.0 | 4.3 | 2.7 | 4.3 | 4.3 | 1.3 | 7.0 | 1.0 | 2.7 | 1.7 | 0.7 | 0.0 | 0.0 |
| | | | 6 | 1.7 | 16.3 | 1.5 | 19.0 | 20.5 | 10.0 | 13.0 | 10.0 | 2.0 | 13.5 | 14.0 | 6.0 | 6.0 | 4.0 | 6.0 | 10.5 | 8.0 | 4.5 | 4.5 | 12.0 |
| | | | 7 | 1.0 | 2.0 | 8.3 | 31.7 | 10.3 | 10.7 | 6.7 | 4.3 | 8.7 | 5.0 | 18.0 | 12.0 | 17.3 | 1.7 | 3.0 | 3.0 | 12.7 | 14.3 | 2.3 | 4.7 |
| | | | 8 | 9.0 | 7.3 | 5.7 | 8.3 | 13.0 | 1.3 | 7.7 | 6.3 | 1.3 | 1.3 | 1.0 | 27.7 | 5.3 | 1.7 | 8.0 | 3.0 | 11.3 | 6.7 | 0.7 | 5.0 |
| | 5 | 24-07-10 | 1 | 8.0 | 16.7 | 4.7 | 7.3 | 7.3 | 12.0 | 3.3 | 5.7 | 4.3 | 0.7 | 8.3 | 11.3 | 6.7 | 2.3 | 2.7 | 6.7 | 6.7 | 3.7 | 4.0 | 3.0 |
| | | | 2 | 13.0 | 2.7 | 6.3 | 22.7 | 22.7 | 7.0 | 4.0 | 5.3 | 2.0 | 2.3 | 2.3 | 7.7 | 6.0 | 4.0 | 8.7 | 6.0 | 7.7 | 4.0 | 10.3 | 0.3 |
| | | | 3 | 10.3 | 8.0 | 2.7 | 6.7 | 9.3 | 8.3 | 8.3 | 9.0 | 6.0 | 7.7 | 6.7 | 10.0 | 12.7 | 6.3 | 8.3 | 7.7 | 6.7 | 10.3 | 9.7 | 6.7 |
| | | | 4 | 15.3 | 20.3 | 4.3 | 8.7 | 5.7 | 14.3 | 1.7 | 2.0 | 4.3 | 7.3 | 4.3 | 6.0 | 3.7 | 1.0 | 1.3 | 3.7 | 3.7 | 7.3 | 6.0 | 2.3 |
| | | | 5 | 15.0 | 2.0 | 4.3 | 10.7 | 5.3 | 5.0 | 8.3 | 20.3 | 1.3 | 1.7 | 22.0 | 11.0 | 2.3 | 1.0 | 7.3 | 8.0 | 1.3 | 8.3 | 16.7 | 2.3 |
| | | | 6 | 7.0 | 8.0 | 1.3 | 15.3 | 22.5 | 6.0 | 5.5 | 13.5 | 1.5 | 1.0 | 12.5 | 12.0 | 8.5 | 3.0 | 14.0 | 7.0 | 1.0 | 12.0 | 12.0 | 2.5 |
| | | | 7 | 13.7 | 16.3 | 5.0 | 6.0 | 6.0 | 5.7 | 2.7 | 2.7 | 5.3 | 8.0 | 2.3 | 10.7 | 6.7 | 2.3 | 5.0 | 5.0 | 6.0 | 7.0 | 4.3 | 2.7 |
| | | | 8 | 13.7 | 8.3 | 2.7 | 6.7 | 6.0 | 18.7 | 11.7 | 1.3 | 2.7 | 18.7 | 11.7 | 2.7 | 7.3 | 3.3 | 3.7 | 8.3 | 4.7 | 5.7 | 6.7 | 1.3 |

| Cross | Rep | Start Date (YYMMDD) | Vial | Days 1 | 2 | 3 | 4 | 5 | 6 | 7 | 8 | 9 | 10 | 11 | 12 | 13 | 14 | 15 | 16 | 17 | 18 | 19 | 20 |
|---|---|---|---|---|---|---|---|---|---|---|---|---|---|---|---|---|---|---|---|---|---|---|---|
| *DJ694*$^{L}$/+;*UAS-EDTP*$^{E}$/+ ♀ x *w*$^{1118}$ ♂ | 6 | 24-09-28 | 1 | 7.7 | 3.0 | 10.0 | 2.0 | 19.3 | 19.0 | 6.3 | 15.3 | 4.3 | 16.0 | 13.0 | 7.7 | 3.3 | 10.0 | 1.0 | 9.7 | 10.3 | 3.3 | 5.3 | 9.3 |
| | | | 2 | 5.0 | 5.0 | 10.3 | 1.0 | 22.3 | 13.0 | 10.3 | 14.0 | 9.3 | 3.0 | 5.7 | 4.7 | 8.7 | 16.7 | 13.3 | 4.0 | 6.7 | 2.0 | 1.3 | 14.7 |
| | | | 3 | 7.3 | 3.3 | 25.7 | 18.3 | 4.3 | 19.3 | 8.3 | 16.7 | 5.7 | 8.0 | 4.7 | 10.0 | 7.0 | 14.7 | 7.7 | 8.3 | 8.3 | 1.3 | 5.0 | 15.0 |
| | | | 4 | 12.7 | 10.3 | 4.3 | 4.0 | 24.0 | 11.3 | 7.7 | 18.3 | 3.0 | 7.0 | 10.3 | 2.0 | 7.0 | 7.3 | 7.3 | 0.7 | 3.3 | 1.3 | 10.0 | 12.0 |
| | | | 5 | 11.0 | 9.7 | 15.3 | 6.7 | 21.3 | 8.0 | 20.3 | 7.3 | 9.3 | 9.0 | 1.0 | 13.3 | 13.3 | 0.3 | 5.3 | 6.7 | 9.3 | 3.0 | 3.0 | 16.3 |
| | | | 6 | 5.0 | 3.3 | 1.7 | 2.3 | 14.0 | 13.7 | 16.3 | 14.3 | 1.7 | 14.7 | 1.0 | 15.0 | 10.0 | 16.3 | 11.7 | 6.3 | 1.0 | 5.0 | 5.3 | 12.7 |
| | | | 7 | 5.0 | 1.3 | 0.3 | 8.7 | 16.7 | 7.3 | 11.3 | 16.7 | 1.0 | 14.0 | 1.0 | 23.0 | 19.7 | 12.7 | 9.7 | 8.3 | 6.0 | 1.7 | 1.0 | 19.7 |
| | | | 8 | 3.7 | 3.7 | 9.7 | 4.0 | 10.3 | 11.7 | 10.0 | 1.7 | 5.7 | 22.7 | 1.0 | 6.3 | 3.3 | 19.3 | 4.0 | 10.0 | 1.3 | 1.7 | 4.7 | 14.3 |
| | 7 | 24-09-28 | 1 | 9.7 | 8.7 | 26.7 | 18.3 | 18.3 | 6.0 | 15.7 | 16.7 | 8.0 | 8.7 | 7.3 | 10.0 | 7.3 | 10.0 | 1.7 | 7.0 | 1.0 | 8.3 | 2.3 | 10.0 |
| | | | 2 | 3.3 | 1.0 | 1.0 | 11.3 | 18.3 | 5.7 | 4.7 | 17.0 | 4.0 | 13.3 | 9.0 | 11.0 | 13.3 | 9.7 | 3.3 | 4.3 | 7.7 | 12.3 | 0.7 | 19.3 |
| | | | 3 | 5.3 | 4.0 | 6.3 | 2.0 | 18.7 | 1.0 | 7.3 | 13.0 | 3.7 | 13.7 | 1.7 | 14.3 | 3.3 | 4.7 | 4.3 | 3.0 | 0.7 | 6.0 | 5.7 | 13.0 |
| | | | 4 | 16.3 | 1.0 | 5.7 | 26.7 | 6.0 | 10.3 | 12.3 | 11.3 | 10.7 | 13.0 | 8.3 | 6.3 | 2.0 | 12.0 | 4.7 | 0.3 | 3.0 | 1.7 | 11.7 | 16.3 |
| | | | 5 | 4.7 | 1.3 | 9.0 | 7.0 | 13.3 | 2.3 | 8.0 | 23.7 | 14.3 | 3.7 | 2.7 | 7.3 | 8.7 | 6.3 | 0.3 | 0.3 | 1.0 | 2.0 | 9.0 | 19.7 |
| | | | 6 | 7.3 | 6.0 | 0.7 | 1.7 | 14.0 | 1.7 | 3.0 | 4.0 | 8.3 | 12.7 | 5.3 | 13.3 | 5.0 | 4.0 | 2.3 | 10.0 | 1.7 | 9.7 | 1.3 | 22.0 |
| | | | 7 | 11.0 | 5.0 | 1.0 | 11.0 | 4.7 | 16.0 | 5.0 | 4.7 | 8.3 | 5.0 | 1.0 | 14.7 | 12.3 | 4.3 | 7.7 | 4.0 | 2.3 | 6.0 | 7.7 | 18.3 |
| | | | 8 | 11.7 | 0.7 | 1.3 | 3.3 | 10.7 | 13.7 | 19.7 | 10.0 | 2.3 | 21.0 | 0.7 | 12.7 | 0.3 | 10.0 | 1.0 | 9.7 | 7.3 | 4.3 | 13.7 | 9.7 |
| *UAS-EDTP*$^{E}$/+ ♀ x *w*$^{1118}$ ♂ | 6 | 24-09-28 | 1 | 11.7 | 1.7 | 25.7 | 22.7 | 23.7 | 10.3 | 21.7 | 13.0 | 9.7 | 11.0 | 6.3 | 8.7 | 5.7 | 5.7 | 7.3 | 8.0 | 7.7 | 3.7 | 10.3 | 4.0 |
| | | | 2 | 7.3 | 0.7 | 50.3 | 24.7 | 21.0 | 12.3 | 19.0 | 7.7 | 8.0 | 11.0 | 4.0 | 8.0 | 6.3 | 4.0 | 5.0 | 9.3 | 5.0 | 2.0 | 1.0 | 11.0 |
| | | | 3 | 17.7 | 11.7 | 15.7 | 17.3 | 11.7 | 16.0 | 20.0 | 12.7 | 9.7 | 12.3 | 4.0 | 11.0 | 9.7 | 5.0 | 5.3 | 4.3 | 4.7 | 1.3 | 14.0 | 2.0 |
| | | | 4 | 7.3 | 10.3 | 30.0 | 22.7 | 17.0 | 14.7 | 10.3 | 11.7 | 3.7 | 16.3 | 4.3 | 15.0 | 2.7 | 13.3 | 1.3 | 6.7 | 1.0 | 4.7 | 10.0 | 3.7 |
| | | | 5 | 4.7 | 1.7 | 39.0 | 21.3 | 14.0 | 11.3 | 15.3 | 13.7 | 4.0 | 11.3 | 5.0 | 6.7 | 6.0 | 4.3 | 7.7 | 4.7 | 4.7 | 4.3 | 11.3 | 10.0 |
| | | | 6 | 22.7 | 14.3 | 7.3 | 19.3 | 13.3 | 8.0 | 16.0 | 10.3 | 5.7 | 13.0 | 4.7 | 13.0 | 9.3 | 6.7 | 5.7 | 3.3 | 1.7 | 2.0 | 5.0 | 8.7 |
| | | | 7 | 15.0 | 6.7 | 17.3 | 17.0 | 14.0 | 14.7 | 14.3 | 11.3 | 9.0 | 12.7 | 1.3 | 4.0 | 8.0 | 1.0 | 10.0 | 2.7 | 0.0 | 2.0 | 12.7 | 5.3 |
| | | | 8 | 18.0 | 9.7 | 21.3 | 14.3 | 10.3 | 10.3 | 23.0 | 5.3 | 10.3 | 6.3 | 11.0 | 5.3 | 0.7 | 22.3 | 0.7 | 0.7 | 1.0 | 9.0 | 6.7 | 19.7 |
| | 7 | 24-09-28 | 1 | 15.0 | 7.3 | 4.7 | 28.7 | 22.0 | 10.7 | 17.7 | 14.0 | 5.7 | 10.0 | 3.3 | 6.7 | 9.0 | 4.7 | 6.0 | 2.3 | 5.0 | 7.7 | 1.0 | 8.3 |
| | | | 2 | 24.7 | 6.3 | 13.0 | 33.7 | 22.7 | 11.0 | 14.3 | 9.3 | 11.7 | 5.0 | 5.0 | 11.7 | 4.7 | 2.7 | 9.3 | 3.0 | 7.0 | 8.7 | 3.0 | 13.3 |
| | | | 3 | 21.0 | 12.7 | 13.3 | 19.7 | 22.0 | 12.0 | 5.7 | 6.0 | 11.7 | 11.7 | 12.7 | 3.7 | 6.7 | 7.7 | 6.7 | 3.7 | 9.3 | 1.0 | 3.0 | 11.7 |
| | | | 4 | 6.7 | 2.3 | 17.0 | 11.3 | 14.0 | 11.3 | 6.7 | 11.3 | 10.3 | 13.3 | 1.0 | 9.7 | 6.7 | 2.7 | 0.3 | 7.3 | 5.3 | 9.7 | 2.0 | 11.3 |
| | | | 5 | 22.0 | 4.0 | 8.0 | 18.7 | 14.3 | 10.0 | 14.3 | 14.3 | 5.0 | 17.3 | 4.3 | 8.7 | 5.3 | 10.0 | 1.0 | 8.0 | 0.3 | 11.3 | 1.3 | 15.0 |
| | | | 6 | 7.7 | 11.3 | 6.3 | 19.3 | 12.7 | 4.7 | 18.7 | 14.7 | 9.0 | 10.0 | 4.0 | 5.0 | 6.0 | 6.7 | 3.7 | 4.7 | 2.3 | 5.3 | 2.7 | 11.0 |
| | | | 7 | 6.3 | 9.3 | 15.0 | 24.3 | 22.0 | 11.3 | 13.0 | 13.3 | 8.3 | 9.0 | 4.7 | 10.0 | 3.0 | 4.7 | 0.3 | 7.0 | 2.0 | 7.7 | 5.0 | 13.0 |
| | | | 8 | 10.0 | 3.3 | 18.3 | 17.0 | 19.0 | 19.7 | 11.7 | 13.7 | 7.0 | 9.0 | 3.7 | 6.7 | 7.0 | 13.0 | 5.7 | 6.3 | 8.0 | 6.7 | 8.3 | 3.7 |

| | | Start Date | | Days | | | | | | | | | | | | | | | | | | | |
|---|---|---|---|---|---|---|---|---|---|---|---|---|---|---|---|---|---|---|---|---|---|---|---|
| Cross | Rep | (YYMMDD) | Vial | 21 | 22 | 23 | 24 | 25 | 26 | 27 | 28 | 29 | 30 | 31 | 32 | 33 | 34 | 35 | 36 | 37 | 38 | 39 | 40 |
| *DJ694*$^{L}$/+;*UAS-EDTP*$^{E}$/+ ♀ x $w^{1118}$ ♂ | 6 | 24-09-28 | 1 | 4.0 | 3.0 | 9.7 | 1.7 | 6.0 | 6.3 | 1.3 | 3.3 | 5.7 | 6.7 | 4.3 | 3.7 | 3.7 | 6.7 | 1.3 | 3.3 | 2.3 | 7.0 | 4.0 | 4.0 |
| | | | 2 | 6.7 | 12.3 | 7.0 | 1.7 | 3.0 | 1.3 | 1.0 | 7.3 | 6.3 | 3.0 | 5.0 | 15.0 | 4.7 | 10.7 | 5.0 | 4.7 | 2.3 | 1.7 | 10.0 | 2.7 |
| | | | 3 | 7.0 | 6.0 | 6.0 | 2.7 | 6.0 | 3.0 | 1.0 | 4.3 | 9.7 | 1.7 | 9.7 | 3.3 | 2.3 | 6.7 | 7.0 | 5.0 | 6.7 | 2.7 | 5.0 | 8.7 |
| | | | 4 | 9.0 | 2.3 | 6.7 | 1.7 | 2.3 | 3.3 | 2.7 | 2.7 | 0.3 | 0.3 | 0.7 | 0.7 | 4.7 | 3.3 | 4.3 | 9.0 | 4.7 | 1.0 | 2.0 | 2.0 |
| | | | 5 | 12.3 | 10.0 | 8.3 | 0.7 | 1.7 | 5.0 | 1.3 | 2.0 | 1.0 | 2.3 | 3.3 | 8.3 | 9.3 | 15.7 | 11.0 | 4.3 | 4.0 | 3.7 | 8.3 | 8.7 |
| | | | 6 | 3.3 | 20.3 | 6.3 | 1.3 | 1.3 | 4.3 | 2.0 | 4.3 | 1.7 | 1.7 | 9.3 | 9.7 | 3.7 | 6.0 | 3.7 | 3.7 | 4.0 | 10.3 | 4.0 | 4.7 |
| | | | 7 | 8.7 | 3.3 | 0.7 | 1.0 | 1.7 | 8.7 | 2.3 | 6.7 | 1.3 | 1.0 | 14.3 | 16.0 | 2.7 | 5.0 | 8.7 | 3.0 | 3.3 | 1.7 | 4.3 | 2.0 |
| | | | 8 | 1.0 | 8.3 | 9.3 | 1.7 | 1.0 | 4.3 | 4.3 | 1.7 | 0.7 | 1.7 | 6.3 | 10.7 | 5.7 | 5.7 | 3.7 | 10.7 | 4.7 | 2.0 | 9.7 | 5.3 |
| | 7 | 24-09-28 | 1 | 2.7 | 5.0 | 5.3 | 2.0 | 5.0 | 0.7 | 3.3 | 2.3 | 1.7 | 4.3 | 6.7 | 3.0 | 1.0 | 3.3 | 4.0 | 3.3 | 3.0 | 1.0 | 3.7 | 3.0 |
| | | | 2 | 8.0 | 9.3 | 8.7 | 0.7 | 10.7 | 4.0 | 1.3 | 14.7 | 3.3 | 1.0 | 2.3 | 8.3 | 8.7 | 8.7 | 8.7 | 11.0 | 8.7 | 4.0 | 15.0 | 7.0 |
| | | | 3 | 12.3 | 6.3 | 9.3 | 2.3 | 4.0 | 2.3 | 2.3 | 8.7 | 6.0 | 0.7 | 1.7 | 12.3 | 8.0 | 3.7 | 2.7 | 10.0 | 2.0 | 3.0 | 8.0 | 7.0 |
| | | | 4 | 0.7 | 3.3 | 3.0 | 0.7 | 3.3 | 1.7 | 2.7 | 8.3 | 5.0 | 2.0 | 4.7 | 8.7 | 5.3 | 5.7 | 7.7 | 3.7 | 6.0 | 5.3 | 7.3 | 5.7 |
| | | | 5 | 3.3 | 12.7 | 8.0 | 1.3 | 4.0 | 1.3 | 3.7 | 6.7 | 2.0 | 4.3 | 18.7 | 3.3 | 4.0 | 6.7 | 2.0 | 4.0 | 6.3 | 4.0 | 10.0 | 1.3 |
| | | | 6 | 19.0 | 7.7 | 9.0 | 2.7 | 3.0 | 4.3 | 4.7 | 6.0 | 7.0 | 1.3 | 7.3 | 13.7 | 8.3 | 5.0 | 2.7 | 5.7 | 4.0 | 5.7 | 5.3 | 5.0 |
| | | | 7 | 2.0 | 13.0 | 4.7 | 1.0 | 5.0 | 2.0 | 4.3 | 5.0 | 4.7 | 2.7 | 2.3 | 9.7 | 12.7 | 6.3 | 5.0 | 6.0 | 3.3 | 6.0 | 10.7 | 4.0 |
| | | | 8 | 3.3 | 6.0 | 3.0 | 5.0 | 1.7 | 2.3 | 2.3 | 7.0 | 2.7 | 1.3 | 2.7 | 6.7 | 1.7 | 3.7 | 1.3 | 0.3 | 1.3 | 0.0 | 2.5 | 1.7 |
| *UAS-EDTP*$^{E}$/+ ♀ x $w^{1118}$ ♂ | 6 | 24-09-28 | 1 | 8.3 | 6.7 | 3.0 | 8.0 | 5.7 | 1.0 | 6.7 | 7.7 | 10.0 | 8.0 | 4.0 | 13.0 | 5.3 | 0.7 | 1.7 | 10.3 | 1.3 | 5.3 | 3.7 | 6.7 |
| | | | 2 | 3.3 | 11.7 | 1.7 | 4.3 | 6.3 | 5.0 | 1.3 | 4.7 | 5.0 | 4.7 | 2.7 | 5.0 | 6.0 | 5.0 | 5.7 | 5.0 | 5.0 | 9.3 | 6.3 | 3.3 |
| | | | 3 | 6.3 | 5.0 | 5.7 | 6.7 | 1.3 | 9.0 | 3.0 | 9.7 | 2.7 | 9.7 | 3.7 | 8.0 | 5.3 | 8.3 | 5.7 | 9.3 | 8.3 | 4.7 | 3.3 | 5.0 |
| | | | 4 | 9.0 | 10.7 | 6.0 | 1.3 | 5.7 | 10.3 | 1.3 | 7.0 | 3.3 | 11.3 | 1.3 | 3.3 | 8.3 | 15.0 | 6.3 | 4.3 | 8.7 | 3.7 | 9.3 | 4.7 |
| | | | 5 | 1.0 | 7.3 | 6.0 | 2.0 | 7.7 | 3.3 | 0.7 | 3.3 | 6.0 | 11.0 | 4.3 | 6.7 | 2.7 | 6.3 | 8.3 | 13.7 | 3.7 | 5.0 | 9.3 | 4.0 |
| | | | 6 | 3.3 | 10.0 | 4.7 | 0.7 | 2.0 | 9.7 | 2.3 | 2.3 | 1.7 | 1.0 | 10.7 | 4.0 | 4.3 | 7.0 | 7.5 | 1.0 | 2.0 | 1.7 | 5.0 | 0.7 |
| | | | 7 | 5.7 | 8.3 | 6.0 | 2.0 | 1.0 | 3.0 | 1.7 | 3.3 | 5.0 | 2.3 | 10.7 | 10.3 | 3.0 | 5.0 | 6.0 | 7.3 | 1.3 | 4.7 | 5.0 | 5.7 |
| | | | 8 | 6.7 | 5.7 | 4.3 | 4.3 | 4.0 | 0.3 | 2.0 | 5.3 | 4.7 | 2.3 | 4.3 | 4.7 | 4.0 | 3.3 | 7.7 | 0.7 | 2.0 | 2.0 | 10.0 | 2.0 |
| | 7 | 24-09-28 | 1 | 6.0 | 6.7 | 0.3 | 4.7 | 3.3 | 1.0 | 4.0 | 3.0 | 4.3 | 10.0 | 6.7 | 4.0 | 4.0 | 6.0 | 4.7 | 4.0 | 2.3 | 9.3 | 2.3 | 2.0 |
| | | | 2 | 2.7 | 6.3 | 0.7 | 5.7 | 7.3 | 2.0 | 4.0 | 8.7 | 1.0 | 7.0 | 4.3 | 5.3 | 3.7 | 3.3 | 5.0 | 3.7 | 1.7 | 7.7 | 4.0 | 8.7 |
| | | | 3 | 9.7 | 3.3 | 2.3 | 5.0 | 4.7 | 4.7 | 3.0 | 10.7 | 6.0 | 1.0 | 6.3 | 8.3 | 2.7 | 7.0 | 5.3 | 9.3 | 3.3 | 11.7 | 4.0 | 4.3 |
| | | | 4 | 7.0 | 8.7 | 3.3 | 2.7 | 11.7 | 8.3 | 4.7 | 4.7 | 9.7 | 4.3 | 6.7 | 3.0 | 8.3 | 4.7 | 4.7 | 6.7 | 1.3 | 5.3 | 9.0 | 5.7 |
| | | | 5 | 1.7 | 3.0 | 1.0 | 8.0 | 2.0 | 6.0 | 3.7 | 9.0 | 9.0 | 6.0 | 2.7 | 5.3 | 6.7 | 4.0 | 10.3 | 4.7 | 6.3 | 8.0 | 5.0 | 1.3 |
| | | | 6 | 2.0 | 8.7 | 1.0 | 3.3 | 7.3 | 2.3 | 3.0 | 11.3 | 6.3 | 8.0 | 5.0 | 6.0 | 4.7 | 5.7 | 5.3 | 6.7 | 7.0 | 2.7 | 7.0 | 3.3 |
| | | | 7 | 1.0 | 6.3 | 3.3 | 5.0 | 5.3 | 1.7 | 4.3 | 0.7 | 2.7 | 2.0 | 6.0 | 6.7 | 1.3 | 3.7 | 4.0 | 8.0 | 1.3 | 2.3 | 7.0 | 4.7 |
| | | | 8 | 1.3 | 6.0 | 4.3 | 3.3 | 6.3 | 3.7 | 2.3 | 8.3 | 5.7 | 11.0 | 1.7 | 10.3 | 1.3 | 7.3 | 9.0 | 8.3 | 8.0 | 9.0 | 11.3 | 5.3 |

Fertility was quantified as the number of eggs laid per female per day for the indicated genetic crosses in 25˚C. For each cross, multiple biological replicates were scored, with each replicate consisting of 3–8 vials (5 or 3 females per vial depends on the replicate) that were monitored daily over the assay period. Values are reported for each day following the start date.
The "Start Date (YYMMDD)" column indicates the date on which fertility test began for each replicate. An "x" indicates that all females within the vial had died, and "–" indicates that data were not applicable.

Table S6. Number of flies that emerged from eggs to adults for the indicated genotypes per each day 

| Cross | Rep | Start Date (YYMMDD) | Vial | Total Egg | Days 1 | 2 | 3 | 4 | 5 | 6 | 7 | 8 | 9 | 10 | 11 | 12 | 13 | 14 | 15 | 16 | 17 | 18 | 19 | 20 | 21 | 22 | 23 | 24 | 25 | 26 | 27 | 28 |
|---|---|---|---|---|---|---|---|---|---|---|---|---|---|---|---|---|---|---|---|---|---|---|---|---|---|---|---|---|---|---|---|---|
| *DJ694* ♀ x *DJ694* ♂ | 1 | 04-03-16 | 1 | 855 | 6 | 28 | 23 | 52 | 29 | 47 | 30 | 64 | 43 | 2 | 9 | 51 | 5 | 17 | 21 | 31 | 6 | 25 | 3 | 6 | 13 | 8 | 1 | 1 | 1 | 1 | 0 | 0 |
| | | | 2 | 1074 | 5 | 57 | 16 | 81 | 20 | 62 | 5 | 40 | 3 | 27 | 24 | 37 | 43 | 8 | 56 | 8 | 3 | 49 | 4 | 4 | 3 | 18 | 1 | 13 | 19 | 2 | 2 | 6 |
| | | | 3 | 454 | 35 | 48 | 37 | 27 | 32 | 23 | 4 | 23 | 5 | 11 | 7 | 7 | 11 | 7 | 8 | 3 | 1 | 9 | 1 | 0 | - | 2 | - | - | 0 | 0 | 0 | 0 |
| | 2 | 05-02-20 | 1 | 1159 | 24 | 29 | 44 | 21 | 29 | 44 | 34 | 9 | 26 | 21 | 33 | 32 | 26 | 49 | 21 | 55 | 21 | 17 | 11 | 15 | 27 | 12 | 17 | 14 | 18 | 8 | 9 | 15 |
| | | | 2 | 1285 | 24 | 43 | 47 | 46 | 31 | 36 | 63 | 22 | 18 | 54 | 12 | 62 | 59 | 26 | 29 | 45 | 15 | 56 | 7 | 34 | 2 | 23 | 1 | 28 | 1 | 16 | 1 | 11 |
| | | | 3 | 1135 | 11 | 21 | 33 | 9 | 23 | 18 | 28 | 26 | 10 | 27 | 10 | 27 | 7 | 13 | 2 | 9 | 3 | 18 | 3 | 0 | 1 | 2 | 3 | 0 | 0 | 0 | 0 | 0 |
| $w^{1118/CS10}$ ♀ x *DJ694* ♂ | 1 | 04-03-16 | 1 | 1258 | 33 | 36 | 42 | 48 | 30 | 23 | 30 | 49 | 44 | 21 | 48 | 35 | 25 | 24 | 38 | 10 | 29 | 24 | 11 | 32 | 19 | 12 | 17 | 18 | 22 | 7 | 14 | 12 |
| | | | 2 | 1595 | 60 | 74 | 53 | 52 | 26 | 52 | 40 | 54 | 38 | 37 | 51 | 30 | 31 | 18 | 20 | 17 | 22 | 17 | 21 | 23 | 22 | 23 | 25 | 13 | 16 | 7 | 23 | 19 |
| | | | 3 | 972 | 37 | 51 | 46 | 37 | 33 | 35 | 43 | 29 | 43 | 35 | 30 | 29 | 23 | 19 | 20 | 23 | 16 | 18 | 6 | 8 | 10 | 9 | 11 | 10 | 8 | 16 | 3 | 10 |
| | 2 | 05-02-20 | 1 | 1959 | 36 | 68 | 30 | 40 | 37 | 38 | 24 | 44 | 50 | 66 | 50 | 49 | 40 | 53 | 34 | 39 | 36 | 31 | 36 | 49 | 35 | 24 | 14 | 30 | 42 | 45 | 34 | 38 |
| | | | 2 | 2019 | 71 | 29 | 46 | 44 | 43 | 28 | 34 | 38 | 48 | 39 | 52 | 49 | 37 | 37 | 34 | 28 | 23 | 24 | 19 | 16 | 22 | 21 | 33 | 36 | 29 | 7 | 59 | 22 |
| | | | 3 | 2090 | 43 | 47 | 38 | 45 | 17 | 54 | 33 | 38 | 48 | 48 | 41 | 34 | 40 | 17 | 38 | 14 | 21 | 61 | 34 | 36 | 50 | 35 | 32 | 23 | 33 | 21 | 54 | 20 |
| *DJ694* ♀ x $w^{1118/CS10}$ ♂ | 1 | 04-03-16 | 1 | 552 | 4 | 52 | 17 | 65 | 7 | 31 | 48 | 10 | 2 | 59 | 36 | 6 | 49 | 3 | 20 | 3 | 3 | 6 | 3 | 1 | 5 | 2 | 0 | 2 | 3 | 2 | 3 | 0 |
| | | | 2 | 883 | 33 | 54 | 60 | 35 | 30 | 56 | 4 | 34 | 57 | 36 | 65 | 10 | 46 | 5 | 52 | 22 | 10 | 28 | 2 | 15 | 7 | 5 | 5 | 13 | 6 | 3 | 6 | 8 |
| | | | 3 | 970 | 5 | 44 | 47 | 25 | 18 | 68 | 34 | 48 | 51 | 12 | 87 | 13 | 45 | 6 | 65 | 2 | 4 | 56 | 4 | 2 | 32 | 18 | 1 | 11 | 32 | 1 | 2 | 1 |
| | 2 | 05-02-20 | 1 | 1099 | 37 | 50 | 44 | 38 | 46 | 39 | 25 | 21 | 75 | 11 | 28 | 64 | 7 | 26 | 42 | 17 | 17 | 43 | 1 | 10 | 18 | 1 | 9 | 15 | 7 | 6 | 15 | 1 |
| | | | 2 | 1231 | 26 | 34 | 57 | 31 | 45 | 64 | 27 | 43 | 29 | 46 | 6 | 49 | 48 | 45 | 7 | 34 | 5 | 85 | 17 | 8 | 23 | 18 | 16 | 32 | 4 | 5 | 30 | 8 |
| | | | 3 | 994 | 7 | 78 | 22 | 13 | 48 | 43 | 23 | 16 | 59 | 15 | 39 | 10 | 69 | 25 | 44 | 16 | 10 | 43 | 5 | 6 | 41 | 7 | 12 | 26 | 13 | 22 | 6 | 3 |
| $w^{1118/CS10}$ ♀ x $w^{1118}$ ♂ | 1 | 04-03-16 | 1 | 1100 | 67 | 58 | 46 | 34 | 36 | 32 | 27 | 40 | 35 | 28 | 35 | 34 | 20 | 18 | 28 | 13 | 22 | 16 | 12 | 19 | 25 | 13 | 15 | 21 | 30 | 16 | 18 | 31 |
| | | | 2 | 1566 | 46 | 66 | 63 | 37 | 40 | 52 | 25 | 42 | 42 | 39 | 39 | 39 | 43 | 26 | 22 | 35 | 27 | 41 | 7 | 35 | 7 | 31 | 18 | 26 | 26 | 17 | 27 | 11 |
| | | | 3 | 1712 | 37 | 80 | 74 | 52 | 35 | 34 | 33 | 53 | 41 | 33 | 38 | 34 | 28 | 30 | 22 | 19 | 28 | 28 | 12 | 16 | 16 | 20 | 16 | 24 | 27 | 14 | 18 | 12 |
| | 2 | 05-02-20 | 1 | 2007 | 30 | 40 | 39 | 26 | 22 | 55 | 38 | 28 | 35 | 56 | 44 | 39 | 48 | 44 | 20 | 68 | 35 | 32 | 50 | 41 | 43 | 36 | 28 | 44 | 26 | 38 | 14 | 9 |
| | | | 2 | 2123 | 58 | 58 | 23 | 39 | 39 | 26 | 52 | 34 | 30 | 50 | 41 | 37 | 44 | 39 | 41 | 45 | 40 | 43 | 21 | 49 | 48 | 23 | 30 | 40 | 48 | 32 | 23 | 57 |
| | | | 3 | 2165 | 61 | 41 | 36 | 19 | 23 | 35 | 35 | 24 | 28 | 25 | 35 | 40 | 26 | 36 | 22 | 36 | 23 | 37 | 30 | 34 | 32 | 21 | 47 | 39 | 30 | 29 | 31 | 47 |
| *DJ694/+* ♀ x $w^{1118/CS10}$ ♂ | 1 | 04-03-16 | 1 | 1910 | 34 | 35 | 41 | 34 | 28 | 44 | 16 | 34 | 17 | 22 | 26 | 19 | 21 | 12 | 13 | 33 | 16 | 19 | 9 | 21 | 13 | 21 | 32 | 17 | 38 | 16 | 24 | 17 |
| | | | 2 | 2085 | 35 | 88 | 35 | 67 | 28 | 58 | 14 | 74 | 31 | 55 | 55 | 48 | 43 | 26 | 43 | 12 | 62 | 35 | 16 | 24 | 47 | 19 | 34 | 20 | 40 | 34 | 19 | 15 |
| | | | 3 | 1955 | 47 | 79 | 30 | 75 | 24 | 60 | 39 | 41 | 47 | 32 | 40 | 37 | 50 | 15 | 29 | 36 | 32 | 27 | 22 | 27 | 16 | 27 | 24 | 35 | 25 | 24 | 24 | 16 |
| $w^{1118/CS10}$ ♀ x *DJ694/+* ♂ | 1 | 04-03-16 | 1 | 1795 | 64 | 69 | 47 | 43 | 35 | 31 | 23 | 40 | 37 | 54 | 33 | 37 | 34 | 22 | 37 | 28 | 27 | 26 | 12 | 18 | 21 | 13 | 32 | 34 | 38 | 32 | 24 | 20 |
| | | | 2 | 1089 | 64 | 60 | 43 | 39 | 34 | 42 | 30 | 35 | 38 | 43 | 34 | 33 | 23 | 15 | 21 | 28 | 23 | 10 | 7 | 26 | 14 | 22 | 11 | 31 | 18 | 18 | 21 | 4 |
| | | | 3 | 1573 | 46 | 57 | 58 | 22 | 36 | 33 | 35 | 32 | 41 | 31 | 33 | 26 | 24 | 12 | 30 | 25 | 33 | 24 | 24 | 17 | 12 | 23 | 21 | 19 | 32 | 18 | 28 | 18 |

Table S6. Number of flies that emerged from eggs to adults for the indicated genotypes per each day 

| Cross | Rep | Start Date (YYMMDD) | Vial | 29 | 30 | 31 | 32 | 33 | 34 | 35 | 36 | 37 | 38 | 39 | 40 | 41 | 42 | 43 | 44 | 45 | 46 | 47 | 48 | 49 | 50 | 51 | 52 | 53 | 54 | 55 | 56 | 57 | 58 |
|---|---|---|---|---|---|---|---|---|---|---|---|---|---|---|---|---|---|---|---|---|---|---|---|---|---|---|---|---|---|---|---|---|---|
| *DJ694* ♀ x *DJ694* ♂ | 1 | 04-03-16 | 1 | 0 | 0 | 0 | 0 | 0 | 0 | 0 | 0 | 0 | - | - | - | 0 | - | - | - | - | - | - | - | - | - | - | - | - | - | - | - | x | x |
| | | | 2 | 3 | 7 | 0 | 2 | 5 | 0 | 0 | 1 | 1 | 1 | 0 | 0 | 0 | 0 | 0 | 0 | 0 | 0 | 0 | 0 | 0 | 0 | - | - | - | - | - | - | - | - |
| | | | 3 | 0 | - | x | x | x | x | x | x | x | x | x | x | x | x | x | x | x | x | x | x | x | x | x | x | x | x | x | x | x | x |
| | 2 | 05-02-20 | 1 | 5 | 15 | 3 | 11 | 1 | 1 | 0 | 3 | 0 | 2 | 0 | 0 | 0 | 0 | 0 | 0 | 0 | 0 | 0 | 0 | 0 | 0 | 0 | 0 | 0 | 0 | 0 | x | x | x |
| | | | 2 | 0 | 1 | 1 | 3 | 1 | 0 | 1 | 0 | 0 | 0 | 0 | 0 | 0 | 0 | 0 | 0 | 0 | 0 | 0 | 0 | 0 | 0 | 0 | 0 | 0 | 0 | 0 | x | x | x |
| | | | 3 | 0 | 0 | 0 | 0 | 0 | 0 | 0 | 0 | 0 | 0 | 0 | 0 | 0 | 0 | 0 | 0 | 0 | 0 | 0 | 0 | 0 | 0 | 0 | 0 | 0 | 0 | 0 | 0 | 0 | 0 |
| *w* $^{1118/CS10}$ ♀ x *DJ694* ♂ | 1 | 04-03-16 | 1 | 8 | 10 | 11 | 2 | 2 | 10 | 7 | 2 | 1 | 1 | 2 | 5 | 0 | 0 | 0 | - | 0 | 0 | 0 | 0 | 0 | 0 | 0 | 0 | 0 | 0 | 0 | 3 | - | 0 |
| | | | 2 | 10 | 22 | 15 | 2 | 2 | 15 | 11 | 6 | 5 | 3 | 8 | 5 | 1 | 2 | 5 | 0 | 1 | 0 | 0 | 0 | 0 | - | 0 | 0 | 0 | 0 | 0 | - | 0 | 0 |
| | | | 3 | 9 | 11 | 6 | 3 | 9 | 8 | 6 | 5 | 1 | 6 | 4 | 2 | 3 | 1 | 0 | - | - | 0 | 0 | - | 0 | - | - | 0 | - | - | 0 | - | 0 | 0 |
| | 2 | 05-02-20 | 1 | 21 | 20 | 13 | 19 | 36 | 13 | 14 | 37 | 11 | 13 | 6 | 10 | 7 | 1 | 0 | 1 | 0 | 0 | 0 | 0 | 0 | 0 | 0 | 0 | 0 | 0 | 0 | 0 | 0 | 0 |
| | | | 2 | 24 | 36 | 31 | 10 | 35 | 21 | 25 | 27 | 6 | 22 | 6 | 12 | 14 | 10 | 8 | 5 | 7 | 9 | 3 | 2 | 1 | 0 | 0 | 5 | 1 | 0 | 3 | 0 | 0 | 0 |
| | | | 3 | 39 | 21 | 30 | 26 | 28 | 22 | 23 | 9 | 17 | 6 | 8 | 3 | 9 | 7 | 2 | 0 | 1 | 0 | 0 | 0 | 0 | 0 | 0 | 0 | 0 | 0 | 2 | 0 | 0 | 0 |
| *DJ694* ♀ x *w* $^{1118/CS10}$ ♂ | 1 | 04-03-16 | 1 | 3 | 1 | 1 | 0 | 0 | 0 | 0 | - | - | - | - | - | - | - | - | - | - | - | - | x | x | x | x | x | x | x | x | x | x | x |
| | | | 2 | 3 | 4 | 6 | 2 | 6 | 1 | 1 | 2 | 4 | 2 | 1 | 0 | 0 | 0 | - | - | - | - | - | - | - | - | - | - | - | x | x | x | x | x |
| | | | 3 | 8 | 4 | 0 | 1 | 10 | 13 | 1 | 4 | 2 | 0 | 7 | 1 | 0 | 3 | 1 | - | - | - | 3 | - | 0 | 0 | 0 | - | - | - | - | - | x | x |
| | 2 | 05-02-20 | 1 | 5 | 13 | 1 | 2 | 0 | 2 | 1 | 0 | 0 | 0 | 0 | 0 | 0 | 0 | 0 | 0 | 0 | 0 | 0 | 0 | 0 | 0 | 0 | 0 | 0 | 0 | x | x | x | x |
| | | | 2 | 6 | 4 | 2 | 2 | 0 | 11 | 0 | 0 | 0 | 0 | 0 | 0 | 0 | 0 | 0 | 0 | 0 | 0 | 0 | 0 | 0 | 0 | 0 | 0 | 0 | 0 | 0 | 0 | 0 | 0 |
| | | | 3 | 5 | 8 | 32 | 0 | 9 | 4 | 11 | 1 | 9 | 0 | 0 | 0 | 0 | 0 | 0 | 0 | 0 | 0 | 0 | 0 | 0 | 0 | 0 | x | x | x | x | x | x | x |
| *w* $^{1118/CS10}$ ♀ x *w* $^{1118}$ ♂ | 1 | 04-03-16 | 1 | 6 | 23 | 19 | 2 | 18 | 10 | 11 | 12 | 14 | 13 | 7 | 7 | 10 | 10 | 7 | 1 | 4 | 3 | 0 | 0 | 0 | 0 | - | - | - | - | 0 | - | - | 0 |
| | | | 2 | 39 | 31 | 36 | 21 | 33 | 14 | 12 | 16 | 12 | 19 | 20 | 15 | 16 | 14 | 14 | 23 | 19 | 19 | 11 | 6 | 16 | 5 | 2 | 1 | 0 | 5 | 11 | 3 | 3 | 0 |
| | | | 3 | 27 | 19 | 20 | 10 | 14 | 9 | 17 | 7 | 12 | 14 | 10 | 14 | 12 | 14 | 19 | 12 | 19 | 13 | 20 | 17 | 18 | 11 | 11 | 6 | 4 | 4 | 7 | 2 | 9 | 2 |
| | 2 | 05-02-20 | 1 | 43 | 34 | 17 | 30 | 23 | 20 | 38 | 7 | 32 | 10 | 18 | 7 | 31 | 10 | 4 | 27 | 19 | 20 | 36 | 20 | 33 | 33 | 22 | 28 | 27 | 16 | 16 | 21 | 2 | 13 |
| | | | 2 | 28 | 36 | 26 | 31 | 26 | 33 | 11 | 9 | 35 | 16 | 7 | 6 | 28 | 13 | 16 | 12 | 15 | 27 | 21 | 17 | 16 | 30 | 33 | 34 | 24 | 19 | 19 | 15 | 6 | 20 |
| | | | 3 | 26 | 34 | 32 | 22 | 36 | 17 | 36 | 10 | 36 | 12 | 8 | 4 | 24 | 25 | 11 | 32 | 26 | 27 | 31 | 28 | 31 | 40 | 36 | 21 | 18 | 22 | 19 | 15 | 4 | 24 |
| *DJ694/+* ♀ x *w* $^{1118/CS10}$ ♂ | 1 | 04-03-16 | 1 | 32 | 26 | 27 | 27 | 34 | 34 | 24 | 27 | 3 | 25 | 33 | 14 | 11 | 25 | 33 | 6 | 7 | 11 | 17 | 10 | 13 | 4 | 28 | 3 | 5 | 14 | 13 | 6 | 17 | 9 |
| | | | 2 | 46 | 41 | 53 | 10 | 60 | 29 | 17 | 18 | 22 | 10 | 17 | 34 | 11 | 28 | 35 | 18 | 15 | 18 | 24 | 28 | 13 | 22 | 19 | 0 | 10 | 18 | 7 | 9 | 37 | 19 |
| | | | 3 | 36 | 26 | 34 | 14 | 37 | 26 | 26 | 25 | 16 | 17 | 24 | 15 | 22 | 29 | 31 | 22 | 14 | 6 | 43 | 15 | 22 | 20 | 26 | 10 | 9 | 12 | 21 | 8 | 28 | 12 |
| *w* $^{1118/CS10}$ ♀ x *DJ694/+* ♂ | 1 | 04-03-16 | 1 | 34 | 33 | 32 | 20 | 29 | 19 | 25 | 23 | 15 | 27 | 16 | 14 | 6 | 25 | 37 | 7 | 25 | 29 | 22 | 27 | 22 | 20 | 17 | 19 | 14 | 11 | 17 | 10 | 13 | 6 |
| | | | 2 | 23 | 10 | 13 | 5 | 31 | 8 | 16 | 8 | 10 | 4 | 1 | 4 | 3 | 4 | 3 | 4 | 4 | 1 | 0 | 1 | 4 | 2 | 0 | - | 0 | 0 | 0 | - | - | - |
| | | | 3 | 27 | 30 | 21 | 13 | 24 | 8 | 13 | 16 | 17 | 13 | 11 | 11 | 8 | 17 | 20 | 8 | 22 | 18 | 12 | 22 | 18 | 11 | 16 | 2 | 13 | 13 | 23 | 4 | 22 | 15 |

Table S6. Number of flies that emerged from eggs to adults for the indicated genotypes per each day 

| Cross | Rep | Start Date (YYMMDD) | Vial | 59 | 60 | 61 | 62 | 63 | 64 | 65 | 66 | 67 | 68 | 69 | 70 | 71 | 72 | 73 | 74 | 75 | 76 | 77 | 78 | 79 | 80 | 81 | 82 | 83 | 84 | 85 |
|---|---|---|---|---|---|---|---|---|---|---|---|---|---|---|---|---|---|---|---|---|---|---|---|---|---|---|---|---|---|---|
| *DJ694* ♀ x *DJ694* ♂ | 1 | 04-03-16 | 1 | x | x | x | x | x | x | x | x | x | x | x | x | x | x | x | x | x | x | x | x | x | x | x | x | x | x | x |
| | | | 2 | - | x | x | x | x | x | x | x | x | x | x | x | x | x | x | x | x | x | x | x | x | x | x | x | x | x | x |
| | | | 3 | x | x | x | x | x | x | x | x | x | x | x | x | x | x | x | x | x | x | x | x | x | x | x | x | x | x | x |
| | 2 | 05-02-20 | 1 | x | x | x | x | x | x | x | x | x | x | x | x | x | x | x | x | x | x | x | x | x | x | x | x | x | x | x |
| | | | 2 | x | x | x | x | x | x | x | x | x | x | x | x | x | x | x | x | x | x | x | x | x | x | x | x | x | x | x |
| | | | 3 | x | x | x | x | x | x | x | x | x | x | x | x | x | x | x | x | x | x | x | x | x | x | x | x | x | x | x |
| $w^{1118/CS10}$ ♀ x *DJ694* ♂ | 1 | 04-03-16 | 1 | 0 | 0 | - | 0 | 0 | 0 | - | - | - | - | - | x | x | x | x | x | x | x | x | x | x | x | x | x | x | x | x |
| | | | 2 | 0 | 0 | - | - | - | - | - | - | - | - | - | - | - | - | - | x | x | x | x | x | x | x | x | x | x | x | x |
| | | | 3 | - | - | - | 0 | - | - | - | 0 | 0 | 0 | - | - | - | - | - | - | - | - | - | - | - | - | - | - | - | - | - |
| | 2 | 05-02-20 | 1 | 0 | 0 | 0 | 0 | 0 | 0 | 0 | 0 | 0 | 0 | 0 | 0 | 0 | 0 | 0 | 0 | 0 | 0 | 0 | 0 | 0 | 0 | 0 | 0 | 0 | 0 | 0 |
| | | | 2 | 0 | 0 | 0 | 0 | 0 | 0 | 0 | 0 | 0 | 0 | 0 | 0 | 0 | 0 | 0 | 0 | 0 | 0 | 0 | 0 | 0 | 0 | 0 | 0 | 0 | 0 | 0 |
| | | | 3 | 0 | 0 | 0 | 0 | 0 | 0 | 0 | 0 | 0 | 0 | 0 | 0 | 0 | 0 | 0 | 0 | 0 | 0 | 0 | 0 | 0 | 0 | 0 | x | x | x | x |
| *DJ694* ♀ x $w^{1118/CS10}$ ♂ | 1 | 04-03-16 | 1 | x | x | x | x | x | x | x | x | x | x | x | x | x | x | x | x | x | x | x | x | x | x | x | x | x | x | x |
| | | | 2 | x | x | x | x | x | x | x | x | x | x | x | x | x | x | x | x | x | x | x | x | x | x | x | x | x | x | x |
| | | | 3 | x | x | x | x | x | x | x | x | x | x | x | x | x | x | x | x | x | x | x | x | x | x | x | x | x | x | x |
| | 2 | 05-02-20 | 1 | x | x | x | x | x | x | x | x | x | x | x | x | x | x | x | x | x | x | x | x | x | x | x | x | x | x | x |
| | | | 2 | x | x | x | x | x | x | x | x | x | x | x | x | x | x | x | x | x | x | x | x | x | x | x | x | x | x | x |
| | | | 3 | x | x | x | x | x | x | x | x | x | x | x | x | x | x | x | x | x | x | x | x | x | x | x | x | x | x | x |
| $w^{1118/CS10}$ ♀ x $w^{1118}$ ♂ | 1 | 04-03-16 | 1 | - | 0 | - | - | - | - | - | - | - | - | - | - | - | - | - | - | - | - | 0 | - | - | - | - | x | x | x | x |
| | | | 2 | 13 | 5 | 10 | 8 | 2 | 0 | - | - | x | x | x | x | x | x | x | x | x | x | x | x | x | x | x | x | x | x | x |
| | | | 3 | 4 | 1 | 12 | 4 | 2 | 4 | 3 | 0 | 0 | 0 | 0 | 0 | 0 | 0 | 2 | 0 | 0 | - | 1 | - | - | - | - | - | - | - | - |
| | 2 | 05-02-20 | 1 | 21 | 17 | 10 | 12 | 10 | 11 | 3 | 2 | 1 | 3 | 2 | 0 | 1 | 0 | 0 | 0 | 0 | 0 | 0 | 0 | 0 | 0 | 0 | 0 | 0 | 0 | x |
| | | | 2 | 11 | 15 | 13 | 10 | 14 | 2 | 1 | 1 | 3 | 1 | 0 | 0 | 0 | 0 | 0 | 0 | 0 | 0 | 0 | 0 | 0 | 0 | x | x | x | x | x |
| | | | 3 | 12 | 10 | 13 | 14 | 12 | 9 | 4 | 5 | 7 | 2 | 3 | 3 | 0 | 0 | 1 | 0 | 0 | 0 | 0 | 0 | 0 | 0 | 0 | 0 | 0 | 0 | x |
| *DJ694/+* ♀ x $w^{1118/CS10}$ ♂ | 1 | 04-03-16 | 1 | 7 | 26 | 2 | 19 | 11 | 22 | 7 | 21 | 14 | 14 | 8 | 1 | 9 | 15 | 6 | 2 | 1 | 2 | 0 | - | 0 | 0 | - | - | - | - | - |
| | | | 2 | 18 | 34 | 12 | 10 | 10 | 25 | 13 | 11 | 3 | 21 | 3 | 0 | 2 | 1 | 0 | 1 | 0 | - | - | - | - | - | - | - | x | x | x |
| | | | 3 | 23 | 19 | 29 | 34 | 20 | 14 | 8 | 19 | 11 | 7 | 1 | 1 | 7 | 5 | 1 | 0 | 1 | 1 | - | - | - | 0 | 0 | - | 0 | 0 | - |
| $w^{1118/CS10}$ ♀ x *DJ694/+* ♂ | 1 | 04-03-16 | 1 | 14 | 11 | 7 | 11 | 9 | 3 | 13 | 11 | 7 | 6 | 2 | 0 | 4 | - | - | 5 | 1 | - | 0 | - | - | - | 0 | - | - | - | - |
| | | | 2 | - | - | - | - | - | - | - | - | - | - | x | x | x | x | x | x | x | x | x | x | x | x | x | x | x | x | x |
| | | | 3 | 23 | 10 | 20 | 13 | 12 | 13 | 4 | 6 | - | 3 | 0 | - | 0 | 0 | 0 | 0 | - | - | - | - | 0 | 0 | - | 0 | - | - | - |

Table S6. Number of flies that emerged from eggs to adults for the indicated genotypes per each day

| Cross | Rep | Start Date (YYMMDD) | Vial | Days | | | | | | | | | | | | | | | | | | | |
|---|---|---|---|---|---|---|---|---|---|---|---|---|---|---|---|---|---|---|---|---|---|---|---|
| | | | | 29 | 30 | 31 | 32 | 33 | 34 | 35 | 36 | 37 | 38 | 39 | 40 | 41 | 42 | 43 | 44 | 45 | 46 | 47 | 48 |
| *w* $^{1118}$ ♀ x *DJ694;UAS-EDTP* $^{A}$ ♂ | 3 | 04-11-30 | 1 | 27 | 13 | 6 | 0 | 1 | 0 | 2 | 0 | 0 | 0 | 0 | 0 | 0 | 0 | 0 | 0 | 0 | 0 | 0 | 0 |
| | | | 2 | 10 | 18 | 15 | 8 | 14 | 8 | 10 | 12 | 9 | 7 | 9 | 0 | 0 | 0 | 0 | 0 | 12 | 4 | 3 | 1 |
| | | | 3 | 10 | 10 | 12 | 4 | 7 | 1 | 2 | 0 | 0 | 0 | 0 | 2 | 0 | 0 | 0 | 0 | 0 | 0 | 0 | 0 |
| *DJ694;UAS-EDTP* $^{A}$ ♀ x *w* $^{1118}$ ♂ | 3 | 04-11-30 | 1 | 13 | 33 | 15 | 18 | 16 | 5 | 21 | 7 | 7 | 22 | 11 | 5 | 0 | 13 | 28 | 1 | 0 | 0 | 3 | 2 |
| | | | 2 | 33 | 7 | 24 | 18 | 14 | 9 | 11 | 9 | 17 | 17 | 18 | 18 | 14 | 12 | 19 | 5 | 5 | 2 | 0 | 0 |
| | | | 3 | 0 | 6 | 16 | 0 | 1 | 0 | 0 | 0 | 0 | 0 | 0 | 0 | 0 | 0 | 0 | 0 | 0 | 0 | 0 | 0 |
| *w* $^{1118}$ ♀ x *DJ694* ♂ | 3 | 04-11-30 | 1 | 1 | 0 | 0 | 0 | 0 | 0 | 0 | 0 | 0 | 0 | 0 | 0 | 0 | 0 | 0 | 0 | 0 | 0 | 0 | 0 |
| | | | 2 | 2 | 0 | 0 | 0 | 0 | 0 | 0 | 0 | 0 | 0 | 0 | 0 | 0 | 0 | 0 | 0 | 0 | 0 | 0 | 0 |
| | | | 3 | 3 | 0 | 1 | 0 | 6 | 3 | 8 | 3 | 1 | 0 | 0 | 0 | 2 | 0 | 2 | 0 | 1 | 1 | 0 | 0 |
| | | | 4 | 2 | 10 | 0 | 10 | 2 | 7 | 13 | 0 | 7 | 13 | 7 | 1 | 2 | 0 | 0 | 0 | 0 | 0 | 0 | 0 |
| *w* $^{1118}$ ♀ x *DJ694;UAS-EDTP* $^{B}$ ♂ | 3 | 04-11-30 | 1 | 17 | 22 | 18 | 24 | 10 | 11 | 14 | 15 | 4 | 1 | 2 | 3 | 4 | 2 | 17 | 15 | 9 | 10 | 7 | 3 |
| | | | 2 | 21 | 19 | 31 | 21 | 15 | 10 | 20 | 12 | 14 | 24 | 13 | 17 | 18 | 2 | 15 | 7 | 8 | 4 | 0 | 0 |
| | | | 3 | 12 | 14 | 0 | 4 | 11 | 15 | 15 | 5 | 10 | 17 | 8 | 10 | 12 | 9 | 12 | 7 | 1 | 0 | 2 | 2 |
| | | | 4 | 16 | 15 | 9 | 4 | 8 | 5 | 8 | 12 | 11 | 8 | 3 | 8 | 13 | 7 | 8 | 4 | 2 | 0 | 8 | 4 |
| *DJ694;UAS-EDTP* $^{B}$ ♀ x *w* $^{1118}$ ♂ | 3 | 04-11-30 | 1 | 8 | 9 | 36 | 9 | 27 | 17 | 17 | 17 | 15 | 12 | 9 | 17 | 12 | 15 | 15 | 6 | 7 | 4 | 1 | 11 |
| | | | 2 | 15 | 10 | 18 | 11 | 4 | 16 | 27 | 15 | 24 | 15 | 16 | 12 | 17 | 20 | 23 | 3 | 2 | 11 | 1 | 9 |
| | | | 3 | 21 | 15 | 24 | 9 | 9 | 16 | 9 | 11 | 3 | 14 | 8 | 14 | 28 | 23 | 16 | 13 | 2 | 5 | 2 | 8 |
| *DJ694* ♀ x *w* $^{1118}$ ♂ | 3 | 04-11-30 | 1 | 0 | 0 | 0 | 0 | 0 | 0 | 0 | 0 | 0 | 0 | 0 | 0 | 0 | 0 | x | x | x | x | x | x |
| | | | 2 | 5 | 0 | 11 | 4 | 8 | 0 | 0 | 0 | 0 | 0 | 0 | 0 | 0 | 0 | 0 | 0 | 0 | 0 | 0 | 0 |
| | | | 3 | 3 | 0 | 2 | 3 | 1 | 1 | 1 | 4 | 0 | 1 | 4 | 1 | 1 | 0 | 0 | 0 | 0 | 0 | 0 | 0 |
| | | | 4 | 1 | 0 | 0 | 0 | 1 | 1 | 0 | 0 | 0 | 0 | 0 | 0 | 0 | 0 | 0 | 0 | 0 | 0 | 0 | 0 |

For each cross, multiple biological replicates and vials were scored, and the number of eggs that successfully emerge as adults are reported for each day following the start date. The “Start Date (YYMMDD)” column indicates the date on which egg laying began for each replicateColumn “Total Egg” indicates the total number of eggs laid, either across the full 85-day period or within days 29 to 48, depending on the replicate. An “x” indicates that all females within the vial had died, and “–” indicates that data were not applicable.

| | | | | Avg sum/♀ at 40d | | vs. OE | | vs.*DJ694/+* | |
|---|---|---|---|---|---|---|---|---|---|
| **Rep** | **Date** | **Female's Genotype** | **n♀** | **Mean** | **SD** | **p** | **Δ** | **p** | **Δ** |
| 1 | 2024-09-28 | *DJ694$^{L}$/+;UAS-EDTP$^{E}$/+* | 24 | 274.25 | 27.57 | - | - | - | - |
| 1 | 2024-09-28 | *DJ694$^{L}$/+* | 24 | 289.13 | 21.50 | 0.3964 | X | - | - |
| 1 | 2024-09-28 | *UAS-EDTP$^{E}$/+* | 24 | 308.27 | 26.54 | 0.0247 | X | 0.0492 | X |
| 2 | 2024-09-28 | *DJ694$^{L}$/+;UAS-EDTP$^{E}$/+* | 24 | 261.15 | 26.67 | - | - | - | - |
| 2 | 2024-09-28 | *DJ694$^{L}$/+* | 24 | 265.08 | 23.33 | 0.7503 | X | - | - |
| 2 | 2024-09-28 | *UAS-EDTP$^{E}$/+* | 24 | 292.96 | 21.27 | 0.0195 | X | 0.0210 | X |

Column “Rep” denotes the experimental ID for each independent biological replicate. Column “Date” indicates the start date of the fertility assay. Column “Female Genotype” indicates the genotype of the females; all females were mated with w males.
The value of n♀ represents the number of females at the start of the assay (3 females per vial, 8 vials per genotype in a given replicate); some females died during the course of the experiment.
“Avg Sum/♀” represents the average cumulative number of eggs laid per female from day 1 to day 40. Mean and standard deviation (SD) are shown for each measurement, calculated from 8 vials per genotype within a single replicate.
Column “vs. OE” presents the statistical comparison to *DJ694/+; UAS-EDTP/+*.
Column “vs. *DJ694/+*” presents the statistical comparison to the heterozygous control. Both using two-sample, two-tailed t-tests assuming equal variance. Within these two sections, column “p” indicates the p-value, and column “Δ” indicates whether a noticeable change relative to the overexpression group is observed. “X” indicates that error bars overlap.

Table S8. Daily female fertility measurements (eggs laid per female per day) for *DJ694/+;UAS-EDTP/+* and controls

| | | Start Date | | Days | | | | | | | | | | | | | |
|---|---|---|---|---|---|---|---|---|---|---|---|---|---|---|---|---|---|
| ***Female's Geno*** | **Rep** | **(YYMMDD)** | **vial** | **1** | **2** | **3** | **4** | **5** | **6** | **7** | **8** | **9** | **10** | **11** | **12** | **13** | **14** |
| *DJ694 $^{L}$ /+;UAS-EDTP $^{E}$ /+* | 1 | 24-09-28 | 1 | 7.67 | 3.00 | 10.00 | 2.00 | 19.33 | 19.00 | 6.33 | 15.33 | 4.33 | 16.00 | 13.00 | 7.67 | 3.33 | 10.00 |
| | | | 2 | 5.00 | 5.00 | 10.33 | 1.00 | 22.33 | 13.00 | 10.33 | 14.00 | 9.33 | 3.00 | 5.67 | 4.67 | 8.67 | 16.67 |
| | | | 3 | 7.33 | 3.33 | 25.67 | 18.33 | 4.33 | 19.33 | 8.33 | 16.67 | 5.67 | 8.00 | 4.67 | 10.00 | 7.00 | 14.67 |
| | | | 4 | 12.67 | 10.33 | 4.33 | 4.00 | 24.00 | 11.33 | 7.67 | 18.33 | 3.00 | 7.00 | 10.33 | 2.00 | 7.00 | 7.33 |
| | | | 5 | 11.00 | 9.67 | 15.33 | 6.67 | 21.33 | 8.00 | 20.33 | 7.33 | 9.33 | 9.00 | 1.00 | 13.33 | 13.33 | 0.33 |
| | | | 6 | 5.00 | 3.33 | 1.67 | 2.33 | 14.00 | 13.67 | 16.33 | 14.33 | 1.67 | 14.67 | 1.00 | 15.00 | 10.00 | 16.33 |
| | | | 7 | 5.00 | 1.33 | 0.33 | 8.67 | 16.67 | 7.33 | 11.33 | 16.67 | 1.00 | 14.00 | 1.00 | 23.00 | 19.67 | 12.67 |
| | | | 8 | 3.67 | 3.67 | 9.67 | 4.00 | 10.33 | 11.67 | 10.00 | 1.67 | 5.67 | 22.67 | 1.00 | 6.33 | 3.33 | 19.33 |
| | 2 | | 1 | 9.67 | 8.67 | 26.67 | 18.33 | 18.33 | 6.00 | 15.67 | 16.67 | 8.00 | 8.67 | 7.33 | 10.00 | 7.33 | 10.00 |
| | | | 2 | 3.33 | 1.00 | 1.00 | 11.33 | 18.33 | 5.67 | 4.67 | 17.00 | 4.00 | 13.33 | 9.00 | 11.00 | 13.33 | 9.67 |
| | | | 3 | 5.33 | 4.00 | 6.33 | 2.00 | 18.67 | 1.00 | 7.33 | 13.00 | 3.67 | 13.67 | 1.67 | 14.33 | 3.33 | 4.67 |
| | | | 4 | 16.33 | 1.00 | 5.67 | 26.67 | 6.00 | 10.33 | 12.33 | 11.33 | 10.67 | 13.00 | 8.33 | 6.33 | 2.00 | 12.00 |
| | | | 5 | 4.67 | 1.33 | 9.00 | 7.00 | 13.33 | 2.33 | 8.00 | 23.67 | 14.33 | 3.67 | 2.67 | 7.33 | 8.67 | 6.33 |
| | | | 6 | 7.33 | 6.00 | 0.67 | 1.67 | 14.00 | 1.67 | 3.00 | 4.00 | 8.33 | 12.67 | 5.33 | 13.33 | 5.00 | 4.00 |
| | | | 7 | 11.00 | 5.00 | 1.00 | 11.00 | 4.67 | 16.00 | 5.00 | 4.67 | 8.33 | 5.00 | 1.00 | 14.67 | 12.33 | 4.33 |
| | | | 8 | 11.67 | 0.67 | 1.33 | 3.33 | 10.67 | 13.67 | 19.67 | 10.00 | 2.33 | 21.00 | 0.67 | 12.67 | 0.33 | 10.00 |

| | | Start Date | | Days | | | | | | | | | | | | | |
|---|---|---|---|---|---|---|---|---|---|---|---|---|---|---|---|---|---|
| ***Female's Geno*** | **Rep** | **(YYMMDD)** | **vial** | **15** | **16** | **17** | **18** | **19** | **20** | **21** | **22** | **23** | **24** | **25** | **26** | **27** | **28** |
| *DJ694*$^{L}$*/+;UAS-EDTP*$^{E}$*/+* | 1 | 24-09-28 | 1 | 1.00 | 9.67 | 10.33 | 3.33 | 5.33 | 9.33 | 4.00 | 3.00 | 9.67 | 1.67 | 6.00 | 6.33 | 1.33 | 3.33 |
| | | | 2 | 13.33 | 4.00 | 6.67 | 2.00 | 1.33 | 14.67 | 6.67 | 12.33 | 7.00 | 1.67 | 3.00 | 1.33 | 1.00 | 7.33 |
| | | | 3 | 7.67 | 8.33 | 8.33 | 1.33 | 5.00 | 15.00 | 7.00 | 6.00 | 6.00 | 2.67 | 6.00 | 3.00 | 1.00 | 4.33 |
| | | | 4 | 7.33 | 0.67 | 3.33 | 1.33 | 10.00 | 12.00 | 9.00 | 2.33 | 6.67 | 1.67 | 2.33 | 3.33 | 2.67 | 2.67 |
| | | | 5 | 5.33 | 6.67 | 9.33 | 3.00 | 3.00 | 16.33 | 12.33 | 10.00 | 8.33 | 0.67 | 1.67 | 5.00 | 1.33 | 2.00 |
| | | | 6 | 11.67 | 6.33 | 1.00 | 5.00 | 5.33 | 12.67 | 3.33 | 20.33 | 6.33 | 1.33 | 1.33 | 4.33 | 2.00 | 4.33 |
| | | | 7 | 9.67 | 8.33 | 6.00 | 1.67 | 1.00 | 19.67 | 8.67 | 3.33 | 0.67 | 1.00 | 1.67 | 8.67 | 2.33 | 6.67 |
| | | | 8 | 4.00 | 10.00 | 1.33 | 1.67 | 4.67 | 14.33 | 1.00 | 8.33 | 9.33 | 1.67 | 1.00 | 4.33 | 4.33 | 1.67 |
| | 2 | | 1 | 1.67 | 7.00 | 1.00 | 8.33 | 2.33 | 10.00 | 2.67 | 5.00 | 5.33 | 2.00 | 5.00 | 0.67 | 3.33 | 2.33 |
| | | | 2 | 3.33 | 4.33 | 7.67 | 12.33 | 0.67 | 19.33 | 8.00 | 9.33 | 8.67 | 0.67 | 10.67 | 4.00 | 1.33 | 14.67 |
| | | | 3 | 4.33 | 3.00 | 0.67 | 6.00 | 5.67 | 13.00 | 12.33 | 6.33 | 9.33 | 2.33 | 4.00 | 2.33 | 2.33 | 8.67 |
| | | | 4 | 4.67 | 0.33 | 3.00 | 1.67 | 11.67 | 16.33 | 0.67 | 3.33 | 3.00 | 0.67 | 3.33 | 1.67 | 2.67 | 8.33 |
| | | | 5 | 0.33 | 0.33 | 1.00 | 2.00 | 9.00 | 19.67 | 3.33 | 12.67 | 8.00 | 1.33 | 4.00 | 1.33 | 3.67 | 6.67 |
| | | | 6 | 2.33 | 10.00 | 1.67 | 9.67 | 1.33 | 22.00 | 19.00 | 7.67 | 9.00 | 2.67 | 3.00 | 4.33 | 4.67 | 6.00 |
| | | | 7 | 7.67 | 4.00 | 2.33 | 6.00 | 7.67 | 18.33 | 2.00 | 13.00 | 4.67 | 1.00 | 5.00 | 2.00 | 4.33 | 5.00 |
| | | | 8 | 1.00 | 9.67 | 7.33 | 4.33 | 13.67 | 9.67 | 3.33 | 6.00 | 3.00 | 5.00 | 1.67 | 2.33 | 2.33 | 7.00 |

| *Female's Geno* | Rep | Start Date (YYMMDD) | vial | 29 | 30 | 31 | 32 | 33 | 34 | 35 | 36 | 37 | 38 | 39 | 40 |
|---|---|---|---|---|---|---|---|---|---|---|---|---|---|---|---|
| *DJ694 $^{L}$ /+;UAS-EDTP $^{E}$ /+* | 1 | 24-09-28 | 1 | 5.67 | 6.67 | 4.33 | 3.67 | 3.67 | 6.67 | 1.33 | 3.33 | 2.33 | 7.00 | 4.00 | 4.00 |
| | | | 2 | 6.33 | 3.00 | 5.00 | 15.00 | 4.67 | 10.67 | 5.00 | 4.67 | 2.33 | 1.67 | 10.00 | 2.67 |
| | | | 3 | 9.67 | 1.67 | 9.67 | 3.33 | 2.33 | 6.67 | 7.00 | 5.00 | 6.67 | 2.67 | 5.00 | 8.67 |
| | | | 4 | 0.33 | 0.33 | 0.67 | 0.67 | 4.67 | 3.33 | 4.33 | 9.00 | 4.67 | 1.00 | 2.00 | 2.00 |
| | | | 5 | 1.00 | 2.33 | 3.33 | 8.33 | 9.33 | 15.67 | 11.00 | 4.33 | 4.00 | 3.67 | 8.33 | 8.67 |
| | | | 6 | 1.67 | 1.67 | 9.33 | 9.67 | 3.67 | 6.00 | 3.67 | 3.67 | 4.00 | 10.33 | 4.00 | 4.67 |
| | | | 7 | 1.33 | 1.00 | 14.33 | 16.00 | 2.67 | 5.00 | 8.67 | 3.00 | 3.33 | 1.67 | 4.33 | 2.00 |
| | | | 8 | 0.67 | 1.67 | 6.33 | 10.67 | 5.67 | 5.67 | 3.67 | 10.67 | 4.67 | 2.00 | 9.67 | 5.33 |
| | 2 | | 1 | 1.67 | 4.33 | 6.67 | 3.00 | 1.00 | 3.33 | 4.00 | 3.33 | 3.00 | 1.00 | 3.67 | 3.00 |
| | | | 2 | 3.33 | 1.00 | 2.33 | 8.33 | 8.67 | 8.67 | 8.67 | 11.00 | 8.67 | 4.00 | 15.00 | 7.00 |
| | | | 3 | 6.00 | 0.67 | 1.67 | 12.33 | 8.00 | 3.67 | 2.67 | 10.00 | 2.00 | 3.00 | 8.00 | 7.00 |
| | | | 4 | 5.00 | 2.00 | 4.67 | 8.67 | 5.33 | 5.67 | 7.67 | 3.67 | 6.00 | 5.33 | 7.33 | 5.67 |
| | | | 5 | 2.00 | 4.33 | 18.67 | 3.33 | 4.00 | 6.67 | 2.00 | 4.00 | 6.33 | 4.00 | 10.00 | 1.33 |
| | | | 6 | 7.00 | 1.33 | 7.33 | 13.67 | 8.33 | 5.00 | 2.67 | 5.67 | 4.00 | 5.67 | 5.33 | 5.00 |
| | | | 7 | 4.67 | 2.67 | 2.33 | 9.67 | 12.67 | 6.33 | 5.00 | 6.00 | 3.33 | 6.00 | 10.67 | 4.00 |
| | | | 8 | 2.67 | 1.33 | 2.67 | 6.67 | 1.67 | 3.67 | 1.33 | 0.33 | 1.33 | 0.00 | 2.50 | 1.67 |

Columns 29–40 are under the spanning header "Days".

| | | Start Date | | Days | | | | | | | | | | | | | |
|---|---|---|---|---|---|---|---|---|---|---|---|---|---|---|---|---|---|
| ***Female's Geno*** | **Rep** | **(YYMMDD)** | **vial** | **1** | **2** | **3** | **4** | **5** | **6** | **7** | **8** | **9** | **10** | **11** | **12** | **13** | **14** |
| *DJ694*$^{L}$*/+* | 1 | 24-09-28 | 1 | 11.00 | 9.33 | 7.33 | 15.67 | 26.00 | 3.00 | 14.33 | 19.33 | 8.33 | 10.33 | 4.00 | 6.67 | 3.67 | 11.33 |
| | | | 2 | 13.67 | 10.67 | 1.33 | 24.33 | 11.00 | 7.00 | 12.33 | 12.33 | 5.00 | 15.33 | 2.00 | 9.33 | 7.33 | 11.33 |
| | | | 3 | 9.67 | 5.33 | 21.00 | 11.67 | 14.00 | 6.00 | 9.33 | 12.67 | 12.00 | 3.33 | 6.00 | 11.00 | 9.67 | 7.33 |
| | | | 4 | 15.00 | 6.33 | 23.00 | 5.67 | 19.33 | 14.67 | 6.33 | 15.67 | 4.33 | 17.33 | 6.00 | 3.67 | 6.67 | 11.67 |
| | | | 5 | 6.33 | 4.67 | 14.33 | 9.67 | 9.67 | 8.00 | 8.67 | 15.67 | 13.33 | 9.00 | 4.33 | 6.33 | 7.00 | 13.33 |
| | | | 6 | 8.67 | 8.33 | 11.33 | 9.33 | 17.67 | 6.00 | 13.33 | 13.67 | 9.67 | 12.33 | 3.33 | 3.33 | 7.33 | 5.00 |
| | | | 7 | 11.33 | 9.33 | 10.00 | 13.00 | 17.33 | 9.33 | 5.00 | 9.00 | 11.00 | 8.33 | 8.67 | 4.33 | 7.67 | 12.00 |
| | | | 8 | 17.00 | 11.00 | 29.00 | 1.33 | 26.00 | 2.67 | 13.67 | 20.33 | 1.00 | 18.33 | 16.67 | 13.00 | 1.33 | 9.67 |
| | 2 | | 1 | 28.00 | 6.67 | 9.33 | 23.67 | 18.00 | 1.00 | 25.67 | 7.00 | 11.33 | 11.33 | 12.00 | 6.67 | 2.67 | 15.00 |
| | | | 2 | 20.67 | 0.67 | 13.00 | 24.33 | 3.67 | 3.33 | 22.33 | 10.00 | 15.00 | 9.33 | 9.33 | 8.00 | 8.33 | 11.33 |
| | | | 3 | 18.67 | 5.67 | 8.67 | 28.00 | 7.33 | 18.33 | 14.67 | 5.00 | 12.33 | 3.67 | 5.67 | 12.00 | 11.00 | 5.00 |
| | | | 4 | 4.67 | 15.33 | 7.33 | 17.67 | 1.00 | 1.00 | 27.33 | 6.00 | 13.67 | 10.00 | 10.67 | 6.00 | 1.00 | 13.33 |
| | | | 5 | 9.00 | 10.67 | 8.67 | 1.33 | 17.33 | 6.00 | 9.33 | 11.67 | 5.33 | 10.00 | 9.67 | 6.67 | 6.67 | 4.00 |
| | | | 6 | 6.67 | 7.67 | 1.00 | 20.33 | 1.00 | 7.67 | 23.67 | 15.33 | 3.67 | 10.67 | 7.33 | 6.67 | 7.33 | 5.00 |
| | | | 7 | 0.67 | 5.33 | 0.67 | 13.00 | 15.00 | 1.33 | 17.33 | 5.00 | 9.00 | 10.33 | 1.00 | 15.33 | 3.33 | 3.33 |
| | | | 8 | 5.33 | 11.67 | 4.67 | 26.00 | 22.67 | 4.67 | 14.00 | 9.33 | 13.67 | 6.33 | 1.00 | 8.33 | 10.33 | 3.00 |

| Female's Geno | Rep | Start Date (YYMMDD) | vial | Days 15 | 16 | 17 | 18 | 19 | 20 | 21 | 22 | 23 | 24 | 25 | 26 | 27 | 28 |
|---|---|---|---|---|---|---|---|---|---|---|---|---|---|---|---|---|---|
| *DJ694 $^{L}$ /+* | 1 | 24-09-28 | 1 | 10.33 | 3.00 | 9.67 | 9.67 | 6.00 | 17.33 | 2.33 | 10.00 | 1.33 | 5.00 | 3.00 | 10.33 | 3.67 | 8.67 |
| | | | 2 | 6.00 | 7.00 | 6.67 | 6.00 | 3.00 | 9.67 | 4.67 | 9.33 | 1.00 | 5.33 | 7.33 | 1.33 | 5.00 | 1.67 |
| | | | 3 | 4.00 | 3.00 | 6.33 | 8.00 | 3.67 | 4.00 | 9.67 | 7.33 | 1.33 | 3.33 | 1.00 | 9.67 | 4.33 | 7.00 |
| | | | 4 | 3.67 | 3.33 | 1.00 | 16.33 | 4.00 | 8.33 | 9.67 | 10.00 | 0.67 | 4.00 | 3.33 | 10.00 | 2.00 | 5.00 |
| | | | 5 | 3.00 | 12.67 | 1.00 | 8.00 | 9.00 | 11.33 | 1.33 | 4.33 | 4.67 | 1.00 | 6.67 | 5.00 | 3.33 | 6.00 |
| | | | 6 | 5.00 | 1.67 | 11.33 | 7.00 | 1.67 | 14.67 | 2.00 | 3.00 | 3.33 | 2.33 | 12.00 | 3.00 | 7.00 | 6.67 |
| | | | 7 | 7.33 | 6.33 | 3.67 | 9.00 | 11.67 | 4.33 | 12.33 | 2.00 | 6.67 | 3.00 | 6.33 | 3.33 | 4.33 | 5.00 |
| | | | 8 | 6.33 | 11.00 | 8.67 | 5.00 | 4.67 | 16.33 | 3.33 | 8.33 | 5.50 | 0.67 | 6.33 | 4.33 | 1.67 | 8.33 |
| | 2 | | 1 | 3.00 | 7.33 | 6.67 | 5.67 | 7.00 | 11.67 | 2.00 | 10.00 | 1.67 | 2.33 | 6.67 | 2.00 | 2.67 | 1.67 |
| | | | 2 | 3.33 | 5.00 | 3.67 | 2.67 | 2.67 | 16.67 | 3.00 | 7.67 | 5.33 | 3.33 | 8.33 | 0.67 | 5.33 | 9.67 |
| | | | 3 | 9.67 | 12.67 | 1.33 | 6.33 | 4.33 | 9.33 | 7.67 | 2.00 | 7.00 | 5.33 | 6.00 | 6.33 | 4.00 | 4.33 |
| | | | 4 | 2.67 | 11.00 | 2.00 | 8.00 | 7.67 | 9.67 | 6.67 | 4.00 | 8.33 | 3.67 | 4.33 | 5.00 | 3.00 | 7.00 |
| | | | 5 | 9.67 | 1.67 | 6.67 | 6.00 | 4.67 | 12.67 | 1.00 | 6.67 | 1.33 | 1.67 | 5.00 | 0.67 | 3.67 | 8.67 |
| | | | 6 | 4.33 | 6.33 | 10.67 | 7.00 | 8.67 | 6.33 | 11.67 | 2.67 | 10.00 | 2.33 | 4.33 | 2.33 | 3.67 | 6.00 |
| | | | 7 | 12.33 | 6.67 | 2.67 | 1.67 | 2.67 | 14.67 | 2.67 | 7.33 | 3.33 | 1.67 | 6.67 | 2.00 | 3.33 | 7.00 |
| | | | 8 | 10.33 | 5.67 | 8.67 | 2.00 | 1.67 | 13.00 | 8.00 | 5.00 | 3.67 | 1.00 | 5.33 | 1.00 | 3.00 | 6.33 |

| Female's Geno | Rep | Start Date (YYMMDD) | vial | 29 | 30 | 31 | 32 | 33 | 34 | 35 | 36 | 37 | 38 | 39 | 40 |
|---|---|---|---|---|---|---|---|---|---|---|---|---|---|---|---|
| *DJ694$^{L}$/+* | 1 | 24-09-28 | 1 | 3.00 | 5.33 | 3.00 | 5.67 | 10.00 | 2.00 | 6.33 | 4.00 | 1.67 | 1.33 | 7.00 | 1.67 |
| | | | 2 | 3.67 | 7.67 | 3.67 | 2.67 | 4.67 | 8.33 | 6.00 | 3.33 | 3.00 | 3.67 | 5.00 | 6.67 |
| | | | 3 | 2.33 | 1.67 | 10.67 | 6.67 | 6.33 | 4.67 | 7.67 | 5.33 | 1.33 | 1.33 | 6.67 | 7.00 |
| | | | 4 | 2.67 | 3.67 | 7.33 | 5.67 | 5.00 | 6.67 | 6.67 | 5.00 | 6.00 | 3.67 | 4.00 | 4.33 |
| | | | 5 | 7.33 | 1.67 | 4.67 | 5.00 | 8.00 | 7.33 | 10.67 | 5.67 | 3.33 | 1.67 | 6.67 | 1.67 |
| | | | 6 | 2.67 | 7.00 | 5.67 | 3.00 | 6.33 | 6.00 | 4.67 | 6.67 | 2.67 | 6.33 | 4.33 | 3.33 |
| | | | 7 | 5.67 | 4.33 | 3.67 | 6.00 | 7.33 | 6.33 | 3.33 | 5.67 | 2.00 | 2.00 | 5.00 | 0.33 |
| | | | 8 | 5.67 | 1.00 | 2.33 | 11.33 | 3.67 | 4.67 | 5.67 | 1.00 | 3.67 | 0.33 | 0.67 | 1.33 |
| | 2 | | 1 | 1.67 | 1.33 | 7.33 | 4.33 | 1.33 | 3.33 | 1.00 | 3.33 | 2.33 | 6.00 | 3.33 | 3.00 |
| | | | 2 | 3.67 | 3.33 | 3.33 | 5.00 | 0.67 | 1.67 | 2.00 | 1.00 | 1.00 | 0.33 | 3.33 | 0.67 |
| | | | 3 | 7.33 | 1.00 | 4.00 | 10.00 | 1.67 | 6.33 | 3.33 | 4.00 | 2.00 | 8.00 | 3.67 | 4.00 |
| | | | 4 | 1.67 | 0.67 | 6.67 | 4.67 | 2.67 | 4.67 | 5.33 | 5.00 | 4.00 | 4.00 | 3.00 | 3.67 |
| | | | 5 | 5.33 | 4.00 | 5.67 | 4.00 | 4.33 | 8.33 | 4.33 | 3.33 | 7.67 | 7.00 | 7.33 | 2.00 |
| | | | 6 | 3.33 | 4.00 | 11.67 | 5.33 | 1.67 | 5.33 | 3.67 | 4.00 | 3.67 | 2.67 | 6.33 | 4.33 |
| | | | 7 | 5.33 | 4.33 | 3.67 | 5.00 | 2.00 | 9.00 | 1.33 | 6.67 | 3.00 | 1.00 | 5.67 | 4.33 |
| | | | 8 | 3.67 | 1.00 | 5.33 | 8.00 | 3.67 | 4.33 | 5.33 | 4.33 | 7.00 | 1.33 | 4.67 | 2.33 |

| *Female's Geno* | Rep | Start Date (YYMMDD) | vial | Days 1 | 2 | 3 | 4 | 5 | 6 | 7 | 8 | 9 | 10 | 11 | 12 | 13 | 14 |
|---|---|---|---|---|---|---|---|---|---|---|---|---|---|---|---|---|---|
| *UAS-EDTP$^E$/+* | 1 | 24-09-28 | 1 | 11.67 | 1.67 | 25.67 | 22.67 | 23.67 | 10.33 | 21.67 | 13.00 | 9.67 | 11.00 | 6.33 | 8.67 | 5.67 | 5.67 |
| | | | 2 | 7.33 | 0.67 | 50.33 | 24.67 | 21.00 | 12.33 | 19.00 | 7.67 | 8.00 | 11.00 | 4.00 | 8.00 | 6.33 | 4.00 |
| | | | 3 | 17.67 | 11.67 | 15.67 | 17.33 | 11.67 | 16.00 | 20.00 | 12.67 | 9.67 | 12.33 | 4.00 | 11.00 | 9.67 | 5.00 |
| | | | 4 | 7.33 | 10.33 | 30.00 | 22.67 | 17.00 | 14.67 | 10.33 | 11.67 | 3.67 | 16.33 | 4.33 | 15.00 | 2.67 | 13.33 |
| | | | 5 | 4.67 | 1.67 | 39.00 | 21.33 | 14.00 | 11.33 | 15.33 | 13.67 | 4.00 | 11.33 | 5.00 | 6.67 | 6.00 | 4.33 |
| | | | 6 | 22.67 | 14.33 | 7.33 | 19.33 | 13.33 | 8.00 | 16.00 | 10.33 | 5.67 | 13.00 | 4.67 | 13.00 | 9.33 | 6.67 |
| | | | 7 | 15.00 | 6.67 | 17.33 | 17.00 | 14.00 | 14.67 | 14.33 | 11.33 | 9.00 | 12.67 | 1.33 | 4.00 | 8.00 | 1.00 |
| | | | 8 | 18.00 | 9.67 | 21.33 | 14.33 | 10.33 | 10.33 | 23.00 | 5.33 | 10.33 | 6.33 | 11.00 | 5.33 | 0.67 | 22.33 |
| | 2 | | 1 | 15.00 | 7.33 | 4.67 | 28.67 | 22.00 | 10.67 | 17.67 | 14.00 | 5.67 | 10.00 | 3.33 | 6.67 | 9.00 | 4.67 |
| | | | 2 | 24.67 | 6.33 | 13.00 | 33.67 | 22.67 | 11.00 | 14.33 | 9.33 | 11.67 | 5.00 | 5.00 | 11.67 | 4.67 | 2.67 |
| | | | 3 | 21.00 | 12.67 | 13.33 | 19.67 | 22.00 | 12.00 | 5.67 | 6.00 | 11.67 | 11.67 | 12.67 | 3.67 | 6.67 | 7.67 |
| | | | 4 | 6.67 | 2.33 | 17.00 | 11.33 | 14.00 | 11.33 | 6.67 | 11.33 | 10.33 | 13.33 | 1.00 | 9.67 | 6.67 | 2.67 |
| | | | 5 | 22.00 | 4.00 | 8.00 | 18.67 | 14.33 | 10.00 | 14.33 | 14.33 | 5.00 | 17.33 | 4.33 | 8.67 | 5.33 | 10.00 |
| | | | 6 | 7.67 | 11.33 | 6.33 | 19.33 | 12.67 | 4.67 | 18.67 | 14.67 | 9.00 | 10.00 | 4.00 | 5.00 | 6.00 | 6.67 |
| | | | 7 | 6.33 | 9.33 | 15.00 | 24.33 | 22.00 | 11.33 | 13.00 | 13.33 | 8.33 | 9.00 | 4.67 | 10.00 | 3.00 | 4.67 |
| | | | 8 | 10.00 | 3.33 | 18.33 | 17.00 | 19.00 | 19.67 | 11.67 | 13.67 | 7.00 | 9.00 | 3.67 | 6.67 | 7.00 | 13.00 |

| Female's Geno | Rep | Start Date (YYMMDD) | vial | 15 | 16 | 17 | 18 | 19 | 20 | 21 | 22 | 23 | 24 | 25 | 26 | 27 | 28 |
|---|---|---|---|---|---|---|---|---|---|---|---|---|---|---|---|---|---|
| *UAS-EDTP^E/+* | 1 | 24-09-28 | 1 | 7.33 | 8.00 | 7.67 | 3.67 | 10.33 | 4.00 | 8.33 | 6.67 | 3.00 | 8.00 | 5.67 | 1.00 | 6.67 | 7.67 |
| | | | 2 | 5.00 | 9.33 | 5.00 | 2.00 | 1.00 | 11.00 | 3.33 | 11.67 | 1.67 | 4.33 | 6.33 | 5.00 | 1.33 | 4.67 |
| | | | 3 | 5.33 | 4.33 | 4.67 | 1.33 | 14.00 | 2.00 | 6.33 | 5.00 | 5.67 | 6.67 | 1.33 | 9.00 | 3.00 | 9.67 |
| | | | 4 | 1.33 | 6.67 | 1.00 | 4.67 | 10.00 | 3.67 | 9.00 | 10.67 | 6.00 | 1.33 | 5.67 | 10.33 | 1.33 | 7.00 |
| | | | 5 | 7.67 | 4.67 | 4.67 | 4.33 | 11.33 | 10.00 | 1.00 | 7.33 | 6.00 | 2.00 | 7.67 | 3.33 | 0.67 | 3.33 |
| | | | 6 | 5.67 | 3.33 | 1.67 | 2.00 | 5.00 | 8.67 | 3.33 | 10.00 | 4.67 | 0.67 | 2.00 | 9.67 | 2.33 | 2.33 |
| | | | 7 | 10.00 | 2.67 | 0.00 | 2.00 | 12.67 | 5.33 | 5.67 | 8.33 | 6.00 | 2.00 | 1.00 | 3.00 | 1.67 | 3.33 |
| | | | 8 | 0.67 | 0.67 | 1.00 | 9.00 | 6.67 | 19.67 | 6.67 | 5.67 | 4.33 | 4.33 | 4.00 | 0.33 | 2.00 | 5.33 |
| | 2 | | 1 | 6.00 | 2.33 | 5.00 | 7.67 | 1.00 | 8.33 | 6.00 | 6.67 | 0.33 | 4.67 | 3.33 | 1.00 | 4.00 | 3.00 |
| | | | 2 | 9.33 | 3.00 | 7.00 | 8.67 | 3.00 | 13.33 | 2.67 | 6.33 | 0.67 | 5.67 | 7.33 | 2.00 | 4.00 | 8.67 |
| | | | 3 | 6.67 | 3.67 | 9.33 | 1.00 | 3.00 | 11.67 | 9.67 | 3.33 | 2.33 | 5.00 | 4.67 | 4.67 | 3.00 | 10.67 |
| | | | 4 | 0.33 | 7.33 | 5.33 | 9.67 | 2.00 | 11.33 | 7.00 | 8.67 | 3.33 | 2.67 | 11.67 | 8.33 | 4.67 | 4.67 |
| | | | 5 | 1.00 | 8.00 | 0.33 | 11.33 | 1.33 | 15.00 | 1.67 | 3.00 | 1.00 | 8.00 | 2.00 | 6.00 | 3.67 | 9.00 |
| | | | 6 | 3.67 | 4.67 | 2.33 | 5.33 | 2.67 | 11.00 | 2.00 | 8.67 | 1.00 | 3.33 | 7.33 | 2.33 | 3.00 | 11.33 |
| | | | 7 | 0.33 | 7.00 | 2.00 | 7.67 | 5.00 | 13.00 | 1.00 | 6.33 | 3.33 | 5.00 | 5.33 | 1.67 | 4.33 | 0.67 |
| | | | 8 | 5.67 | 6.33 | 8.00 | 6.67 | 8.33 | 3.67 | 1.33 | 6.00 | 4.33 | 3.33 | 6.33 | 3.67 | 2.33 | 8.33 |

| Female's Geno | Rep | Start Date (YYMMDD) | vial | 29 | 30 | 31 | 32 | 33 | 34 | 35 | 36 | 37 | 38 | 39 | 40 |
|---|---|---|---|---|---|---|---|---|---|---|---|---|---|---|---|
| *UAS-EDTP $^{E}$ /+* | 1 | 24-09-28 | 1 | 10.00 | 8.00 | 4.00 | 13.00 | 5.33 | 0.67 | 1.67 | 10.33 | 1.33 | 5.33 | 3.67 | 6.67 |
| | | | 2 | 5.00 | 4.67 | 2.67 | 5.00 | 6.00 | 5.00 | 5.67 | 5.00 | 5.00 | 9.33 | 6.33 | 3.33 |
| | | | 3 | 2.67 | 9.67 | 3.67 | 8.00 | 5.33 | 8.33 | 5.67 | 9.33 | 8.33 | 4.67 | 3.33 | 5.00 |
| | | | 4 | 3.33 | 11.33 | 1.33 | 3.33 | 8.33 | 15.00 | 6.33 | 4.33 | 8.67 | 3.67 | 9.33 | 4.67 |
| | | | 5 | 6.00 | 11.00 | 4.33 | 6.67 | 2.67 | 6.33 | 8.33 | 13.67 | 3.67 | 5.00 | 9.33 | 4.00 |
| | | | 6 | 1.67 | 1.00 | 10.67 | 4.00 | 4.33 | 7.00 | 7.50 | 1.00 | 2.00 | 1.67 | 5.00 | 0.67 |
| | | | 7 | 5.00 | 2.33 | 10.67 | 10.33 | 3.00 | 5.00 | 6.00 | 7.33 | 1.33 | 4.67 | 5.00 | 5.67 |
| | | | 8 | 4.67 | 2.33 | 4.33 | 4.67 | 4.00 | 3.33 | 7.67 | 0.67 | 2.00 | 2.00 | 10.00 | 2.00 |
| | 2 | | 1 | 4.33 | 10.00 | 6.67 | 4.00 | 4.00 | 6.00 | 4.67 | 4.00 | 2.33 | 9.33 | 2.33 | 2.00 |
| | | | 2 | 1.00 | 7.00 | 4.33 | 5.33 | 3.67 | 3.33 | 5.00 | 3.67 | 1.67 | 7.67 | 4.00 | 8.67 |
| | | | 3 | 6.00 | 1.00 | 6.33 | 8.33 | 2.67 | 7.00 | 5.33 | 9.33 | 3.33 | 11.67 | 4.00 | 4.33 |
| | | | 4 | 9.67 | 4.33 | 6.67 | 3.00 | 8.33 | 4.67 | 4.67 | 6.67 | 1.33 | 5.33 | 9.00 | 5.67 |
| | | | 5 | 9.00 | 6.00 | 2.67 | 5.33 | 6.67 | 4.00 | 10.33 | 4.67 | 6.33 | 8.00 | 5.00 | 1.33 |
| | | | 6 | 6.33 | 8.00 | 5.00 | 6.00 | 4.67 | 5.67 | 5.33 | 6.67 | 7.00 | 2.67 | 7.00 | 3.33 |
| | | | 7 | 2.67 | 2.00 | 6.00 | 6.67 | 1.33 | 3.67 | 4.00 | 8.00 | 1.33 | 2.33 | 7.00 | 4.67 |
| | | | 8 | 5.67 | 11.00 | 1.67 | 10.33 | 1.33 | 7.33 | 9.00 | 8.33 | 8.00 | 9.00 | 11.33 | 5.33 |

Egg-laying was quantified as the number of eggs laid per female per day. For each genotype, multiple biological replicates were scored, with each replicate consisting of multiple vials (8 vials per genotype, 3 female in each vial) that were monitored over the assay period. All female are mated with *w* males.

The "Start Date (YYMMDD)" column indicates the date on which egg laying began for each replicate. An "x" indicates that all females within the vial had died.

| Exp | Date yymmdd | Genotype | Sex | n | Mean | SD | Med | Max | %↓DJ694 mean | %↓DJ694 P | %↓DJ694 Δ | %↑Rescue mean | %↑Rescue P(hom) | %↑Rescue P(het) | %↑Rescue Δ |
|---|---|---|---|---|---|---|---|---|---|---|---|---|---|---|---|
| 1 | 00-06-05 | *DJ694* | ♀ | 475 | 37.02 | 2.93 | 39.00 | 48.47 | - | - | - | - | - | - | - |
| 1 | 00-06-05 | $w^{1118}$ | ♀ | 548 | 52.82 | 5.76 | 56.00 | 72.00 | -15.11 | <2E-16 | ↓ | - | - | - | - |
| 1 | 00-06-05 | *DJ694* | ♂ | 512 | 34.90 | 3.26 | 35.00 | 43.06 | - | - | - | - | - | - | - |
| 1 | 00-06-05 | $w^{1118}$ | ♂ | 576 | 44.21 | 3.95 | 42.00 | 59.88 | -5.19 | <2E-16 | ↓ | - | - | - | - |
| 2 | 04-12-01 | *DJ694* | ♀ | 123 | 53.12 | 2.26 | 55.00 | 60.00 | - | - | - | - | - | - | - |
| 2 | 04-12-01 | *DJ694/+* | ♀ | 103 | 74.87 | 5.53 | 79.00 | 91.50 | -20.12 | <2E-16 | ↓ | - | - | - | - |
| 2 | 04-12-01 | *DJ694;UAS-EDTP$^A$/+* | ♀ | 74 | 68.32 | 1.67 | 72.00 | 79.25 | - | - | - | 20.34 | <2E-16 | 2E-09 | ↑ |
| 2 | 04-12-01 | *DJ694;UAS-EDTP$^B$/+* | ♀ | 99 | 72.44 | 4.57 | 72.00 | 82.00 | - | - | - | 22.54 | <2E-16 | 6E-05 | ↑ |
| 2 | 04-12-01 | *DJ694;UAS-EDTP$^A$/UAS-EDTP$^B$* | ♀ | 115 | 71.84 | 4.27 | 76.00 | 83.75 | - | - | - | 22.01 | <2E-16 | 4E-05 | ↑ |
| 2 | 04-12-01 | *DJ694* | ♂ | 131 | 51.65 | 1.07 | 51.00 | 56.50 | - | - | - | - | - | - | - |
| 2 | 04-12-01 | *DJ694/+* | ♂ | 94 | 70.09 | 4.54 | 69.00 | 78.50 | -19.58 | <2E-16 | ↓ | - | - | - | - |
| 2 | 04-12-01 | *DJ694;UAS-EDTP$^A$/+* | ♂ | 84 | 56.68 | 2.14 | 60.00 | 69.00 | - | - | - | 3.47 | 5E-12 | <2E-16 | ↑ |
| 2 | 04-12-01 | *DJ694;UAS-EDTP$^B$/+* | ♂ | 86 | 70.53 | 3.62 | 72.00 | 81.25 | - | - | - | 26.93 | <2E-16 | 0.3 | ↑ |
| 2 | 04-12-01 | *DJ694;UAS-EDTP$^A$/UAS-EDTP$^B$* | ♂ | 108 | 62.75 | 4.14 | 69.00 | 75.75 | - | - | - | 11.19 | <2E-16 | 1E-05 | ↑ |
| 3 | 04-12-04 | *DJ694* | ♀ | 110 | 44.87 | 1.67 | 45.00 | 53.25 | - | - | - | - | - | - | - |
| 3 | 04-12-04 | *DJ694/+* | ♀ | 112 | 74.08 | 8.22 | 80.00 | 88.75 | -29.35 | <2E-16 | ↓ | - | - | - | - |
| 3 | 04-12-04 | *DJ694;UAS-EDTP$^A$/+* | ♀ | 109 | 58.75 | 3.50 | 59.00 | 72.00 | - | - | - | 18.74 | <2E-16 | <2E-16 | ↑ |
| 3 | 04-12-04 | *DJ694;UAS-EDTP$^B$/+* | ♀ | 155 | 54.16 | 4.96 | 55.00 | 68.50 | - | - | - | 5.74 | 4E-16 | <2E-16 | ↑ |
| 3 | 04-12-04 | *DJ694;UAS-EDTP$^A$/UAS-EDTP$^B$* | ♀ | 130 | 59.91 | 5.86 | 59.00 | 73.50 | - | - | - | 16.14 | <2E-16 | <2E-16 | ↑ |
| 3 | 04-12-04 | *DJ694* | ♂ | 104 | 50.06 | 2.16 | 52.00 | 56.00 | - | - | - | - | - | - | - |
| 3 | 04-12-04 | *DJ694/+* | ♂ | 96 | 66.93 | 3.60 | 69.00 | 79.00 | -17.53 | <2E-16 | ↓ | - | - | - | - |
| 3 | 04-12-04 | *DJ694;UAS-EDTP$^A$/+* | ♂ | 86 | 54.50 | 4.14 | 55.00 | 65.00 | - | - | - | 0 | 2E-08 | <2E-16 | X |
| 3 | 04-12-04 | *DJ694;UAS-EDTP$^B$/+* | ♂ | 145 | 62.05 | 3.46 | 66.00 | 72.00 | - | - | - | 12.19 | <2E-16 | 2E-08 | ↑ |
| 3 | 04-12-04 | *DJ694;UAS-EDTP$^A$/UAS-EDTP$^B$* | ♂ | 148 | 59.79 | 3.55 | 59.00 | 75.50 | - | - | - | 7.69 | <2E-16 | 2E-07 | ↑ |
| 4 | 04-12-01 | *DJ694* | ♀ | 120 | 54.23 | 1.67 | 55.00 | 59.00 | - | - | - | - | - | - | - |
| 4 | 04-12-01 | *DJ694/+* | ♀ | 86 | 76.84 | 8.22 | 81.00 | 90.75 | -18.53 | <2E-16 | ↓ | - | - | - | - |
| 4 | 04-12-01 | *DJ694;UAS-EDTP$^A$/+* | ♀ | 123 | 64.71 | 3.51 | 65.00 | 77.75 | - | - | - | 9.48 | <2E-16 | <2E-16 | ↑ |
| 4 | 04-12-01 | *DJ694;UAS-EDTP$^B$/+* | ♀ | 95 | 69.92 | 4.96 | 72.00 | 81.75 | - | - | - | 16.21 | <2E-16 | 1E-08 | ↑ |
| 4 | 04-12-01 | *DJ694;UAS-EDTP$^A$/UAS-EDTP$^B$* | ♀ | 113 | 74.60 | 5.87 | 76.00 | 84.75 | - | - | - | 22.95 | <2E-16 | 0.001 | ↑ |
| 4 | 04-12-01 | *DJ694* | ♂ | 118 | 53.30 | 2.17 | 55.00 | 59.00 | - | - | - | - | - | - | - |
| 4 | 04-12-01 | *DJ694/+* | ♂ | 84 | 65.03 | 3.62 | 69.00 | 81.75 | -9.68 | <2E-16 | ↓ | - | - | - | - |
| 4 | 04-12-01 | *DJ694;UAS-EDTP$^A$/+* | ♂ | 100 | 57.83 | 4.14 | 58.00 | 69.75 | - | - | - | 0 | 1E-08 | 1E-08 | X |
| 4 | 04-12-01 | *DJ694;UAS-EDTP$^B$/+* | ♂ | 82 | 66.98 | 4.92 | 69.00 | 77.75 | - | - | - | 11.90 | <2E-16 | 0.3 | ↑ |
| 4 | 04-12-01 | *DJ694;UAS-EDTP$^A$/UAS-EDTP$^B$* | ♂ | 79 | 65.36 | 3.55 | 69.00 | 75.00 | - | - | - | 11.43 | <2E-16 | 0.5 | ↑ |
| 5 | 04-12-04 | *DJ694* | ♀ | 127 | 46.32 | 0.83 | 48.00 | 57.00 | - | - | - | - | - | - | - |
| 5 | 04-12-04 | *DJ694/+* | ♀ | 121 | 55.86 | 7.86 | 59.00 | 78.25 | -1.78 | 3E-11 | ↓ | - | - | - | - |
| 5 | 04-12-04 | *DJ694;UAS-EDTP$^A$/+* | ♀ | 129 | 54.66 | 2.94 | 59.00 | 68.50 | - | - | - | 9.69 | 1E-13 | 0.003 | ↑ |
| 5 | 04-12-04 | *DJ694;UAS-EDTP$^B$/+* | ♀ | 129 | 55.97 | 5.07 | 55.00 | 72.00 | - | - | - | 7.95 | 8E-15 | 0.04 | ↑ |
| 5 | 04-12-04 | *DJ694;UAS-EDTP$^A$/UAS-EDTP$^B$* | ♀ | 140 | 65.97 | 5.95 | 66.00 | 79.75 | - | - | - | 27.31 | <2E-16 | 4E-04 | ↑ |
| 5 | 04-12-04 | *DJ694* | ♂ | 112 | 47.25 | 2.05 | 48.00 | 55.00 | - | - | - | - | - | - | - |
| 5 | 04-12-04 | *DJ694/+* | ♂ | 122 | 57.15 | 3.78 | 59.00 | 76.50 | -7.63 | 3E-16 | ↓ | - | - | - | - |
| 5 | 04-12-04 | *DJ694;UAS-EDTP$^A$/+* | ♂ | 110 | 58.38 | 4.23 | 59.00 | 68.50 | - | - | - | 9.85 | <2E-16 | 0.4 | ↑ |
| 5 | 04-12-04 | *DJ694;UAS-EDTP$^B$/+* | ♂ | 114 | 63.63 | 2.94 | 66.00 | 77.25 | - | - | - | 23.11 | <2E-16 | 6E-04 | ↑ |

| Exp | Date yymmdd | Genotype | Sex | n | Mean | SD | Med | Max | %↓DJ694 mean | %↓DJ694 P | %↓DJ694 Δ | %↑Rescue mean | %↑Rescue P(hom) | %↑Rescue P(het) | %↑Rescue Δ |
|---|---|---|---|---|---|---|---|---|---|---|---|---|---|---|---|
| 5 | 04-12-04 | *DJ694;UAS-EDTP*$^{A}$/*UAS-EDTP*$^{B}$ | ♂ | 122 | 59.97 | 4.91 | 62.00 | 76.50 | - | - | - | 11.68 | <2E-16 | 0.1 | ↑ |
| 6 | 04-08-08 | *DJ694* | ♀ | 113 | 46.27 | 1.55 | 48.00 | 56.50 | - | - | - | - | - | - | - |
| 6 | 04-08-08 | $w^{1118}$ | ♀ | 104 | 66.34 | 9.48 | 73.00 | 84.25 | -15.90 | <2E-16 | ↓ | - | - | - | - |
| 6 | 04-08-08 | *DJ694;UAS-EDTP*$^{A}$/+ | ♀ | 102 | 69.46 | 3.01 | 73.00 | 78.25 | - | - | - | 38.97 | <2E-16 | - | ↑ |
| 6 | 04-08-08 | *DJ694;UAS-EDTP*$^{B}$/+ | ♀ | 110 | 69.48 | 5.16 | 73.00 | 83.50 | - | - | - | 34.51 | <2E-16 | - | ↑ |
| 6 | 04-08-08 | *DJ694* | ♂ | 107 | 48.44 | 0.86 | 52.00 | 55.75 | - | - | - | - | - | - | - |
| 6 | 04-08-08 | $w^{1118}$ | ♂ | 43 | 59.49 | 2.81 | 52.00 | 66.25 | -13.01 | 4E-13 | ↓ | - | - | - | - |
| 6 | 04-08-08 | *DJ694;UAS-EDTP*$^{A}$/+ | ♂ | 103 | 74.11 | 1.96 | 76.00 | 83.50 | - | - | - | 46.35 | <2E-16 | - | ↑ |
| 6 | 04-08-08 | *DJ694;UAS-EDTP*$^{B}$/+ | ♂ | 99 | 66.05 | 5.85 | 69.00 | 76.75 | - | - | - | 22.10 | <2E-16 | - | ↑ |
| 7 | 04-08-12 | *DJ694* | ♀ | 97 | 45.92 | 1.05 | 48.00 | 54.00 | - | - | - | - | - | - | - |
| 7 | 04-08-12 | $w^{1118}$ | ♀ | 153 | 66.07 | 4.04 | 69.00 | 81.75 | -24.27 | <2E-16 | ↓ | - | - | - | - |
| 7 | 04-08-12 | *DJ694;UAS-EDTP*$^{A}$/+ | ♀ | 124 | 70.03 | 3.72 | 72.00 | 77.25 | - | - | - | 41.17 | <2E-16 | - | ↑ |
| 7 | 04-08-12 | *DJ694;UAS-EDTP*$^{B}$/+ | ♀ | 131 | 70.24 | 2.44 | 75.00 | 83.25 | - | - | - | 44.35 | <2E-16 | - | ↑ |
| 7 | 04-08-12 | *DJ694* | ♂ | 107 | 49.73 | 0.83 | 51.00 | 55.50 | - | - | - | - | - | - | - |
| 7 | 04-08-12 | $w^{1118}$ | ♂ | 120 | 58.16 | 3.96 | 61.00 | 75.00 | -6.73 | 4E-16 | ↓ | - | - | - | - |
| 7 | 04-08-12 | *DJ694;UAS-EDTP*$^{A}$/+ | ♂ | 111 | 69.13 | 2.68 | 72.00 | 81.00 | - | - | - | 31.44 | <2E-16 | - | ↑ |
| 7 | 04-08-12 | *DJ694;UAS-EDTP*$^{B}$/+ | ♂ | 98 | 66.14 | 3.03 | 67.00 | 75.75 | - | - | - | 24.84 | <2E-16 | - | ↑ |
| 8 | 04-08-15 | *DJ694* | ♀ | 83 | 44.70 | 2.77 | 48.00 | 53.00 | - | - | - | - | - | - | - |
| 8 | 04-08-15 | $w^{1118}$ | ♀ | 108 | 58.84 | 3.71 | 62.00 | 72.75 | -13.89 | <2E-16 | ↓ | - | - | - | - |
| 8 | 04-08-15 | *DJ694;UAS-EDTP*$^{A}$/+ | ♀ | 127 | 59.31 | 6.27 | 62.00 | 71.00 | - | - | - | 11.74 | <2E-16 | - | ↑ |
| 8 | 04-08-15 | *DJ694;UAS-EDTP*$^{B}$/+ | ♀ | 137 | 64.96 | 4.12 | 66.00 | 76.50 | - | - | - | 28.18 | <2E-16 | - | ↑ |
| 8 | 04-08-15 | *DJ694* | ♂ | 74 | 46.05 | 2.46 | 48.00 | 53.00 | - | - | - | - | - | - | - |
| 8 | 04-08-15 | $w^{1118}$ | ♂ | 104 | 57.58 | 2.86 | 58.00 | 72.75 | -11.37 | <2E-16 | ↓ | - | - | - | - |
| 8 | 04-08-15 | *DJ694;UAS-EDTP*$^{A}$/+ | ♂ | 117 | 65.43 | 2.67 | 69.00 | 75.75 | - | - | - | 29.38 | <2E-16 | - | ↑ |
| 8 | 04-08-15 | *DJ694;UAS-EDTP*$^{B}$/+ | ♂ | 129 | 64.12 | 4.43 | 66.00 | 75.75 | - | - | - | 23.08 | <2E-16 | - | ↑ |
| 9 | 04-08-17 | *DJ694* | ♀ | 93 | 36.21 | 5.65 | 36.00 | 52.33 | - | - | - | - | - | - | - |
| 9 | 04-08-17 | $w^{1118}$ | ♀ | 104 | 70.22 | 3.53 | 70.00 | 84.00 | -37.23 | <2E-16 | ↓ | - | - | - | - |
| 9 | 04-08-17 | *DJ694;UAS-EDTP*$^{A}$/+ | ♀ | 73 | 64.76 | 4.18 | 67.00 | 74.00 | - | - | - | 44.74 | <2E-16 | - | ↑ |
| 9 | 04-08-17 | *DJ694;UAS-EDTP*$^{B}$/+ | ♀ | 39 | 62.51 | 8.94 | 64.00 | 83.00 | - | - | - | 27.99 | <2E-16 | - | ↑ |
| 9 | 04-08-17 | *DJ694* | ♂ | 68 | 37.98 | 0.23 | 43.00 | 54.00 | - | - | - | - | - | - | - |
| 9 | 04-08-17 | $w^{1118}$ | ♂ | 107 | 56.56 | 2.09 | 56.00 | 73.75 | -29.86 | <2E-16 | ↓ | - | - | - | - |
| 9 | 04-08-17 | *DJ694;UAS-EDTP*$^{A}$/+ | ♂ | 64 | 69.53 | 1.67 | 70.00 | 79.00 | - | - | - | 77.59 | <2E-16 | - | ↑ |
| 9 | 04-08-17 | *DJ694;UAS-EDTP*$^{B}$/+ | ♂ | 34 | 63.98 | 3.46 | 67.00 | 73.00 | - | - | - | 58.39 | 5E-16 | - | ↑ |
| 10 | 03-09-22 | *DJ694* | ♀ | 53 | 50.01 | 3.71 | 51.57 | 55.25 | - | - | - | - | - | - | - |
| 10 | 03-09-22 | $w^{1118}$ | ♀ | 110 | 67.79 | 8.31 | 68.47 | 77.14 | -9.70 | <2E-16 | ↓ | - | - | - | - |
| 11 | 03-09-22 | *DJ694* | ♀ | 65 | 58.67 | 1.14 | 58.39 | 64.25 | - | - | - | - | - | - | - |
| 11 | 03-09-22 | $w^{1118}$ | ♀ | 71 | 69.82 | 7.72 | 70.68 | 81.50 | -3.70 | <2E-16 | ↓ | - | - | - | - |
| 11 | 03-09-22 | *DJ694* | ♂ | 68 | 48.98 | 1.21 | 47.39 | 55.50 | - | - | - | - | - | - | - |
| 11 | 03-09-22 | $w^{1118}$ | ♂ | 62 | 52.63 | 2.08 | 50.15 | 65.90 | -0.71 | 0.007 | ↓ | - | - | - | - |
| 12 | 03-09-25 | *DJ694* | ♀ | 40 | 49.68 | 8.17 | 49.79 | 57.50 | - | - | - | - | - | - | - |
| 12 | 03-09-25 | $w^{1118}$ | ♀ | 93 | 60.61 | 6.18 | 61.96 | 72.86 | 0 | 5E-10 | X | - | - | - | - |
| 12 | 03-09-25 | *DJ694* | ♂ | 49 | 39.55 | 6.97 | 35.05 | 51.95 | - | - | - | - | - | - | - |
| 12 | 03-09-25 | $w^{1118}$ | ♂ | 103 | 59.99 | 4.49 | 59.94 | 73.27 | -16.18 | <2E-16 | ↓ | - | - | - | - |

| Exp | Date yymmdd | Genotype | Sex | n | Mean | SD | Med | Max | %↓DJ694 mean | %↓DJ694 P | %↓DJ694 ∆ | %↑Rescue mean | %↑Rescue P(hom) | %↑Rescue P(het) | %↑Rescue ∆ |
|---|---|---|---|---|---|---|---|---|---|---|---|---|---|---|---|
| 13 | 03-09-25 | *DJ694* | ♀ | 53 | 50.58 | 5.23 | 48.88 | 60.51 | - | - | - | - | - | - | - |
| 13 | 03-09-25 | $w^{1118}$ | ♀ | 61 | 73.37 | 1.84 | 73.04 | 86.21 | -21.98 | <2E-16 | ↓ | - | - | - | - |
| 13 | 03-09-25 | *DJ694* | ♂ | 68 | 43.65 | 6.98 | 42.56 | 72.69 | - | - | - | - | - | - | - |
| 13 | 03-09-25 | $w^{1118}$ | ♂ | 55 | 63.90 | 0.80 | 62.67 | 77.20 | -19.76 | 3E-16 | ↓ | - | - | - | - |
| 14 | 04-03-14 | *DJ694* | ♀ | 76 | 60.64 | 2.28 | 59.69 | 73.98 | - | - | - | - | - | - | - |
| 14 | 04-03-14 | $w^{1118}$ | ♀ | 77 | 81.28 | 1.68 | 79.70 | 87.19 | -20.96 | <2E-16 | ↓ | - | - | - | - |
| 14 | 04-03-14 | *DJ694/+* | ♀ | 71 | 71.07 | 6.42 | 74.17 | 84.07 | -2.68 | 1E-06 | ↓ | - | - | - | - |
| 14 | 04-03-14 | *DJ694* | ♂ | 93 | 52.25 | 3.65 | 51.02 | 59.31 | - | - | - | - | - | - | - |
| 14 | 04-03-14 | $w^{1118}$ | ♂ | 73 | 60.69 | 2.47 | 66.89 | 77.56 | -3.97 | 8E-11 | ↓ | - | - | - | - |
| 14 | 04-03-14 | *DJ694/+* | ♂ | 90 | 65.40 | 5.59 | 67.37 | 80.00 | -6.53 | <2E-16 | ↓ | - | - | - | - |
| 15 | 04-03-14 | *DJ694* | ♀ | 76 | 65.51 | 7.70 | 67.00 | 77.15 | - | - | - | - | - | - | - |
| 15 | 04-03-14 | $w^{1118}$ | ♀ | 69 | 81.44 | 2.59 | 82.67 | 86.98 | -7.16 | <2E-16 | ↓ | - | - | - | - |
| 15 | 04-03-14 | *DJ694/+* | ♀ | 79 | 71.80 | 4.89 | 78.33 | 84.91 | 0 | 4E-06 | X | - | - | - | - |
| 15 | 04-03-14 | *DJ694* | ♂ | 72 | 56.80 | 6.57 | 57.67 | 69.40 | - | - | - | - | - | - | - |
| 15 | 04-03-14 | $w^{1118}$ | ♂ | 76 | 58.88 | 3.99 | 59.00 | 69.92 | 0 | 0.9 | X | - | - | - | - |
| 15 | 04-03-14 | *DJ694/+* | ♂ | 77 | 72.32 | 7.74 | 78.33 | 86.92 | -1.87 | <2E-16 | ↓ | - | - | - | - |
| 16 | 04-03-18 | *DJ694* | ♀ | 132 | 60.45 | 2.49 | 59.00 | 75.67 | - | - | - | - | - | - | - |
| 16 | 04-03-18 | $w^{1118}$ | ♀ | 102 | 74.83 | 1.10 | 79.75 | 81.61 | -14.63 | <2E-16 | ↓ | - | - | - | - |
| 16 | 04-03-18 | *DJ694* | ♂ | 136 | 51.00 | 4.09 | 52.00 | 57.50 | - | - | - | - | - | - | - |
| 16 | 04-03-18 | $w^{1118}$ | ♂ | 115 | 60.68 | 6.63 | 62.00 | 73.94 | 0 | <2E-16 | X | - | - | - | - |
| 17 | 19-10-09 | *DJ694* | ♀ | 142 | 40.59 | 2.57 | 42.00 | 51.50 | - | - | - | - | - | - | - |
| 17 | 19-10-09 | *DJ694/+* | ♀ | 126 | 48.41 | 1.03 | 49.00 | 57.25 | -8.91 | 3E-13 | ↓ | - | - | - | - |
| 17 | 19-10-09 | *DJ694* | ♂ | 146 | 42.4 | 1.64 | 44.00 | 50.8 | - | - | - | - | - | - | - |
| 17 | 19-10-09 | *DJ694/+* | ♂ | 112 | 51.2 | 2.14 | 54.00 | 57.3 | -10.39 | <2E-16 | ↓ | - | - | - | - |
| 18 | 19-10-11 | *DJ694* | ♀ | 129 | 39.49 | 0.89 | 40.00 | 49.75 | - | - | - | - | - | - | - |
| 18 | 19-10-11 | *DJ694/+* | ♀ | 141 | 46.60 | 0.75 | 47.00 | 60.00 | -11.94 | 2E-13 | ↓ | - | - | - | - |
| 18 | 19-10-11 | *DJ694* | ♂ | 114 | 42.2 | 1.04 | 45.00 | 47.50 | - | - | - | - | - | - | - |
| 18 | 19-10-11 | *DJ694/+* | ♂ | 103 | 54.7 | 2.28 | 52.00 | 66 | -17.59 | <2E-16 | ↓ | - | - | - | - |
| 19 | 20-02-05 | *DJ694* | ♀ | 78 | 39.64 | 1.01 | 40.00 | 52.00 | - | - | - | - | - | - | - |
| 19 | 20-02-05 | *DJ694/+* | ♀ | 101 | 54.46 | 5.04 | 54.00 | 71.25 | -17.73 | 1E-16 | ↓ | - | - | - | - |
| 19 | 20-02-05 | *DJ694* | ♂ | 114 | 43.1 | 1.41 | 47.00 | 50.50 | - | - | - | - | - | - | - |
| 19 | 20-02-05 | *DJ694/+* | ♂ | 51 | 63.58 | 10.74 | 63.00 | 85.50 | -15.73 | <2E-16 | ↓ | - | - | - | - |
| 20 | 20-02-07 | *DJ694* | ♀ | 99 | 42.97 | 2.84 | 45.00 | 52.50 | - | - | - | - | - | - | - |
| 20 | 20-02-07 | *DJ694/+* | ♀ | 91 | 56.82 | 2.91 | 56.00 | 79.25 | -15.04 | 2E-16 | ↓ | - | - | - | - |
| 20 | 20-02-07 | *DJ694* | ♂ | 98 | 46.19 | 1.25 | 47.00 | 52.00 | - | - | - | - | - | - | - |
| 20 | 20-02-07 | *DJ694/+* | ♂ | 81 | 61.28 | 2.82 | 59.00 | 89.75 | -18.84 | <2E-16 | ↓ | - | - | - | - |
| 21 | 23-03-06 | $DJ694^{L}$ | ♀ | 54 | 45.06 | 7.79 | 44.75 | 65.30 | - | - | - | - | - | - | - |
| 21 | 23-03-06 | $w^{1118}$ | ♀ | 71 | 53.24 | 3.45 | 49.31 | 74.95 | 0 | 0.03 | X | - | - | - | - |
| 21 | 23-03-06 | $DJ694^{L}/+$ | ♀ | 77 | 52.56 | 8.26 | 54.96 | 81.48 | 0 | 0.004 | X | - | - | - | - |
| 21 | 23-03-06 | $DJ694^{L}$ | ♂ | 44 | 47.63 | 3.33 | 46.47 | 60.17 | - | - | - | - | - | - | - |
| 21 | 23-03-06 | $w^{1118}$ | ♂ | 66 | 53.34 | 4.73 | 50.67 | 78.20 | 0 | 5E-04 | X | - | - | - | - |
| 21 | 23-03-06 | $DJ694^{L}/+$ | ♂ | 75 | 57.17 | 4.50 | 50.73 | 82.62 | -3.25 | 5E-07 | ↓ | - | - | - | - |
| 22 | 23-03-08 | $DJ694^{L}$ | ♀ | 36 | 38.70 | 19.65 | 38.15 | 58.53 | - | - | - | - | - | - | - |

| Exp | Date yymmdd | Genotype | Sex | n | Mean | SD | Med | Max | %↓DJ694 mean | %↓DJ694 P | %↓DJ694 ∆ | %↑Rescue mean | %↑Rescue P(hom) | %↑Rescue P(het) | %↑Rescue ∆ |
|---|---|---|---|---|---|---|---|---|---|---|---|---|---|---|---|
| 22 | 23-03-08 | $w^{1118}$ | ♀ | 55 | 43.90 | 10.97 | 43.47 | 58.78 | 0 | 0.003 | X | - | - | - | - |
| 22 | 23-03-08 | $DJ694^{L}$/+ | ♀ | 74 | 46.24 | 11.92 | 45.59 | 70.31 | 0 | 0.002 | X | - | - | - | - |
| 22 | 23-03-08 | $DJ694^{L}$ | ♂ | 45 | 38.86 | 12.69 | 41.56 | 53.97 | - | - | - | - | - | - | - |
| 22 | 23-03-08 | $w^{1118}$ | ♂ | 47 | 45.97 | 7.85 | 47.42 | 64.67 | 0 | 0.1 | X | - | - | - | - |
| 22 | 23-03-08 | $DJ694^{L}$/+ | ♂ | 68 | 50.26 | 2.95 | 49.17 | 67.33 | 0 | 5E-04 | X | - | - | - | - |
| 23 | 24-07-09 | $DJ694^{H}$ | ♀ | 123 | 46.75 | 2.61 | 45.37 | 56.52 | - | - | - | - | - | - | - |
| 23 | 24-07-09 | $DJ694^{H}$/+ | ♀ | 130 | 61.69 | 1.87 | 57.94 | 72.27 | -17.49 | <2E-16 | ↓ | - | - | - | - |
| 23 | 24-07-09 | $DJ694^{H}$ | ♂ | 94 | 48.75 | 1.36 | 0.00 | 53.52 | - | - | - | - | - | - | - |
| 23 | 24-07-09 | $DJ694^{H}$/+ | ♂ | 115 | 53.56 | 0.76 | 52.11 | 63.92 | -5.07 | 3E-11 | ↓ | - | - | - | - |
| 23 | 24-07-09 | $DJ694^{L}$ | ♀ | 113 | 47.46 | 2.43 | 46.68 | 57.56 | - | - | - | - | - | - | - |
| 23 | 24-07-09 | $w^{1118}$ | ♀ | 114 | 64.54 | 2.03 | 62.88 | 72.11 | -20.19 | <2E-16 | ↓ | - | - | - | - |
| 23 | 24-07-09 | $DJ694^{L}$/+ | ♀ | 160 | 59.65 | 2.97 | 57.06 | 69.62 | -11.98 | <2E-16 | ↓ | - | - | - | - |
| 23 | 24-07-09 | $DJ694^{L}$;$UAS\text{-}EDTP^{E}$/+ | ♀ | 122 | 59.22 | 2.01 | 56.90 | 69.67 | - | - | - | 14.66 | <2E-16 | 0.2 | ↑ |
| 23 | 24-07-09 | $DJ694^{L}$ | ♂ | 79 | 48.46 | 1.97 | 46.64 | 52.70 | - | - | - | - | - | - | - |
| 23 | 24-07-09 | $w^{1118}$ | ♂ | 129 | 49.51 | 4.98 | 50.32 | 63.56 | 0 | 2E-06 | X | - | - | - | - |
| 23 | 24-07-09 | $DJ694^{L}$/+ | ♂ | 148 | 55.03 | 1.88 | 53.26 | 65.55 | -5.13 | <2E-16 | ↓ | - | - | - | - |
| 23 | 24-07-09 | $DJ694^{L}$;$UAS\text{-}EDTP^{E}$/+ | ♂ | 84 | 55.34 | 2.91 | 55.07 | 66.00 | - | - | - | 3.97 | 1E-15 | 0.2 | ↑ |
| 23 | 24-07-09 | $DJ694^{O}$ | ♀ | 77 | 53.98 | 2.40 | 54.19 | 58.41 | - | - | - | - | - | - | - |
| 23 | 24-07-09 | $DJ694^{O}$/+ | ♀ | 136 | 61.16 | 1.76 | 57.86 | 71.11 | -5.08 | 2E-12 | ↓ | - | - | - | - |
| 23 | 24-07-09 | $DJ694^{O}$ | ♂ | 59 | 52.44 | 1.26 | 51.52 | 55.53 | - | - | - | - | - | - | - |
| 23 | 24-07-09 | $DJ694^{O}$/+ | ♂ | 124 | 58.73 | 1.41 | 56.92 | 67.53 | -6.31 | <2E-16 | ↓ | - | - | - | - |
| 23 | 24-07-09 | $DJ694^{P}$ | ♀ | 67 | 54.31 | 4.94 | 54.52 | 59.21 | - | - | - | - | - | - | - |
| 23 | 24-07-09 | $DJ694^{P}$/+ | ♀ | 149 | 63.23 | 1.36 | 59.14 | 69.84 | -4.25 | <2E-16 | ↓ | - | - | - | - |
| 23 | 24-07-09 | $DJ694^{P}$ | ♂ | 92 | 50.91 | 2.56 | 51.13 | 56.16 | - | - | - | - | - | - | - |
| 23 | 24-07-09 | $DJ694^{P}$/+ | ♂ | 145 | 57.24 | 1.24 | 55.65 | 67.11 | -4.52 | 3E-14 | ↓ | - | - | - | - |
| 24 | 24-07-12 | $DJ694^{H}$ | ♀ | 179 | 44.05 | 5.81 | 41.56 | 55.99 | - | - | - | - | - | - | - |
| 24 | 24-07-12 | $DJ694^{H}$/+ | ♀ | 161 | 62.02 | 1.23 | 58.33 | 73.15 | -17.97 | <2E-16 | ↓ | - | - | - | - |
| 24 | 24-07-12 | $DJ694^{H}$ | ♂ | 162 | 44.20 | 2.90 | 44.96 | 51.52 | - | - | - | - | - | - | - |
| 24 | 24-07-12 | $DJ694^{H}$/+ | ♂ | 112 | 52.98 | 1.78 | 51.16 | 60.97 | -8.01 | <2E-16 | ↓ | - | - | - | - |
| 24 | 24-07-12 | $DJ694^{L}$ | ♀ | 166 | 50.05 | 2.12 | 49.97 | 57.90 | - | - | - | - | - | - | - |
| 24 | 24-07-12 | $w^{1118}$ | ♀ | 128 | 62.98 | 0.79 | 59.30 | 72.56 | -16.11 | <2E-16 | ↓ | - | - | - | - |
| 24 | 24-07-12 | $DJ694^{L}$/+ | ♀ | 148 | 58.81 | 3.47 | 57.35 | 74.25 | -5.73 | <2E-16 | ↓ | - | - | - | - |
| 24 | 24-07-12 | $DJ694^{L}$;$UAS\text{-}EDTP^{E}$/+ | ♀ | 122 | 55.83 | 6.70 | 54.15 | 65.21 | - | - | - | 0 | 1E-07 | 2E-05 | X |
| 24 | 24-07-12 | $DJ694^{L}$ | ♂ | 152 | 45.65 | 0.86 | 46.66 | 52.57 | - | - | - | - | - | - | - |
| 24 | 24-07-12 | $w^{1118}$ | ♂ | 103 | 51.54 | 0.67 | 49.48 | 58.38 | -8.57 | 1E-07 | ↓ | - | - | - | - |
| 24 | 24-07-12 | $DJ694^{L}$/+ | ♂ | 93 | 55.03 | 1.83 | 51.87 | 67.11 | -12.57 | 2E-16 | ↓ | - | - | - | - |
| 24 | 24-07-12 | $DJ694^{L}$;$UAS\text{-}EDTP^{E}$/+ | ♂ | 102 | 51.22 | 2.38 | 50.23 | 60.71 | - | - | - | 5.00 | 3E-08 | 5E-04 | ↑ |
| 24 | 24-07-12 | $DJ694^{O}$ | ♀ | 92 | 52.25 | 2.44 | 51.20 | 57.51 | - | - | - | - | - | - | - |
| 24 | 24-07-12 | $DJ694^{O}$/+ | ♀ | 178 | 62.74 | 3.01 | 59.35 | 73.71 | -8.44 | <2E-16 | ↓ | - | - | - | - |
| 24 | 24-07-12 | $DJ694^{O}$ | ♂ | 67 | 51.05 | 2.26 | 51.52 | 57.27 | - | - | - | - | - | - | - |
| 24 | 24-07-12 | $DJ694^{O}$/+ | ♂ | 129 | 54.14 | 1.95 | 52.59 | 65.40 | 0 | 0.03 | X | - | - | - | - |
| 24 | 24-07-12 | $DJ694^{P}$ | ♀ | 91 | 53.29 | 7.58 | 51.67 | 56.90 | - | - | - | - | - | - | - |
| 24 | 24-07-12 | $DJ694^{P}$/+ | ♀ | 163 | 64.34 | 0.84 | 59.69 | 74.59 | -4.15 | <2E-16 | ↓ | - | - | - | - |

| Exp | Date yymmdd | Genotype | Sex | n | Mean | SD | Med | Max | %↓DJ694 | | | %↑Rescue | | | |
|---|---|---|---|---|---|---|---|---|---|---|---|---|---|---|---|
| | | | | | | | | | mean | P | △ | mean | P(hom) | P(het) | △ |
| 24 | 24-07-12 | *DJ694* $^{P}$ | ♂ | 74 | 52.31 | 0.79 | 51.57 | 57.41 | - | - | - | - | - | - | - |
| 24 | 24-07-12 | *DJ694* $^{P}$ /+ | ♂ | 131 | 57.82 | 2.29 | 55.87 | 69.28 | -4.38 | 3E-10 | ↓ | - | - | - | - |

All p-values from this table come from survivial longrank analysis.
Column labeled "n" show longevity experiment sample size.
Column “Mean” represents the average lifespan per replicate, calculated as the mean of the vial average lifespans (2–4 vials). For each vial, average lifespan was calculated as the total lifespan of all flies divided by the number of flies in that vial. Total lifespan was calculated as ∑(number of deaths on day i × day i).
Column labeled "SD" represent standard deviation of the mean lifespan.
Column labeled "Med" is the median lifespan represent the time taken for the survival percentage to be 50%.
Column labeled "Max" is the max lifespan represent the time taken for the survival percentage to be 5%.
The column '% DJ694' represents the percentage decline in lifespan of homozygous *DJ694* relative to the relevant control (*DJ694/+* or *w*), calculated as the reduction from the lower error bar of the control (control mean minus SD) to the upper error bar of *DJ694* (*DJ694* mean plus SD), expressed as a percentage of the control lower error bar. This provides a conservative estimate of the minimum difference in lifespan between genotypes.
Column “△” under “%↓DJ694” indicates whether *DJ694* show a noticeble decrease when compared to the control. “↓” denotes that *DJ694* mean lifespan + its SD is lower than the control’s mean lifespan − its SD, whereas “X” denotes overlapping error bars.
The "mean" under column labeled "%↑Rescue" show the percent extention in lifespan when UAS-*EDTP* was added to the mutant. The "P(hom)" is the p-value comparing to homozygous *DJ694*. The "P(het)" is the p-value comparing to heterozygous *DJ694*.
Column “△” under “%↑Rescue” indicates whether a noticeable increase relative to *DJ694* is observed. “↑” denotes that the rescue genotype’s mean lifespan minus the rescue genotype's SD is greater than *DJ694* mean lifespan plus the *DJ694*'s SD, whereas “X” indicates error bar overlaps.

| Exp | Date (yymmdd) | Genotype | Sex | n | Mean | SD | Med | Max | vs. DJ694/+ P | vs. UAS/+ P | △ |
|---|---|---|---|---|---|---|---|---|---|---|---|
| 1 | 04-11-07 | *DJ694/+* | ♀ | 86 | 63.75 | 5.31 | 65.00 | 79.75 | - | - | - |
| 1 | 04-11-07 | *UAS-EDTP^A /+* | ♀ | 94 | 61.02 | 9.28 | 61.00 | 84.25 | - | - | - |
| 1 | 04-11-07 | *DJ694/+;UAS-EDTP^A /+* | ♀ | 109 | 59.97 | 5.97 | 61.00 | 83.25 | 1 | 1 | X |
| 1 | 04-11-07 | *DJ694/+* | ♂ | 108 | 59.84 | 6.65 | 58.00 | 71.75 | - | - | - |
| 1 | 04-11-07 | *UAS-EDTP^A /+* | ♂ | 97 | 46.40 | 2.07 | 47.00 | 59.75 | - | - | - |
| 1 | 04-11-07 | *DJ694/+;UAS-EDTP^A /+* | ♂ | 116 | 70.51 | 5.35 | 72.00 | 84.00 | 6E-14 | <2E-16 | X |
| 2 | 04-11-09 | *DJ694/+* | ♀ | 126 | 66.59 | 6.02 | 70.00 | 83.75 | - | - | - |
| 2 | 04-11-09 | *UAS-EDTP^A /+* | ♀ | 112 | 55.17 | 2.94 | 56.00 | 79.50 | - | - | - |
| 2 | 04-11-09 | *DJ694/+;UAS-EDTP^A /+* | ♀ | 105 | 69.20 | 3.21 | 73.00 | 84.75 | 0.5512 | 4E-09 | X |
| 2 | 04-11-09 | *DJ694/+* | ♂ | 124 | 62.01 | 5.87 | 63.00 | 82.25 | - | - | - |
| 2 | 04-11-09 | *UAS-EDTP^A /+* | ♂ | 108 | 50.44 | 7.39 | 50.50 | 63.75 | - | - | - |
| 2 | 04-11-09 | *DJ694/+;UAS-EDTP^A /+* | ♂ | 91 | 70.37 | 4.62 | 77.00 | 90.00 | 1E-07 | <2E-16 | X |
| 3 | 04-11-07 | *DJ694/+* | ♀ | 129 | 67.09 | 7.03 | 68.00 | 86.75 | - | - | - |
| 3 | 04-11-07 | *UAS-EDTP^A /+* | ♀ | 78 | 53.53 | 3.46 | 51.00 | 78.75 | - | - | - |
| 3 | 04-11-07 | *DJ694/+;UAS-EDTP^A /+* | ♀ | 101 | 42.36 | 3.47 | 37.00 | 74.25 | <2E-16 | 0.0064 | ↓ |
| 3 | 04-11-07 | *DJ694/+* | ♂ | 112 | 63.19 | 9.14 | 65.00 | 77.00 | - | - | - |
| 3 | 04-11-07 | *UAS-EDTP^A /+* | ♂ | 68 | 42.41 | 3.23 | 37.00 | 54.33 | - | - | - |
| 3 | 04-11-07 | *DJ694/+;UAS-EDTP^A /+* | ♂ | 86 | 53.26 | 10.91 | 54.00 | 78.00 | 0.0037 | 1E-05 | X |
| 4 | 04-11-09 | *DJ694/+* | ♀ | 116 | 58.83 | 3.68 | 59.00 | 87.25 | - | - | - |
| 4 | 04-11-09 | *UAS-EDTP^A /+* | ♀ | 133 | 58.80 | 3.06 | 59.00 | 84.75 | - | - | - |
| 4 | 04-11-09 | *DJ694/+;UAS-EDTP^A /+* | ♀ | 122 | 35.44 | 1.57 | 35.00 | 65.25 | <2E-16 | <2E-16 | ↓ |
| 4 | 04-11-09 | *DJ694/+* | ♂ | 93 | 54.64 | 2.92 | 56.00 | 76.00 | - | - | - |
| 4 | 04-11-09 | *UAS-EDTP^A /+* | ♂ | 117 | 49.81 | 3.15 | 52.00 | 67.25 | - | - | - |
| 4 | 04-11-09 | *DJ694/+;UAS-EDTP^A /+* | ♂ | 98 | 51.71 | 6.10 | 52.00 | 79.25 | 1 | 0.0104 | X |
| 5 | 04-12-01 | *DJ694/+* | ♀ | 103 | 74.87 | 5.53 | 79.00 | 91.50 | - | - | - |
| 5 | 04-12-01 | *UAS-EDTP^A /+* | ♀ | 104 | 52.93 | 4.75 | 55.00 | 70.50 | - | - | - |
| 5 | 04-12-01 | *DJ694/+;UAS-EDTP^A /+* | ♀ | 85 | 62.06 | 6.96 | 69.00 | 79.25 | 4E-13 | 0.0002 | X |
| 5 | 04-12-01 | *UAS-EDTP^B /+* | ♀ | 38 | 47.58 | 1.09 | 42.50 | 75.50 | - | - | - |
| 5 | 04-12-01 | *DJ694/+;UAS-EDTP^B /+* | ♀ | 101 | 68.94 | 7.55 | 76.00 | 84.50 | 8E-04 | 1E-07 | X |
| 5 | 04-12-01 | *DJ694/+* | ♂ | 94 | 70.09 | 4.54 | 69.00 | 78.50 | - | - | - |
| 5 | 04-12-01 | *UAS-EDTP^A /+* | ♂ | 108 | 56.42 | 9.41 | 62.00 | 74.00 | - | - | - |
| 5 | 04-12-01 | *DJ694/+;UAS-EDTP^A /+* | ♂ | 116 | 61.88 | 4.79 | 65.00 | 75.00 | 8E-06 | 1 | X |
| 5 | 04-12-01 | *UAS-EDTP^B /+* | ♂ | 64 | 45.03 | 5.29 | 41.00 | 61.67 | - | - | - |
| 5 | 04-12-01 | *DJ694/+;UAS-EDTP^B /+* | ♂ | 112 | 70.04 | 5.90 | 72.00 | 86.25 | 1 | <2E-16 | X |
| 6 | 04-12-04 | *DJ694/+* | ♀ | 112 | 74.08 | 8.22 | 80.00 | 88.75 | - | - | - |
| 6 | 04-12-04 | *UAS-EDTP^A /+* | ♀ | 112 | 46.76 | 3.57 | 48.00 | 67.75 | - | - | - |
| 6 | 04-12-04 | *DJ694/+;UAS-EDTP^A /+* | ♀ | 89 | 38.23 | 11.68 | 38.00 | 57.00 | <2E-16 | 0.0008 | X |
| 6 | 04-12-04 | *UAS-EDTP^B /+* | ♀ | 55 | 37.54 | 7.40 | 38.00 | 69.33 | - | - | - |
| 6 | 04-12-04 | *DJ694/+;UAS-EDTP^B /+* | ♀ | 118 | 48.50 | 8.91 | 50.00 | 68.50 | <2E-16 | 0.0139 | X |
| 6 | 04-12-04 | *DJ694/+* | ♂ | 96 | 66.93 | 3.62 | 69.00 | 79.00 | - | - | - |
| 6 | 04-12-04 | *UAS-EDTP^A /+* | ♂ | 114 | 56.80 | 1.28 | 59.00 | 78.00 | - | - | - |
| 6 | 04-12-04 | *DJ694/+;UAS-EDTP^A /+* | ♂ | 86 | 45.76 | 4.25 | 45.00 | 65.00 | <2E-16 | 1E-07 | ↓ |

| Exp | Date (yymmdd) | Genotype | Sex | n | Mean | SD | Med | Max | vs. DJ694/+ P | vs. UAS/+ P | ∆ |
|---|---|---|---|---|---|---|---|---|---|---|---|
| 6 | 04-12-04 | *UAS-EDTP$^B$/+* | ♂ | 68 | 35.93 | 6.39 | 38.00 | 66.00 | - | - | - |
| 6 | 04-12-04 | *DJ694/+;UAS-EDTP$^B$/+* | ♂ | 116 | 55.09 | 2.31 | 55.00 | 79.00 | 4E-05 | 5E-11 | X |
| 7 | 04-12-01 | *DJ694/+* | ♀ | 86 | 76.84 | 6.41 | 81.00 | 90.75 | - | - | - |
| 7 | 04-12-01 | *UAS-EDTP$^A$/+* | ♀ | 113 | 55.90 | 7.76 | 58.00 | 88.25 | - | - | - |
| 7 | 04-12-01 | *DJ694/+;UAS-EDTP$^A$/+* | ♀ | 104 | 64.69 | 6.00 | 62.00 | 83.75 | 2E-09 | 1 | X |
| 7 | 04-12-01 | *UAS-EDTP$^B$/+* | ♀ | 59 | 41.87 | 6.24 | 41.00 | 71.33 | - | - | - |
| 7 | 04-12-01 | *DJ694/+;UAS-EDTP$^B$/+* | ♀ | 106 | 66.84 | 4.39 | 69.00 | 88.75 | 2E-03 | 4E-10 | X |
| 7 | 04-12-01 | *DJ694/+* | ♂ | 84 | 65.03 | 5.54 | 69.00 | 81.75 | - | - | - |
| 7 | 04-12-01 | *UAS-EDTP$^A$/+* | ♂ | 109 | 52.34 | 2.30 | 55.00 | 77.75 | - | - | - |
| 7 | 04-12-01 | *DJ694/+;UAS-EDTP$^A$/+* | ♂ | 104 | 59.67 | 6.00 | 58.00 | 75.75 | 0.1267 | 0.1397 | X |
| 7 | 04-12-01 | *UAS-EDTP$^B$/+* | ♂ | 58 | 48.89 | 5.70 | 48.00 | 70.00 | - | - | - |
| 7 | 04-12-01 | *DJ694/+;UAS-EDTP$^B$/+* | ♂ | 101 | 67.81 | 3.91 | 69.00 | 83.00 | 1 | 1E-13 | X |
| 8 | 04-12-04 | *DJ694/+* | ♀ | 121 | 55.862 | 7.86 | 59 | 78.25 | - | - | - |
| 8 | 04-12-04 | *UAS-EDTP$^A$/+* | ♀ | 108 | 41.07 | 1.80 | 41.00 | 61.50 | - | - | - |
| 8 | 04-12-04 | *DJ694/+;UAS-EDTP$^A$/+* | ♀ | 108 | 45.38 | 5.06 | 45.00 | 70.25 | 9E-05 | 0.0752 | X |
| 8 | 04-12-04 | *UAS-EDTP$^B$/+* | ♀ | 55 | 47.75 | 7.88 | 41.00 | 75.33 | - | - | - |
| 8 | 04-12-04 | *DJ694/+;UAS-EDTP$^B$/+* | ♀ | 124 | 54.20 | 8.05 | 59.00 | 73.00 | 1 | 1 | X |
| 8 | 04-12-04 | *DJ694/+* | ♂ | 122 | 57.15 | 3.78 | 59.00 | 76.50 | - | - | - |
| 8 | 04-12-04 | *UAS-EDTP$^A$/+* | ♂ | 131 | 47.25 | 2.06 | 48.00 | 67.75 | - | - | - |
| 8 | 04-12-04 | *DJ694/+;UAS-EDTP$^A$/+* | ♂ | 68 | 50.31 | 1.65 | 52.00 | 70.67 | 2E-09 | 1 | X |
| 8 | 04-12-04 | *UAS-EDTP$^B$/+* | ♂ | 137 | 46.41 | 4.42 | 45.00 | 66.00 | - | - | - |
| 8 | 04-12-04 | *DJ694/+;UAS-EDTP$^B$/+* | ♂ | 125 | 65.62 | 3.22 | 66.00 | 87.00 | 8E-06 | 3E-10 | ↑ |

All p-values from this table come from survivial longrank analysis.
Column labeled "n" show longevity experiment sample size
Column labeled "Mean" represent average lifespan
Column labeled "SD" represent standard deviation of the mean lifespan.
Column labeled "Med" is the median lifespan represent the time taken for the survival percentage to be 50%
Column labeled "Max" is the max lifespan represent the time taken for the survival percentage to be 5%
Column labeled "vs." show the p-value comparing *DJ694/+;UAS-EDTP/+* to "*DJ694/+*" or "*UAS-EDTP/+*" controls.
Column labeled "∆" indicates whether the experimental genotype (*DJ694/+;UAS-EDTP/+)* showed a consistent change in lifespan relative to both controls (*DJ694/+ and UAS-EDTP/+).* A change was only declared when the experimental genotype showed the same direction of difference relative to both controls, with non-overlapping mean ± standard deviation ranges between the experimental group and each control.

Table S11. Longevity of *Q72/+;;MHC/UAS-EDTP* and control genotypes. 

| Exp | Date YYMMDD | Genotype | Sex | n | Mean | SD | Med | Max | Changes %Mean | Changes P-values | Changes Δ |
|---|---|---|---|---|---|---|---|---|---|---|---|
| 1 | 19-10-09 | *Q72/+;;MHC/+* | ♀ | 111 | 22.55 | 1.06 | 23.00 | 26.00 | 0 | 0.3 | X |
| 1 | 19-10-09 | *Q72/+;;MHC/UAS-EDTP* | ♀ | 145 | 22.84 | 0.25 | 23.00 | 26.00 | - | - | - |
| 1 | 19-10-09 | *UAS-EDTP/+* | ♀ | 134 | 36.98 | 2.33 | 35.00 | 57.25 | -33.36 | <2E-16 | ↓ |
| 2 | 19-10-11 | *Q72/+;;MHC/+* | ♀ | 119 | 21.59 | 0.44 | 21.00 | 24.50 | 0 | 7E-04 | X |
| 2 | 19-10-11 | *Q72/+;;MHC/UAS-EDTP* | ♀ | 103 | 22.36 | 1.10 | 24.00 | 25.50 | - | - | - |
| 2 | 19-10-11 | *UAS-EDTP/+* | ♀ | 116 | 46.58 | 3.75 | 45.00 | 66.00 | -45.23 | <2E-16 | ↓ |
| 3 | 20-02-05 | *Q72/+;;MHC/+* | ♀ | 106 | 22.63 | 0.48 | 23.00 | 26.50 | 0 | 0.05 | X |
| 3 | 20-02-05 | *Q72/+;;MHC/UAS-EDTP* | ♀ | 79 | 24.59 | 1.52 | 26.00 | 28.00 | - | - | - |
| 3 | 20-02-05 | *UAS-EDTP/+* | ♀ | 96 | 54.57 | 7.27 | 54.00 | 76.00 | -44.79 | <2E-16 | ↓ |
| 4 | 20-02-07 | *Q72/+;;MHC/+* | ♀ | 86 | 23.43 | 0.33 | 24.00 | 25.50 | 6.79 | 4E-13 | ↑ |
| 4 | 20-02-07 | *Q72/+;;MHC/UAS-EDTP* | ♀ | 94 | 25.86 | 0.49 | 26.00 | 30.25 | - | - | - |
| 4 | 20-02-07 | *UAS-EDTP/+* | ♀ | 98 | 60.81 | 7.15 | 60.00 | 85.00 | -50.88 | <2E-16 | ↓ |
| 1 | 19-10-09 | *Q72/+;;MHC/+* | ♂ | 138 | 27.90 | 0.48 | 27.35 | 31.44 | 0 | <2E-16 | X |
| 1 | 19-10-09 | *Q72/+;;MHC/+* | ♂ | 132 | 21.44 | 0.97 | 21.00 | 26.00 | | <2E-16 | |
| 1 | 19-10-09 | *Q72/+;;MHC/UAS-EDTP* | ♂ | 134 | 24.68 | 0.99 | 26.00 | 28.00 | - | - | - |
| 1 | 19-10-09 | *UAS-EDTP/+* | ♂ | 158 | 47.43 | 6.05 | 48.00 | 66.25 | -37.98 | <2E-16 | ↓ |
| 2 | 19-10-11 | *Q72/+;;MHC/+* | ♂ | 135 | 28.12 | 1.40 | 27.50 | 31.28 | 0 | 2E-08 | X |
| 2 | 19-10-11 | *Q72/+;;MHC/+* | ♂ | 132 | 22.37 | 1.36 | 21.00 | 25.50 | | 1E-12 | |
| 2 | 19-10-11 | *Q72/+;;MHC/UAS-EDTP* | ♂ | 92 | 24.84 | 0.90 | 24.00 | 30.25 | - | - | - |
| 2 | 19-10-11 | *UAS-EDTP/+* | ♂ | 125 | 58.72 | 11.45 | 52.00 | 69.25 | -45.55 | <2E-16 | ↓ |
| 3 | 20-02-05 | *Q72/+;;MHC/+* | ♂ | 114 | 29.51 | 0.38 | 28.16 | 33.36 | 0 | <2E-16 | X |
| 3 | 20-02-05 | *Q72/+;;MHC/+* | ♂ | 71 | 24.69 | 0.82 | 26.00 | 28.50 | | 0.003 | |
| 3 | 20-02-05 | *Q72/+;;MHC/UAS-EDTP* | ♂ | 51 | 26.48 | 0.74 | 26.00 | 30.00 | - | - | - |
| 3 | 20-02-05 | *UAS-EDTP/+* | ♂ | 94 | 71.59 | 4.48 | 75.00 | 90.00 | -59.44 | <2E-16 | ↓ |
| 4 | 20-02-07 | *Q72/+;;MHC/+* | ♂ | 90 | 28.85 | 1.74 | 27.14 | 34.95 | 0 | 3E-06 | X |
| 4 | 20-02-07 | *Q72/+;;MHC/+* | ♂ | 99 | 24.94 | 0.66 | 24.00 | 30.00 | | 2E-04 | |
| 4 | 20-02-07 | *Q72/+;;MHC/UAS-EDTP* | ♂ | 97 | 26.42 | 0.77 | 26.00 | 32.00 | - | - | - |
| 4 | 20-02-07 | *UAS-EDTP/+* | ♂ | 105 | 65.62 | 8.39 | 61.00 | 80.50 | -52.48 | <2E-16 | ↓ |

All flies are treated with RU486.
Column labeled "n" show longevity experiment sample size. Column labeled "Mean" represent average lifespan. Column labeled "SD" represent standard deviation of the mean lifespan. Column labeled "Med" is the median lifespan represent the time taken for the survival percentage to be 50%. Column labeled "Max" is the max lifespan represent the time taken for the survival percentage drop to 5%.
The column "Δ" under "Change" indicates the direction of difference between *Q72/+;;MHC/UAS-EDTP* and the controls. "↓" denotes that the experimental mean plus its SD is lower than the positive control mean minus its SD. "↑" denotes that the experimental mean minus its SD is higher than the negative control mean plus its SD. "X" denotes overlapping error bars between the experimental group and both controls. The column "%mean" reports the magnitude of the difference when error bars do not overlap. For "↓", it is calculated as the difference between the positive control lower error bar (mean minus SD) and the experimental upper error bar (mean plus SD), divided by the positive control mean minus its SD. For "↑", it is calculated as the difference between the experimental lower error bar (mean minus SD) and the negative control upper error bar (mean plus SD), divided by the negative control mean plus its SD. When "X" is indicated, "%mean" is reported as 0. The column "P-value" shows the result of statistical comparisons between *Q72/+;;MHC/UAS-EDTP* and each respective control. All p-values from this table come from survival logrank analysis. For males, an additional negative control was included; if the mean lifespan of *Q72/+;;MHC/UAS-EDTP* fell between the two negative controls, no change was declared.

| Genotypes used in Experiment | Cross: Female | Cross: Male |
|---|---|---|
| *$w^{1118}$* | *$w^{1118}$* | *$w^{1118}$* |
| *DJ694* | *DJ694* | *DJ694* |
| *$DJ694^{H}$* | *$DJ694^{H}$* | *$DJ694^{H}$* |
| *$DJ694^{L}$* | *$DJ694^{L}$* | *$DJ694^{L}$* |
| *$DJ694^{O}$* | *$DJ694^{O}$* | *$DJ694^{O}$* |
| *$DJ694^{P}$* | *$DJ694^{P}$* | *$DJ694^{P}$* |
| *DJ694/+* | *DJ694* | *$w^{1118}$* |
| *$DJ694^{H}$/+* | *$w^{1118}$* | *$DJ694^{H}$* |
| *$DJ694^{L}$/+* | *$w^{1118}$* | *$DJ694^{L}$* |
| *$DJ694^{O}$/+* | *$w^{1118}$* | *$DJ694^{O}$* |
| *$DJ694^{P}$/+* | *$w^{1118}$* | *$DJ694^{P}$* |
| *DJ694;UAS-$EDTP^{A}$/+* | *DJ694* | *DJ694;UAS-$EDTP^{A}$* |
| *DJ694;UAS-$EDTP^{B}$/+* | *DJ694* | *DJ694;UAS-$EDTP^{B}$* |
| *DJ694;UAS-$EDTP^{A}$/UAS-$EDTP^{B}$* | *DJ694;UAS-$EDTP^{A}$* | *DJ694;UAS-$EDTP^{B}$* |
| *DJ694;UAS-$EDTP^{E}$/+* | *$DJ694^{L}$* | *DJ694;UAS-$EDTP^{E}$* |
| *UAS-$EDTP^{A}$/+* | *$w^{1118}$* | *UAS-$EDTP^{A}$* |
| *UAS-$EDTP^{B}$/+* | *$w^{1118}$* | *+/Cyo;UAS-$EDTP^{B}$* |
| *DJ694/+;UAS-$EDTP^{A}$/+* | *$w^{1118}$* | *DJ694;UAS-$EDTP^{A}$* |
| *DJ694/+;UAS-$EDTP^{B}$/+* | *$w^{1118}$* | *DJ694;UAS-$EDTP^{B}$* |
| *UAS-EDTP/+* | *$w^{1118}$* | *UAS-$EDTP^{E}$* |
| *Q72/+;;MHC/+* | *Q72;;MHC* | *$w^{1118}$* |
| *Q72/Y;;MHC/+* | *Q72;;MHC* | *Q72* |
| *Q72/+;;MHC/UAS-EDTP* | *Q72;;MHC* | *UAS-$EDTP^{E}$* |

Each row describes the parental cross used to generate a specific experimental genotype. Female parent is listed first, followed by male parent.

| | | | | Females | | | | | | | | | Males | | | | | | | | |
|---|---|---|---|---|---|---|---|---|---|---|---|---|---|---|---|---|---|---|---|---|---|
| | | | Date | Age (days) | | | | | | | | | Age (days) | | | | | | | | |
| Genotype | Parameter | Rep | YYMMDD | 2 | 4 | 8 | 15 | 22 | 29 | 30 | 36 | 43 | 2 | 4 | 8 | 15 | 22 | 29 | 30 | 36 | 43 |
| *w1118* | %Immobile | Rep 1 | 24-07-11 | 4.0 | 11.3 | 8.4 | 6.1 | 8.9 | 33.5 | | 99.4 | 36.6 | 35.9 | 19.8 | 4.5 | 65.6 | 68.5 | 21.3 | | 61.9 | 64.4 |
| | | | | 5.3 | 14.8 | 10.3 | 11.3 | 20.7 | 11.4 | | 88.5 | 57.9 | 8.1 | 56.6 | 86.3 | 5.4 | 27.0 | 64.0 | | 93.6 | 10.2 |
| | | | | 3.5 | 11.0 | 4.3 | 51.0 | 24.9 | 41.7 | | 97.6 | 68.0 | 7.0 | 10.5 | 29.9 | 73.9 | 76.6 | 47.8 | | 95.5 | 55.0 |
| | | | | 5.6 | 2.3 | 75.2 | 8.9 | 26.2 | 78.4 | | 92.7 | 35.0 | 5.8 | 22.1 | 26.2 | 81.1 | 11.0 | 77.3 | | 90.6 | 58.3 |
| | | | | 5.6 | 5.6 | 64.4 | 8.0 | 22.9 | 71.6 | | 83.5 | 29.3 | 10.3 | 3.4 | 21.1 | 1.9 | 70.8 | 72.6 | | | 69.9 |
| | | Rep 2 | 24-07-14 | 4.5 | 9.2 | 13.7 | | 5.8 | 50.0 | | 90.6 | 9.9 | 10.7 | 11.3 | 69.9 | 1.6 | 88.7 | 83.1 | | 76.2 | 14.6 |
| | | | | 6.5 | 8.2 | 75.7 | | 21.3 | 3.3 | | 95.2 | 12.2 | 10.5 | 8.3 | 7.0 | 83.4 | 18.3 | 85.6 | | 94.8 | 14.7 |
| | | | | 4.9 | 3.8 | 15.8 | | 10.7 | 14.7 | | 91.4 | 35.5 | 78.1 | 75.9 | 84.3 | 57.0 | 82.2 | 69.1 | | 80.6 | 1.7 |
| | | | | 1.9 | 15.9 | 78.6 | | 10.8 | 20.0 | | 79.3 | 43.9 | 3.6 | 67.8 | 51.3 | 82.4 | 7.5 | 80.5 | | 64.2 | 43.1 |
| | | | | | | | | 17.1 | 47.1 | | 94.3 | 24.5 | 4.0 | 11.3 | 80.3 | 72.6 | 17.3 | 32.2 | | 48.9 | 36.9 |
| | | Rep 3 | 23-03-09 | | | 16.9 | 10.8 | 6.6 | | 26.3 | | | | | 23.4 | 7.4 | 8.6 | | 5.3 | | |
| | | | | | | 14.7 | 8.1 | 4.2 | | 23.4 | | | | | 13.8 | 3.4 | 6.7 | | 4.9 | | |
| | | | | | | 11.6 | 11.3 | 11.2 | | 15.5 | | | | | 21.5 | 7.2 | 1.5 | | 5.3 | | |
| | | | | | | 4.2 | 11.9 | 9.7 | | 24.0 | | | | | 8.9 | 6.6 | 5.3 | | 15.8 | | |
| | | | | | | 20.0 | 5.0 | 8.6 | | 20.3 | | | | | 95.8 | 6.0 | 9.9 | | 5.2 | | |
| | | Rep 4 | 23-03-13 | | | 0.2 | 14.5 | 6.9 | | 5.1 | | | | | 17.6 | 27.9 | 11.5 | | 21.7 | | |
| | | | | | | 4.6 | 16.4 | 5.0 | | 10.0 | | | | | 4.4 | 44.0 | 7.5 | | 11.1 | | |
| | | | | | | 2.9 | 9.8 | 5.4 | | 7.9 | | | | | 4.4 | 16.4 | 20.4 | | 86.3 | | |
| | | | | | | 3.9 | 16.7 | 14.1 | | 7.5 | | | | | 5.3 | 48.4 | 64.0 | | 75.0 | | |
| | | | | | | 21.4 | 50.5 | 22.4 | | 14.6 | | | | | 3.6 | 41.5 | 5.9 | | 79.2 | | |
| | | Rep 5 | 23-03-13 | | | 7.2 | 6.3 | 13.2 | | 9.2 | | | | | 12.3 | 53.9 | 10.6 | | 2.9 | | |
| | | | | | | 11.6 | 21.7 | 21.4 | | 5.0 | | | | | 10.1 | 63.3 | 11.0 | | 9.2 | | |
| | | | | | | 6.8 | 12.2 | 2.4 | | 10.4 | | | | | 6.2 | 8.9 | 5.7 | | 5.1 | | |
| | | | | | | 2.3 | 10.4 | 9.7 | | 43.9 | | | | | 11.2 | 13.0 | 7.6 | | 9.9 | | |
| | | | | | | 10.7 | 12.0 | 3.8 | | 4.3 | | | | | 2.9 | 34.0 | 7.1 | | 11.4 | | |
| | | Rep 6 | 23-03-15 | | | 26.6 | 13.0 | 5.1 | | 13.7 | | | | | 13.3 | 12.7 | 11.8 | | 12.1 | | |
| | | | | | | 7.0 | 19.7 | 20.4 | | 36.5 | | | | | 42.5 | 32.7 | 12.8 | | 19.1 | | |
| | | | | | | 9.3 | 10.2 | 8.6 | | 22.3 | | | | | 9.1 | 90.4 | 8.3 | | 39.1 | | |
| | | | | | | 10.6 | 69.2 | 7.1 | | 12.8 | | | | | 35.7 | 62.9 | 3.9 | | 38.2 | | |
| | | | | | | 10.7 | 60.8 | 1.9 | | 5.6 | | | | | 9.9 | 21.2 | 9.6 | | 53.2 | | |

| Genotype | Parameter | Rep | Date YYMMDD | Females Age (days) 2 | 4 | 8 | 15 | 22 | 29 | 30 | 36 | 43 | Males Age (days) 2 | 4 | 8 | 15 | 22 | 29 | 30 | 36 | 43 |
|---|---|---|---|---|---|---|---|---|---|---|---|---|---|---|---|---|---|---|---|---|---|
| *w1118* | Distance | Rep 1 | 24-07-11 | 7719.3 | 8634.9 | 8855.1 | 9070.4 | 9553.6 | 4661.7 | | 169.0 | 5485.2 | 3711.1 | 8793.9 | 7419.3 | 1514.2 | 1866.3 | 8382.5 | | 4001.6 | 2443.5 |
| | | | | 9140.7 | 7882.9 | 8261.0 | 8988.3 | 4985.5 | 7684.1 | | 708.3 | 2769.7 | 5614.4 | 2366.4 | 918.5 | 9479.9 | 4690.7 | 2553.1 | | 625.0 | 9469.0 |
| | | | | 11343.7 | 7523.5 | 9481.2 | 2557.9 | 3764.2 | 4251.8 | | 291.3 | 2100.3 | 9214.5 | 8724.4 | 5778.6 | 1229.5 | 1027.6 | 5403.8 | | 552.4 | 3333.1 |
| | | | | 7362.0 | 7795.2 | 1451.6 | 7098.1 | 3820.0 | 940.4 | | 504.1 | 5031.3 | 9287.8 | 8531.8 | 7465.6 | 856.0 | 10801.0 | 871.8 | | 712.1 | 2759.6 |
| | | | | 7653.2 | 8653.8 | 2103.3 | 10920.0 | 5053.7 | 1327.9 | | 892.4 | 6233.3 | 8448.2 | 11959.2 | 10326.9 | 11895.7 | 950.6 | 1205.3 | | | 2580.9 |
| | | Rep 2 | 24-07-14 | 12091.6 | 8363.9 | 7535.3 | | 6930.9 | 4139.2 | | 587.9 | 10108.7 | 8528.1 | 10012.0 | 1990.7 | 11504.8 | 717.5 | 800.5 | | 1280.3 | 8918.6 |
| | | | | 8697.6 | 11033.1 | 1146.5 | | 4268.0 | 9314.1 | | 504.5 | 7726.1 | 7153.6 | 11619.7 | 8788.2 | 842.4 | 5757.2 | 739.5 | | 598.2 | 11169.6 |
| | | | | 9144.6 | 11133.1 | 7034.4 | | 7552.5 | 8273.6 | | 520.4 | 4876.9 | 1156.6 | 912.0 | 777.6 | 1923.9 | 1355.5 | 2838.3 | | 1155.3 | 15846.2 |
| | | | | 11853.2 | 8988.3 | 1005.1 | | 6711.5 | 5279.6 | | 930.5 | 4118.9 | 8842.9 | 2507.0 | 3133.1 | 1015.0 | 11295.7 | 948.6 | | 2832.2 | 2385.7 |
| | | | | | | | | 5284.8 | 4435.6 | | 421.9 | 6693.7 | 10569.3 | 8789.6 | 1372.1 | 1255.7 | 6945.8 | 7253.8 | | 4418.0 | 4788.5 |
| | | Rep 3 | 23-03-09 | | | 10207.6 | 14787.8 | 8753.5 | | 9968.3 | | | | | 9788.2 | 19287.3 | 12741.3 | | 17208.9 | | |
| | | | | | | 9789.3 | 11418.3 | 12441.5 | | 9639.5 | | | | | 10082.8 | 14451.6 | 13214.6 | | 17639.8 | | |
| | | | | | | 5858.7 | 9615.4 | 14476.1 | | 9259.6 | | | | | 8929.7 | 17667.8 | 16956.0 | | 12794.6 | | |
| | | | | | | 12678.8 | 12149.4 | 5730.3 | | 10274.0 | | | | | 15190.4 | 13911.3 | 19073.6 | | 10178.1 | | |
| | | | | | | 12402.0 | 14794.1 | 10857.5 | | 8179.6 | | | | | 636.6 | 14432.3 | 11337.4 | | 15694.7 | | |
| | | Rep 4 | 23-03-13 | | | 19341.3 | 11361.7 | 15385.3 | | 16010.2 | | | | | 11966.2 | 8615.2 | 12954.7 | | 10745.4 | | |
| | | | | | | 15775.3 | 11057.5 | 12130.4 | | 12792.7 | | | | | 17821.5 | 3490.4 | 13553.2 | | 10414.0 | | |
| | | | | | | 15210.3 | 10637.4 | 10437.8 | | 12522.8 | | | | | 16610.9 | 8380.4 | 9209.8 | | 761.2 | | |
| | | | | | | 14338.8 | 9481.9 | 9098.8 | | 10096.1 | | | | | 15816.8 | 3496.6 | 4501.9 | | 1335.5 | | |
| | | | | | | 9498.8 | 2838.0 | 7959.3 | | 9862.4 | | | | | 18194.9 | 6234.2 | 14947.6 | | 1207.5 | | |
| | | Rep 5 | 23-03-13 | | | 11013.1 | 13377.1 | 9166.5 | | 12274.1 | | | | | 17588.9 | 3680.1 | 11998.1 | | 17848.5 | | |
| | | | | | | 9530.4 | 7205.9 | 9840.4 | | 14794.7 | | | | | 11954.8 | 2134.7 | 15274.6 | | 13848.1 | | |
| | | | | | | 12621.5 | 6071.0 | 12425.6 | | 13602.3 | | | | | 14972.5 | 10896.8 | 12803.1 | | 16228.6 | | |
| | | | | | | 10585.5 | 11652.9 | 11977.8 | | 3807.8 | | | | | 14772.1 | 10552.6 | 12502.6 | | 12138.2 | | |
| | | | | | | 11204.9 | 11275.8 | 15343.9 | | 10726.3 | | | | | 15443.0 | 5372.1 | 11618.2 | | 10085.0 | | |
| | | Rep 6 | 23-03-15 | | | 5968.8 | 8063.7 | 14760.4 | | 10801.8 | | | | | 13125.2 | 13878.5 | 11679.7 | | 10782.7 | | |
| | | | | | | 15214.4 | 5889.5 | 10155.0 | | 4660.6 | | | | | 6477.7 | 5015.1 | 10906.7 | | 8432.9 | | |
| | | | | | | 10824.6 | 8156.8 | 11719.5 | | 7025.9 | | | | | 13660.3 | 914.2 | 12972.5 | | 3014.8 | | |
| | | | | | | 10532.5 | 2307.6 | 16933.4 | | 10892.3 | | | | | 9842.3 | 3625.5 | 14281.9 | | 5143.0 | | |
| | | | | | | 13302.8 | 2283.4 | 12886.8 | | 15815.0 | | | | | 11687.5 | 11065.2 | 11468.1 | | 2866.1 | | |

| Genotype | Parameter | Rep | YYMMDD Date | Females Age (days) 2 | 4 | 8 | 15 | 22 | 29 | 30 | 36 | 43 | Males Age (days) 2 | 4 | 8 | 15 | 22 | 29 | 30 | 36 | 43 |
|---|---|---|---|---|---|---|---|---|---|---|---|---|---|---|---|---|---|---|---|---|---|
| *w1118* | Velocity | Rep 1 | 24-07-11 | 8.6 | 10.3 | 9.3 | 10.4 | 11.2 | 6.8 | | 1.3 | 5.7 | 5.4 | 11.4 | 7.5 | 3.3 | 4.8 | 11.2 | | 8.6 | 2.3 |
| | | | | 9.9 | 10.0 | 8.8 | 11.1 | 6.3 | 9.3 | | 1.6 | 2.1 | 6.1 | 3.8 | 1.6 | 11.0 | 6.5 | 6.4 | | 1.6 | 7.4 |
| | | | | 12.3 | 9.0 | 9.5 | 5.1 | 4.8 | 7.2 | | 1.3 | 2.1 | 10.7 | 10.1 | 7.6 | 2.9 | 2.1 | 10.1 | | 1.4 | 2.7 |
| | | | | 7.9 | 8.5 | 2.0 | 8.4 | 5.1 | 2.0 | | 2.5 | 3.3 | 10.2 | 11.4 | 9.9 | 1.7 | 13.1 | 1.8 | | 1.6 | 2.2 |
| | | | | 8.2 | 9.6 | 2.5 | 13.2 | 6.1 | 2.5 | | 2.3 | 6.3 | 9.0 | 13.0 | 12.7 | 12.9 | 2.5 | 1.7 | | | 3.2 |
| | | Rep 2 | 24-07-14 | 13.1 | 9.8 | 8.3 | | 7.3 | 7.0 | | 1.6 | 8.8 | 10.6 | 12.4 | 5.7 | 12.4 | 2.4 | 2.0 | | 1.9 | 7.5 |
| | | | | 9.5 | 13.1 | 2.2 | | 4.9 | 9.4 | | 2.0 | 6.6 | 8.7 | 13.6 | 9.6 | 2.5 | 6.8 | 1.6 | | 1.9 | 9.7 |
| | | | | 10.0 | 12.7 | 7.8 | | 8.2 | 9.1 | | 1.5 | 4.0 | 2.7 | 2.0 | 1.6 | 4.0 | 5.2 | 7.7 | | 1.8 | 13.0 |
| | | | | 12.6 | 11.4 | 1.9 | | 6.9 | 5.7 | | 2.5 | 2.7 | 10.0 | 7.0 | 6.0 | 3.4 | 12.0 | 1.8 | | 5.5 | 2.8 |
| | | | | | | | | 5.7 | 6.9 | | 1.8 | 5.5 | 12.0 | 10.6 | 5.7 | 2.3 | 7.8 | 9.8 | | 7.2 | 3.5 |
| | | Rep 3 | 23-03-09 | | | 9.9 | 13.3 | 7.6 | | 11.9 | | | | | 10.1 | 17.8 | 12.2 | | 16.2 | | |
| | | | | | | 8.6 | 8.9 | 11.1 | | 10.9 | | | | | 9.5 | 12.0 | 12.3 | | 16.5 | | |
| | | | | | | 4.9 | 5.6 | 14.3 | | 9.4 | | | | | 9.0 | 15.9 | 15.4 | | 11.6 | | |
| | | | | | | 10.3 | 9.1 | 4.8 | | 11.6 | | | | | 13.5 | 11.6 | 17.8 | | 10.2 | | |
| | | | | | | 12.4 | 10.3 | 10.1 | | 8.8 | | | | | 1.4 | 11.7 | 10.6 | | 14.4 | | |
| | | Rep 4 | 23-03-13 | | | 16.4 | 11.1 | 13.5 | | 11.7 | | | | | 12.4 | 10.1 | 11.9 | | 12.8 | | |
| | | | | | | 14.2 | 11.0 | 10.2 | | 8.4 | | | | | 16.3 | 3.7 | 12.1 | | 11.0 | | |
| | | | | | | 12.3 | 9.9 | 8.7 | | 7.2 | | | | | 14.7 | 8.4 | 9.1 | | 2.1 | | |
| | | | | | | 10.9 | 9.4 | 8.0 | | 7.0 | | | | | 13.9 | 3.0 | 9.2 | | 2.3 | | |
| | | | | | | 9.8 | 2.7 | 7.9 | | 7.8 | | | | | 16.2 | 8.3 | 13.4 | | 1.8 | | |
| | | Rep 5 | 23-03-13 | | | 9.4 | 12.3 | 8.3 | | 6.3 | | | | | 16.6 | 5.6 | 10.9 | | 15.7 | | |
| | | | | | | 8.5 | 7.1 | 9.8 | | 11.6 | | | | | 10.5 | 2.5 | 14.3 | | 11.5 | | |
| | | | | | | 11.0 | 5.1 | 10.3 | | 10.4 | | | | | 13.1 | 10.0 | 10.8 | | 13.1 | | |
| | | | | | | 8.8 | 10.6 | 10.9 | | 3.6 | | | | | 13.4 | 10.0 | 10.9 | | 9.2 | | |
| | | | | | | 10.2 | 10.2 | 13.3 | | 8.1 | | | | | 13.1 | 5.2 | 9.8 | | 8.2 | | |
| | | Rep 6 | 23-03-15 | | | 6.1 | 7.6 | 13.1 | | 9.4 | | | | | 12.9 | 13.6 | 10.8 | | 10.1 | | |
| | | | | | | 14.0 | 5.8 | 10.4 | | 3.1 | | | | | 8.9 | 5.6 | 10.4 | | 8.4 | | |
| | | | | | | 9.7 | 7.2 | 10.4 | | 3.8 | | | | | 12.9 | 1.5 | 11.8 | | 3.1 | | |
| | | | | | | 10.1 | 4.7 | 15.3 | | 7.4 | | | | | 12.9 | 7.1 | 12.1 | | 6.2 | | |
| | | | | | | 12.4 | 3.5 | 9.4 | | 10.9 | | | | | 10.9 | 11.8 | 10.2 | | 4.0 | | |

| | | | Date | Females | | | | | | | | | Males | | | | | | | | |
|---|---|---|---|---|---|---|---|---|---|---|---|---|---|---|---|---|---|---|---|---|---|
| | | | | Age (days) | | | | | | | | | Age (days) | | | | | | | | |
| Genotype | Parameter | Rep | YYMMDD | 2 | 4 | 8 | 15 | 22 | 29 | 30 | 36 | 43 | 2 | 4 | 8 | 15 | 22 | 29 | 30 | 36 | 43 |
| *DJ694*[H] | % Immobile | Rep 1 | 24-07-11 | 64.1 | 13.6 | 24.9 | 56.6 | 14.0 | | | 88.9 | 91.5 | 15.8 | 4.8 | 6.0 | 8.6 | 22.0 | 79.9 | | 28.7 | 36.0 |
| | | | | 6.3 | 19.2 | 7.4 | 15.5 | 28.7 | | | 86.9 | 91.5 | 8.2 | 13.9 | 37.3 | 20.4 | 22.9 | 63.2 | | 23.1 | 50.0 |
| | | | | 22.3 | 8.5 | 38.8 | 86.8 | 65.6 | | | 30.4 | 94.8 | 33.7 | 10.3 | 29.8 | 53.4 | 9.3 | 63.2 | | 38.3 | 37.6 |
| | | | | 7.9 | 21.1 | 18.1 | 14.6 | 7.7 | | | 74.6 | 93.3 | 14.7 | 5.9 | 64.0 | 3.8 | 22.3 | 50.7 | | 31.8 | 31.7 |
| | | | | 21.6 | 17.5 | 6.8 | 40.7 | 4.1 | | | 84.0 | 87.3 | 9.2 | 11.5 | 22.4 | 13.1 | 7.4 | 10.4 | | 55.4 | 26.6 |
| | | Rep 2 | 24-07-14 | 5.1 | 19.4 | 18.1 | | 6.6 | | | 56.6 | 46.7 | 28.6 | 8.7 | 20.0 | 24.1 | 7.5 | 84.6 | | 9.2 | 51.5 |
| | | | | 9.1 | 7.4 | 25.8 | | 7.2 | | | 49.7 | 16.6 | 8.5 | 8.3 | 29.8 | 43.8 | 2.6 | 69.7 | | 13.2 | 31.1 |
| | | | | 10.2 | 19.9 | 12.0 | | 13.7 | | | 87.4 | 31.0 | 5.1 | 6.8 | 81.1 | 12.1 | 15.7 | 82.0 | | 23.0 | 39.9 |
| | | | | 7.6 | 17.1 | 52.6 | | 6.3 | | | 48.1 | 62.2 | 54.8 | 10.4 | 50.2 | 24.6 | 58.7 | 21.8 | | 14.7 | 49.5 |
| | | | | 10.2 | 17.4 | 37.8 | | 77.2 | | | 50.0 | 45.8 | 7.8 | 6.9 | 58.3 | 22.5 | 60.8 | 34.0 | | 5.0 | 2.2 |
| | Distance | Rep 1 | 24-07-11 | 2015.5 | 6758.1 | 4438.8 | 1911.8 | 6552.7 | | | 756.7 | 864.9 | 6068.2 | 7349.1 | 10385.3 | 7520.7 | 6124.4 | 934.3 | | 4344.5 | 4842.7 |
| | | | | 7074.3 | 6920.3 | 8054.6 | 6226.4 | 3733.4 | | | 852.4 | 759.5 | 10204.2 | 8692.4 | 6428.3 | 7222.5 | 6354.9 | 1459.2 | | 5606.7 | 3655.3 |
| | | | | 7253.0 | 8418.3 | 4107.0 | 760.2 | 1449.9 | | | 3529.0 | 612.4 | 4345.7 | 9010.4 | 5845.0 | 2060.7 | 8787.9 | 1453.2 | | 3583.2 | 6066.3 |
| | | | | 8492.7 | 5795.2 | 7020.7 | 7649.4 | 6862.6 | | | 1098.5 | 537.1 | 7613.8 | 9110.8 | 2640.5 | 11936.6 | 6742.6 | 2316.2 | | 4994.7 | 8106.7 |
| | | | | 5376.2 | 6649.5 | 10030.2 | 2892.2 | 7245.7 | | | 937.3 | 965.4 | 9547.4 | 8160.0 | 7915.0 | 10858.3 | 10435.8 | 6502.6 | | 2780.6 | 8662.5 |
| | | Rep 2 | 24-07-14 | 8197.5 | 7959.8 | 7004.7 | | 6693.4 | | | 1760.7 | 3683.4 | 6079.4 | 8414.0 | 5973.7 | 3959.3 | 10590.6 | 789.6 | | 7302.5 | 4413.6 |
| | | | | 8696.2 | 9441.0 | 5091.4 | | 8044.0 | | | 1521.5 | 11725.5 | 7475.6 | 9699.5 | 5574.1 | 4811.5 | 10528.0 | 2597.5 | | 6497.8 | 7983.5 |
| | | | | 7071.5 | 7240.2 | 7873.9 | | 7593.2 | | | 781.5 | 10576.2 | 9806.6 | 9808.8 | 892.2 | 10100.0 | 9223.3 | 847.4 | | 6025.1 | 6472.7 |
| | | | | 9171.1 | 7594.0 | 2768.6 | | 8335.6 | | | 1659.5 | 4031.2 | 2775.1 | 9591.7 | 4073.4 | 4707.1 | 3515.8 | 6723.0 | | 8151.3 | 4572.1 |
| | | | | 9955.4 | 7642.6 | 3447.5 | | 1187.8 | | | 1717.1 | 3443.8 | 10229.4 | 12005.9 | 2964.7 | 5169.6 | 3980.8 | 4073.0 | | 8536.6 | 17180.0 |
| | Velocity | Rep 1 | 24-07-11 | 3.6 | 8.3 | 5.0 | 2.5 | 8.2 | | | 1.5 | 1.7 | 7.3 | 7.5 | 11.4 | 8.9 | 8.3 | 1.5 | | 4.9 | 3.0 |
| | | | | 6.2 | 9.1 | 8.4 | 8.0 | 5.2 | | | 1.5 | 1.5 | 12.0 | 10.4 | 10.1 | 9.8 | 8.8 | 2.2 | | 6.1 | 2.6 |
| | | | | 9.1 | 10.0 | 4.5 | 1.5 | 2.7 | | | 4.2 | 1.5 | 6.0 | 10.4 | 8.3 | 3.4 | 10.7 | 1.8 | | 4.4 | 4.1 |
| | | | | 9.9 | 7.7 | 7.6 | 10.0 | 8.0 | | | 1.9 | 1.6 | 9.2 | 10.1 | 5.9 | 13.8 | 9.4 | 3.6 | | 5.9 | 5.0 |
| | | | | 4.6 | 8.6 | 10.6 | 3.9 | 8.0 | | | 1.7 | 1.5 | 10.5 | 9.5 | 10.3 | 13.2 | 12.5 | 7.5 | | 4.7 | 5.1 |
| | | Rep 2 | 24-07-14 | 8.5 | 10.7 | 8.0 | | 7.1 | | | 2.8 | 2.5 | 9.2 | 9.9 | 7.3 | 5.0 | 11.6 | 1.5 | | 7.2 | 3.9 |
| | | | | 9.9 | 11.1 | 6.2 | | 8.7 | | | 1.9 | 5.7 | 8.9 | 11.5 | 7.9 | 8.5 | 10.9 | 7.2 | | 6.7 | 5.1 |
| | | | | 7.9 | 9.4 | 8.0 | | 8.8 | | | 1.6 | 6.9 | 11.5 | 11.4 | 1.6 | 12.1 | 11.0 | 1.9 | | 6.9 | 5.0 |
| | | | | 10.3 | 9.8 | 4.4 | | 8.9 | | | 2.1 | 5.0 | 4.6 | 11.7 | 7.8 | 6.2 | 7.6 | 8.3 | | 8.6 | 4.2 |
| | | | | 10.9 | 10.0 | 4.5 | | 2.3 | | | 2.1 | 2.2 | 13.0 | 14.1 | 6.4 | 6.4 | 9.0 | 4.1 | | 8.2 | 13.5 |

| Genotype | Parameter | Rep | Date YYMMDD | Females Age (days) 2 | 4 | 8 | 15 | 22 | 29 | 30 | 36 | 43 | Males Age (days) 2 | 4 | 8 | 15 | 22 | 29 | 30 | 36 | 43 |
|---|---|---|---|---|---|---|---|---|---|---|---|---|---|---|---|---|---|---|---|---|---|
| *DJ694^H/+* | %Immobile | Rep 1 | 24-07-11 | 1.9 | 9.1 | 6.1 | 8.2 | 15.2 | | | 85.9 | 22.8 | 7.0 | 6.0 | 10.7 | 5.3 | 6.5 | 73.1 | | 46.0 | 25.5 |
| | | | | 9.9 | 11.1 | 5.8 | 9.6 | 13.8 | | | 83.0 | 75.0 | 7.7 | 2.9 | 22.1 | 20.5 | 6.6 | 66.2 | | 56.1 | 36.0 |
| | | | | 11.8 | 5.5 | 6.6 | 6.2 | 7.4 | | | 85.4 | 3.2 | 10.8 | 2.1 | 4.9 | 8.3 | 7.9 | 65.0 | | 41.7 | 3.5 |
| | | | | 7.8 | 6.4 | 6.1 | 6.8 | 8.0 | | | 74.0 | 90.7 | 3.2 | 5.0 | 9.1 | 76.0 | 2.4 | 58.0 | | 63.1 | 20.7 |
| | | | | 5.3 | 5.9 | 6.0 | 14.3 | 13.4 | | | 79.1 | 95.6 | 6.6 | 3.3 | 9.5 | 6.1 | 18.7 | 68.3 | | 76.0 | 26.2 |
| | | Rep 2 | 24-07-14 | 11.2 | 9.0 | 5.4 | | 10.2 | | | 5.6 | 2.5 | 9.7 | 12.0 | 5.7 | 7.0 | 8.7 | 58.6 | | 18.4 | 4.0 |
| | | | | 11.2 | 4.8 | 10.3 | | 11.5 | | | 44.0 | 29.7 | 4.4 | 4.0 | 5.0 | 12.4 | 8.2 | 84.9 | | 18.7 | 2.6 |
| | | | | 13.2 | 2.0 | 5.2 | | 15.4 | | | 13.0 | 23.4 | 3.1 | 4.2 | 6.7 | 2.0 | 1.9 | 84.0 | | 20.5 | 29.4 |
| | | | | 16.2 | 10.9 | 4.3 | | 14.8 | | | 38.5 | 21.3 | 4.4 | 3.9 | 27.0 | 3.4 | 9.2 | 74.4 | | 12.4 | 18.7 |
| | | | | 7.9 | 10.4 | 5.8 | | 6.4 | | | 2.8 | 5.3 | 10.4 | 2.3 | 2.9 | 3.6 | 7.3 | 6.6 | | 9.9 | 2.7 |
| | Distance | Rep 1 | 24-07-11 | 9922.8 | 12537.3 | 10816.4 | 5676.7 | 6190.3 | | | 814.9 | 4831.2 | 10061.6 | 10644.7 | 11914.5 | 11034.9 | 10833.8 | 1066.4 | | 3941.0 | 10535.0 |
| | | | | 6565.8 | 9214.5 | 9182.8 | 7223.4 | 9380.9 | | | 949.3 | 1559.6 | 8999.2 | 13654.7 | 8416.6 | 8177.2 | 13266.4 | 1505.6 | | 4284.1 | 5491.7 |
| | | | | 8873.1 | 7921.7 | 9863.4 | 8158.6 | 10953.9 | | | 877.1 | 12469.9 | 9122.0 | 11399.3 | 16520.4 | 12736.9 | 12983.6 | 1500.5 | | 5041.6 | 16798.3 |
| | | | | 7171.0 | 11837.8 | 8570.5 | 7906.2 | 7583.1 | | | 1095.8 | 1014.0 | 11425.5 | 9621.4 | 10811.1 | 1050.6 | 12700.7 | 2147.3 | | 3289.2 | 8071.2 |
| | | | | 9239.1 | 10127.3 | 10031.2 | 7319.1 | 6999.9 | | | 1306.1 | 615.8 | 11244.2 | 12656.7 | 9801.3 | 10633.3 | 8627.7 | 1283.5 | | 2329.9 | 9241.5 |
| | | Rep 2 | 24-07-14 | 8163.1 | 9507.5 | 9967.1 | | 7497.5 | | | 9506.0 | 13592.8 | 9692.0 | 10859.3 | 14411.3 | 12888.4 | 9709.6 | 3356.4 | | 8260.6 | 13802.4 |
| | | | | 11004.6 | 10953.8 | 8248.0 | | 8045.7 | | | 1680.1 | 6416.4 | 12466.8 | 13005.8 | 11994.8 | 10356.3 | 15317.2 | 1014.8 | | 9895.7 | 21805.5 |
| | | | | 8158.5 | 10476.8 | 10557.7 | | 7723.2 | | | 9509.5 | 12898.1 | 14963.7 | 11415.1 | 14074.5 | 13071.5 | 14257.9 | 954.7 | | 7764.1 | 8877.4 |
| | | | | 7093.5 | 7205.8 | 9809.0 | | 6927.9 | | | 6700.3 | 7662.7 | 9934.6 | 13446.0 | 10640.0 | 12394.7 | 11368.4 | 1834.1 | | 7978.9 | 9141.4 |
| | | | | 10505.5 | 8380.8 | 7319.8 | | 9276.5 | | | 7492.2 | 13081.9 | 12828.6 | 16617.5 | 14116.9 | 11872.1 | 14191.2 | 12403.5 | | 10503.9 | 14023.5 |
| | Velocity | Rep 1 | 24-07-11 | 10.5 | 12.5 | 11.2 | 6.3 | 7.7 | | | 1.7 | 4.6 | 11.5 | 11.7 | 13.7 | 12.7 | 12.8 | 1.6 | | 5.7 | 11.2 |
| | | | | 6.5 | 11.0 | 9.5 | 8.7 | 11.9 | | | 1.8 | 3.0 | 10.4 | 14.8 | 11.0 | 11.1 | 15.7 | 1.8 | | 7.9 | 4.1 |
| | | | | 10.0 | 9.0 | 10.2 | 9.6 | 13.1 | | | 1.7 | 10.6 | 10.9 | 11.9 | 18.0 | 15.4 | 15.7 | 2.3 | | 7.1 | 11.4 |
| | | | | 7.6 | 13.8 | 8.7 | 9.3 | 8.8 | | | 1.8 | 3.0 | 12.8 | 10.3 | 12.1 | 1.8 | 14.4 | 3.1 | | 7.0 | 5.5 |
| | | | | 9.9 | 11.4 | 10.3 | 9.1 | 8.4 | | | 3.7 | 1.6 | 12.6 | 13.7 | 10.9 | 12.2 | 11.5 | 1.8 | | 7.0 | 5.5 |
| | | Rep 2 | 24-07-14 | 9.2 | 11.2 | 10.1 | | 8.2 | | | 9.9 | 11.3 | 11.9 | 13.4 | 16.0 | 14.7 | 10.6 | 7.0 | | 9.2 | 11.5 |
| | | | | 13.0 | 12.4 | 8.7 | | 9.1 | | | 2.0 | 3.6 | 14.5 | 14.8 | 13.1 | 12.3 | 17.1 | 3.0 | | 11.2 | 18.9 |
| | | | | 9.9 | 11.3 | 10.7 | | 9.2 | | | 10.6 | 7.3 | 17.5 | 12.9 | 15.7 | 14.1 | 14.9 | 2.1 | | 8.8 | 5.9 |
| | | | | 8.6 | 8.5 | 9.9 | | 8.0 | | | 10.4 | 3.9 | 11.4 | 15.1 | 15.1 | 13.5 | 12.7 | 4.5 | | 8.3 | 6.3 |
| | | | | 11.4 | 10.0 | 7.1 | | 8.6 | | | 7.5 | 9.7 | 15.6 | 18.6 | 15.2 | 13.0 | 15.5 | 13.1 | | 10.7 | 12.1 |

| | | | | Females | | | | | | | | | Males | | | | | | | | |
|---|---|---|---|---|---|---|---|---|---|---|---|---|---|---|---|---|---|---|---|---|---|
| | | | Date | Age (days) | | | | | | | | | Age (days) | | | | | | | | |
| Genotype | Parameter | Rep | YYMMDD | 2 | 4 | 8 | 15 | 22 | 29 | 30 | 36 | 43 | 2 | 4 | 8 | 15 | 22 | 29 | 30 | 36 | 43 |
| *DJ694*[L] | %Immobile | Rep 1 | 24-07-11 | 7.6 | 14.4 | 14.6 | 15.9 | 28.3 | | | 82.6 | 92.0 | 12.8 | 11.6 | 7.8 | 25.6 | 10.0 | 32.0 | | 31.3 | 83.0 |
| | | | | 12.8 | 9.3 | 8.5 | 69.9 | 78.5 | | | 88.6 | 24.7 | 9.1 | 9.1 | 14.8 | 4.8 | 8.6 | 76.8 | | 65.8 | 19.7 |
| | | | | 56.6 | 9.5 | 16.6 | 17.9 | 70.1 | | | 70.3 | 93.6 | 9.8 | 4.4 | 5.6 | 12.2 | 6.5 | 68.3 | | 88.7 | 40.0 |
| | | | | 21.7 | 7.6 | 11.8 | 91.0 | 19.6 | | | 80.1 | 50.2 | 4.8 | 7.4 | 7.5 | 22.9 | 12.6 | 52.5 | | 75.5 | 21.1 |
| | | | | 5.5 | 19.7 | 29.8 | 11.5 | 13.8 | | | 84.6 | 89.7 | 57.7 | 25.6 | 22.9 | 12.6 | 5.7 | 48.1 | | 77.5 | 22.4 |
| | | Rep 2 | 24-07-14 | 45.6 | 7.7 | 11.4 | | 39.0 | | | 6.5 | 53.7 | 13.2 | 12.0 | 5.1 | 62.6 | 86.5 | 75.1 | | 10.5 | 23.7 |
| | | | | 4.8 | 11.0 | 8.4 | | 24.3 | | | 96.4 | 28.9 | 19.3 | 24.7 | 19.4 | 8.9 | 27.8 | 88.4 | | 22.0 | 38.6 |
| | | | | 14.7 | 15.0 | 16.9 | | 28.1 | | | 78.8 | 39.1 | 15.8 | 4.3 | 8.9 | 1.3 | 40.4 | 90.1 | | 26.8 | 24.6 |
| | | | | 10.2 | 36.1 | 9.4 | | 19.3 | | | 88.7 | 25.4 | 6.4 | 68.5 | 15.7 | 76.9 | 45.9 | 13.5 | | 11.5 | 36.5 |
| | | | | 4.3 | 13.8 | 11.6 | | 13.4 | | | 87.9 | 24.4 | 15.2 | 8.7 | 19.0 | 26.3 | 15.2 | 71.0 | | 16.4 | 15.3 |
| | | Rep 3 | 23-03-09 | | | 48.1 | 3.3 | 5.5 | | 4.7 | | | | | 77.5 | 4.3 | 12.8 | | 15.3 | | |
| | | | | | | 13.7 | 6.3 | 19.9 | | 70.6 | | | | | 21.2 | 6.3 | 13.9 | | 10.7 | | |
| | | | | | | 41.6 | 2.6 | 7.1 | | 57.2 | | | | | 37.2 | 3.1 | 8.6 | | 23.1 | | |
| | | | | | | 27.3 | 4.6 | 4.5 | | 41.0 | | | | | 38.7 | 38.6 | 7.4 | | 65.5 | | |
| | | | | | | 16.6 | 1.2 | 2.3 | | 17.9 | | | | | 21.9 | 35.1 | 5.6 | | 62.8 | | |
| | | Rep 4 | 23-03-13 | | | 6.6 | 12.3 | 11.5 | | 7.0 | | | | | 36.2 | 33.8 | 30.5 | | 22.0 | | |
| | | | | | | 4.4 | 13.4 | 11.1 | | 14.4 | | | | | 7.4 | 13.9 | 82.5 | | 49.3 | | |
| | | | | | | 6.1 | 41.0 | 13.4 | | 30.8 | | | | | 10.1 | 6.5 | 1.8 | | 6.9 | | |
| | | | | | | 14.6 | 11.2 | 26.4 | | 37.6 | | | | | 31.0 | 15.4 | 7.0 | | 35.2 | | |
| | | | | | | 10.7 | 41.6 | 55.9 | | 41.0 | | | | | 12.3 | 14.9 | 40.8 | | 38.8 | | |
| | | Rep 5 | 23-03-13 | | | 14.7 | 4.7 | 4.9 | | 7.1 | | | | | 3.4 | 8.7 | 27.2 | | 34.1 | | |
| | | | | | | 5.1 | 12.1 | 2.8 | | 12.9 | | | | | 7.7 | 9.0 | 4.4 | | 0.9 | | |
| | | | | | | 21.9 | 6.6 | 6.9 | | 27.7 | | | | | 8.6 | 16.6 | 10.5 | | 24.5 | | |
| | | | | | | 22.0 | 5.5 | 24.6 | | 11.1 | | | | | 74.6 | 10.1 | 32.7 | | 14.7 | | |
| | | | | | | 15.5 | 6.7 | 21.7 | | 2.1 | | | | | 24.5 | 29.9 | 21.6 | | 2.1 | | |
| | | Rep 6 | 23-03-15 | | | 6.2 | 4.1 | 8.0 | | 5.5 | | | | | 76.1 | 14.4 | 14.7 | | 81.8 | | |
| | | | | | | 28.5 | 7.1 | 10.9 | | 9.4 | | | | | 23.7 | 31.5 | 30.2 | | 4.9 | | |
| | | | | | | 20.9 | 48.5 | 10.3 | | 1.2 | | | | | 96.4 | 10.7 | 7.4 | | 76.8 | | |
| | | | | | | 19.6 | 5.3 | 11.4 | | 3.7 | | | | | 14.5 | 7.7 | 12.0 | | 16.2 | | |
| | | | | | | 7.4 | 8.2 | 5.5 | | 22.7 | | | | | 96.5 | 7.8 | 15.4 | | 94.1 | | |

| | | | | Females | | | | | | | | | Males | | | | | | | | |
|---|---|---|---|---|---|---|---|---|---|---|---|---|---|---|---|---|---|---|---|---|---|
| | | | Date | Age (days) | | | | | | | | | Age (days) | | | | | | | | |
| Genotype | Parameter | Rep | YYMMDD | 2 | 4 | 8 | 15 | 22 | 29 | 30 | 36 | 43 | 2 | 4 | 8 | 15 | 22 | 29 | 30 | 36 | 43 |
| $DJ694^L$ | Distance | Rep 1 | 24-07-11 | 9364.2 | 5461.7 | 6698.0 | 4896.3 | 5356.9 | | | 949.7 | 951.3 | 7338.9 | 7673.5 | 7232.8 | 5801.1 | 8828.2 | 6674.2 | | 4374.4 | 1329.5 |
| | | | | 7521.1 | 7139.0 | 9328.9 | 1229.4 | 981.0 | | | 663.1 | 5546.6 | 7549.8 | 8379.6 | 9397.5 | 8982.2 | 9434.3 | 1027.2 | | 1385.5 | 8224.2 |
| | | | | 2496.1 | 5755.4 | 6707.9 | 5493.6 | 1030.4 | | | 1199.0 | 684.2 | 7523.8 | 8598.6 | 10298.3 | 6389.8 | 13643.0 | 1392.8 | | 847.4 | 5570.3 |
| | | | | 6362.6 | 7946.3 | 7897.9 | 534.8 | 5644.6 | | | 1027.7 | 3303.2 | 11096.2 | 11238.4 | 9179.9 | 6269.8 | 7329.3 | 3043.6 | | 1144.0 | 8803.6 |
| | | | | 7987.5 | 6696.7 | 6572.3 | 10520.7 | 4881.3 | | | 771.1 | 749.7 | 3063.6 | 6859.9 | 10325.7 | 9418.3 | 9004.9 | 2388.1 | | 1141.7 | 9812.6 |
| | | Rep 2 | 24-07-14 | 4485.9 | 8836.3 | 8100.9 | | 3885.5 | | | 6666.2 | 3453.5 | 8994.7 | 8527.3 | 10113.4 | 2345.6 | 850.5 | 1131.8 | | 6098.1 | 8232.3 |
| | | | | 10350.4 | 8693.2 | 8462.0 | | 4013.2 | | | 331.0 | 5924.4 | 6663.1 | 4765.1 | 7970.1 | 7171.6 | 8244.8 | 757.3 | | 6468.1 | 4367.6 |
| | | | | 7464.8 | 6375.8 | 6912.7 | | 4736.1 | | | 968.0 | 7211.0 | 8171.0 | 11704.0 | 7888.7 | 10461.7 | 2116.3 | 704.7 | | 5171.6 | 9654.2 |
| | | | | 7609.0 | 5880.1 | 7842.2 | | 5296.7 | | | 565.0 | 8339.2 | 10642.8 | 1742.6 | 7418.1 | 1062.7 | 3343.0 | 9009.3 | | 6811.7 | 6639.5 |
| | | | | 9630.8 | 7672.8 | 7663.2 | | 7124.1 | | | 689.8 | 8070.7 | 8457.2 | 10718.1 | 7085.5 | 4342.4 | 8264.8 | 1425.3 | | 3976.7 | 10957.4 |
| | | Rep 3 | 23-03-09 | | | 4957.8 | 14941.5 | 12398.2 | | 11812.7 | | | | | 1154.4 | 18919.9 | 14396.0 | | 13447.7 | | |
| | | | | | | 9759.6 | 14985.7 | 7862.7 | | 1741.5 | | | | | 7227.9 | 17419.3 | 15055.7 | | 15352.4 | | |
| | | | | | | 4817.8 | 14990.9 | 11775.8 | | 2742.4 | | | | | 4904.3 | 15616.3 | 17568.0 | | 10052.9 | | |
| | | | | | | 7511.6 | 13813.6 | 12030.9 | | 4360.2 | | | | | 5035.1 | 5347.5 | 16508.3 | | 2304.8 | | |
| | | | | | | 8811.0 | 14895.8 | 14466.4 | | 8063.8 | | | | | 9165.9 | 5807.5 | 17903.8 | | 3675.2 | | |
| | | Rep 4 | 23-03-13 | | | 13951.4 | 9247.0 | 13378.2 | | 13906.8 | | | | | 5020.8 | 9544.1 | 9406.2 | | 8296.9 | | |
| | | | | | | 12725.5 | 8533.2 | 9545.7 | | 11559.9 | | | | | 11754.5 | 13107.7 | 1298.9 | | 4710.7 | | |
| | | | | | | 12479.0 | 6510.4 | 7484.2 | | 7046.5 | | | | | 13362.5 | 17539.2 | 17690.7 | | 13334.2 | | |
| | | | | | | 11768.6 | 5457.9 | 7454.3 | | 5337.2 | | | | | 7573.0 | 14324.8 | 16236.9 | | 7210.5 | | |
| | | | | | | 11084.7 | 5408.8 | 3607.2 | | 5002.2 | | | | | 11468.1 | 14615.6 | 4308.2 | | 6299.4 | | |
| | | Rep 5 | 23-03-13 | | | 11996.1 | 12763.0 | 12991.6 | | 15112.5 | | | | | 15176.3 | 15990.5 | 11072.5 | | 5927.6 | | |
| | | | | | | 11015.4 | 13213.0 | 21415.6 | | 8847.4 | | | | | 19465.6 | 14797.6 | 13643.9 | | 13699.0 | | |
| | | | | | | 7736.2 | 12318.4 | 11755.5 | | 4407.6 | | | | | 11450.7 | 10111.9 | 14898.8 | | 7887.7 | | |
| | | | | | | 8131.5 | 12793.5 | 7565.6 | | 12069.5 | | | | | 1748.4 | 16437.4 | 5534.6 | | 13619.3 | | |
| | | | | | | 9240.7 | 12004.5 | 11519.4 | | 16095.6 | | | | | 9265.7 | 8549.0 | 11614.8 | | 15239.4 | | |
| | | Rep 6 | 23-03-15 | | | 14085.1 | 13729.4 | 12160.7 | | 13090.0 | | | | | 1254.1 | 12686.2 | 10980.9 | | 1078.0 | | |
| | | | | | | 6880.4 | 14170.6 | 11404.1 | | 13451.7 | | | | | 6724.1 | 10893.4 | 10448.2 | | 11249.1 | | |
| | | | | | | 7653.9 | 2269.8 | 8329.1 | | 21341.8 | | | | | 502.7 | 11976.3 | 11756.9 | | 3818.0 | | |
| | | | | | | 10290.2 | 14601.4 | 10863.0 | | 21321.6 | | | | | 14932.2 | 17130.5 | 14814.4 | | 8024.5 | | |
| | | | | | | 13146.7 | 8943.1 | 14803.5 | | 11767.4 | | | | | 512.9 | 14954.6 | 14103.5 | | 652.1 | | |

| | | | | Females | | | | | | | | | Males | | | | | | | | |
|---|---|---|---|---|---|---|---|---|---|---|---|---|---|---|---|---|---|---|---|---|---|
| | | | Date | Age (days) | | | | | | | | | Age (days) | | | | | | | | |
| Genotype | Parameter | Rep | YYMMDD | 2 | 4 | 8 | 15 | 22 | 29 | 30 | 36 | 43 | 2 | 4 | 8 | 15 | 22 | 29 | 30 | 36 | 43 |
| $DJ694^{L}$/+ | Velocity | Rep 1 | 24-07-11 | 9.9 | 6.7 | 7.0 | 5.3 | 7.9 | | | 1.8 | 3.1 | 8.9 | 8.9 | 7.8 | 8.1 | 10.8 | 10.2 | | 5.2 | 1.6 |
| | | | | 7.2 | 8.4 | 9.8 | 2.7 | 1.7 | | | 1.5 | 4.7 | 8.9 | 9.7 | 11.2 | 10.3 | 11.4 | 1.6 | | 2.2 | 5.3 |
| | | | | 3.6 | 6.7 | 6.7 | 7.2 | 1.8 | | | 1.8 | 1.7 | 8.9 | 9.5 | 11.2 | 7.7 | 16.4 | 1.7 | | 1.8 | 3.8 |
| | | | | 8.7 | 9.4 | 8.3 | 1.6 | 7.4 | | | 1.8 | 4.8 | 12.8 | 12.9 | 10.1 | 8.3 | 9.1 | 5.1 | | 2.0 | 4.3 |
| | | | | 8.4 | 8.2 | 8.3 | 13.4 | 5.4 | | | 1.7 | 1.9 | 3.5 | 9.7 | 13.8 | 11.3 | 10.4 | 3.4 | | 2.1 | 5.9 |
| | | Rep 2 | 24-07-14 | 7.6 | 10.2 | 8.6 | | 6.2 | | | 6.8 | 3.0 | 11.6 | 10.7 | 11.1 | 5.2 | 2.6 | 2.0 | | 6.0 | 4.4 |
| | | | | 11.5 | 10.1 | 8.7 | | 5.1 | | | 1.4 | 5.0 | 9.1 | 6.3 | 10.3 | 8.2 | 11.3 | 1.4 | | 7.4 | 2.8 |
| | | | | 9.1 | 7.4 | 7.7 | | 6.3 | | | 2.6 | 5.4 | 10.9 | 13.6 | 9.1 | 11.3 | 2.5 | 2.1 | | 6.2 | 5.3 |
| | | | | 8.8 | 9.6 | 8.1 | | 6.3 | | | 1.6 | 4.7 | 13.0 | 3.5 | 9.0 | 1.6 | 5.5 | 10.3 | | 7.0 | 4.6 |
| | | | | 10.5 | 9.6 | 7.8 | | 6.1 | | | 1.6 | 4.4 | 8.8 | 12.3 | 8.9 | 5.6 | 9.7 | 2.9 | | 3.8 | 8.8 |
| | | Rep 3 | 23-03-09 | | | 7.2 | 11.9 | 11.7 | | 10.9 | | | | | 2.1 | 17.4 | 14.4 | | 13.9 | | |
| | | | | | | 9.0 | 12.6 | 8.3 | | 2.6 | | | | | 7.2 | 16.0 | 15.5 | | 15.0 | | |
| | | | | | | 5.9 | 9.7 | 11.0 | | 3.6 | | | | | 6.0 | 13.0 | 17.3 | | 11.3 | | |
| | | | | | | 8.0 | 11.4 | 10.6 | | 5.7 | | | | | 6.3 | 5.4 | 15.9 | | 3.7 | | |
| | | | | | | 8.4 | 11.9 | 13.1 | | 8.1 | | | | | 9.3 | 5.9 | 17.3 | | 6.6 | | |
| | | Rep 4 | 23-03-13 | | | 8.6 | 9.4 | 12.6 | | 8.3 | | | | | 3.8 | 12.4 | 11.0 | | 9.8 | | |
| | | | | | | 10.7 | 11.0 | 8.2 | | 8.1 | | | | | 10.1 | 13.4 | 1.6 | | 8.0 | | |
| | | | | | | 10.0 | 10.4 | 6.3 | | 6.7 | | | | | 11.7 | 16.7 | 15.5 | | 13.4 | | |
| | | | | | | 11.3 | 13.4 | 7.9 | | 5.8 | | | | | 7.2 | 15.0 | 14.8 | | 10.2 | | |
| | | | | | | 8.7 | 7.5 | 3.3 | | 4.7 | | | | | 8.5 | 15.1 | 2.8 | | 9.1 | | |
| | | Rep 5 | 23-03-13 | | | 11.7 | 11.7 | 11.6 | | 10.5 | | | | | 13.0 | 15.4 | 12.6 | | 5.8 | | |
| | | | | | | 10.0 | 13.0 | 18.9 | | 5.5 | | | | | 17.6 | 14.3 | 12.1 | | 11.1 | | |
| | | | | | | 8.2 | 11.1 | 10.0 | | 3.9 | | | | | 10.9 | 10.3 | 14.1 | | 7.7 | | |
| | | | | | | 8.2 | 11.8 | 7.5 | | 10.1 | | | | | 3.0 | 16.0 | 3.4 | | 13.5 | | |
| | | | | | | 8.8 | 11.1 | 11.7 | | 10.7 | | | | | 10.0 | 10.1 | 11.9 | | 12.7 | | |
| | | Rep 6 | 23-03-15 | | | 12.9 | 12.2 | 11.2 | | 9.8 | | | | | 2.0 | 12.5 | 10.8 | | 1.8 | | |
| | | | | | | 7.8 | 12.9 | 10.6 | | 6.4 | | | | | 7.3 | 13.4 | 12.4 | | 9.7 | | |
| | | | | | | 8.0 | 2.8 | 5.1 | | 12.6 | | | | | 1.7 | 11.4 | 10.4 | | 12.3 | | |
| | | | | | | 10.6 | 13.1 | 8.8 | | 9.7 | | | | | 15.0 | 16.1 | 14.4 | | 7.8 | | |
| | | | | | | 12.2 | 8.1 | 11.7 | | 8.0 | | | | | 2.4 | 13.9 | 13.5 | | 1.5 | | |

| | | | Date | Females | | | | | | | | | Males | | | | | | | | |
|---|---|---|---|---|---|---|---|---|---|---|---|---|---|---|---|---|---|---|---|---|---|
| | | | | Age (days) | | | | | | | | | Age (days) | | | | | | | | |
| Genotype | Parameter | Rep | YYMMDD | 2 | 4 | 8 | 15 | 22 | 29 | 30 | 36 | 43 | 2 | 4 | 8 | 15 | 22 | 29 | 30 | 36 | 43 |
| *DJ694*$^{L}$/+ | %Immobile | Rep 1 | 24-07-11 | 6.9 | 7.3 | 2.2 | 70.5 | 11.5 | | | 92.2 | 11.5 | 10.9 | 12.0 | 14.2 | 6.6 | 2.9 | 68.1 | | 10.4 | 4.5 |
| | | | | 17.1 | 11.8 | 4.2 | 3.9 | 1.3 | | | 53.8 | 71.9 | 9.6 | 2.7 | 1.1 | 2.6 | 7.6 | 62.0 | | 51.5 | 3.0 |
| | | | | 17.9 | 9.2 | 5.5 | 4.4 | 67.3 | | | 81.1 | 25.8 | 8.7 | 3.0 | 1.5 | 3.1 | 6.6 | 43.5 | | 92.0 | 18.1 |
| | | | | 8.4 | 12.1 | 2.9 | 3.4 | 10.8 | | | 48.3 | 33.8 | 3.5 | 6.4 | 28.1 | 6.8 | 7.1 | 16.1 | | 59.4 | 1.4 |
| | | | | 6.1 | 5.5 | 21.0 | 3.2 | 8.0 | | | 42.2 | 11.0 | 6.2 | 6.5 | 28.0 | 2.4 | 12.7 | 3.8 | | 76.6 | 34.0 |
| | | Rep 2 | 24-07-14 | 20.3 | 16.6 | 5.0 | | 22.3 | | | 5.1 | 16.9 | 9.1 | 13.8 | 3.6 | 2.2 | 8.1 | 51.8 | | 45.3 | 43.4 |
| | | | | 12.1 | 10.2 | 1.8 | | 13.4 | | | 2.1 | 3.2 | 36.0 | 9.7 | 5.5 | 2.3 | 2.3 | 78.6 | | 3.3 | 1.2 |
| | | | | 19.0 | 9.9 | 5.1 | | 12.0 | | | 85.7 | 52.2 | 9.0 | 8.0 | 3.1 | 5.7 | 7.5 | 73.0 | | 10.3 | 2.6 |
| | | | | 11.9 | 17.5 | 8.9 | | 6.8 | | | 22.1 | 1.6 | 8.0 | 5.7 | 1.7 | 5.9 | 13.6 | 79.7 | | 8.4 | 7.3 |
| | | | | 6.8 | 12.6 | 4.2 | | 6.2 | | | 16.2 | 1.7 | 12.4 | 7.7 | 7.5 | 3.4 | 1.8 | 52.0 | | 11.3 | 14.1 |
| | | Rep 3 | 23-03-09 | | | 8.5 | 0.4 | 17.4 | | 6.0 | | | | | 6.1 | 5.1 | 6.7 | | 20.1 | | |
| | | | | | | 15.3 | 5.9 | 11.1 | | 20.0 | | | | | 40.9 | 6.6 | 5.1 | | 11.9 | | |
| | | | | | | 24.5 | 16.7 | 2.4 | | 2.4 | | | | | 9.2 | 3.9 | 4.2 | | 4.4 | | |
| | | | | | | 2.8 | 1.3 | 2.9 | | 60.0 | | | | | 10.2 | 3.5 | 3.4 | | 50.6 | | |
| | | | | | | 7.7 | 1.8 | 18.5 | | 9.9 | | | | | 51.6 | 31.2 | 9.1 | | 4.4 | | |
| | | Rep 4 | 23-03-13 | | | 0.5 | 16.8 | 12.4 | | 7.0 | | | | | 10.4 | 9.5 | 28.4 | | 71.8 | | |
| | | | | | | 4.0 | 8.1 | 8.1 | | 15.4 | | | | | 6.8 | 11.9 | 44.7 | | 27.2 | | |
| | | | | | | 7.5 | 7.8 | 9.1 | | 11.3 | | | | | 5.4 | 17.8 | 15.9 | | 59.4 | | |
| | | | | | | 9.3 | 14.7 | 29.1 | | 11.2 | | | | | 2.1 | 22.6 | 6.8 | | 20.6 | | |
| | | | | | | 10.7 | 20.0 | 10.6 | | 4.4 | | | | | 3.6 | 8.0 | 10.2 | | 31.7 | | |
| | | Rep 5 | 23-03-13 | | | 6.6 | 3.3 | 74.3 | | 7.3 | | | | | 9.3 | 59.7 | 2.9 | | 13.1 | | |
| | | | | | | 8.2 | 4.6 | 17.1 | | 17.4 | | | | | 7.3 | 7.9 | 8.3 | | 67.0 | | |
| | | | | | | 5.9 | 13.4 | 8.9 | | 24.6 | | | | | 12.0 | 5.4 | 2.6 | | 32.4 | | |
| | | | | | | 7.9 | 5.9 | 4.1 | | 12.0 | | | | | 7.0 | 41.5 | 15.6 | | 6.7 | | |
| | | | | | | 5.0 | 2.5 | 11.8 | | 47.3 | | | | | 10.9 | 18.6 | 18.7 | | 3.1 | | |
| | | Rep 6 | 23-03-15 | | | 1.6 | 4.2 | 7.5 | | 2.7 | | | | | 15.7 | 16.9 | 2.1 | | 1.9 | | |
| | | | | | | 3.7 | 1.9 | 0.9 | | 6.0 | | | | | 8.6 | 9.4 | 5.5 | | 0.7 | | |
| | | | | | | 6.0 | 3.1 | 7.0 | | 2.0 | | | | | 11.9 | 5.9 | 1.0 | | 5.4 | | |
| | | | | | | 16.0 | 5.1 | 7.5 | | 1.3 | | | | | 34.1 | 12.2 | 6.5 | | 7.2 | | |
| | | | | | | 2.8 | 59.4 | 6.5 | | 3.3 | | | | | 32.5 | 14.5 | 9.6 | | 6.2 | | |

| | | | | Females | | | | | | | | | Males | | | | | | | | |
|---|---|---|---|---|---|---|---|---|---|---|---|---|---|---|---|---|---|---|---|---|---|
| | | | Date | Age (days) | | | | | | | | | Age (days) | | | | | | | | |
| Genotype | Parameter | Rep | YYMMDD | 2 | 4 | 8 | 15 | 22 | 29 | 30 | 36 | 43 | 2 | 4 | 8 | 15 | 22 | 29 | 30 | 36 | 43 |
| $DJ694^{L}/+$ | Distance | Rep 1 | 24-07-11 | 8575.8 | 6980.2 | 9158.8 | 1390.6 | 8070.4 | | | 472.1 | 10749.9 | 9008.7 | 9493.0 | 8293.9 | 13569.7 | 10244.1 | 1415.4 | | 9006.7 | 9760.7 |
| | | | | 5854.2 | 8807.8 | 11234.4 | 8201.9 | 10213.0 | | | 2119.6 | 1596.8 | 9773.7 | 12255.4 | 13315.3 | 14618.3 | 11295.1 | 1583.4 | | 3490.0 | 14124.6 |
| | | | | 6611.2 | 8038.6 | 10724.3 | 10027.3 | 1301.9 | | | 976.3 | 7235.4 | 10162.0 | 11941.0 | 11418.4 | 12593.7 | 14644.1 | 3382.1 | | 618.8 | 10486.5 |
| | | | | 8419.6 | 8109.4 | 10091.4 | 8715.2 | 8672.6 | | | 2850.3 | 5348.7 | 10509.9 | 11741.4 | 7653.8 | 12349.4 | 15859.5 | 10412.1 | | 2596.9 | 17942.9 |
| | | | | 9186.8 | 10236.6 | 6294.8 | 9883.4 | 8234.6 | | | 4916.2 | 9546.7 | 10205.0 | 12843.0 | 6401.5 | 12174.7 | 7354.5 | 16899.0 | | 866.2 | 5289.5 |
| | | Rep 2 | 24-07-14 | 6985.2 | 7761.1 | 10082.1 | | 5083.6 | | | 11068.7 | 6568.0 | 12040.2 | 9722.9 | 13489.0 | 12700.8 | 9449.9 | 3259.5 | | 3737.0 | 4195.9 |
| | | | | 9999.0 | 10665.0 | 10671.6 | | 7312.8 | | | 10640.5 | 9714.7 | 6495.1 | 12099.2 | 13678.3 | 11981.6 | 17131.2 | 1262.4 | | 13216.2 | 15060.8 |
| | | | | 6140.4 | 10363.1 | 11584.3 | | 7151.9 | | | 906.4 | 3724.9 | 9356.6 | 12469.0 | 12200.1 | 10194.3 | 14638.1 | 1536.7 | | 9569.8 | 18389.8 |
| | | | | 9045.3 | 8317.9 | 6638.1 | | 11383.3 | | | 9045.2 | 15201.3 | 7671.5 | 11423.4 | 13843.7 | 13574.2 | 12511.9 | 1075.0 | | 12167.5 | 12599.6 |
| | | | | 10445.4 | 10026.6 | 10790.7 | | 8712.4 | | | 6439.3 | 12269.5 | 10203.1 | 11072.8 | 11295.3 | 11537.0 | 14426.5 | 3299.5 | | 11942.3 | 8811.7 |
| | | Rep 3 | 23-03-09 | | | 11410.3 | 21150.3 | 3863.6 | | 13193.3 | | | | | 12894.6 | 13818.0 | 13218.9 | | 12538.0 | | |
| | | | | | | 14388.2 | 14656.7 | 13585.9 | | 9378.6 | | | | | 8210.9 | 15289.3 | 20893.2 | | 11760.3 | | |
| | | | | | | 9879.0 | 10398.0 | 16255.3 | | 12317.7 | | | | | 8832.9 | 16823.4 | 17340.2 | | 11241.5 | | |
| | | | | | | 15210.5 | 16766.2 | 15107.8 | | 2101.9 | | | | | 9964.9 | 16028.5 | 18258.1 | | 8331.6 | | |
| | | | | | | 12983.0 | 17914.9 | 9919.9 | | 14646.8 | | | | | 6451.3 | 5520.4 | 12690.2 | | 17016.7 | | |
| | | Rep 4 | 23-03-13 | | | 18453.7 | 14749.8 | 14136.5 | | 13814.1 | | | | | 12440.7 | 11050.6 | 8974.7 | | 1625.2 | | |
| | | | | | | 15443.7 | 13635.1 | 11526.7 | | 12180.2 | | | | | 14903.2 | 13766.9 | 7367.6 | | 9093.3 | | |
| | | | | | | 14167.0 | 13251.9 | 11376.0 | | 12160.1 | | | | | 17991.2 | 11553.5 | 10844.9 | | 2395.1 | | |
| | | | | | | 13788.0 | 11187.1 | 9070.1 | | 11910.5 | | | | | 19330.2 | 12624.8 | 13688.8 | | 12725.6 | | |
| | | | | | | 12449.1 | 10149.8 | 15275.4 | | 14486.9 | | | | | 18173.3 | 15605.3 | 13692.6 | | 8794.7 | | |
| | | Rep 5 | 23-03-13 | | | 11597.7 | 13125.3 | 1821.2 | | 14069.6 | | | | | 15128.9 | 3216.7 | 19241.2 | | 8689.3 | | |
| | | | | | | 9933.1 | 15294.3 | 9785.5 | | 6101.0 | | | | | 14561.3 | 14925.8 | 14854.8 | | 1162.3 | | |
| | | | | | | 12865.8 | 9571.5 | 12240.7 | | 9427.0 | | | | | 15280.2 | 16447.8 | 19243.2 | | 6426.6 | | |
| | | | | | | 12588.7 | 10708.7 | 13274.4 | | 10273.0 | | | | | 14361.5 | 6756.7 | 15872.1 | | 16046.9 | | |
| | | | | | | 12881.8 | 12047.4 | 11659.2 | | 3415.5 | | | | | 15426.4 | 12745.5 | 11054.3 | | 14757.6 | | |
| | | Rep 6 | 23-03-15 | | | 14656.9 | 12166.7 | 14338.0 | | 12727.4 | | | | | 10488.1 | 9283.9 | 19709.3 | | 13991.9 | | |
| | | | | | | 12267.0 | 12912.5 | 19997.7 | | 12375.5 | | | | | 11376.6 | 13881.9 | 17826.7 | | 20211.5 | | |
| | | | | | | 13732.4 | 11868.9 | 10085.3 | | 20063.7 | | | | | 12271.6 | 19187.0 | 24513.9 | | 14608.2 | | |
| | | | | | | 8403.0 | 12291.0 | 13071.5 | | 22836.5 | | | | | 10020.9 | 11808.3 | 13959.6 | | 13888.4 | | |
| | | | | | | 17316.9 | 1699.9 | 11033.9 | | 20525.2 | | | | | 8956.5 | 13624.2 | 18089.0 | | 13829.2 | | |

| | | | | Females | | | | | | | | | Males | | | | | | | | |
|---|---|---|---|---|---|---|---|---|---|---|---|---|---|---|---|---|---|---|---|---|---|
| | | | Date | Age (days) | | | | | | | | | Age (days) | | | | | | | | |
| Genotype | Parameter | Rep | YYMMDD | 2 | 4 | 8 | 15 | 22 | 29 | 30 | 36 | 43 | 2 | 4 | 8 | 15 | 22 | 29 | 30 | 36 | 43 |
| *DJ694*$^{L}$/+ | Velocity | Rep 1 | 24-07-11 | 8.6 | 8.0 | 8.9 | 2.4 | 9.9 | | | 1.5 | 10.0 | 10.7 | 11.4 | 9.9 | 16.1 | 11.5 | 2.1 | | 8.6 | 7.0 |
| | | | | 5.8 | 10.8 | 10.7 | 9.4 | 11.4 | | | 3.2 | 2.3 | 11.6 | 13.6 | 13.9 | 16.7 | 13.4 | 2.1 | | 5.6 | 11.9 |
| | | | | 8.0 | 9.5 | 10.2 | 11.7 | 2.4 | | | 1.9 | 7.6 | 12.1 | 13.2 | 11.9 | 14.4 | 17.5 | 4.9 | | 1.5 | 9.9 |
| | | | | 10.1 | 9.8 | 9.4 | 9.9 | 10.5 | | | 4.1 | 6.2 | 11.8 | 13.4 | 10.7 | 14.5 | 18.9 | 12.8 | | 4.8 | 15.4 |
| | | | | 9.7 | 11.6 | 5.7 | 11.3 | 9.5 | | | 7.2 | 8.8 | 11.0 | 14.8 | 8.8 | 13.5 | 9.1 | 19.1 | | 2.1 | 3.3 |
| | | Rep 2 | 24-07-14 | 8.6 | 10.0 | 9.1 | | 6.3 | | | 11.5 | 5.0 | 14.9 | 12.3 | 14.4 | 13.6 | 10.2 | 5.7 | | 5.4 | 3.1 |
| | | | | 11.8 | 12.8 | 9.9 | | 8.2 | | | 10.7 | 7.4 | 10.9 | 14.7 | 15.1 | 12.8 | 17.9 | 3.1 | | 12.6 | 12.3 |
| | | | | 7.8 | 12.5 | 11.6 | | 7.9 | | | 2.6 | 2.9 | 11.5 | 14.7 | 12.9 | 11.2 | 15.9 | 3.4 | | 9.7 | 15.7 |
| | | | | 10.6 | 10.9 | 5.6 | | 12.4 | | | 11.2 | 12.8 | 9.2 | 13.3 | 14.6 | 15.2 | 14.6 | 2.1 | | 12.4 | 11.4 |
| | | | | 11.3 | 12.4 | 9.4 | | 7.9 | | | 7.3 | 10.5 | 12.4 | 13.0 | 12.6 | 12.4 | 14.7 | 5.0 | | 12.5 | 6.6 |
| | | Rep 3 | 23-03-09 | | | 10.1 | 16.7 | 3.5 | | 12.1 | | | | | 11.2 | 11.6 | 12.7 | | 13.8 | | |
| | | | | | | 13.8 | 10.0 | 13.8 | | 10.2 | | | | | 11.0 | 12.5 | 19.7 | | 11.6 | | |
| | | | | | | 10.3 | 8.3 | 14.6 | | 10.9 | | | | | 7.8 | 14.3 | 16.3 | | 10.3 | | |
| | | | | | | 12.8 | 13.2 | 14.0 | | 2.0 | | | | | 8.9 | 13.0 | 17.2 | | 14.0 | | |
| | | | | | | 11.4 | 13.9 | 10.6 | | 14.3 | | | | | 10.4 | 3.6 | 12.5 | | 15.9 | | |
| | | Rep 4 | 23-03-13 | | | 15.1 | 15.4 | 13.0 | | 8.0 | | | | | 12.1 | 10.4 | 10.2 | | 4.0 | | |
| | | | | | | 14.3 | 12.9 | 10.2 | | 10.0 | | | | | 13.1 | 13.5 | 10.4 | | 11.6 | | |
| | | | | | | 10.8 | 12.7 | 10.4 | | 6.7 | | | | | 15.5 | 12.1 | 10.3 | | 2.6 | | |
| | | | | | | 11.5 | 11.4 | 10.1 | | 7.2 | | | | | 18.2 | 14.2 | 12.3 | | 15.1 | | |
| | | | | | | 10.1 | 10.6 | 14.3 | | 10.6 | | | | | 16.0 | 14.8 | 13.2 | | 12.0 | | |
| | | Rep 5 | 23-03-13 | | | 10.9 | 11.4 | 2.6 | | 10.1 | | | | | 13.8 | 4.7 | 16.8 | | 6.3 | | |
| | | | | | | 9.2 | 13.9 | 9.7 | | 4.4 | | | | | 12.9 | 13.9 | 13.6 | | 2.2 | | |
| | | | | | | 11.2 | 9.1 | 11.0 | | 8.4 | | | | | 14.3 | 14.9 | 16.9 | | 4.9 | | |
| | | | | | | 11.1 | 9.6 | 11.3 | | 9.0 | | | | | 12.5 | 8.7 | 16.0 | | 18.6 | | |
| | | | | | | 11.0 | 10.0 | 10.8 | | 3.3 | | | | | 14.2 | 13.4 | 11.2 | | 12.2 | | |
| | | Rep 6 | 23-03-15 | | | 12.8 | 10.7 | 13.2 | | 10.0 | | | | | 10.6 | 9.5 | 17.4 | | 11.9 | | |
| | | | | | | 10.9 | 11.2 | 17.4 | | 8.1 | | | | | 10.6 | 13.2 | 16.2 | | 17.4 | | |
| | | | | | | 12.6 | 10.3 | 8.5 | | 11.5 | | | | | 11.9 | 17.8 | 21.4 | | 12.9 | | |
| | | | | | | 8.3 | 10.7 | 11.9 | | 14.4 | | | | | 12.7 | 11.5 | 12.5 | | 12.5 | | |
| | | | | | | 15.2 | 2.0 | 8.1 | | 11.5 | | | | | 11.1 | 13.4 | 17.3 | | 12.4 | | |

| | | | | Females | | | | | | | | | Males | | | | | | | | |
|---|---|---|---|---|---|---|---|---|---|---|---|---|---|---|---|---|---|---|---|---|---|
| | | | Date | Age (days) | | | | | | | | | Age (days) | | | | | | | | |
| Genotype | Parameter | Rep | YYMMDD | 2 | 4 | 8 | 15 | 22 | 29 | 30 | 36 | 43 | 2 | 4 | 8 | 15 | 22 | 29 | 30 | 36 | 43 |
| $DJ694^{O}$ | %Immobile | Rep 1 | 24-07-11 | 18.3 | 6.1 | 34.3 | 9.7 | 41.5 | 87.9 | | 80.5 | 98.2 | 12.8 | 16.2 | 16.1 | 15.1 | 10.9 | 41.3 | | 55.1 | 46.2 |
| | | | | 8.0 | 15.5 | 15.2 | 12.4 | 24.1 | 86.2 | | 85.6 | 90.9 | 13.5 | 9.8 | 22.0 | 20.5 | 8.9 | 49.9 | | 94.5 | 37.1 |
| | | | | 56.9 | 10.7 | 15.4 | 52.0 | 19.8 | 80.8 | | 92.0 | 82.4 | 7.1 | 11.7 | 20.4 | 12.0 | 81.5 | 62.4 | | 26.9 | 47.3 |
| | | | | 20.9 | 14.1 | 13.4 | 24.6 | 17.1 | 27.7 | | 85.5 | 82.5 | 12.1 | 24.7 | 17.3 | 5.6 | 6.9 | 13.3 | | 72.1 | 32.0 |
| | | | | 67.3 | 20.7 | 35.9 | 10.7 | 12.8 | 67.1 | | 71.4 | 43.1 | 15.7 | 8.4 | 10.6 | 7.7 | 24.9 | 75.1 | | 7.7 | 10.4 |
| | | Rep 2 | 24-07-14 | 12.4 | 17.4 | 35.3 | | 43.4 | 53.2 | | 94.2 | 32.8 | 9.9 | 6.3 | 24.3 | 27.8 | 14.2 | 38.9 | | 37.5 | 4.9 |
| | | | | 13.2 | 10.6 | 22.3 | | 19.0 | 86.9 | | 80.4 | 43.1 | 16.5 | 42.3 | 21.2 | 14.8 | 9.3 | 84.2 | | 26.3 | 11.0 |
| | | | | 27.8 | 8.6 | 13.8 | | 36.2 | 42.4 | | 94.3 | 50.5 | 5.9 | 30.2 | 27.3 | 17.1 | 13.7 | 51.7 | | 15.0 | 28.4 |
| | | | | 8.4 | 9.4 | 26.7 | | 58.7 | 70.2 | | 79.4 | 42.7 | 13.9 | 15.0 | 33.4 | 21.2 | 31.8 | 25.3 | | 19.9 | 17.1 |
| | | | | 12.7 | 7.0 | 11.9 | | 57.5 | 20.6 | | 88.1 | 47.2 | 4.2 | 3.5 | 40.2 | 11.1 | 1.5 | 37.5 | | 10.8 | 9.3 |
| | Distance | Rep 1 | 24-07-11 | 7076.4 | 7193.1 | 4602.9 | 6533.5 | 1976.8 | 679.9 | | 1002.4 | 503.5 | 6223.8 | 6690.3 | 6250.7 | 8097.3 | 7964.6 | 2715.2 | | 2250.7 | 3578.3 |
| | | | | 7693.8 | 6312.6 | 6460.0 | 5046.3 | 3096.8 | 746.7 | | 881.4 | 793.0 | 6400.9 | 8081.3 | 6278.5 | 6049.1 | 7384.0 | 2014.9 | | 589.4 | 4494.7 |
| | | | | 2322.5 | 5374.9 | 4584.5 | 2024.0 | 2979.3 | 864.2 | | 629.4 | 1701.7 | 7776.3 | 8439.1 | 6120.7 | 6475.5 | 797.9 | 1708.4 | | 5056.1 | 3557.3 |
| | | | | 4947.7 | 6385.4 | 6194.0 | 5219.8 | 4634.1 | 3041.9 | | 839.2 | 1227.1 | 6912.6 | 6282.7 | 7590.4 | 7385.5 | 8453.4 | 6075.5 | | 1524.3 | 5315.3 |
| | | | | 1684.5 | 4377.7 | 4720.5 | 5443.2 | 5484.2 | 1456.5 | | 1639.4 | 2416.3 | 6993.4 | 7231.5 | 7945.2 | 7394.2 | 4745.2 | 1084.2 | | 6071.9 | 11597.7 |
| | | Rep 2 | 24-07-14 | 8078.1 | 6381.6 | 5786.9 | | 1849.1 | 3274.6 | | 469.2 | 5204.0 | 7638.8 | 10987.0 | 7709.8 | 5291.6 | 7024.4 | 4662.7 | | 3662.9 | 17168.2 |
| | | | | 7073.6 | 7699.2 | 6342.5 | | 3329.1 | 928.2 | | 895.0 | 4144.6 | 8985.2 | 5854.4 | 6281.7 | 4430.1 | 8867.0 | 886.3 | | 5188.7 | 8356.6 |
| | | | | 3591.9 | 7097.1 | 6311.9 | | 4238.5 | 3128.5 | | 504.3 | 3377.1 | 11547.3 | 6621.3 | 6356.9 | 8387.1 | 8592.5 | 3135.6 | | 6914.7 | 5649.9 |
| | | | | 8286.7 | 9880.1 | 6827.7 | | 1962.6 | 1477.6 | | 1085.0 | 4245.0 | 7434.1 | 8813.6 | 6087.7 | 5792.3 | 5204.7 | 6861.3 | | 7010.4 | 8134.3 |
| | | | | 8212.6 | 6873.3 | 9757.2 | | 4117.1 | 7336.3 | | 726.5 | 3683.6 | 11495.6 | 14250.6 | 4559.9 | 10561.2 | 14076.7 | 3725.8 | | 8108.1 | 9309.6 |
| | Velocity | Rep 1 | 24-07-11 | 8.3 | 8.2 | 5.9 | 7.5 | 2.7 | 1.6 | | 1.8 | 1.2 | 7.3 | 8.3 | 7.2 | 10.3 | 9.7 | 3.8 | | 3.6 | 2.4 |
| | | | | 6.6 | 7.9 | 7.0 | 6.1 | 4.0 | 1.7 | | 1.5 | 1.5 | 7.7 | 9.4 | 7.6 | 7.9 | 8.8 | 2.7 | | 1.4 | 2.8 |
| | | | | 3.7 | 6.3 | 4.4 | 3.6 | 3.6 | 1.7 | | 1.4 | 3.3 | 9.0 | 10.1 | 7.3 | 7.9 | 2.3 | 2.5 | | 5.7 | 2.2 |
| | | | | 5.6 | 7.9 | 6.0 | 7.4 | 5.8 | 4.2 | | 1.4 | 1.5 | 8.3 | 8.6 | 8.9 | 8.3 | 9.9 | 6.9 | | 3.5 | 2.9 |
| | | | | 2.9 | 5.4 | 5.7 | 6.5 | 6.5 | 3.6 | | 2.4 | 2.8 | 7.7 | 8.2 | 8.7 | 8.1 | 6.5 | 1.6 | | 5.5 | 9.9 |
| | | Rep 2 | 24-07-14 | 9.2 | 8.3 | 6.2 | | 2.3 | 6.0 | | 1.4 | 2.8 | 9.3 | 12.9 | 10.4 | 7.3 | 8.1 | 7.1 | | 5.0 | 15.4 |
| | | | | 8.3 | 9.1 | 5.3 | | 3.7 | 1.9 | | 2.2 | 2.6 | 12.1 | 10.5 | 7.9 | 5.2 | 9.8 | 1.7 | | 5.7 | 5.7 |
| | | | | 4.7 | 8.3 | 5.2 | | 6.1 | 4.5 | | 1.4 | 2.2 | 13.8 | 10.2 | 8.7 | 10.4 | 9.9 | 5.7 | | 7.4 | 3.9 |
| | | | | 9.4 | 12.0 | 7.4 | | 3.5 | 2.3 | | 3.3 | 2.6 | 9.4 | 11.2 | 9.2 | 7.5 | 7.1 | 8.9 | | 7.7 | 6.3 |
| | | | | 8.2 | 8.0 | 7.7 | | 5.1 | 6.6 | | 1.4 | 2.5 | 13.0 | 16.5 | 7.6 | 12.5 | 14.7 | 4.6 | | 8.2 | 6.8 |

| Genotype | Parameter | Rep | YYMMDD Date | Females Age (days) 2 | 4 | 8 | 15 | 22 | 29 | 30 | 36 | 43 | Males Age (days) 2 | 4 | 8 | 15 | 22 | 29 | 30 | 36 | 43 |
|---|---|---|---|---|---|---|---|---|---|---|---|---|---|---|---|---|---|---|---|---|---|
| *DJ694*$^{O}$/+ | %Immobile | Rep 1 | 24-07-11 | 8.2 | 18.1 | 5.8 | 4.9 | 8.5 | 58.9 | | 85.8 | 11.8 | 5.8 | 7.7 | 13.6 | 2.6 | 5.4 | 72.8 | | 72.7 | 9.1 |
| | | | | 4.7 | 10.7 | 20.9 | 4.6 | 5.3 | 9.0 | | 75.2 | 53.1 | 3.5 | 8.1 | 7.8 | 4.0 | 6.4 | 54.3 | | 93.6 | 34.1 |
| | | | | 6.7 | 9.2 | 7.1 | 5.9 | 16.2 | 27.3 | | 93.2 | 9.2 | 8.0 | 9.4 | 9.3 | 2.4 | 3.0 | 62.6 | | 50.8 | 6.9 |
| | | | | 13.7 | 22.4 | 14.7 | 8.8 | 9.1 | 23.9 | | 44.3 | 31.4 | 6.9 | 15.4 | 3.7 | 4.1 | 16.3 | 10.2 | | 59.9 | 40.2 |
| | | | | 5.3 | 15.8 | 8.2 | 4.2 | 6.5 | 87.5 | | 77.8 | 21.9 | 5.1 | 4.7 | 5.0 | 9.1 | 6.5 | 56.8 | | 81.4 | 10.1 |
| | | Rep 2 | 24-07-14 | 7.5 | 4.9 | 6.2 | | 6.9 | 16.7 | | 13.8 | 7.0 | 11.7 | 5.7 | 14.4 | 2.3 | 9.1 | 11.8 | | 69.4 | 5.7 |
| | | | | 21.3 | 9.3 | 15.1 | | 5.3 | 52.6 | | 92.6 | 21.4 | 9.0 | 8.0 | 12.2 | 4.0 | 7.2 | 59.5 | | 53.8 | 2.6 |
| | | | | 14.1 | 16.0 | 7.6 | | 6.8 | 6.6 | | 83.3 | 9.0 | 10.1 | 2.8 | 8.9 | 2.2 | 21.8 | 72.7 | | 73.3 | 7.4 |
| | | | | 19.8 | 11.8 | 16.9 | | 7.5 | 55.9 | | 87.2 | 5.4 | 22.4 | 14.0 | 10.3 | 4.3 | 2.3 | 42.0 | | 59.7 | 26.2 |
| | | | | 7.2 | 13.1 | 8.0 | | 8.1 | 5.6 | | 87.3 | 5.7 | 8.1 | 11.5 | 4.1 | 2.4 | 2.3 | 10.1 | | 18.4 | 2.4 |
| | Distance | Rep 1 | 24-07-11 | 7957.9 | 5255.5 | 9666.4 | 8044.5 | 6055.3 | 2005.4 | | 801.9 | 7549.6 | 9068.0 | 10942.1 | 12673.2 | 11787.0 | 10361.2 | 1189.3 | | 2286.8 | 13969.4 |
| | | | | 10008.6 | 8094.4 | 7312.8 | 7415.6 | 7843.3 | 9846.5 | | 1154.0 | 3319.3 | 10111.7 | 12254.4 | 11534.4 | 13175.6 | 11380.5 | 2061.6 | | 620.4 | 5991.7 |
| | | | | 8085.8 | 8389.6 | 9995.0 | 6850.7 | 7041.0 | 6590.2 | | 488.4 | 9596.9 | 10359.8 | 11223.4 | 11671.8 | 14085.1 | 12059.6 | 1574.4 | | 3986.7 | 12455.1 |
| | | | | 6035.6 | 5387.8 | 8957.0 | 9338.5 | 6824.7 | 4802.4 | | 2674.2 | 6006.8 | 9682.7 | 9257.1 | 12216.6 | 12859.3 | 10002.7 | 8335.9 | | 2597.1 | 5277.2 |
| | | | | 8191.5 | 8875.8 | 9239.1 | 9736.8 | 7944.6 | 698.6 | | 1118.1 | 7676.4 | 11310.4 | 12024.4 | 12045.3 | 10630.7 | 13284.7 | 2064.8 | | 1441.6 | 11794.7 |
| | | Rep 2 | 24-07-14 | 8105.9 | 9546.8 | 9805.9 | | 10378.0 | 5783.3 | | 5777.8 | 11172.7 | 9969.0 | 12429.3 | 9242.0 | 12352.8 | 14678.1 | 12856.0 | | 2227.0 | 12808.4 |
| | | | | 6009.3 | 9141.6 | 6420.9 | | 9938.3 | 2837.4 | | 511.8 | 9738.6 | 9352.4 | 12251.2 | 9442.5 | 11570.0 | 14527.6 | 3643.2 | | 3109.0 | 15028.8 |
| | | | | 6776.7 | 8848.1 | 7873.6 | | 8999.1 | 8070.1 | | 986.0 | 10177.5 | 8982.8 | 12551.6 | 11791.4 | 13479.9 | 11286.6 | 1553.3 | | 1920.6 | 15744.5 |
| | | | | 6745.2 | 9090.3 | 8070.7 | | 8274.5 | 2911.9 | | 965.0 | 13391.1 | 7256.7 | 9152.5 | 11280.4 | 12791.2 | 10994.2 | 5307.6 | | 2721.5 | 8051.4 |
| | | | | 8742.0 | 8267.2 | 8944.6 | | 8686.2 | 9403.3 | | 865.3 | 13132.8 | 11351.4 | 10096.7 | 10644.7 | 11673.7 | 14944.5 | 10226.9 | | 6650.0 | 14086.5 |
| | Velocity | Rep 1 | 24-07-11 | 7.8 | 6.4 | 9.8 | 8.7 | 7.0 | 4.1 | | 1.6 | 4.8 | 10.2 | 12.4 | 14.6 | 13.2 | 11.9 | 1.7 | | 6.1 | 11.9 |
| | | | | 11.2 | 9.7 | 8.7 | 8.4 | 8.8 | 11.8 | | 2.2 | 2.8 | 11.3 | 14.3 | 12.4 | 15.0 | 13.4 | 3.0 | | 1.4 | 3.7 |
| | | | | 8.5 | 9.9 | 10.4 | 7.8 | 8.9 | 9.8 | | 1.6 | 7.1 | 12.1 | 13.3 | 12.7 | 15.7 | 13.7 | 2.0 | | 6.5 | 9.4 |
| | | | | 7.2 | 7.2 | 10.0 | 11.2 | 8.1 | 6.6 | | 3.5 | 3.4 | 11.2 | 11.5 | 12.3 | 14.5 | 13.1 | 9.3 | | 4.8 | 3.8 |
| | | | | 8.2 | 11.3 | 9.4 | 11.0 | 9.0 | 1.6 | | 1.7 | 3.9 | 11.9 | 13.4 | 12.4 | 12.5 | 15.8 | 3.0 | | 4.0 | 7.3 |
| | | Rep 2 | 24-07-14 | 9.0 | 10.8 | 9.7 | | 11.3 | 6.6 | | 6.4 | 6.2 | 12.5 | 14.5 | 11.2 | 13.1 | 16.4 | 14.7 | | 4.6 | 7.2 |
| | | | | 7.7 | 10.9 | 6.8 | | 10.6 | 4.8 | | 1.6 | 5.0 | 11.4 | 14.5 | 11.1 | 12.6 | 15.8 | 7.7 | | 5.4 | 11.3 |
| | | | | 8.1 | 11.5 | 8.1 | | 9.7 | 8.5 | | 3.2 | 5.7 | 11.1 | 14.3 | 13.5 | 14.5 | 14.4 | 3.7 | | 4.9 | 13.0 |
| | | | | 8.6 | 11.1 | 9.0 | | 9.0 | 5.5 | | 4.2 | 7.7 | 10.2 | 11.6 | 13.1 | 14.0 | 11.3 | 8.3 | | 4.4 | 5.1 |
| | | | | 8.7 | 10.3 | 8.3 | | 8.9 | 7.9 | | 3.1 | 6.9 | 13.1 | 12.3 | 11.4 | 12.4 | 15.5 | 10.9 | | 7.2 | 9.9 |

| Genotype | Parameter | Rep | Date YYMMDD | Females | | | | | | | | | Males | | | | | | | | |
|---|---|---|---|---|---|---|---|---|---|---|---|---|---|---|---|---|---|---|---|---|---|
| | | | | Age (days) | | | | | | | | | Age (days) | | | | | | | | |
| | | | | 2 | 4 | 8 | 15 | 22 | 29 | 30 | 36 | 43 | 2 | 4 | 8 | 15 | 22 | 29 | 30 | 36 | 43 |
| *$DJ694^P$* | %Immobile | Rep 1 | 24-07-11 | 9.3 | 74.9 | 16.1 | 14.9 | 29.1 | 68.1 | | 80.5 | 14.7 | 15.3 | 9.8 | 30.1 | 45.7 | 9.4 | 55.9 | | 50.9 | 28.4 |
| | | | | 12.9 | 8.5 | 79.9 | 15.4 | 18.5 | 79.2 | | 84.6 | 20.3 | 17.2 | 21.3 | 81.8 | 23.3 | 16.9 | 17.0 | | 91.8 | 21.9 |
| | | | | 77.2 | 7.5 | 9.5 | 32.5 | 15.4 | 82.3 | | 86.1 | 15.8 | 11.2 | 11.5 | 65.8 | 49.8 | 17.7 | 56.7 | | 36.0 | 52.5 |
| | | | | 72.7 | 14.3 | 15.2 | 14.2 | 17.7 | 73.7 | | 84.8 | 13.6 | 7.9 | 18.0 | 17.4 | 62.7 | 10.5 | 48.2 | | 64.3 | 26.3 |
| | | | | 3.0 | 18.2 | 9.6 | 14.9 | 19.2 | 59.0 | | 77.9 | 67.3 | 10.7 | 13.9 | 12.4 | 7.1 | 10.0 | 68.1 | | 80.6 | 43.2 |
| | | Rep 2 | 24-07-14 | 6.6 | 10.7 | 8.6 | | 83.7 | 88.3 | | 87.4 | 30.8 | 13.5 | 9.9 | 22.7 | 39.6 | 9.3 | 52.2 | | 53.4 | 22.8 |
| | | | | 13.7 | 27.9 | 15.5 | | 36.0 | 65.3 | | 84.8 | 19.8 | 17.9 | 15.3 | 3.6 | 17.3 | 16.6 | 73.8 | | 41.7 | 42.7 |
| | | | | 21.5 | 10.3 | 76.2 | | 12.0 | 44.6 | | 82.7 | 13.2 | 28.7 | 42.6 | 37.8 | 32.6 | 15.2 | 63.8 | | 73.4 | 4.4 |
| | | | | 11.8 | 47.7 | 13.1 | | 18.3 | 32.2 | | 86.6 | 18.7 | 21.1 | 18.3 | 18.3 | 66.1 | 16.6 | 82.1 | | 41.9 | 7.3 |
| | | | | 11.4 | 28.2 | 32.7 | | 40.3 | 26.8 | | 55.3 | 16.5 | 9.5 | 49.0 | 11.9 | 7.5 | 12.0 | 62.2 | | 16.8 | 5.0 |
| | Distance | Rep 1 | 24-07-11 | 8791.3 | 1330.6 | 6832.4 | 4583.1 | 3221.7 | 1227.5 | | 971.4 | 9417.0 | 6430.1 | 8897.9 | 6714.8 | 2502.9 | 6959.8 | 1704.4 | | 2972.0 | 6377.0 |
| | | | | 6949.7 | 6118.1 | 1220.8 | 3879.1 | 5462.8 | 928.2 | | 906.8 | 7993.1 | 6477.5 | 5491.5 | 1035.5 | 5233.0 | 8683.5 | 6258.2 | | 676.7 | 8364.4 |
| | | | | 1149.4 | 8071.1 | 6116.7 | 2753.0 | 4424.3 | 804.2 | | 810.2 | 16838.4 | 6065.4 | 9180.8 | 1509.5 | 2562.4 | 7429.8 | 1967.3 | | 4170.9 | 6192.7 |
| | | | | 1135.8 | 6720.9 | 6275.7 | 3891.7 | 4033.7 | 973.6 | | 930.7 | 11067.2 | 6941.2 | 7398.8 | 8550.6 | 1799.7 | 8946.1 | 2489.2 | | 1613.5 | 7754.3 |
| | | | | 9576.9 | 5897.1 | 6148.9 | 5215.3 | 5929.9 | 1540.3 | | 1091.0 | 2260.5 | 9830.3 | 6990.7 | 7622.2 | 8507.7 | 9415.6 | 1354.2 | | 1176.3 | 8106.0 |
| | | Rep 2 | 24-07-14 | 7976.5 | 6843.5 | 7270.6 | | 890.9 | 819.9 | | 628.9 | 6689.0 | 7618.9 | 9050.5 | 6709.2 | 4465.9 | 8296.5 | 3330.4 | | 3532.1 | 8245.4 |
| | | | | 6070.3 | 5341.1 | 6649.4 | | 2646.1 | 1824.8 | | 798.3 | 11628.8 | 6840.4 | 7059.2 | 7748.9 | 5428.0 | 7125.4 | 1217.9 | | 4075.3 | 3963.8 |
| | | | | 4872.4 | 8330.6 | 1229.8 | | 5889.7 | 3719.4 | | 1503.1 | 15442.6 | 4988.9 | 4776.4 | 5970.2 | 3170.5 | 7682.6 | 2075.0 | | 1475.4 | 15100.6 |
| | | | | 8684.5 | 2363.2 | 7629.5 | | 5501.4 | 3712.9 | | 711.2 | 9338.5 | 6828.3 | 8821.9 | 7309.9 | 1423.9 | 8942.7 | 903.8 | | 4043.2 | 13546.7 |
| | | | | 7223.5 | 6150.5 | 5185.0 | | 4486.4 | 6334.8 | | 2573.0 | 11342.6 | 10854.9 | 2140.6 | 7558.5 | 8920.9 | 7631.7 | 1753.5 | | 7368.7 | 15005.6 |
| | Velocity | Rep 1 | 24-07-11 | 8.0 | 3.3 | 7.7 | 4.3 | 4.4 | 2.1 | | 1.6 | 6.0 | 7.9 | 10.3 | 9.2 | 3.6 | 8.2 | 3.0 | | 4.4 | 3.6 |
| | | | | 6.1 | 7.0 | 2.6 | 4.0 | 7.1 | 1.5 | | 1.5 | 4.0 | 8.3 | 7.1 | 3.3 | 6.8 | 11.3 | 7.5 | | 1.6 | 5.0 |
| | | | | 2.5 | 9.4 | 6.4 | 3.3 | 5.4 | 1.5 | | 1.5 | 8.4 | 7.1 | 10.8 | 2.4 | 4.2 | 9.8 | 3.1 | | 5.3 | 6.2 |
| | | | | 2.5 | 8.4 | 6.9 | 4.4 | 5.0 | 1.9 | | 1.8 | 5.1 | 7.9 | 9.3 | 10.0 | 3.0 | 11.0 | 3.3 | | 3.1 | 4.3 |
| | | | | 10.1 | 7.3 | 6.0 | 5.5 | 7.6 | 2.9 | | 1.9 | 2.1 | 10.8 | 8.4 | 8.4 | 10.1 | 11.4 | 1.8 | | 3.0 | 6.2 |
| | | Rep 2 | 24-07-14 | 8.8 | 8.2 | 7.3 | | 1.7 | 1.5 | | 1.5 | 4.1 | 9.7 | 10.9 | 8.8 | 7.2 | 9.1 | 5.7 | | 6.2 | 4.4 |
| | | | | 7.2 | 7.6 | 7.2 | | 3.5 | 3.5 | | 1.4 | 6.2 | 9.0 | 8.9 | 8.4 | 6.5 | 8.4 | 2.4 | | 5.7 | 2.6 |
| | | | | 6.2 | 10.1 | 2.5 | | 6.5 | 6.0 | | 4.2 | 7.3 | 7.4 | 8.7 | 9.4 | 4.2 | 9.0 | 4.2 | | 2.5 | 12.3 |
| | | | | 10.3 | 3.9 | 8.3 | | 6.4 | 4.7 | | 1.8 | 4.8 | 9.4 | 11.7 | 9.1 | 1.9 | 10.7 | 1.6 | | 5.5 | 10.1 |
| | | | | 7.9 | 8.9 | 5.0 | | 5.2 | 6.8 | | 5.0 | 5.6 | 12.4 | 2.7 | 8.8 | 9.4 | 8.6 | 2.2 | | 7.9 | 13.0 |

| Genotype | Parameter | Rep | YYMMDD Date | Females Age (days) | | | | | | | | | Males Age (days) | | | | | | | | |
|---|---|---|---|---|---|---|---|---|---|---|---|---|---|---|---|---|---|---|---|---|---|
| | | | | 2 | 4 | 8 | 15 | 22 | 29 | 30 | 36 | 43 | 2 | 4 | 8 | 15 | 22 | 29 | 30 | 36 | 43 |
| *DJ694*$^{P}$/+ | %Immobile | Rep 1 | 24-07-11 | 72.2 | 10.5 | 9.5 | 26.4 | 12.2 | 72.5 | | 67.1 | 37.7 | 4.1 | 7.9 | 5.5 | 12.6 | 37.5 | 21.5 | | 3.4 | 10.9 |
| | | | | 5.7 | 18.3 | 8.0 | 11.1 | 5.9 | 86.1 | | 84.7 | 27.6 | 6.9 | 18.1 | 7.1 | 17.1 | 5.7 | 51.0 | | 7.9 | 4.4 |
| | | | | 10.5 | 8.2 | 10.8 | 8.1 | 6.8 | 82.6 | | 74.3 | 7.9 | 4.6 | 6.7 | 19.5 | 11.2 | 6.4 | 62.1 | | 90.8 | 5.9 |
| | | | | 17.2 | 2.9 | 15.1 | 12.7 | 5.3 | 80.5 | | 88.8 | 34.6 | 3.1 | 10.5 | 8.0 | 8.9 | 6.2 | 58.3 | | 63.3 | 13.7 |
| | | | | 25.4 | 11.0 | 11.9 | 7.3 | 4.6 | 85.6 | | 91.5 | 72.0 | 6.4 | 9.5 | 9.1 | 2.9 | 11.3 | 63.7 | | 84.8 | 25.8 |
| | | Rep 2 | 24-07-14 | 20.3 | 13.7 | 12.4 | | 73.2 | 20.0 | | 6.7 | 29.3 | 9.9 | 16.2 | 10.8 | 6.6 | 1.7 | 4.8 | | 17.4 | 9.6 |
| | | | | 12.9 | 6.5 | 8.6 | | 1.4 | 13.3 | | 44.2 | 6.0 | 17.5 | 19.1 | 5.8 | 11.5 | 1.3 | 52.6 | | 43.9 | 0.4 |
| | | | | 6.8 | 15.1 | 16.1 | | 2.9 | 63.6 | | 20.1 | 13.1 | 7.0 | 33.7 | 18.5 | 14.9 | 2.1 | 48.5 | | 84.1 | 5.8 |
| | | | | 7.3 | 8.7 | 9.7 | | 5.4 | 6.9 | | 87.9 | 39.8 | 22.9 | 32.4 | 18.7 | 13.6 | 4.1 | 1.8 | | 19.5 | 2.4 |
| | | | | 11.2 | 16.3 | 11.1 | | 10.0 | 2.7 | | 15.9 | 15.1 | 5.9 | 12.5 | 9.8 | 4.2 | 6.5 | 56.6 | | 2.7 | 2.1 |
| | Distance | Rep 1 | 24-07-11 | 606.9 | 7321.8 | 6229.0 | 5507.6 | 6725.1 | 1247.7 | | 1562.6 | 5258.3 | 10323.6 | 10926.1 | 12027.6 | 10268.0 | 3348.0 | 6891.5 | | 12074.4 | 13238.0 |
| | | | | 8371.2 | 7192.1 | 9044.2 | 8336.5 | 6297.3 | 715.1 | | 789.5 | 7245.3 | 12187.9 | 9647.1 | 13371.4 | 10677.4 | 11608.4 | 3364.9 | | 10629.9 | 16733.8 |
| | | | | 8656.2 | 8019.5 | 9061.2 | 8576.8 | 8965.3 | 872.1 | | 1071.4 | 12158.4 | 12203.7 | 12824.5 | 6035.1 | 4947.9 | 11711.1 | 1666.1 | | 857.6 | 11417.1 |
| | | | | 7851.8 | 10756.8 | 8666.0 | 8076.9 | 9210.3 | 894.1 | | 766.7 | 6094.5 | 10832.0 | 8983.2 | 11004.5 | 12575.2 | 11845.2 | 2031.9 | | 2609.3 | 12873.0 |
| | | | | 4845.8 | 7275.7 | 8407.4 | 7815.8 | 8046.9 | 731.7 | | 603.1 | 2128.0 | 9905.3 | 11146.4 | 11625.5 | 11886.2 | 11681.9 | 1514.4 | | 1268.1 | 7464.4 |
| | | Rep 2 | 24-07-14 | 7252.7 | 8913.7 | 6722.8 | | 1156.1 | 6527.4 | | 11089.1 | 6276.7 | 10675.0 | 10064.4 | 10812.2 | 10818.2 | 15289.6 | 14022.2 | | 9783.9 | 13205.6 |
| | | | | 8723.5 | 9039.7 | 9778.8 | | 9653.8 | 7393.8 | | 3590.4 | 11247.3 | 9095.0 | 7720.5 | 10491.9 | 10287.4 | 14810.3 | 4253.1 | | 4538.9 | 21949.2 |
| | | | | 9717.7 | 7200.7 | 7236.8 | | 9787.1 | 2753.4 | | 5417.2 | 8214.0 | 11192.9 | 5811.9 | 9813.8 | 7420.9 | 17654.7 | 3827.8 | | 1173.5 | 15042.8 |
| | | | | 9479.4 | 9535.4 | 8581.1 | | 10539.7 | 9916.8 | | 729.0 | 5262.1 | 7527.0 | 8146.6 | 12323.0 | 10634.1 | 12148.6 | 12969.4 | | 8508.7 | 14535.6 |
| | | | | 7447.8 | 6956.7 | 7563.8 | | 10862.1 | 11892.1 | | 5095.6 | 9083.4 | 10656.8 | 9317.3 | 13556.9 | 11328.6 | 9512.2 | 2515.5 | | 11908.4 | 17528.6 |
| | Velocity | Rep 1 | 24-07-11 | 3.5 | 8.4 | 6.4 | 7.4 | 8.0 | 2.1 | | 3.3 | 3.5 | 11.4 | 12.5 | 12.6 | 12.9 | 5.1 | 9.1 | | 10.9 | 12.0 |
| | | | | 8.2 | 9.3 | 9.4 | 10.2 | 7.0 | 1.7 | | 1.8 | 4.4 | 14.2 | 12.1 | 14.1 | 14.2 | 13.4 | 6.0 | | 10.0 | 14.6 |
| | | | | 10.2 | 9.3 | 9.8 | 10.2 | 10.3 | 2.6 | | 1.8 | 5.9 | 13.9 | 14.6 | 7.2 | 5.3 | 13.7 | 2.5 | | 3.2 | 8.8 |
| | | | | 10.2 | 12.1 | 9.8 | 10.2 | 10.6 | 2.6 | | 1.5 | 3.9 | 12.0 | 10.4 | 11.7 | 15.1 | 13.8 | 3.2 | | 5.3 | 12.2 |
| | | | | 6.1 | 8.3 | 9.0 | 9.1 | 8.9 | 2.2 | | 1.5 | 2.6 | 10.7 | 13.0 | 12.6 | 13.4 | 14.4 | 2.1 | | 4.6 | 4.9 |
| | | Rep 2 | 24-07-14 | 8.9 | 11.1 | 7.0 | | 1.7 | 7.9 | | 11.8 | 4.7 | 13.2 | 14.7 | 12.4 | 12.1 | 15.7 | 14.7 | | 10.8 | 11.9 |
| | | | | 10.3 | 11.0 | 10.3 | | 9.9 | 8.2 | | 5.7 | 9.2 | 12.1 | 12.0 | 11.5 | 12.2 | 15.3 | 8.1 | | 6.8 | 18.8 |
| | | | | 10.8 | 8.9 | 7.9 | | 10.1 | 5.9 | | 6.4 | 4.9 | 13.4 | 9.1 | 12.5 | 8.9 | 18.4 | 6.4 | | 2.2 | 12.6 |
| | | | | 10.6 | 11.3 | 9.1 | | 11.1 | 10.4 | | 2.0 | 3.5 | 10.7 | 12.7 | 15.8 | 12.9 | 12.7 | 13.0 | | 9.5 | 9.2 |
| | | | | 8.4 | 8.7 | 7.8 | | 11.8 | 11.7 | | 5.7 | 6.6 | 12.2 | 11.6 | 15.7 | 12.3 | 9.9 | 2.8 | | 11.5 | 14.5 |

| | | | | Females | | | | | | | | | Males | | | | | | | | |
|---|---|---|---|---|---|---|---|---|---|---|---|---|---|---|---|---|---|---|---|---|---|
| | | | Date | Age (days) | | | | | | | | | Age (days) | | | | | | | | |
| Genotype | Parameter | Rep | YYMMDD | 2 | 4 | 8 | 15 | 22 | 29 | 30 | 36 | 43 | 2 | 4 | 8 | 15 | 22 | 29 | 30 | 36 | 43 |

Individual locomotion measurements were collected from flies of the indicated genotypes at multiple ages across the lifespan. For each genotype, locomotion parameters were measured in independent replicates, with values reported at the corresponding ages (days). Locomotion parameters include percent time spent immobile (%), velocity (mm/s), and distance traveled (mm). The "Date" column indicates the adult emergence date.

Table S14. Age-specific t-test p-values for locomotion comparisons between *DJ694* and *w* 

| | | | Female | | | Male | | |
|---|---|---|---|---|---|---|---|---|
| | | | P-value vs. $w^{1118}$ | | | P-value vs. $w^{1118}$ | | |
| Age | Rep | *GAL4* | %Immobile | Distance | Velocity | %Immobile | Distance | Velocity |
| 2 | Rep1 | $DJ694^{H}$ | 0.0972 | 0.0893 | 0.1016 | 0.6991 | 0.8512 | 0.6472 |
| | | $DJ694^{L}$ | 0.1250 | 0.2071 | 0.2111 | 0.6446 | 0.9729 | 0.8626 |
| | | $DJ694^{O}$ | 0.0356 | 0.0254 | 0.0131 | 0.8459 | 0.7393 | 0.8018 |
| | | $DJ694^{P}$ | 0.1026 | 0.1537 | 0.0698 | 0.8729 | 0.9370 | 0.9211 |
| | Rep2 | $DJ694^{H}$ | 0.0223 | 0.0963 | 0.1194 | 0.9823 | 0.9915 | 0.7730 |
| | | $DJ694^{L}$ | 0.2292 | 0.1121 | 0.1508 | 0.6223 | 0.4650 | 0.3336 |
| | | $DJ694^{O}$ | 0.0308 | 0.0323 | 0.0314 | 0.4578 | 0.2746 | 0.1841 |
| | | $DJ694^{P}$ | 0.0205 | 0.0155 | 0.0249 | 0.8310 | 0.9277 | 0.6728 |
| 4 | Rep1 | $DJ694^{H}$ | 0.0584 | 0.0392 | 0.1850 | 0.1954 | 0.8134 | 0.8552 |
| | | $DJ694^{L}$ | 0.3548 | 0.0186 | 0.0316 | 0.3049 | 0.7904 | 0.9011 |
| | | $DJ694^{O}$ | 0.2182 | 0.0037 | 0.0064 | 0.4125 | 0.6631 | 0.5519 |
| | | $DJ694^{P}$ | 0.2591 | 0.0662 | 0.0588 | 0.4445 | 0.7836 | 0.6906 |
| | Rep2 | $DJ694^{H}$ | 0.0376 | 0.0194 | 0.0342 | 0.1165 | 0.1931 | 0.2762 |
| | | $DJ694^{L}$ | 0.1690 | 0.0156 | 0.0132 | 0.5727 | 0.8048 | 0.9578 |
| | | $DJ694^{O}$ | 0.4698 | 0.0191 | 0.0185 | 0.3867 | 0.3611 | 0.2306 |
| | | $DJ694^{P}$ | 0.0525 | 0.0059 | 0.0077 | 0.6563 | 0.8772 | 0.8364 |
| 8 | Rep1 | $DJ694^{H}$ | 0.4425 | 0.7446 | 0.7075 | 0.9218 | 0.8996 | 0.5348 |
| | | $DJ694^{L}$ | 0.3322 | 0.4626 | 0.4002 | 0.1631 | 0.1161 | 0.1874 |
| | | $DJ694^{O}$ | 0.5660 | 0.7002 | 0.7200 | 0.2779 | 0.7828 | 0.9661 |
| | | $DJ694^{P}$ | 0.7610 | 0.7352 | 0.7939 | 0.6963 | 0.5752 | 0.6366 |
| | Rep2 | $DJ694^{H}$ | 0.3800 | 0.6014 | 0.5384 | 0.5638 | 0.7011 | 0.7921 |
| | | $DJ694^{L}$ | 0.0673 | 0.0588 | 0.0802 | 0.0141 | 0.0132 | 0.0190 |
| | | $DJ694^{O}$ | 0.1908 | 0.1544 | 0.4496 | 0.0782 | 0.0870 | 0.0568 |
| | | $DJ694^{P}$ | 0.4553 | 0.5148 | 0.6167 | 0.0312 | 0.0319 | 0.0389 |
| | Rep3 | $DJ694^{L}$ | 0.0593 | 0.0928 | 0.2974 | 0.7357 | 0.2401 | 0.3085 |
| | Rep4 | $DJ694^{L}$ | 0.6681 | 0.1798 | 0.0564 | 0.0925 | 0.0109 | 0.0032 |
| | Rep5 | $DJ694^{L}$ | 0.0491 | 0.1930 | 0.8164 | 0.2857 | 0.2896 | 0.3683 |
| 9 | Rep6 | $DJ694^{L}$ | 0.5180 | 0.7297 | 0.9172 | 0.0730 | 0.0799 | 0.0547 |
| 15 | Rep3 | $DJ694^{L}$ | 0.0059 | 0.0676 | 0.1618 | 0.1919 | 0.3162 | 0.4478 |
| | Rep4 | $DJ694^{L}$ | 0.8280 | 0.2835 | 0.4385 | 0.0358 | 0.0019 | 0.0011 |
| | Rep5 | $DJ694^{L}$ | 0.0927 | 0.0919 | 0.0785 | 0.1244 | 0.0246 | 0.0098 |
| | Rep6 | $DJ694^{L}$ | 0.2253 | 0.0795 | 0.0935 | 0.0850 | 0.0373 | 0.0428 |

| | | | Female | | | Male | | |
|---|---|---|---|---|---|---|---|---|
| | | | P-value vs. $w^{1118}$ | | | P-value vs. $w^{1118}$ | | |
| Age | Rep | *GAL4* | %Immobile | Distance | Velocity | %Immobile | Distance | Velocity |
| 22 | Rep1 | $DJ694^{H}$ | 0.7847 | 0.8672 | 0.8662 | 0.0383 | 0.0978 | 0.0880 |
| | | $DJ694^{L}$ | 0.1598 | 0.2513 | 0.3191 | 0.0135 | 0.0271 | 0.0384 |
| | | $DJ694^{O}$ | 0.7002 | 0.1834 | 0.1483 | 0.2471 | 0.4175 | 0.5238 |
| | | $DJ694^{P}$ | 0.8538 | 0.5035 | 0.5719 | 0.0223 | 0.0503 | 0.0614 |
| | Rep2 | $DJ694^{H}$ | 0.5368 | 0.8829 | 0.6969 | 0.5436 | 0.3746 | 0.1089 |
| | | $DJ694^{L}$ | 0.0512 | 0.2117 | 0.3665 | 0.9869 | 0.8004 | 0.8263 |
| | | $DJ694^{O}$ | 0.0052 | 0.0047 | 0.0226 | 0.1538 | 0.1855 | 0.1745 |
| | | $DJ694^{P}$ | 0.0885 | 0.0756 | 0.1045 | 0.1398 | 0.2038 | 0.1951 |
| | Rep3 | $DJ694^{L}$ | 0.9528 | 0.5167 | 0.4638 | 0.1671 | 0.3394 | 0.1245 |
| | Rep4 | $DJ694^{L}$ | 0.1974 | 0.2239 | 0.3094 | 0.5706 | 0.7468 | 0.5293 |
| | Rep5 | $DJ694^{L}$ | 0.7258 | 0.6214 | 0.5107 | 0.0756 | 0.4160 | 0.8060 |
| | Rep6 | $DJ694^{L}$ | 0.8625 | 0.2919 | 0.2029 | 0.1440 | 0.8842 | 0.1856 |
| 29 | Rep3 | $DJ694^{L}$ | 0.2192 | 0.0852 | 0.0298 | 0.0479 | 0.0880 | 0.1785 |
| 30 | Rep4 | $DJ694^{L}$ | 0.0360 | 0.1165 | 0.1601 | 0.2024 | 0.2951 | 0.1521 |
| | Rep5 | $DJ694^{L}$ | 0.9043 | 0.8534 | 0.7729 | 0.2859 | 0.2651 | 0.5112 |
| | Rep6 | $DJ694^{L}$ | 0.1768 | 0.0560 | 0.2393 | 0.2900 | 0.6844 | 0.9231 |
| 36 | Rep1 | $DJ694^{H}$ | 0.1250 | 0.1280 | 0.5692 | 0.0261 | 0.0579 | 0.2749 |
| | | $DJ694^{L}$ | 0.0305 | 0.0359 | 0.7434 | 0.9151 | 0.4644 | 0.4091 |
| | | $DJ694^{O}$ | 0.0718 | 0.0551 | 0.7962 | 0.2739 | 0.4019 | 0.7587 |
| | | $DJ694^{P}$ | 0.0198 | 0.0154 | 0.5302 | 0.7540 | 0.9118 | 0.8393 |
| | Rep2 | $DJ694^{H}$ | 0.0101 | 0.0063 | 0.8307 | 0.0001 | 0.0003 | 0.0124 |
| | | $DJ694^{L}$ | 0.3024 | 0.3326 | 0.3905 | 0.0002 | 0.0029 | 0.1013 |
| | | $DJ694^{O}$ | 0.5176 | 0.3572 | 0.8852 | 0.0005 | 0.0044 | 0.0422 |
| | | $DJ694^{P}$ | 0.1452 | 0.1235 | 0.2802 | 0.0514 | 0.1204 | 0.2233 |
| 43 | Rep1 | $DJ694^{H}$ | 0.0003 | 0.0022 | 0.0300 | 0.9945 | 0.6210 | 0.9721 |
| | | $DJ694^{L}$ | 0.1564 | 0.1350 | 0.5756 | 0.2133 | 0.1686 | 0.5859 |
| | | $DJ694^{O}$ | 0.0227 | 0.0088 | 0.1010 | 0.2554 | 0.4061 | 0.7408 |
| | | $DJ694^{P}$ | 0.1725 | 0.0704 | 0.3894 | 0.4868 | 0.0874 | 0.1806 |
| | Rep2 | $DJ694^{H}$ | 0.1705 | 0.9953 | 0.4665 | 0.3147 | 0.8854 | 0.7284 |
| | | $DJ694^{L}$ | 0.3179 | 0.9415 | 0.3987 | 0.5472 | 0.8110 | 0.3513 |
| | | $DJ694^{O}$ | 0.0366 | 0.0488 | 0.0230 | 0.3822 | 0.7292 | 0.9106 |
| | | $DJ694^{P}$ | 0.4753 | 0.0478 | 0.9392 | 0.6057 | 0.4525 | 0.6869 |

P-values were calculated using two-tailed, two-sample equal varience t-tests comparing each *DJ694* to *w* at the indicated ages and replicates for % immobile, distance traveled, and velocity.

| Female | Male | Exp | YYMMDD | Ovary#1 ($\mu m^2$) | Ovary#2 ($\mu m^2$) | Total ($\mu m^2$) | Mean | SD | P (vs. Wt) |
|---|---|---|---|---|---|---|---|---|---|
| *DJ694 $^H$* | *$w^{1118}$* | 1 | 25-07-09 | 65.36 | 83.82 | 149.18 | 125.74 | 40.82 | 0.8855 |
| | | | 25-07-09 | 68.94 | 80.52 | 149.46 | | | |
| | | | 25-07-09 | 31.85 | 39.23 | 71.08 | | | |
| | | | 25-07-09 | 67.24 | 72.72 | 139.96 | | | |
| | | | 25-07-09 | 48.65 | 47.14 | 95.78 | | | |
| | | | 25-07-09 | 58.70 | 58.16 | 116.86 | | | |
| | | | 25-07-09 | 80.77 | 79.40 | 160.16 | | | |
| | | | 25-07-09 | 34.79 | 31.23 | 66.01 | | | |
| | | | 25-07-09 | 98.27 | 84.90 | 183.16 | | | |
| | | 2 | 25-08-15 | 113.89 | 118.55 | 232.44 | 211.32 | 47.80 | 0.0571 |
| | | | 25-08-15 | 123.81 | 115.43 | 239.23 | | | |
| | | | 25-08-15 | 86.89 | 118.18 | 205.07 | | | |
| | | | 25-08-15 | 106.55 | 113.73 | 220.28 | | | |
| | | | 25-08-15 | 91.61 | 100.53 | 192.14 | | | |
| | | | 25-08-15 | 118.86 | 115.67 | 234.53 | | | |
| | | | 25-08-15 | 123.39 | 125.58 | 248.97 | | | |
| | | | 25-08-15 | 91.64 | 77.92 | 169.56 | | | |
| | | | 25-08-15 | 91.79 | 100.55 | 192.34 | | | |
| | | | 25-08-15 | 47.96 | 56.07 | 104.03 | | | |
| | | | 25-08-15 | 126.43 | 159.48 | 285.91 | | | |
| *DJ694 $^H$* | *DJ694 $^H$* | 3 | 25-09-26 | 86.46 | 72.56 | 159.01 | 159.41 | 55.30 | 0.9761 |
| | | | 25-09-26 | 85.52 | 92.16 | 177.68 | | | |
| | | | 25-09-26 | 87.70 | 84.88 | 172.58 | | | |
| | | | 25-09-26 | 153.58 | 128.23 | 281.81 | | | |
| | | | 25-09-26 | 66.67 | 98.89 | 165.55 | | | |
| | | | 25-09-26 | 93.26 | 107.89 | 201.15 | | | |
| | | | 25-09-26 | 56.36 | 48.92 | 105.29 | | | |
| | | | 25-09-26 | 55.83 | 52.92 | 108.75 | | | |
| | | | 25-09-26 | 62.85 | 55.27 | 118.12 | | | |
| | | | 25-09-26 | 49.49 | 54.66 | 104.16 | | | |
| | | 4 | 25-09-30 | 106.89 | 52.02 | 158.91 | 101.52 | 43.73 | 0.6578 |
| | | | 25-09-30 | 37.72 | 50.05 | 87.76 | | | |
| | | | 25-09-30 | 40.49 | 37.35 | 77.84 | | | |
| | | | 25-09-30 | 87.71 | 66.28 | 153.98 | | | |
| | | | 25-09-30 | 37.44 | 36.31 | 73.74 | | | |
| | | | 25-09-30 | 27.88 | 28.98 | 56.86 | | | |
| | | 5 | 25-10-10 | 101.90 | 90.69 | 192.59 | 186.03 | 22.74 | 8.22E-08 |
| | | | 25-10-10 | 92.12 | 99.25 | 191.37 | | | |
| | | | 25-10-10 | 75.67 | 74.35 | 150.01 | | | |
| | | | 25-10-10 | 95.84 | 90.45 | 186.29 | | | |
| | | | 25-10-10 | 71.27 | 107.36 | 178.64 | | | |
| | | | 25-10-10 | 110.44 | 73.54 | 183.98 | | | |
| | | | 25-10-10 | 105.42 | 116.98 | 222.40 | | | |
| | | | 25-10-10 | 114.02 | 96.76 | 210.78 | | | |
| | | | 25-10-10 | 85.25 | 72.97 | 158.23 | | | |

| Female | Male | Exp | YYMMDD | Ovary#1 (μm$^2$) | Ovary#2 (μm$^2$) | Total (μm$^2$) | Mean | SD | P (vs. Wt) |
|---|---|---|---|---|---|---|---|---|---|
| *DJ694$^H$* | *DJ694$^H$* | 6 | 25-10-13 | 71.37 | 78.73 | 150.10 | 114.42 | 27.87 | 0.1231 |
| | | | 25-10-13 | 73.73 | 58.12 | 131.85 | | | |
| | | | 25-10-13 | 78.26 | 80.37 | 158.63 | | | |
| | | | 25-10-13 | 46.29 | 60.00 | 106.29 | | | |
| | | | 25-10-13 | 28.09 | 29.57 | 57.66 | | | |
| | | | 25-10-13 | 49.12 | 46.97 | 96.09 | | | |
| | | | 25-10-13 | 49.05 | 52.44 | 101.49 | | | |
| | | | 25-10-13 | 49.54 | 49.77 | 99.31 | | | |
| | | | 25-10-13 | 55.51 | 61.64 | 117.16 | | | |
| | | | 25-10-13 | 80.63 | 47.85 | 128.48 | | | |
| | | | 25-10-13 | 53.43 | 58.12 | 111.55 | | | |
| *DJ694$^L$* | *$w^{1118}$* | 1 | 25-07-09 | 36.88 | 41.97 | 78.85 | 147.07 | 56.55 | 0.2688 |
| | | | 25-07-09 | 119.79 | 106.97 | 226.76 | | | |
| | | | 25-07-09 | 47.22 | 50.12 | 97.33 | | | |
| | | | 25-07-09 | 97.85 | 107.67 | 205.53 | | | |
| | | | 25-07-09 | 41.45 | 57.37 | 98.82 | | | |
| | | | 25-07-09 | 59.17 | 74.94 | 134.12 | | | |
| | | | 25-07-09 | 129.28 | 88.71 | 217.99 | | | |
| | | | 25-07-09 | 69.41 | 43.85 | 113.26 | | | |
| | | | 25-07-09 | 79.87 | 71.09 | 150.97 | | | |
| | | 2 | 25-08-15 | 78.19 | 73.92 | 152.11 | 210.61 | 52.57 | 0.1376 |
| | | | 25-08-15 | 151.54 | 158.64 | 310.17 | | | |
| | | | 25-08-15 | 93.18 | 90.75 | 183.94 | | | |
| | | | 25-08-15 | 125.68 | 128.98 | 254.66 | | | |
| | | | 25-08-15 | 151.92 | 100.97 | 252.89 | | | |
| | | | 25-08-15 | 75.60 | 56.34 | 131.95 | | | |
| | | | 25-08-15 | 103.42 | 122.97 | 226.39 | | | |
| | | | 25-08-15 | 95.63 | 111.04 | 206.67 | | | |
| | | | 25-08-15 | 106.86 | 92.59 | 199.45 | | | |
| | | | 25-08-15 | 106.22 | 81.65 | 187.87 | | | |
| *DJ694$^L$* | *DJ694$^L$* | 3 | 25-09-26 | 121.52 | 92.66 | 214.18 | 195.84 | 33.92 | 0.0048 |
| | | | 25-09-26 | 98.22 | 125.22 | 223.44 | | | |
| | | | 25-09-26 | 130.99 | 95.17 | 226.16 | | | |
| | | | 25-09-26 | 103.29 | 105.39 | 208.68 | | | |
| | | | 25-09-26 | 123.22 | 111.83 | 235.05 | | | |
| | | | 25-09-26 | 117.67 | 81.57 | 199.24 | | | |
| | | | 25-09-27 | 71.62 | 73.48 | 145.11 | | | |
| | | | 25-09-27 | 68.35 | 85.23 | 153.59 | | | |
| | | | 25-09-27 | 79.93 | 59.77 | 139.70 | | | |
| | | | 25-09-27 | 102.06 | 109.86 | 211.92 | | | |
| | | | 25-09-27 | 83.71 | 113.42 | 197.13 | | | |
| | | 4 | 25-09-30 | 60.78 | 66.61 | 127.39 | 111.28 | 36.02 | 0.7847 |
| | | | 25-09-30 | 59.64 | 63.78 | 123.42 | | | |
| | | | 25-09-30 | 41.75 | 34.17 | 75.92 | | | |
| | | | 25-09-30 | 47.72 | 66.46 | 114.18 | | | |
| | | | 25-09-30 | 74.17 | 56.81 | 130.98 | | | |
| | | | 25-09-30 | 33.72 | 39.97 | 73.69 | | | |
| | | | 25-09-30 | 51.44 | 56.16 | 107.60 | | | |
| | | | 25-09-30 | 20.14 | 39.86 | 60.00 | | | |
| | | | 25-09-30 | 42.70 | 71.11 | 113.81 | | | |
| | | | 25-09-30 | 104.66 | 81.20 | 185.86 | | | |

| Female | Male | Exp | YYMMDD | Ovary#1 (μm²) | Ovary#2 (μm²) | Total (μm²) | Mean | SD | P (vs. Wt) |
|---|---|---|---|---|---|---|---|---|---|
| *DJ694 L* | *DJ694 L* | 5 | 25-10-10 | 103.43 | 94.50 | 197.93 | 180.33 | 20.66 | 7.39E-08 |
| | | | 25-10-10 | 95.52 | 101.20 | 196.71 | | | |
| | | | 25-10-10 | 118.10 | 101.18 | 219.28 | | | |
| | | | 25-10-10 | 84.89 | 76.48 | 161.37 | | | |
| | | | 25-10-10 | 94.33 | 79.16 | 173.49 | | | |
| | | | 25-10-10 | 74.77 | 103.64 | 178.40 | | | |
| | | | 25-10-10 | 87.60 | 72.66 | 160.26 | | | |
| | | | 25-10-10 | 103.38 | 85.28 | 188.66 | | | |
| | | | 25-10-10 | 83.31 | 92.65 | 175.96 | | | |
| | | | 25-10-10 | 90.22 | 61.03 | 151.25 | | | |
| | | 6 | 25-10-13 | 91.41 | 85.45 | 176.86 | 138.17 | 33.82 | 0.3827 |
| | | | 25-10-13 | 78.39 | 84.44 | 162.83 | | | |
| | | | 25-10-13 | 84.87 | 83.08 | 167.95 | | | |
| | | | 25-10-13 | 76.66 | 73.07 | 149.74 | | | |
| | | | 25-10-13 | 66.42 | 68.01 | 134.43 | | | |
| | | | 25-10-13 | 34.41 | 35.76 | 70.17 | | | |
| | | | 25-10-13 | 61.11 | 63.53 | 124.64 | | | |
| | | | 25-10-13 | 55.92 | 27.85 | 83.77 | | | |
| | | | 25-10-13 | 75.50 | 65.99 | 141.49 | | | |
| | | | 25-10-13 | 86.52 | 64.21 | 150.73 | | | |
| | | | 25-10-13 | 77.80 | 79.52 | 157.32 | | | |
| *DJ694 O* | *w 1118* | 1 | 25-07-09 | 28.30 | 21.43 | 49.73 | 96.19 | 35.77 | 0.0759 |
| | | | 25-07-09 | 58.45 | 57.56 | 116.00 | | | |
| | | | 25-07-09 | 63.09 | 57.22 | 120.31 | | | |
| | | | 25-07-09 | 30.47 | 45.96 | 76.43 | | | |
| | | | 25-07-09 | 75.99 | 69.74 | 145.73 | | | |
| | | | 25-07-09 | 30.28 | 37.56 | 67.84 | | | |
| | | | 25-07-09 | 41.53 | 25.80 | 67.33 | | | |
| | | | 25-07-09 | 69.06 | 66.96 | 136.02 | | | |
| | | | 25-07-09 | 56.21 | 68.71 | 124.92 | | | |
| | | | 25-07-09 | 31.72 | 25.85 | 57.57 | | | |
| | | 2 | 25-08-15 | 42.59 | 79.68 | 122.27 | 182.56 | 55.27 | 0.6274 |
| | | | 25-08-15 | 101.09 | 97.28 | 198.37 | | | |
| | | | 25-08-15 | 86.10 | 124.36 | 210.46 | | | |
| | | | 25-08-15 | 105.61 | 134.87 | 240.47 | | | |
| | | | 25-08-15 | 116.27 | 115.46 | 231.73 | | | |
| | | | 25-08-15 | 114.65 | 86.19 | 200.83 | | | |
| | | | 25-08-15 | 70.93 | 72.74 | 143.67 | | | |
| | | | 25-08-15 | 98.66 | 117.55 | 216.20 | | | |
| | | | 25-08-15 | 98.96 | 97.88 | 196.83 | | | |
| | | | 25-08-15 | 42.49 | 22.27 | 64.77 | | | |
| *DJ694 O* | *DJ694 O* | 3 | 25-09-26 | 70.43 | 108.08 | 178.50 | 186.13 | 55.31 | 0.1486 |
| | | | 25-09-26 | 32.21 | 61.58 | 93.78 | | | |
| | | | 25-09-26 | 64.74 | 58.94 | 123.68 | | | |
| | | | 25-09-26 | 88.42 | 93.13 | 181.56 | | | |
| | | | 25-09-26 | 135.22 | 83.30 | 218.52 | | | |
| | | | 25-09-26 | 140.31 | 144.04 | 284.35 | | | |
| | | | 25-09-26 | 83.32 | 106.96 | 190.28 | | | |
| | | | 25-09-26 | 122.85 | 109.13 | 231.98 | | | |
| | | | 25-09-26 | 130.60 | 81.27 | 211.88 | | | |
| | | | 25-09-26 | 46.38 | 100.39 | 146.77 | | | |

| Female | Male | Exp | YYMMDD | Ovary#1 ($\mu m^2$) | Ovary#2 ($\mu m^2$) | Total ($\mu m^2$) | Mean | SD | P (vs. Wt) |
|---|---|---|---|---|---|---|---|---|---|
| *DJ694* $^{O}$ | *DJ694* $^{O}$ | 4 | 25-09-30 | 89.70 | 97.38 | 187.08 | 180.42 | 26.53 | 2.72E-07 |
| | | | 25-09-30 | 91.67 | 83.42 | 175.09 | | | |
| | | | 25-09-30 | 68.90 | 69.40 | 138.30 | | | |
| | | | 25-09-30 | 85.50 | 112.28 | 197.78 | | | |
| | | | 25-09-30 | 96.63 | 82.85 | 179.48 | | | |
| | | | 25-09-30 | 71.81 | 102.23 | 174.04 | | | |
| | | | 25-09-30 | 82.95 | 109.95 | 192.89 | | | |
| | | | 25-09-30 | 77.75 | 119.07 | 196.83 | | | |
| | | | 25-09-30 | 70.38 | 67.85 | 138.22 | | | |
| | | | 25-09-30 | 109.63 | 114.83 | 224.46 | | | |
| | | 5 | 25-10-10 | 92.73 | 58.36 | 151.09 | 186.05 | 41.87 | 6.79E-06 |
| | | | 25-10-10 | 46.55 | 64.05 | 110.59 | | | |
| | | | 25-10-10 | 65.02 | 120.08 | 185.09 | | | |
| | | | 25-10-10 | 98.58 | 73.87 | 172.45 | | | |
| | | | 25-10-10 | 123.19 | 87.39 | 210.58 | | | |
| | | | 25-10-10 | 116.41 | 138.80 | 255.21 | | | |
| | | | 25-10-11 | 105.11 | 99.33 | 204.44 | | | |
| | | | 25-10-11 | 115.17 | 72.27 | 187.44 | | | |
| | | | 25-10-11 | 80.36 | 73.47 | 153.83 | | | |
| | | | 25-10-11 | 119.12 | 110.66 | 229.78 | | | |
| | | 6 | 25-10-13 | 83.71 | 89.67 | 173.38 | 140.76 | 25.28 | 0.1830 |
| | | | 25-10-13 | 81.00 | 70.72 | 151.72 | | | |
| | | | 25-10-13 | 90.63 | 81.76 | 172.38 | | | |
| | | | 25-10-13 | 64.84 | 66.89 | 131.73 | | | |
| | | | 25-10-13 | 90.83 | 71.28 | 162.11 | | | |
| | | | 25-10-13 | 56.70 | 42.26 | 98.96 | | | |
| | | | 25-10-13 | 54.17 | 58.54 | 112.71 | | | |
| | | | 25-10-13 | 72.90 | 78.42 | 151.32 | | | |
| | | | 25-10-13 | 79.74 | 65.80 | 145.53 | | | |
| | | | 25-10-13 | 86.95 | 52.45 | 139.40 | | | |
| | | | 25-10-13 | 60.00 | 49.14 | 109.13 | | | |
| *DJ694* $^{P}$ | *w* $^{1118}$ | 1 | 25-07-09 | 45.95 | 43.72 | 89.67 | 169.43 | 42.73 | 0.0206 |
| | | | 25-07-09 | 99.86 | 98.87 | 198.73 | | | |
| | | | 25-07-09 | 77.37 | 81.00 | 158.37 | | | |
| | | | 25-07-09 | 95.89 | 92.36 | 188.25 | | | |
| | | | 25-07-09 | 83.85 | 90.83 | 174.68 | | | |
| | | | 25-07-09 | 94.94 | 111.96 | 206.91 | | | |
| | | 2 | 25-08-15 | 93.74 | 105.05 | 198.79 | 199.86 | 29.19 | 0.0739 |
| | | | 25-08-15 | 113.82 | 110.72 | 224.53 | | | |
| | | | 25-08-15 | 76.37 | 83.01 | 159.38 | | | |
| | | | 25-08-15 | 99.78 | 124.21 | 224.00 | | | |
| | | | 25-08-15 | 87.77 | 117.41 | 205.18 | | | |
| | | | 25-08-15 | 83.54 | 80.99 | 164.53 | | | |
| | | | 25-08-15 | 101.60 | 106.39 | 207.99 | | | |
| | | | 25-08-15 | 78.09 | 93.46 | 171.54 | | | |
| | | | 25-08-15 | 126.42 | 116.35 | 242.77 | | | |

| Female | Male | Exp | YYMMDD | Ovary#1 (μm²) | Ovary#2 (μm²) | Total (μm²) | Mean | SD | P (vs. Wt) |
|---|---|---|---|---|---|---|---|---|---|
| *DJ694 P* | *DJ694 P* | 3 | 25-09-26 | 76.94 | 101.51 | 178.45 | 202.48 | 46.17 | 0.0106 |
| | | | 25-09-26 | 127.55 | 118.11 | 245.65 | | | |
| | | | 25-09-26 | 119.39 | 120.45 | 239.84 | | | |
| | | | 25-09-26 | 122.69 | 90.17 | 212.86 | | | |
| | | | 25-09-26 | 121.63 | 83.71 | 205.34 | | | |
| | | | 25-09-26 | 111.24 | 118.91 | 230.15 | | | |
| | | | 25-09-27 | 111.67 | 136.43 | 248.10 | | | |
| | | | 25-09-27 | 97.04 | 115.61 | 212.65 | | | |
| | | | 25-09-27 | 77.39 | 65.09 | 142.48 | | | |
| | | | 25-09-27 | 49.64 | 59.65 | 109.28 | | | |
| | | 4 | 25-09-30 | 49.26 | 55.53 | 104.79 | 177.62 | 48.86 | 0.0002 |
| | | | 25-09-30 | 81.45 | 81.16 | 162.62 | | | |
| | | | 25-09-30 | 108.61 | 102.10 | 210.70 | | | |
| | | | 25-09-30 | 98.05 | 129.97 | 228.01 | | | |
| | | | 25-09-30 | 88.79 | 112.36 | 201.15 | | | |
| | | | 25-09-30 | 136.86 | 104.75 | 241.61 | | | |
| | | | 25-09-30 | 67.31 | 59.80 | 127.12 | | | |
| | | | 25-09-30 | 116.63 | 106.50 | 223.13 | | | |
| | | | 25-09-30 | 67.45 | 72.46 | 139.91 | | | |
| | | | 25-09-30 | 79.58 | 57.62 | 137.20 | | | |
| | | 5 | 25-10-10 | 73.35 | 95.05 | 168.40 | 182.35 | 25.74 | 2.02E-07 |
| | | | 25-10-10 | 69.12 | 76.92 | 146.04 | | | |
| | | | 25-10-10 | 94.25 | 45.08 | 139.34 | | | |
| | | | 25-10-10 | 141.40 | 61.63 | 203.03 | | | |
| | | | 25-10-10 | 140.18 | 53.01 | 193.19 | | | |
| | | | 25-10-11 | 90.30 | 91.64 | 181.94 | | | |
| | | | 25-10-11 | 114.85 | 99.18 | 214.03 | | | |
| | | | 25-10-11 | 125.36 | 84.70 | 210.06 | | | |
| | | | 25-10-11 | 111.93 | 59.97 | 171.91 | | | |
| | | | 25-10-11 | 90.41 | 105.19 | 195.60 | | | |
| | | 6 | 25-10-13 | 93.80 | 94.06 | 187.86 | 175.79 | 23.34 | 7.36E-06 |
| | | | 25-10-13 | 76.48 | 103.86 | 180.34 | | | |
| | | | 25-10-13 | 73.71 | 102.56 | 176.27 | | | |
| | | | 25-10-13 | 111.94 | 84.78 | 196.72 | | | |
| | | | 25-10-13 | 73.46 | 141.39 | 214.85 | | | |
| | | | 25-10-13 | 90.03 | 65.99 | 156.02 | | | |
| | | | 25-10-13 | 67.74 | 59.16 | 126.90 | | | |
| | | | 25-10-13 | 78.64 | 79.93 | 158.56 | | | |
| | | | 25-10-13 | 107.44 | 76.25 | 183.69 | | | |
| | | | 25-10-13 | 120.68 | 64.93 | 185.61 | | | |
| | | | 25-10-13 | 67.68 | 99.15 | 166.83 | | | |
| *w 1118* | *w 1118* | 1 | 25-07-09 | 74.34 | 70.32 | 144.66 | 123.39 | 25.42 | |
| | | | 25-07-09 | 81.19 | 48.91 | 130.11 | | | |
| | | | 25-07-09 | 28.26 | 40.78 | 69.05 | | | |
| | | | 25-07-09 | 64.67 | 72.16 | 136.83 | | | |
| | | | 25-07-09 | 50.51 | 60.35 | 110.86 | | | |
| | | | 25-07-09 | 66.66 | 69.94 | 136.60 | | | |
| | | | 25-07-09 | 65.53 | 47.30 | 112.83 | | | |
| | | | 25-07-09 | 55.00 | 59.39 | 114.39 | | | |
| | | | 25-07-09 | 71.58 | 83.67 | 155.24 | | | |

| Female | Male | Exp | YYMMDD | Ovary#1 ($\mu m^2$) | Ovary#2 ($\mu m^2$) | Total ($\mu m^2$) | Mean | SD | P (vs. Wt) |
|---|---|---|---|---|---|---|---|---|---|
| $w^{1118}$ | $w^{1118}$ | 2 | 25-08-15 | 74.87 | 69.83 | 144.70 | 171.69 | 31.25 | |
| | | | 25-08-15 | 56.87 | 59.04 | 115.91 | | | |
| | | | 25-08-15 | 98.33 | 119.01 | 217.34 | | | |
| | | | 25-08-15 | 110.16 | 80.20 | 190.36 | | | |
| | | | 25-08-15 | 89.66 | 94.85 | 184.51 | | | |
| | | | 25-08-15 | 80.44 | 86.43 | 166.87 | | | |
| | | | 25-08-15 | 80.35 | 83.86 | 164.21 | | | |
| | | | 25-08-15 | 83.61 | 105.99 | 189.60 | | | |
| $w^{1118}$ | $w^{1118}$ | 3 | 25-09-26 | 92.57 | 85.78 | 178.35 | 158.86 | 14.32 | |
| | | | 25-09-26 | 89.59 | 62.66 | 152.25 | | | |
| | | | 25-09-26 | 83.71 | 74.44 | 158.16 | | | |
| | | | 25-09-26 | 87.81 | 73.01 | 160.82 | | | |
| | | | 25-09-26 | 86.51 | 59.52 | 146.03 | | | |
| | | | 25-09-26 | 85.00 | 95.42 | 180.41 | | | |
| | | | 25-09-26 | 60.55 | 77.07 | 137.62 | | | |
| | | | 25-09-26 | 86.36 | 86.93 | 173.29 | | | |
| | | | 25-09-26 | 63.06 | 87.82 | 150.87 | | | |
| | | | 25-09-26 | 62.79 | 88.03 | 150.81 | | | |
| $w^{1118}$ | $w^{1118}$ | 4 | 25-09-30 | 47.56 | 56.81 | 104.37 | 108.00 | 18.23 | |
| | | | 25-09-30 | 51.58 | 71.33 | 122.92 | | | |
| | | | 25-09-30 | 54.52 | 61.62 | 116.14 | | | |
| | | | 25-09-30 | 64.50 | 62.10 | 126.61 | | | |
| | | | 25-09-30 | 41.64 | 51.46 | 93.10 | | | |
| | | | 25-09-30 | 59.78 | 51.59 | 111.37 | | | |
| | | | 25-09-30 | 30.78 | 47.31 | 78.09 | | | |
| | | | 25-09-30 | 47.57 | 59.21 | 106.78 | | | |
| | | | 25-09-30 | 55.63 | 64.23 | 119.85 | | | |
| | | | 25-09-30 | 60.23 | 52.34 | 112.58 | | | |
| | | | 25-09-30 | 67.76 | 63.02 | 130.79 | | | |
| | | | 25-09-30 | 33.09 | 40.33 | 73.42 | | | |
| $w^{1118}$ | $w^{1118}$ | 5 | 25-10-10 | 49.22 | 46.84 | 96.06 | 111.78 | 22.30 | |
| | | | 25-10-10 | 49.07 | 56.68 | 105.75 | | | |
| | | | 25-10-10 | 44.70 | 29.07 | 73.77 | | | |
| | | | 25-10-10 | 38.99 | 46.15 | 85.14 | | | |
| | | | 25-10-10 | 36.50 | 47.35 | 83.85 | | | |
| | | | 25-10-10 | 68.89 | 60.45 | 129.34 | | | |
| | | | 25-10-10 | 68.17 | 45.38 | 113.55 | | | |
| | | | 25-10-11 | 90.60 | 54.88 | 145.49 | | | |
| | | | 25-10-11 | 72.10 | 64.80 | 136.91 | | | |
| | | | 25-10-11 | 51.63 | 64.86 | 116.50 | | | |
| | | | 25-10-11 | 59.11 | 62.89 | 121.99 | | | |
| | | | 25-10-11 | 74.59 | 71.46 | 146.05 | | | |
| | | | 25-10-11 | 71.68 | 48.22 | 119.89 | | | |
| | | | 25-10-11 | 57.20 | 50.04 | 107.24 | | | |
| | | | 25-10-11 | 45.44 | 49.68 | 95.13 | | | |

| Female | Male | Exp | YYMMDD | Ovary#1 ($\mu m^2$) | Ovary#2 ($\mu m^2$) | Total ($\mu m^2$) | Mean | SD | P (vs. Wt) |
|---|---|---|---|---|---|---|---|---|---|
| $w^{1118}$ | $w^{1118}$ | 6 | 25-10-13 | 59.87 | 60.82 | 120.69 | 128.97 | 18.64 | |
| | | | 25-10-13 | 76.61 | 67.02 | 143.63 | | | |
| | | | 25-10-13 | 84.04 | 51.21 | 135.25 | | | |
| | | | 25-10-13 | 68.05 | 97.41 | 165.45 | | | |
| | | | 25-10-13 | 40.84 | 52.98 | 93.82 | | | |
| | | | 25-10-13 | 70.50 | 75.36 | 145.86 | | | |
| | | | 25-10-13 | 66.21 | 60.18 | 126.39 | | | |
| | | | 25-10-13 | 56.06 | 75.22 | 131.28 | | | |
| | | | 25-10-13 | 69.66 | 51.80 | 121.47 | | | |
| | | | 25-10-13 | 61.51 | 80.94 | 142.44 | | | |
| | | | 25-10-13 | 61.73 | 85.61 | 147.33 | | | |
| | | | 25-10-13 | 63.60 | 59.77 | 123.36 | | | |
| | | | 25-10-13 | 54.60 | 48.44 | 103.05 | | | |
| | | | 25-10-13 | 45.93 | 63.37 | 109.29 | | | |
| | | | 25-10-13 | 50.98 | 74.32 | 125.30 | | | |

Column "Female" indicates the genotype of the dissected female, and column "Male" indicates the genotype of the mating male. Column "Exp" denotes the experiment ID for each independent biological replicate, and column "YYMMDD" indicates the dissection date. Columns "Ovary #1 (µm²)" and "Ovary #2 (µm²)" report the measured areas of the two ovaries from a single female; ovary numbering does not correspond to left or right. Column "Total (µm²)" represents the sum of both ovary areas. Columns "Mean" and "SD" indicate the mean and standard deviation of total ovary size within each replicate. Column "P (vs. WT)" reports p-values from two-tailed, two-sample equal varience t-tests comparing total ovary size between homozygous *DJ694* and *w1118* females within each replicate.

Table S16. Stage-specific *EDTP* expression fertility data

| | | | | | Age (days) | | | | | | | | | | | | | | | | | |
|---|---|---|---|---|---|---|---|---|---|---|---|---|---|---|---|---|---|---|---|---|---|---|
| | **Start Date** | | | | **5** | | **10** | | **15** | | **20** | | **25** | | **30** | | **40** | | **50** | | **60** | |
| **Rep** | **YYMMDD** | **Temp** | **Genotype** | **n♀** | **mean** | **sd** | **mean** | **sd** | **mean** | **sd** | **mean** | **sd** | **mean** | **sd** | **mean** | **sd** | **mean** | **sd** | **mean** | **sd** | **mean** | **sd** |
| 1 | 25-04-09 | 18-30 | *DJ694*$^{L}$*;GAL80/UAS-EDTP*$^{E}$ | 15 | 36.1 | 14.6 | 93.3 | 20.1 | 113.9 | 20.0 | 134.1 | 24.0 | 162.5 | 35.2 | 176.7 | 48.1 | - | - | - | - | - | - |
| | | | *DJ694*$^{L}$*;GAL80/+* | 15 | 41.7 | 11.7 | 89.0 | 2.4 | 109.4 | 10.6 | 124.2 | 13.4 | 143.7 | 23.6 | 156.6 | 37.2 | - | - | - | - | - | - |
| | | | *DJ694*$^{L}$*/+* | 15 | 49.1 | 10.3 | 116.2 | 17.4 | 135.4 | 17.2 | 170.5 | 20.7 | 222.9 | 29.7 | 267.0 | 45.0 | - | - | - | - | - | - |
| | 25-04-06 | 30-30 | *DJ694*$^{L}$*;GAL80/UAS-EDTP*$^{E}$ | 15 | 20.8 | 9.0 | 37.3 | 14.5 | 43.3 | 19.2 | 44.2 | 20.3 | - | - | - | - | - | - | - | - | - | - |
| | | | *DJ694*$^{L}$*;GAL80/+* | 15 | 7.3 | 7.0 | 10.5 | 10.7 | 11.3 | 11.4 | 12.2 | 13.0 | - | - | - | - | - | - | - | - | - | - |
| | | | *DJ694*$^{L}$*/+* | 15 | 23.9 | 9.7 | 58.9 | 18.1 | 74.5 | 25.6 | 82.3 | 39.2 | - | - | - | - | - | - | - | - | - | - |
| | 25-04-09 | 18-18 | *DJ694*$^{L}$*;GAL80/UAS-EDTP*$^{E}$ | 15 | - | - | 54.5 | 11.3 | - | - | 109.1 | 9.3 | - | - | 137.6 | 10.7 | 164.3 | 16.7 | 190.7 | 16.7 | 213.6 | 19.2 |
| | | | *DJ694*$^{L}$*;GAL80/+* | 15 | - | - | 48.5 | 19.4 | - | - | 100.7 | 19.5 | - | - | 127.8 | 26.1 | 156.9 | 28.7 | 175.1 | 33.5 | 196.0 | 36.9 |
| | | | *DJ694*$^{L}$*/+* | 15 | - | - | 48.4 | 18.5 | - | - | 118.7 | 17.0 | - | - | 154.8 | 19.7 | 191.0 | 20.6 | 221.2 | 24.7 | 252.2 | 30.2 |
| | 25-04-06 | 30-18 | *DJ694*$^{L}$*;GAL80/UAS-EDTP*$^{E}$ | 15 | - | - | 32.2 | 10.4 | - | - | 68.6 | 17.3 | - | - | 97.5 | 24.1 | 120.2 | 31.8 | - | - | - | - |
| | | | *DJ694*$^{L}$*;GAL80/+* | 15 | - | - | 16.5 | 5.2 | - | - | 42.6 | 8.6 | - | - | 62.4 | 9.2 | 74.2 | 14.7 | - | - | - | - |
| | | | *DJ694*$^{L}$*/+* | 15 | - | - | 37.1 | 6.8 | - | - | 84.6 | 17.3 | - | - | 112.6 | 25.3 | 137.9 | 32.9 | - | - | - | - |
| 2 | 25-04-14 | 18-30 | *DJ694*$^{L}$*;GAL80/UAS-EDTP*$^{E}$ | 15 | 58.9 | 17.0 | 85.0 | 20.1 | 102.9 | 24.9 | 124.3 | 31.5 | 132.8 | 33.3 | - | - | - | - | - | - | - | - |
| | | | *DJ694*$^{L}$*;GAL80/+* | 15 | 54.2 | 8.4 | 82.2 | 6.6 | 110.4 | 4.5 | 131.4 | 5.6 | 139.6 | 9.2 | - | - | - | - | - | - | - | - |
| | | | *DJ694*$^{L}$*/+* | 15 | 57.6 | 18.4 | 98.8 | 25.5 | 131.7 | 26.1 | 158.9 | 28.8 | 182.3 | 25.8 | - | - | - | - | - | - | - | - |
| | 25-04-07 | 30-30 | *DJ694*$^{L}$*;GAL80/UAS-EDTP*$^{E}$ | 15 | 23.7 | 8.9 | 41.4 | 14.7 | 50.9 | 24.9 | 56.4 | 35.4 | - | - | - | - | - | - | - | - | - | - |
| | | | *DJ694*$^{L}$*;GAL80/+* | 15 | 7.3 | 4.4 | 16.9 | 7.2 | 18.1 | 7.7 | 18.3 | 7.6 | - | - | - | - | - | - | - | - | - | - |
| | | | *DJ694*$^{L}$*/+* | 15 | 30.7 | 16.8 | 60.8 | 22.0 | 80.2 | 24.4 | 97.3 | 26.8 | - | - | - | - | - | - | - | - | - | - |
| | 25-04-13 | 18-18 | *DJ694*$^{L}$*;GAL80/UAS-EDTP*$^{E}$ | 15 | - | - | 50.7 | 9.0 | - | - | 83.5 | 11.0 | - | - | 116.2 | 14.6 | 146.3 | 15.8 | 171.7 | 15.0 | 196.9 | 17.0 |
| | | | *DJ694*$^{L}$*;GAL80/+* | 15 | - | - | 49.7 | 6.4 | - | - | 84.6 | 6.6 | - | - | 118.9 | 6.0 | 147.3 | 9.6 | 169.9 | 6.8 | 193.7 | 10.7 |
| | | | *DJ694*$^{L}$*/+* | 15 | - | - | 75.2 | 12.0 | - | - | 112.0 | 16.9 | - | - | 147.9 | 14.0 | 178.0 | 12.0 | 214.3 | 14.9 | 250.1 | 20.6 |
| | 25-04-07 | 30-18 | *DJ694*$^{L}$*;GAL80/UAS-EDTP*$^{E}$ | 15 | - | - | 19.9 | 18.5 | - | - | 48.0 | 29.8 | - | - | 61.8 | 36.6 | 68.6 | 40.2 | 70.6 | 40.6 | - | - |
| | | | *DJ694*$^{L}$*;GAL80/+* | 15 | - | - | 14.9 | 6.9 | - | - | 38.7 | 11.4 | - | - | 53.2 | 11.0 | 61.2 | 13.4 | 64.1 | 16.4 | - | - |
| | | | *DJ694*$^{L}$*/+* | 15 | - | - | 20.2 | 3.2 | - | - | 51.2 | 13.3 | - | - | 77.0 | 25.0 | 99.4 | 34.5 | 119.5 | 46.8 | - | - |

| | Start Date | | | | Age (days) 5 | | 10 | | 15 | | 20 | | 25 | | 30 | | 40 | | 50 | | 60 | |
|---|---|---|---|---|---|---|---|---|---|---|---|---|---|---|---|---|---|---|---|---|---|---|
| Rep | YYMMDD | Temp | Genotype | n♀ | mean | sd | mean | sd | mean | sd | mean | sd | mean | sd | mean | sd | mean | sd | mean | sd | mean | sd |
| 3 | 25-04-24 | 18-30 | *DJ694*$^{L}$*;GAL80/UAS-EDTP*$^{E}$ | 15 | 73.5 | 20.2 | 145.1 | 32.6 | 185.2 | 31.3 | 201.9 | 28.4 | 203.3 | 28.3 | - | - | - | - | - | - | - | - |
| | | | *DJ694*$^{L}$*;GAL80/+* | 15 | 57.7 | 7.9 | 121.4 | 10.9 | 162.4 | 28.0 | 179.7 | 29.8 | 184.8 | 30.8 | - | - | - | - | - | - | - | - |
| | | | *DJ694*$^{L}$*/+* | 15 | 87.8 | 10.9 | 174.8 | 11.1 | 233.5 | 15.5 | 249.8 | 19.1 | 264.2 | 19.5 | - | - | - | - | - | - | - | - |
| | 25-04-20 | 30-30 | *DJ694*$^{L}$*;GAL80/UAS-EDTP*$^{E}$ | 15 | 28.1 | 13.9 | 39.5 | 17.0 | 41.6 | 17.9 | 41.7 | 18.0 | - | - | - | - | - | - | - | - | - | - |
| | | | *DJ694*$^{L}$*;GAL80/+* | 15 | 23.1 | 19.2 | 35.7 | 22.3 | 39.3 | 24.3 | 39.3 | 24.3 | - | - | - | - | - | - | - | - | - | - |
| | | | *DJ694*$^{L}$*/+* | 15 | 49.8 | 20.8 | 79.7 | 31.1 | 106.4 | 34.2 | 128.6 | 43.0 | - | - | - | - | - | - | - | - | - | - |
| | 25-04-24 | 18-18 | *DJ694*$^{L}$*;GAL80/UAS-EDTP*$^{E}$ | 15 | - | - | 58.2 | 3.3 | - | - | 99.0 | 10.6 | - | - | 118.9 | 11.9 | 143.8 | 13.2 | 163.6 | 13.5 | 185.9 | 19.2 |
| | | | *DJ694*$^{L}$*;GAL80/+* | 15 | - | - | 71.4 | 10.2 | - | - | 115.8 | 14.2 | - | - | 146.4 | 10.6 | 178.1 | 12.0 | 203.5 | 10.0 | 230.3 | 6.3 |
| | | | *DJ694*$^{L}$*/+* | 15 | - | - | 81.1 | 10.8 | - | - | 130.4 | 16.8 | - | - | 167.1 | 18.7 | 206.7 | 15.8 | 240.0 | 20.6 | 282.5 | 28.2 |
| | 25-04-19 | 30-18 | *DJ694*$^{L}$*;GAL80/UAS-EDTP*$^{E}$ | 15 | - | - | 25.3 | 11.1 | - | - | 45.2 | 22.6 | - | - | 61.3 | 21.6 | 66.7 | 19.1 | 68.6 | 17.8 | - | - |
| | | | *DJ694*$^{L}$*;GAL80/+* | 15 | - | - | 19.3 | 9.1 | - | - | 44.6 | 9.3 | - | - | 59.5 | 14.9 | 72.7 | 20.4 | 77.0 | 26.1 | - | - |
| | | | *DJ694*$^{L}$*/+* | 15 | - | - | 27.9 | 6.1 | - | - | 58.7 | 17.1 | - | - | 83.8 | 21.8 | 111.5 | 22.0 | 133.3 | 32.2 | - | - |
| 4 | 25-04-28 | 18-30 | *DJ694*$^{L}$*;GAL80/UAS-EDTP*$^{E}$ | 15 | 89.8 | 11.2 | 156.5 | 13.7 | 182.7 | 18.2 | 199.5 | 32.2 | 205.0 | 38.0 | - | - | - | - | - | - | - | - |
| | | | *DJ694*$^{L}$*;GAL80/+* | 15 | 77.8 | 11.6 | 140.2 | 21.0 | 172.0 | 25.2 | 193.0 | 30.4 | 196.2 | 30.0 | - | - | - | - | - | - | - | - |
| | | | *DJ694*$^{L}$*/+* | 15 | 112.9 | 6.4 | 193.6 | 11.9 | 220.5 | 21.0 | 235.4 | 23.4 | 249.3 | 30.8 | - | - | - | - | - | - | - | - |
| | 25-04-20 | 30-30 | *DJ694*$^{L}$*;GAL80/UAS-EDTP*$^{E}$ | 15 | 32.7 | 6.4 | 50.9 | 17.2 | 56.1 | 18.9 | 57.4 | 19.3 | - | - | - | - | - | - | - | - | - | - |
| | | | *DJ694*$^{L}$*;GAL80/+* | 15 | 18.5 | 8.8 | 25.3 | 10.1 | 27.1 | 11.2 | 27.1 | 11.2 | - | - | - | - | - | - | - | - | - | - |
| | | | *DJ694*$^{L}$*/+* | 15 | 73.3 | 22.1 | 110.6 | 36.5 | 148.9 | 36.8 | 175.3 | 40.0 | - | - | - | - | - | - | - | - | - | - |
| | 25-04-28 | 18-18 | *DJ694*$^{L}$*;GAL80/UAS-EDTP*$^{E}$ | 15 | - | - | 77.4 | 10.7 | - | - | 119.2 | 14.0 | - | - | 144.7 | 11.7 | 181.0 | 12.7 | 213.1 | 17.4 | 241.5 | 23.6 |
| | | | *DJ694*$^{L}$*;GAL80/+* | 15 | - | - | 64.3 | 8.0 | - | - | 106.4 | 11.4 | - | - | 132.2 | 10.9 | 166.2 | 15.0 | 195.0 | 13.6 | 221.8 | 13.4 |
| | | | *DJ694*$^{L}$*/+* | 15 | - | - | 89.3 | 6.8 | - | - | 135.3 | 9.4 | - | - | 174.4 | 20.4 | 214.5 | 34.0 | 256.0 | 51.4 | 295.0 | 53.6 |
| | 25-04-19 | 30-18 | *DJ694*$^{L}$*;GAL80/UAS-EDTP*$^{E}$ | 15 | - | - | 30.3 | 15.0 | - | - | 53.8 | 25.1 | - | - | 71.1 | 33.0 | 81.1 | 39.1 | 87.5 | 43.7 | - | - |
| | | | *DJ694*$^{L}$*;GAL80/+* | 15 | - | - | 13.5 | 2.3 | - | - | 26.3 | 5.3 | - | - | 35.5 | 11.5 | 38.7 | 14.3 | 41.7 | 17.4 | - | - |
| | | | *DJ694*$^{L}$*/+* | 15 | - | - | 32.7 | 8.5 | - | - | 56.1 | 10.9 | - | - | 84.0 | 13.0 | 108.9 | 17.8 | 130.2 | 19.2 | - | - |

Column “Rep” denotes the experiment ID for each independent biological replicate, and “Start Date (YYMMDD)” indicates the start date of each fertility assay. Column “Temp” indicates the temperature regime, with the first value representing developmental temperature and the second value representing adult temperature (°C). Column “Genotype” indicates the genotype of the egg-laying females; all females were mated with w males.
Columns “Mean” and “SD” represent the mean and standard deviation of the cumulative number of eggs laid per female at each time point (age in days). Values are averaged across 5 vials per genotype within each replicate, with each vial containing 3 females. Values highlighted in yellow indicate the data points used for statistical comparisons.

Table S17. Statistic (p-value) table for stage-specific *EDTP* expression fertility assay 

| | | | Age (days) | | | | | | | | |
|---|---|---|---|---|---|---|---|---|---|---|---|
| Rep | Temp | Comp | 5 | 10 | 15 | 20 | 25 | 30 | 40 | 50 | 60 |
| 1 | 18-30 | Re vs. Ho | 0.5180 | 0.6503 | 0.6704 | 0.4435 | 0.3502 | 0.4827 | - | - | - |
| | | Ho vs. He | 0.3191 | 0.0086 | 0.0206 | 0.0030 | 0.0016 | 0.0029 | - | - | - |
| | | Re vs. He | 0.1406 | 0.0903 | 0.1050 | 0.0330 | 0.0190 | 0.0155 | - | - | - |
| | 30-30 | Re vs. Ho | 0.0283 | 0.0105 | 0.0126 | 0.0181 | - | - | - | - | - |
| | | Ho vs. He | 0.0144 | 0.0009 | 0.0010 | 0.0052 | - | - | - | - | - |
| | | Re vs. He | 0.6213 | 0.0711 | 0.0616 | 0.0895 | - | - | - | - | - |
| | 18-18 | Re vs. Ho | - | 0.5672 | - | 0.4068 | - | 0.4607 | 0.6342 | 0.3764 | 0.3722 |
| | | Ho vs. He | - | 0.9914 | - | 0.1569 | - | 0.1023 | 0.0628 | 0.0382 | 0.0300 |
| | | Re vs. He | - | 0.5449 | - | 0.2997 | - | 0.1243 | 0.0538 | 0.0518 | 0.0426 |
| | 30-18 | Re vs. Ho | - | 0.0165 | - | 0.0170 | - | 0.0159 | 0.0189 | - | - |
| | | Ho vs. He | - | 0.0006 | - | 0.0013 | - | 0.0031 | 0.0042 | - | - |
| | | Re vs. He | - | 0.3998 | - | 0.1812 | - | 0.3621 | 0.4111 | - | - |
| 2 | 18-30 | Re vs. Ho | 0.5924 | 0.7720 | 0.5276 | 0.6331 | 0.6726 | - | - | - | - |
| | | Ho vs. He | 0.7173 | 0.1963 | 0.1097 | 0.0694 | 0.0081 | - | - | - | - |
| | | Re vs. He | 0.9084 | 0.3707 | 0.1122 | 0.1074 | 0.0301 | - | - | - | - |
| | 30-30 | Re vs. Ho | 0.0060 | 0.0101 | 0.0229 | 0.0462 | - | - | - | - | - |
| | | Ho vs. He | 0.0166 | 0.0028 | 0.0006 | 0.0002 | - | - | - | - | - |
| | | Re vs. He | 0.4326 | 0.1390 | 0.0968 | 0.0731 | - | - | - | - | - |
| | 18-18 | Re vs. Ho | - | 0.8450 | - | 0.8484 | - | 0.7160 | 0.9068 | 0.8129 | 0.7358 |
| | | Ho vs. He | - | 0.0030 | - | 0.0097 | - | 0.0028 | 0.0021 | 0.0003 | 0.0006 |
| | | Re vs. He | - | 0.0064 | - | 0.0134 | - | 0.0081 | 0.0073 | 0.0020 | 0.0021 |
| | 30-18 | Re vs. Ho | - | 0.5914 | - | 0.5299 | - | 0.6299 | 0.7088 | 0.7473 | - |
| | | Ho vs. He | - | 0.1559 | - | 0.1477 | - | 0.0875 | 0.0499 | 0.0372 | - |
| | | Re vs. He | - | 0.9662 | - | 0.8319 | - | 0.4651 | 0.2287 | 0.1160 | - |
| 3 | 18-30 | Re vs. Ho | 0.1423 | 0.1620 | 0.2592 | 0.2630 | 0.3527 | - | - | - | - |
| | | Ho vs. He | 0.0011 | 0.0001 | 0.0011 | 0.0022 | 0.0012 | - | - | - | - |
| | | Re vs. He | 0.2008 | 0.0899 | 0.0149 | 0.0140 | 0.0042 | - | - | - | - |
| | 30-30 | Re vs. Ho | 0.6457 | 0.7712 | 0.8726 | 0.8674 | - | - | - | - | - |
| | | Ho vs. He | 0.0677 | 0.0331 | 0.0072 | 0.0037 | - | - | - | - | - |
| | | Re vs. He | 0.0886 | 0.0348 | 0.0056 | 0.0031 | - | - | - | - | - |
| | 18-18 | Re vs. Ho | - | 0.0247 | - | 0.0663 | - | 0.0049 | 0.0026 | 0.0007 | 0.0012 |
| | | Ho vs. He | - | 0.1806 | - | 0.1756 | - | 0.0639 | 0.0123 | 0.0074 | 0.0037 |
| | | Re vs. He | - | 0.0019 | - | 0.0076 | - | 0.0013 | 0.0001 | 0.0001 | 0.0002 |
| | 30-18 | Re vs. Ho | - | 0.3717 | - | 0.96 | - | 0.8819 | 0.6463 | 0.5683 | - |
| | | Ho vs. He | - | 0.1147 | - | 0.1423 | - | 0.0742 | 0.0201 | 0.0160 | - |
| | | Re vs. He | - | 0.6573 | - | 0.3156 | - | 0.1405 | 0.0089 | 0.0043 | - |

| Rep | Temp | Comp | Age (days) 5 | 10 | 15 | 20 | 25 | 30 | 40 | 50 | 60 |
|---|---|---|---|---|---|---|---|---|---|---|---|
| 4 | 18-30 | Re vs. Ho | 0.1341 | 0.1829 | 0.4613 | 0.7488 | 0.6953 | - | - | - | - |
| | | Ho vs. He | 0.0004 | 0.0011 | 0.0107 | 0.0383 | 0.0246 | - | - | - | - |
| | | Re vs. He | 0.0039 | 0.0018 | 0.0160 | 0.0787 | 0.0776 | - | - | - | - |
| | 30-30 | Re vs. Ho | 0.0199 | 0.0213 | 0.0185 | 0.0161 | - | - | - | - | - |
| | | Ho vs. He | 0.0009 | 0.0010 | 0.0001 | 4E-05 | - | - | - | - | - |
| | | Re vs. He | 0.0042 | 0.0107 | 0.0010 | 0.0004 | - | - | - | - | - |
| | 18-18 | Re vs. Ho | - | 0.0594 | - | 0.1524 | - | 0.1178 | 0.1320 | 0.1051 | 0.1424 |
| | | Ho vs. He | - | 0.0007 | - | 0.0024 | - | 0.0036 | 0.0197 | 0.0334 | 0.0181 |
| | | Re vs. He | - | 0.0694 | - | 0.0652 | - | 0.0226 | 0.0727 | 0.1148 | 0.0755 |
| | 30-18 | Re vs. Ho | - | 0.0383 | - | 0.0437 | - | 0.0520 | 0.0525 | 0.0610 | - |
| | | Ho vs. He | - | 0.0012 | - | 0.0006 | - | 0.0003 | 0.0001 | 0.0001 | - |
| | | Re vs. He | - | 0.7569 | - | 0.8580 | - | 0.4406 | 0.1853 | 0.0801 | - |

Column “Rep” denotes the experiment ID for each independent biological replicate.
Column “Temp” indicates the temperature regime, with the first value representing developmental temperature and the second value representing adult temperature (°C).
Column “Comp” denotes the genotype comparison, where “Re vs. Ho” indicates *DJ694;GAL80/UAS-EDTP* compared to *DJ694;GAL80/+,* “Ho vs. He” indicates *DJ694;GAL80/+* compared to *DJ694/*+, and “Re vs. He” indicates *DJ694;GAL80/UAS-EDTP* compared to *DJ694/+.*
Statistical comparisons were performed at each time point (age in days), comparing the cumulative mean number of eggs laid per female between genotypes. P-values were calculated using two-sample, two-tailed t-tests assuming equal variance. For each genotype within a replicate, measurements were obtained from 5 vials, with 3 females per vial. P-values highlighted in yellow are those used to determine the statistical outcome for each comparison.

**Female fertility of homozygous *DJ694* and $w^{1118}$ at different time intervals**

| | | Homozygous *DJ694* | | $w^{1118}$ | | T-Test | |
|---|---|---|---|---|---|---|---|
| **Rep** | **Time** | **Mean** | **SD** | **Mean** | **SD** | **p-value** | **% Decline** |
| | 1-10d | 79.27 | 5.32 | 98.93 | 8.66 | 0.0285 | -19.88 |
| | 1-20d | 132.75 | 16.97 | 156.87 | 16.92 | 0.1563 | -15.37 |
| | 1-29d | 152.61 | 27.95 | 202.17 | 25.16 | 0.0846 | -24.51 |
| | 1-40d | 164.04 | 33.81 | 255.69 | 46.24 | 0.0503 | -35.84 |
| | 1-50d | 166.68 | 36.90 | 312.49 | 97.31 | 0.0722 | -46.66 |
| | 1-60d | 166.84 | 37.11 | 355.66 | 140.91 | 0.0882 | -53.09 |
| | 1-70d | 166.84 | 37.11 | 399.16 | 198.43 | 0.1170 | -58.20 |
| 1 | 1-80d | 166.84 | 37.11 | 409.83 | 208.89 | 0.1183 | -59.29 |
| | 10-20d | 62.98 | 7.14 | 65.80 | 10.57 | 0.7215 | -4.28 |
| | 20-29d | 21.14 | 12.73 | 50.97 | 9.25 | 0.0304 | -58.51 |
| | 29-40d | 13.57 | 7.76 | 60.93 | 26.14 | 0.0396 | -77.73 |
| | 40-50d | 3.00 | 4.36 | 60.14 | 55.48 | 0.1499 | -95.01 |
| | 50-60d | 0.50 | 0.87 | 47.83 | 49.87 | 0.1756 | -98.95 |
| | 60-70d | 0.00 | 0.00 | 47.33 | 61.51 | 0.2535 | -100.00 |
| | 70-80d | 0.00 | 0.00 | 11.33 | 14.01 | 0.2338 | -100.00 |
| | 1-10d | 86.00 | 9.30 | 96.60 | 8.49 | 0.2186 | -10.97 |
| | 1-20d | 158.57 | 15.98 | 183.47 | 8.93 | 0.0780 | -13.57 |
| | 1-29d | 208.11 | 15.84 | 253.20 | 12.60 | 0.0182 | -17.81 |
| | 1-40d | 233.20 | 13.20 | 306.80 | 14.40 | 0.0028 | -23.99 |
| | 1-50d | 233.20 | 13.20 | 360.87 | 13.36 | 0.0003 | -35.38 |
| | 1-60d | 233.20 | 13.20 | 408.81 | 8.71 | 4.31E-05 | -42.96 |
| | 1-70d | 233.20 | 13.20 | 427.53 | 4.22 | 1.71E-05 | -45.45 |
| 2 | 1-80d | 233.20 | 13.20 | 428.47 | 4.22 | 1.67E-05 | -45.57 |
| | 10-20d | 78.77 | 11.34 | 96.47 | 4.23 | 0.0644 | -18.35 |
| | 20-29d | 52.54 | 2.92 | 79.87 | 5.41 | 0.0015 | -34.21 |
| | 29-40d | 28.38 | 13.19 | 61.07 | 0.90 | 0.0128 | -53.53 |
| | 40-50d | 0.13 | 0.23 | 55.33 | 9.00 | 0.0004 | -99.76 |
| | 50-60d | 0.00 | 0.00 | 56.01 | 4.19 | 2.05E-05 | -100.00 |
| | 60-70d | 0.00 | 0.00 | 23.48 | 8.51 | 0.0088 | -100.00 |
| | 70-80d | 0.00 | 0.00 | 1.31 | 1.25 | 0.1445 | -100.00 |

Cumulative egg output was measured in two independent replicates (Rep 1 and Rep 2). In each replicate, groups of five homozygous *DJ694* or $w^{1118}$ females were housed with five $w^{1118}$ males per vial, with three vials per genotype. The column "Mean" represents the cumulative number of eggs laid per female over each indicated time interval, averaged across three vials, with the corresponding standard deviation shown in the column "SD". Statistical comparisons between *DJ694* and $w^{1118}$ were made using a two-sample, two-tailed, equal-variance t-test. The percentage decline in mean egg output of *DJ694* relative to $w^{1118}$ is shown in the final column (%Decline).

| | **Egg/♀ for *DJ694*** | | | **Egg/♀ for $w^{1118}$** | | |
|---|---|---|---|---|---|---|
| **Age (Days)** | **Vial #1** | **Vial #2** | **Vial #3** | **Vial #1** | **Vial #2** | **Vial #3** |
| 1 | 0.00 | 0.00 | 0.00 | 0.00 | 0.00 | 0.00 |
| 2 | 0.00 | 0.00 | 0.33 | 0.67 | 6.67 | 0.00 |
| 3 | 0.00 | 0.33 | 0.00 | 13.33 | 0.33 | 0.67 |
| 4 | 0.67 | 0.67 | 0.67 | 17.33 | 14.67 | 0.33 |
| 5 | 1.00 | 0.33 | 0.67 | 1.00 | 12.67 | 12.67 |
| 6 | 0.00 | 0.67 | 0.00 | 0.00 | 1.00 | 3.33 |
| 7 | 0.67 | 0.67 | 0.67 | 9.00 | 2.33 | 13.33 |
| 8 | 4.33 | 0.33 | 0.67 | 3.00 | 14.67 | 17.33 |
| 9 | 0.33 | 2.00 | 0.67 | 15.67 | 3.67 | 1.00 |
| 10 | 0.67 | 1.00 | 0.33 | 2.00 | 0.33 | 2.33 |
| 11 | 5.00 | 0.00 | 0.33 | 3.00 | 5.00 | 0.33 |
| 12 | 2.67 | 3.00 | 3.33 | 4.67 | 4.00 | 11.67 |
| 13 | 6.67 | 1.33 | 1.00 | 6.67 | 6.33 | 12.00 |
| 14 | 1.67 | 3.00 | 1.33 | 9.67 | 13.33 | 5.67 |
| 15 | 4.00 | 1.33 | 1.00 | 2.67 | 1.67 | 0.33 |
| 16 | 0.33 | 0.67 | 1.00 | 5.00 | 11.67 | 4.00 |
| 17 | 7.33 | 4.67 | 1.67 | 4.67 | 5.33 | 7.33 |
| 18 | 7.33 | 15.67 | 3.33 | 8.00 | 7.33 | 9.33 |
| 19 | 4.67 | 0.00 | 0.33 | 0.00 | 0.00 | 0.00 |
| 20 | 0.33 | 1.00 | 5.00 | 4.00 | 6.00 | 8.33 |
| 21 | 2.33 | 1.33 | 7.00 | 8.00 | 6.67 | 11.33 |
| 22 | 2.33 | 11.33 | 0.00 | 0.33 | 0.00 | 6.00 |
| 23 | 2.67 | 6.67 | 1.00 | 2.67 | 9.33 | 0.33 |
| 24 | 8.33 | 1.00 | 0.00 | 11.00 | 3.33 | 6.00 |
| 25 | 8.33 | 3.00 | 0.67 | 4.67 | 1.00 | 4.33 |
| 26 | 1.33 | 1.00 | 0.33 | 1.00 | 4.33 | 2.00 |
| 27 | 0.00 | 1.67 | 0.33 | 0.00 | 0.33 | 0.33 |
| 28 | 3.67 | 1.00 | 0.33 | 3.33 | 1.00 | 2.00 |
| 29 | 0.00 | 10.00 | 0.00 | 6.67 | 0.00 | 0.33 |
| 30 | 5.67 | 0.00 | 0.33 | 7.00 | 1.00 | 8.00 |
| 31 | 4.33 | 1.00 | 1.67 | 6.00 | 11.33 | 9.33 |
| 32 | 1.67 | 1.50 | 1.33 | 3.67 | 4.33 | 3.00 |
| 33 | 1.67 | 3.50 | 1.00 | 3.00 | 4.67 | 1.67 |
| 34 | 0.33 | 1.50 | 0.50 | 0.00 | 0.33 | 0.33 |
| 35 | 0.33 | 0.50 | 0.00 | 1.33 | 0.00 | 0.33 |
| 36 | 2.00 | 2.50 | 0.50 | 0.00 | 0.67 | 14.67 |
| 37 | 0.00 | 1.50 | 4.00 | 7.67 | 10.33 | 0.00 |
| 38 | 3.33 | 0.50 | 0.00 | 4.33 | 0.00 | 1.33 |
| 39 | 1.67 | 1.50 | 1.00 | 1.67 | 5.00 | 10.67 |
| 40 | 2.33 | 2.00 | 0.00 | 4.00 | 1.67 | 2.33 |
| 41 | 0.00 | 0.50 | 1.00 | 4.67 | 1.33 | 4.33 |
| 42 | 0.00 | 0.00 | 0.50 | 1.00 | 0.67 | 3.67 |
| 43 | 9.00 | 0.50 | 1.50 | 1.00 | 0.33 | 0.00 |

| | Egg/♀ for *DJ694* | | | Egg/♀ for $w^{1118}$ | | |
|---|---|---|---|---|---|---|
| **Age (Days)** | **Vial #1** | **Vial #2** | **Vial #3** | **Vial #1** | **Vial #2** | **Vial #3** |
| 44 | 0.00 | 0.00 | 0.50 | 2.33 | 0.67 | 0.33 |
| 45 | 0.67 | 0.50 | 4.50 | 1.33 | 7.00 | 2.33 |
| 46 | 2.00 | 0.00 | 2.50 | 1.33 | 11.67 | 2.67 |
| 47 | 3.33 | 0.00 | 0.00 | 0.33 | 0.00 | 4.33 |
| 48 | 0.00 | 0.00 | 0.00 | 0.00 | 1.00 | 1.00 |
| 49 | 0.67 | 3.00 | 0.50 | 1.00 | 6.67 | 4.33 |
| 50 | 3.33 | 0.00 | 1.00 | 0.67 | 2.67 | 0.67 |
| 51 | 0.33 | 2.00 | 1.00 | 0.00 | 2.00 | 6.33 |
| 52 | 3.33 | 2.00 | 2.50 | 0.67 | 2.00 | 1.00 |
| 53 | 2.00 | 2.00 | 0.50 | 0.33 | 2.00 | 2.00 |
| 54 | 1.00 | 11.00 | 0.50 | 1.00 | 1.33 | 2.00 |
| 55 | 1.00 | 7.00 | 1.00 | 2.00 | 13.33 | 7.33 |
| 56 | 1.00 | 9.00 | 5.00 | 0.00 | 1.67 | 1.67 |
| 57 | 0.67 | 1.00 | 1.00 | 0.67 | 0.00 | 0.00 |
| 58 | 1.33 | 1.00 | 0.00 | 5.33 | 4.00 | 4.00 |
| 59 | 0.00 | 0.00 | 2.00 | 1.00 | 1.33 | 0.00 |
| 60 | 0.33 | 0.00 | 0.50 | 0.00 | 8.33 | 0.00 |
| 61 | 0.00 | 7.00 | 0.00 | 1.00 | 0.67 | 1.67 |
| 62 | 1.00 | 4.00 | 1.00 | 6.67 | 7.00 | 6.67 |
| 63 | 1.00 | 2.00 | 0.50 | 1.33 | 6.67 | 1.67 |
| 64 | 0.00 | 0.00 | 0.00 | 0.00 | 0.00 | 0.00 |
| 65 | 0.67 | 1.00 | 0.00 | 0.33 | 0.67 | 0.33 |
| 66 | 0.33 | 0.00 | 0.50 | 0.33 | 0.00 | 0.00 |
| 67 | 0.00 | 4.00 | 0.50 | 1.00 | 4.33 | 3.67 |
| 68 | 0.67 | 2.00 | 0.50 | 0.00 | 0.67 | 1.33 |
| 69 | 1.00 | 4.00 | 0.50 | 0.00 | 0.00 | 0.00 |
| 70 | 0.00 | 0.00 | 1.50 | 3.33 | 5.00 | 2.67 |
| 71 | 0.33 | 2.00 | 1.50 | 1.33 | 1.33 | 0.00 |
| 72 | 0.00 | 2.00 | 1.00 | 0.00 | 0.00 | 0.00 |
| 73 | 0.00 | 0.00 | 0.00 | 3.33 | 3.67 | 3.00 |
| 74 | 0.33 | 5.00 | 0.00 | 1.67 | 1.33 | 1.67 |
| 75 | 0.00 | 2.00 | 1.00 | 0.00 | 0.00 | 2.33 |
| 76 | 0.67 | 1.00 | 0.00 | 2.67 | 5.33 | 0.67 |
| 77 | 0.33 | 4.00 | 0.50 | 3.00 | 1.33 | 1.00 |
| 78 | 1.00 | 3.00 | 0.50 | 0.67 | 1.33 | 0.67 |
| 79 | 0.00 | 1.00 | 0.00 | 3.00 | 1.67 | 3.00 |
| 80 | 0.33 | 1.00 | 0.00 | 0.67 | 0.67 | 2.33 |
| 81 | 0.33 | 2.00 | 0.00 | 1.33 | 0.33 | 1.33 |
| 82 | 0.00 | 0.00 | 0.00 | 0.67 | 0.67 | 0.33 |
| 83 | 0.33 | 0.00 | 0.00 | 0.00 | 5.00 | 1.67 |
| 84 | 0.33 | 0.00 | 0.00 | 1.00 | 1.67 | 1.00 |
| 85 | 0.33 | 0.00 | 0.00 | 2.00 | 2.33 | 0.67 |
| 86 | 0.33 | 0.00 | 0.00 | 0.00 | 0.00 | 0.67 |

| | **Egg/♀ for *DJ694*** | | | **Egg/♀ for *w*[1118]** | | |
|---|---|---|---|---|---|---|
| **Age (Days)** | **Vial #1** | **Vial #2** | **Vial #3** | **Vial #1** | **Vial #2** | **Vial #3** |
| 87 | 0.67 | 0.00 | 0.00 | 0.33 | 1.67 | 1.00 |
| 88 | 0.00 | 0.00 | 0.00 | 0.33 | 1.67 | 1.00 |
| 89 | 0.33 | 1.00 | 0.00 | 2.67 | 3.33 | 3.00 |
| 90 | 0.67 | 1.00 | 0.00 | 5.00 | 2.00 | 3.00 |
| 91 | 0.00 | 1.00 | 0.00 | 2.00 | 0.67 | 2.00 |

Fertility assay started at 2024-Aug-8th.

Columns indicate age in days (Age, Days) and the number of eggs laid per female (Egg/♀) for each genotype across individual vials (Vial #1–3). Each vial initially contained three females (all mated with equal amount of wt males); some females may have died over the course of the assay.

Table S20. *DJ694* and *W1118*'s Daily Fertility Data at 29˚ C

| | Egg/♀ for *DJ694* | | Egg/♀ for $w^{1118}$ | | |
|---|---|---|---|---|---|
| **Age (Days)** | **Vial #1** | **Vial #2** | **Vial #1** | **Vial #2** | **Vial #3** |
| 1 | 0.00 | 0.00 | 5.67 | 0.00 | 0.00 |
| 2 | 1.67 | 1.00 | 7.33 | 8.67 | 4.33 |
| 3 | 1.33 | 0.67 | 1.33 | 2.67 | 0.67 |
| 4 | 5.67 | 2.67 | 12.67 | 17.33 | 12.00 |
| 5 | 15.33 | 3.00 | 12.67 | 17.67 | 11.33 |
| 6 | 5.33 | 0.67 | 1.67 | 1.00 | 1.00 |
| 7 | 2.00 | 2.33 | 9.33 | 7.33 | 3.67 |
| 8 | 2.67 | 1.00 | 3.00 | 1.67 | 10.33 |
| 9 | 7.33 | 9.00 | 1.00 | 6.00 | 0.33 |
| 10 | 3.67 | 0.33 | 8.67 | 14.33 | 10.67 |
| 11 | 5.67 | 1.33 | 2.33 | 8.33 | 5.33 |
| 12 | 4.33 | 1.33 | 1.67 | 6.33 | 2.00 |
| 13 | 0.67 | 0.00 | 2.33 | 1.00 | 1.33 |
| 14 | 0.00 | 0.33 | 2.00 | 3.00 | 0.00 |
| 15 | 0.00 | 0.33 | 6.00 | 2.67 | 3.00 |
| 16 | 0.00 | 0.00 | 3.00 | 2.33 | 7.00 |
| 17 | 0.00 | 0.33 | 8.33 | 4.67 | 4.33 |
| 18 | 0.00 | 0.00 | 1.00 | 0.00 | 0.33 |
| 19 | 0.00 | 0.00 | 1.00 | 0.00 | 0.00 |
| 20 | 0.00 | 0.00 | 1.00 | 0.00 | 0.00 |
| 21 | 0.00 | 0.00 | 0.00 | 1.33 | 0.00 |
| 22 | 0.00 | 0.00 | 0.00 | 1.67 | 0.33 |
| 23 | 0.00 | 0.00 | 0.00 | 1.67 | 0.00 |

Fertility assay started at 2024-Sep-18th.

Columns indicate age in days (Age, Days) and the number of eggs laid per female (Egg/♀) for each genotype across individual vials (Vial #1–3). Each vial initially contained three females (all mated with equal amount of wt males); some females may have died over the course of the assay.

Table S21. Day 5 female fertility comparison between homozygous *DJ694* and *w1118* controls. 

| Rep | 694 Strains | Homozygous *DJ694* | | $w^{1118}$ | | T-test at day 5 | |
|---|---|---|---|---|---|---|---|
| | | Mean | SD | Mean | SD | p value | Δ |
| 1 | *DJ694*$^{H}$ | 48.38 | 14.43 | 85.13 | 17.98 | 0.0005 | ↓ |
| | *DJ694*$^{L}$ | 50.08 | 16.84 | | | 0.0013 | ↓ |
| | *DJ694*$^{O}$ | 21.04 | 16.11 | | | 2.85E-06 | ↓ |
| | *DJ694*$^{P}$ | 19.25 | 8.85 | | | 2.29E-07 | ↓ |
| 2 | *DJ694*$^{H}$ | 36.08 | 14.57 | 47.79 | 8.30 | 0.0683 | ↓ |
| | *DJ694*$^{L}$ | 21.88 | 11.45 | | | 0.0001 | ↓ |
| | *DJ694*$^{O}$ | 4.25 | 2.56 | | | 1.54E-06 | ↓ |
| | *DJ694*$^{P}$ | 4.42 | 1.34 | | | 1.40E-06 | ↓ |

Columns indicate the experimental replicate (Rep), Homozygous *DJ694* strains, and summary statistics for homozygous *DJ694* and *w1118* females (mean and standard deviation). Fertility is assessed as the cumulative number of eggs laid per female at day 5. Mean values represent the average cumulative egg counts per female within each replicate. The “Δ” column indicates the difference in mean fertility, a "↓ " means the mean of the *DJ694* is lower than the mean of the *w1118*. The “T-test at day 5” column shows p-values from two-tailed, two-sample t-tests assuming equal variance, comparing homozygous *DJ694* to *w1118* controls within each replicate.
